# Cold adaptation drives population genomic divergence in the ecological specialist, *Drosophila montana*

**DOI:** 10.1101/2020.04.20.050450

**Authors:** R. A. W. Wiberg, V. Tyukmaeva, A. Hoikkala, M. G. Ritchie, M. Kankare

**Affiliations:** Centre for Biological Diversity, School of Biology, University of St Andrews, St Andrews, KY16 9TH, Scotland, UK; Department of Biological and Environmental Science, University of Jyväskylä, Jyväskylä, 40014, Finland

**Keywords:** Environmental adaptation, genomic divergence, cold tolerance, CTmin, CCRT, cline populations, *D. montana*

## Abstract

Detecting signatures of ecological adaptation in comparative genomics is challenging, but analysing population samples with characterised geographic distributions, such as clinal variation, can help identify genes showing covariation with important ecological variation. Here we analysed patterns of geographic variation in the cold-adapted species *Drosophila montana* across phenotypes, genotypes and environmental conditions and tested for signatures of cold adaptation in population genomic divergence. We first derived the climatic variables associated with the geographic distribution of 24 populations across two continents to trace the scale of environmental variation experienced by the species, and measured variation in the cold tolerance of the flies of six populations from different geographic contexts. We then performed pooled whole genome sequencing of these six populations, and used Bayesian methods to identify SNPs where genetic differentiation is associated with both climatic variables and the population phenotypic measurements, while controlling for effects of demography and population structure. The top candidate SNPs were enriched on the X and 4^th^ chromosomes, and they also lay near genes implicated in other studies of cold tolerance and population divergence in this species and its close relatives. We conclude that ecological adaptation has contributed to the divergence of *D. montana* populations throughout the genome and in particular on the X and 4^th^ chromosomes, which also showed highest interpopulation F_ST_. This study demonstrates that ecological selection can drive genomic divergence at different scales, from candidate genes to chromosome-wide effects.

## INTRODUCTION

The geographic structure of a species is a result of its phylogeographic history, influenced by past and present dispersal, population demography, and selection. Obtaining genome-wide data on genetic polymorphisms across multiple populations of a species is becoming relatively easy, but interpreting the patterns of geographic variation in such data and identifying genes which vary primarily due to selection remains challenging. Often simple ‘outlier’ approaches using genome scans which measure genetic differentiation such as F_ST_ or Dxy are adopted, but results are difficult to interpret due to confounding effects of selection, drift and population structure, or genomic features such as inversions and other causes of variation in recombination rate (Noor & Bennet, 2009; Cruickshank & Hahn, 2014; Wolf & Ellegren, 2016; Ravinet et al., 2017). If environmental data are available, we can use associations with such factors to help identify loci where genetic differentiation covaries with this environmental variation. Some genome scan methods can incorporate environmental variation and simultaneously fit effects for covariance with environmental factors, while controlling for effects of population demography (Foll & Gaggiotti 2008; de Villemereuil & Gaggiotti 2015). This approach has successfully identified genetic variation associated with altitude in humans, among other examples (Foll, Gaggiotti, Daub, Vatsiou, & Excoffier, 2014; de Villemereuil & Gaggiotti, 2015; Gautier, 2015) and has become a useful approach to investigate the ecological adaptations underlying population divergence.

Clinal patterns of variation in phenotypes or gene frequencies have a long history of being used to infer selection along ecotones, and analyses of cline shape can sometimes identify loci under direct selection from others showing clinal variation for other reasons, such as phylogeographic history (Barton & Gale, 1993). Such studies can be very powerful, especially when independent parallel clines are available. For example, Kolaczkowski, Kern, Holloway & Begun (2011) sampled isofemale lines from extremes of a cline in Australian populations of *D. melanogaster*, and found many genes implicated in clinally varying phenotypes to show highest differentiation. Also, Bergland, Behrman, O’Brien, Schmidt & Petrov (2014), and Kapun, Fabian, Goudet & Flatt (2016) sampled North American clines in *D. melanogaster* and *D. simulans* over several years to uncover clinal variation at a genome scale. Bergland et al., (2014) also found consistent fluctuations in allele frequencies for a population sampled over several seasons, indicating a regular response to seasonally varying selection pressures. On the other hand, Machado et al. (2015) concluded that migration and gene flow play a greater role than adaptation in the overall clinality of genomic variants in *D. simulans* than *D. melanogaster*. While the two species share a significant proportion of the genes showing clinal variation, their differences in overwintering ability, migration and population bottlenecks probably act as additional drivers of differences in patterns of variation between them (Machado et al., 2015). Similar studies of clinal variation in phenotypes and allele frequencies have also been carried out in other insects (e.g. Paolucci, Salis, Vermeulen, Beukeboom, & van de Zande, 2016), plants (e.g. Chen et al., 2012; Bradbury, Smithson, & Krauss, 2013), mammals (e.g. Hoekstra, Drumm, & Nachman, 2004; Carneiro et al., 2013), fish (e.g. Vines et al., 2016), and other organisms (Endler, 1973; Endler, 1977; Takahashi, 2015). However, more studies are needed that simultaneously compare patterns of variation in allele frequencies and potentially causal environmental variation or ecologically important traits while controlling for population structure, as such studies are necessary to determine if adaptation to climate is directly driving patterns of genetic differentiation.

Here we investigated geographic variation at both the phenotypic and genetic levels in *Drosophila montana* samples from two continents. This species has spread around the northern hemisphere (Throckmorton, 1969) and both mtDNA and microsatellite data have revealed genetically distinct Finnish and North American populations (Mirol et al., 2007). Moreover, more recent modelling of genome-wide SNP frequencies suggest that the Finnish-North American split happened around 1.75MYA and that of the North American populations shortly after that (Garlovsky et al., 2020). It is one of the most cold-tolerant *Drosophila* species (Kellermann et al., 2012;) and the basic cold tolerance of *D. montana* flies can increase towards the cold seasons through two mechanisms, photoperiodic reproductive diapause (Vesala & Hoikkala, 2011) and cold-acclimation induced by a decrease in day length and/or temperature (Vesala, Salminen, Laiho, Hoikkala, & Kankare, 2012; Kauranen et al., 2019). *D. montana* populations have been found to show clinal variation in the critical day length required for diapause induction (CDL; Tyukmaeva, Salminen, Kankare, Knott, & Hoikkala 2011; Lankinen, Tyukmaeva, & Hoikkala 2013). There is also a correlation between CDL and latitudinally co-varying climatic factors such as the mean temperature of the coldest month (Tyukmaeva, Lankinen, Kinnunen, Kauranen, & Hoikkala, 2020). In addition, *D. montana* populations from different geographic regions show variation in their courtship cues and mate choice (Routtu et al., 2007; Klappert, Mazzi, Hoikkala, & Ritchie, 2007), which has led to partial reproductive isolation between some distant populations (Jennings, Mazzi, Ritchie, & Hoikkala, 2011; Jennings, Snook, & Hoikkala, 2014). At the genetic level, differential gene expression studies have identified candidate genes underlying diapause (Kankare, Salminen, Laiho, Vesala, & Hoikkala, 2010; Kankare, Parker, Merisalo, Salminen, & Hoikkala, 2016), perception of day length (Parker, Ritchie, & Kankare, 2016), and cold acclimation (Parker et al., 2015). Furthermore, a quasi-natural selection experiment for shorter CDL, accompanied by a decrease in cold-tolerance, induced widespread changes in loci with potential roles with these traits (Kauranen et al., 2019). Finally, population genomic analyses have identified several outlier loci when examining differentiation between North American and European populations (Parker et al., 2018). All this makes *D. montana* an interesting example of nascent speciation, potentially influenced by ecological adaptation.

Here we sought to ask to what extent patterns in the genomic divergence of *D. montana* populations across two continents are correlated with climatic variation and phenotypic responses to cold adaptation. We performed pooled whole-genome sequencing (pool-seq) on six different populations and used Bayesian methods to examine the association between genomic differentiation between populations and environmental variables across both continents. We also phenotyped populations for two different cold tolerance measures; critical thermal minimum (CTmin), and chill coma recovery time (CCRT), and investigated the associations between them and the genetic and climatic data. Ultimately, we asked if the genomic loci showing an association between genetic and environmental differentiation also showed association with population differentiation in cold tolerance phenotypes, and examined the possible overlap between the set of genes close to candidate SNPs with sets of candidate genes from previous studies of cold adaptation in *D. montana*. If population differentiation is driven by ecological selection then we would predict the extreme cold adaptation of *D montana* to have left a signature of genomic divergence associated with environmental and phenotypic differentiation across these loci.

## MATERIALS AND METHODS

### Sample collections and DNA extraction

We collected samples of 49-50 wild-caught flies from six *D. montana* populations from a range of latitudes from 66°N to 38°N in the spring of 2013 or 2014. Four of these populations represented a range of latitudes in North America (N. A.), and two populations were from a range in Finland (figure 1; table 1). Samples of wild-caught flies from the six populations were stored in ethanol (the male/female ratio varied across samples; table 1) and DNA of individual flies was extracted using CTAB solution and phenol-chloroform-isoamylalcohol purifications in 2016. Genomic DNA was extracted from individual flies and quantified using Qubit (Thermo Fisher Scientific), and an equal amount of DNA from each individual (50 ng) was pooled into the final sample. Sequencing was performed at the Finnish Functional Genomics Centre in Turku, Finland (www.btk.fi/functional-genomics) on the Illumina HiSeq3000 platform (paired-end reads, read length = 150bp, estimated coverage ~121x).

**Table 1.** The sources of genomic samples (coordinates and the name of the nearest town), altitude of the sampling site, the year in which sampling was performed, and the number of males and females sampled (M/F) for each pool.

| Source | Sampling Site | Year | M/F |
| --- | --- | --- | --- |
| USA,<br>Alaska | Seward<br>60°9'N; 149°27'W<br>Altitude 35 m | 2013 | 30/20 |
| Canada,<br>British Columbia | Terrace<br>54°27'N; 128°34'W<br>Altitude 217 m | 2014 | 22/27 |
| USA,<br>Washington | Ashford,<br>46°45'N; 121°57'W<br>Altitude 573 m | 2013 | 16/34 |
| USA,<br>Colorado | Crested Butte<br>38°54'N; 106°57'W<br>Altitude 2900 m | 2013 | 36/13 |
| Finland | Oulanka<br>66°40'N; 29°20'E<br>Altitude 337 m | 2013 | 25/25 |
| Finland | Korpilahti<br>62°20'N; 25°34'E<br>Altitude 133 m | 2013 | 27/23 |

**Figure 1.**
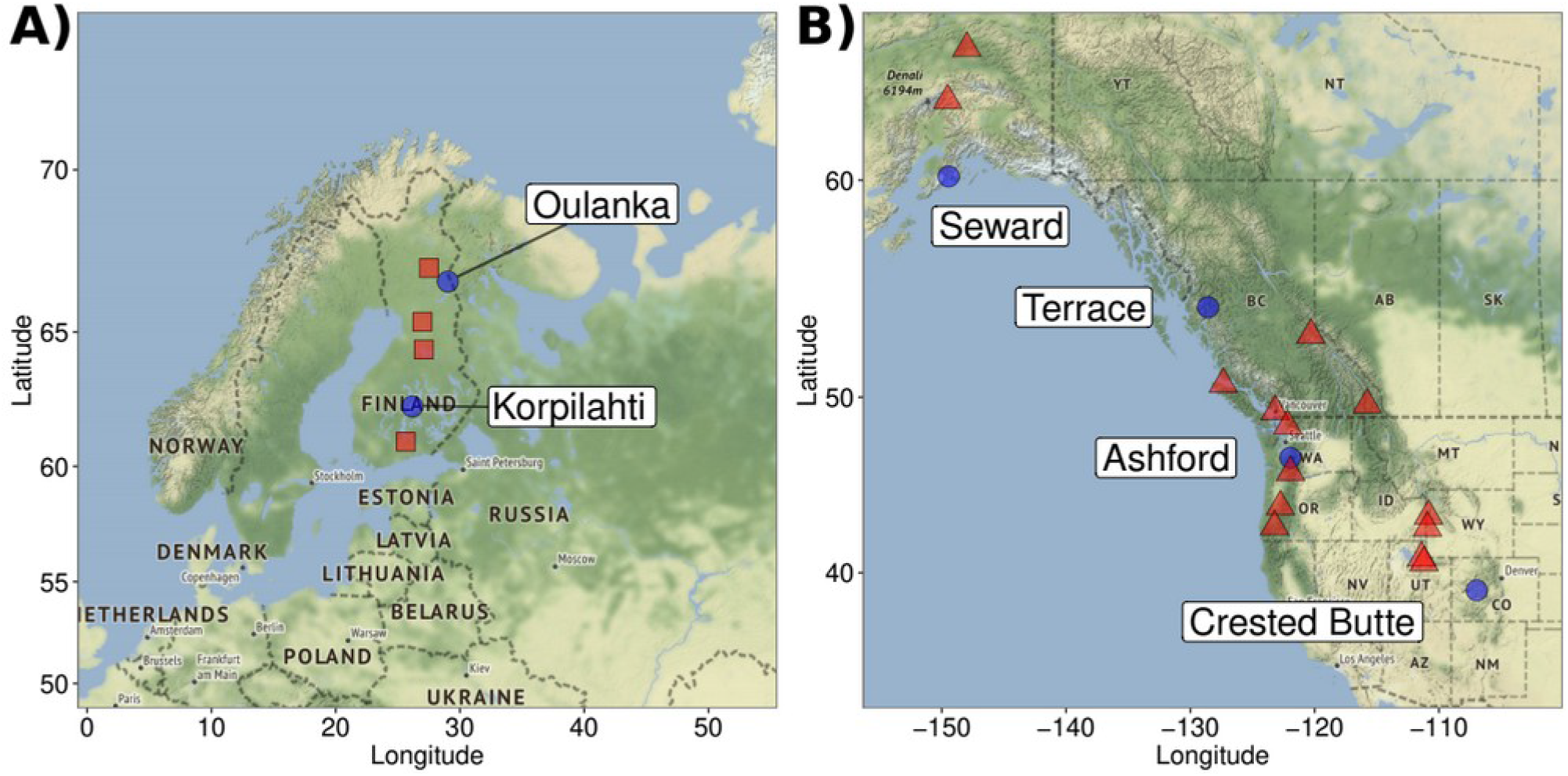
Maps of all *D. montana* populations in this study. Panels show population locations from **A)** Finland and **B)** North America showing the locations of all populations sampled. Labelled, blue circles give the locations of populations sampled for phenotyping and sequencing.

### Phenotyping

We measured the critical thermal minimum (CTmin) and chill coma recovery time (CCRT) of flies from six populations. Fly samples for these tests were collected for five populations (Seward, Terrace, Ashford, Crested Butte, and Korpilahti) from mass-bred populations that have been maintained in the lab since 2013-2014. For the Oulanka population, flies were collected from three isofemale strains (established in 2014), because the mass-bred population had been contaminated by another species. The mass-bred populations were originally established from F4 progenies of 20 isofemale strains each (400 flies) and have been maintained in constant temperature (19 ± 1°C) and light (24h of light, to prevent the flies from entering diapause) regimes for about 20-25 generations before the experiment. All flies were supplied with fresh malt medium in half-pint bottles every week (Lakovaara, 1969). The newly emerged flies were collected using light CO_2_ anaesthesia within 24 hours after emergence, separated by sex and placed in malt-vials in the same conditions until sexual maturity (20-21 days old) and were then used in CTmin and CCRT tests. The same individual flies were first assayed for CTmin and then for CCRT, the flies were not anaesthetised before these tests.

We assayed a total of 328 females and 302 males for CTmin and CCRT. These assays were done in batches of between 22 and 30 flies, and split evenly by sex (21 batches in total). Between 32 and 46 (mean 39) flies per population per sex were tested (for the Oulanka isofemale strains, between 15 and 39 flies per strain per sex were used). CTmin tests are based on detecting the temperature (CTmin) at which flies lose neuromuscular function and enter reversible state of chill coma (Andersen et al., 2015). In these tests, the flies were placed into tubes sealed with parafilm and submerged into a 30% glycol-water mixture in Julabo F32-HL chamber. The temperature was decreased at the rate of 0.5°C per minute (from 19°C to −6°C) and CTmin was determined by eye, as the temperature at which a fly was unable to stand on its legs. Immediately after the Ctmin test, the temperature was set to −6°C and the flies were left in this temperature for 16 hours. Then vials were quickly taken out of the glycol-water bath and the flies’ CCRT was determined as the time required for the flies to recover from chill coma and stand on their legs. The ambient room temperatures was recorded during trials but in an initial analysis including source population and room temperature there was no statistically significant effect of room temperature on CCRT (F_5,569_ = 0.72, p = 0.4), this variable was therefore left out of further analyses.

To investigate population differences in CTmin and CCRT phenotypes we fit General Additive Mixed Models (GAMMs) in R (v. 3.6.3; R Development Core Team, 2020) using the “mgcv” R package (Wood, 2004). In simple linear models including population, sex, and experimental batch as a fixed effects, experimental batch had an effect on CTmin (F_20,533_ = 2.8, p < 0.01), although compared to population (F_5,533_=12, p < 0.01), and sex (F_1,533_=11.5, p < 0.01) this effect was not strong. Experimental batch had a stronger effect on CCRT (F_20,533_ = 2.83, p < 0.01), compared to population (F_5,533_=4.5, p < 0.01) and sex (F_1,533_=1.5, p < 0.22). We therefore included it as a random effect in GAMM analyses. The full GAMMs included altitude, latitude, and sex as fixed effects and experimental batch as a random effect. We used a cubic regression spline as the basis smoothing function for both altitude and latitude. The raw data for all phenotyping are given in table S1. The full models are shown in the Supplementary Materials.

### Bioclimatic variables and population geography

We obtained representative climate data from the WorldClim database (Hijmans, Cameron, Parra, Jones, & Jarvis, 2005) for each *D. montana* population sampled for the pool-seq (see above), as well as for 18 additional populations of this species (table S1; Tyukmaeva et al., 2020). We downloaded the climate data and extracted the values corresponding to population coordinates using the R package “raster” (v. 2.5-8; Hijmans et al., 2016). In total this amounts to 55 bioclimatic variables for each population (table S1). To reduce the number of variables in the dataset a principle components analysis (PCA) was performed using the “PCA()” function from the R package “FactoMineR” (v. 1.28; Lê, Josse, & Husson, 2008). Principle components were kept for further analysis if their eigenvalues were > 1. PCA scores for each population were z-transformed using the “scale()” function in base R. Additionally, CTmin and CCRT were summarised to a mean value for each population. In total, this gives four “environmental” variables measured for each population (PC1, PC2, CTmin, and CCRT).

### Mapping, SNP calling and genomic analysis

Quality of raw reads was checked with FASTQC (v. 0.11.5; Andrews, 2015) and reads were trimmed using trimmomatic (v. 0.32; Bolger, Lohse, & Usadel, 2014; see Supplementary Materials for full trimming parameters). Trimmed reads were mapped to the *D. montana* reference genome (Parker et al., 2018) using BWA mem (v. 0.7.7; Li, 2013) with the default options but keeping only alignments with a mapping quality of > 20 following best practice guidelines for pool-seq (Schlötterer, Tobler, Kofler, & Nolte, 2014). Duplicate alignments were removed with samtools rmdup (v 1.3.1; Li et al., 2009) and regions around indels were re-aligned using picard (v. 1.118; Broad Institute), GATK (v. 3.2-2; McKenna et al., 2010) and samtools. Separate .bam files for each of the sequenced populations were finally merged using bamtools (v. 2.4.0; Barnett, Garrison, Quinlan, Strömberg, & Marth, 2011).

Over 80% of reads were properly mapped and retained in all samples. The mean coverage for Seward samples was nearly twice that of the other samples (figure S1 and figure S2). To remove the potential for this difference to cause artefacts in downstream analyses, the .bam files for Seward were down-sampled to contain 94.1 million reads (the average across the 5 remaining populations). Median empirical coverage was between ~62 and 88x (table S2; figure S3) and much less variable among the populations, allowing common maximum and minimum thresholds to be set based on the aggregate distribution. Allele counts for each population at each genomic position were obtained with samtools mpileup (v. 1.3.1; Li et al., 2009) using options to skip indel calling as well as ignoring reads with a mapping quality < 20 and sites with a base quality < 15. This was followed by the heuristic SNP calling software PoolSNP using a minimum count of 5 to call an allele, and a minimum coverage of 37 and a maximum coverage < 95^th^ percentile of the scaffold-wide coverage distribution to call a SNP (Kapun et al., 2019). Even if all these filters were passed, an allele was not considered if its frequency was < 0.001. Finally, we only considered SNPs on scaffolds > 10kb in length. The final set consisted of 2,190,511 biallelic SNPs that could be placed on scaffolds ordered along the chromosomes and were used in downstream analyses.

To test for an association between the four environmental variables and genetic differentiation we used BayeScEnv (v. 1.1; Foll & Gaggiotti, 2008; de Villemereuil & Gaggiotti, 2015). BayeScEnv fits a model of F_ST_ to population differentiation for each locus, incorporating environmental differentiation as a predictor (included as the parameter *g*) while fitting two locus-specific effects, one for environmental selection, and the other for other processes (demography, other types of selection). It therefore controls for confounding effects of population structure/relatedness in testing for an association with environmental variables. BayeScEnv was run with five pilot runs of 1000 iterations each, followed by a main chain of 4,000 iterations of which 2,000 were discarded as burn-in. Four MCMC chains were run for each analysis to evaluate convergence of the chains to common parameter estimates. Because of the unbalanced number of males (and the resulting variation in ratios of X:Y chromosomes) in the pools, BayeScEnv analyses were performed separately on SNPs that could be assigned to the autosomal linkage groups (chromosomes) and the X chromosome. Raw count data were used for the autosomal data. For X linked SNPs, allele count data were scaled to the known number of X chromosomes in the pool using *n_eff_*, the effective sample size taking into account the multiple rounds of binomial sampling inherent to a pool-seq design (Kolaczkowski et al., 2011; Feder, Petrov, & Bergland, 2012). *N_eff_* scaled the allele counts at each SNP downwards based on the known number of chromosomes in the pool (see table 1).

Chains were assessed for convergence with the “coda” R package (v. 0.19-1; Plummer, Best, Cowles, & Vines, 2006). Convergence was reached across the four chains for most analyses and parameters (potential scale reduction factors (PSRFs) of ~1 in a Gelman-Rubin diagnostic test; figures S4-S7), except for analyses of autosomal SNPs and PC2 as the environmental variable which showed mild convergence problems (PSRF = 1.71), although parameter estimates agreed well with the other chains. Thus, this first chain (black line in figures S4-S7) was discounted for all parameters and only estimates from the remaining 3 chains were used. The union of significant SNPs (using q-values for the *g* parameter, describing the association between genetic differentiation at a locus and environmental differentiation, to control the FDR at 0.05, i.e. SNPs with q-values < 0.05) across these chains were taken as the final candidate SNPs.

Finally, population genetic statistics (π, and Tajima’s D) were computed in windows of 10kb with a step size of 5kb using methods implemented for pool-seq data (Kapun et al., 2019). Windows contained a mean of 488.2 ±1.2 SNPs (mean ±SE). These statistics were only computed for scaffolds with a length > 50kb. SNP-wise F_ST_ was computed for each population with the R package “poolfstat” (v. 1.1.1; Hivert, Leblois, Petit, Gautier, & Vitalis, 2018) by first computing all pairwise values, and then deriving population specific F_ST_ values by averaging across all pairwise values where a population was included. We computed 95% confidence intervals for mean SNP-wise F_ST_ on each chromosome from the distribution of mean F_ST_ values across 100 bootstrap samples of SNPs. See the supplementary material for pseudocode commands of the key pipeline steps.

We performed GO term enrichment analyses with DAVID (v. 6.8; Huang et al., 2009a, b) and GOwinda (v. 1.12; Kofler & Schlötterer 2012), which accounts for differences in gene length. For DAVID, since the *D. montana* annotation contains information about orthologs in *D. virilis*, we extracted all genes within 10kb upstream or downstream of a candidate SNP with an ortholog in *D. virilis* and submitted them to DAVID. For analyses with GOwinda, because the gene-sets need to be given manually, we obtained gene-sets from FuncAssociate3 (Berriz, Beaver, Cenik, Tasan & Roth 2009). As above, we considered SNPs within 10kb of a gene (--gene-definition updownstream10000) and performed 1 million simulations to obtain the empirical p-value distribution. Regulatory elements, such as enhancers, and transcription factor binding sites, can occur up to 1Mb up- or downstream from a target gene in other species (e.g. Maston, Evans & Green. 2006; Chan et al., 2010; Werner, Koshikawa, Williams & Carroll 2010; Pennacchio, Bickmore, Dean, Nobrega, & Bejerano 2013) but generally lie within 2kb of a gene region in *D. melanogaster* (Arnosti, 2003), thus 10kb represents a compromise.

## RESULTS

### Cold tolerance measures, bioclimatic variables and population geography

Across individuals, there was no evidence of an association between minimum critical temperature (CTmin; pooled across sexes) and the chill coma recovery time (CCRT; cor = 0.08, p = 0.07). Neither was there evidence for an association between these traits across populations (cor = 0.22 p = 0.68). CTmin was significantly different between sexes, with females being more cold tolerant (t = 3.14, p = 0.002). While latitude appears to have a non-linear effect on CTmin (figure 2A and figure S8A), the Effective Degrees of Freedom (edf = 1) of the partial effect of latitude suggested a largely linear effect after accounting for altitude (F = 40.4, p < 0.001). Altitude had a more complex effect (edf = 1.262, F = 24.0, p < 0.001), but with a strong altitude outlier in Crested Butte (table 1), this result should be taken with some caution. Moreover, the overall adjusted R-squared was low at only 0.09. For CCRT, only latitude had a significant effect (edf = 1, F = 17.2, p < 0.001; figure 2B). Taken together, CTmin was lower for populations at higher latitudes as well as for populations at higher altitudes (figure 2A and figure S8A). As one would expect, CTmin showed on average lower values (figure 2A and figure S8A), and CCRTs were shorter (figure 2B and figure S8B), at higher latitudes, meaning that more northern and higher altitude populations show higher cold tolerance.

**Figure 2.**
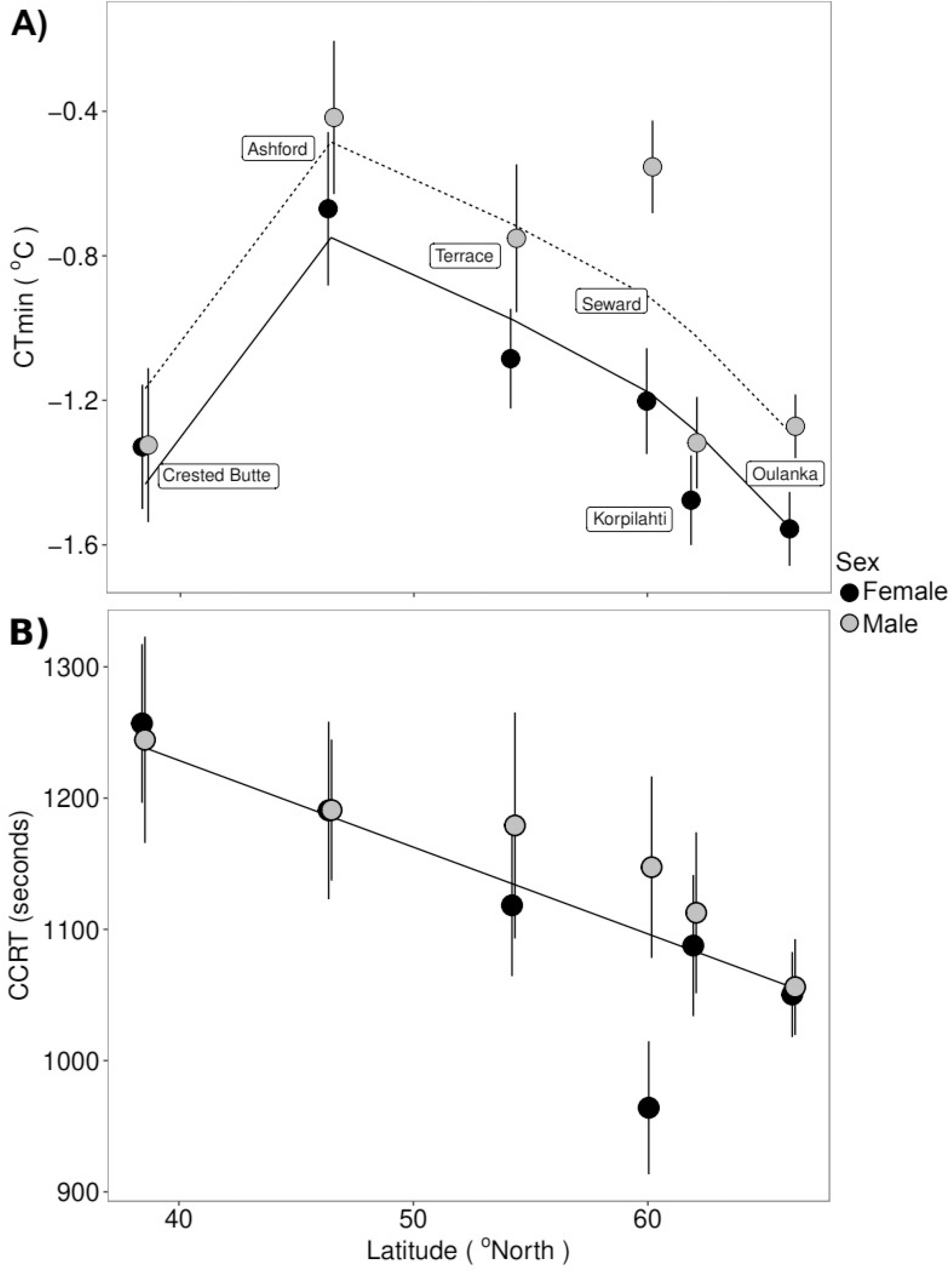
Population phenotypes. **A)** and **B)** show the variation in CTmin and CCRT across populations and latitude, respectively. Solid and dashed lines show the predicted values from the partial effect of latitude from the best model (see Results) for males and females, respectively. In **B)**, although the points are plotted separately for males and females, the best model only included latitude as a covariate. In both panels the points and error bars give the mean and standard errors, respectively. Figure S8 shows the full data for each population.

We performed a Principal Component Analysis (PCA) of the WorldClim climate data for a total of 24 *D. montana* populations, from which we had collected samples and where climate data were available. We wanted to examine whether the populations chosen for the cold tolerance tests and genome sequencing were representative of the range of environmental variation experienced by the species. These analyses identified four Principal Components (PCs) that together explained about 98% of the variation (figure 3A and figure S9). The first two PCs separated the populations roughly by a measure of distance from the coast (PC1) and then by latitude and altitude (PC2). PC1 explained ~54% of the variation (figure 3A and B) and loaded heavily on climate and biological variables associated with precipitation and temperature such as “Mean Temperature of Coldest Quarter”, “Precipitation of Wettest Month”, “Annual Precipitation” (see figure S9 and table S3). Meanwhile, PC2 explained ~23% of the variation (figure 3A and C) and loaded heavily on biological variables that are associated with latitudinal clinality, e.g. “Mean Diurnal [Temperature] Range,” and “Isothermality” which is the diurnal range divided by the mean “Annual [Temperature] Range”. The remaining PCs (PC3 and PC4) explained about 11.5 and 5% of the variation, respectively and did not capture as much of the climatic variation, and were therefore not considered further. Latitude (Spearman’s Rank Correlation: rho = −0.50, p = 0.01) but not altitude (rho = −0.19, p = 0.39) correlated significantly with PC1. However, both altitude (rho = −0.50, p = 0.02) and latitude (rho = −0.59, p = 0.003) correlated with PC2 (see also figure 3C). Importantly, these patterns were fairly robust also when performing PCA using only the six populations for which genomic data were collected. Loadings on PC1 were highly comparable, and although correlations with latitude and altitude did not achieve statistical significance, they were similar in magnitude and direction (Latitude vs. PC1: rho = 0.43, Altitude vs. PC1: rho = −0.14, p=0.8). For PC2 the correlations were also not statistically significant and differed for latitude both in magnitude and direction (rho = 0.6, p = 0.2), while for altitude, the correlation remained similar and marginally non-significant (rho = −0.82, p=0.06). Moreover, while the range of the sequenced populations covered ~70% of the range of all populations for PC1, they covered only ~50% of the range for PC2 (figure 3A). This suggests that environmental variation across the six populations selected for sequencing reflects that experienced by all 24 populations at least for the PC1 axis. Therefore, the relationship between environmental variables and genetic differentiation in the samples selected for pool-seq is likely to reflect true patterns across populations of *D. montana*, though power might be somewhat reduced for PC2. For all subsequent analyses, we used the results of the PCA using all 24 populations, and to examine the association between climate and phenotype, we compared these across the six populations. CTmin was positively correlated with PC1 (Pearson’s correlation coefficient (cor) = 0.94, p < 0.01) and had a marginally non-significant association with PC2 (cor = 0.75, p = 0.09). However, CCRT showed no relationship with either PC1 (cor = 0.09, p = 0.87) or PC2 (cor = −0.46, p = 0.36) although the small sample sizes (N = 6 in all cases) make reliable conclusions difficult (see figure S10).

**Figure 3.**
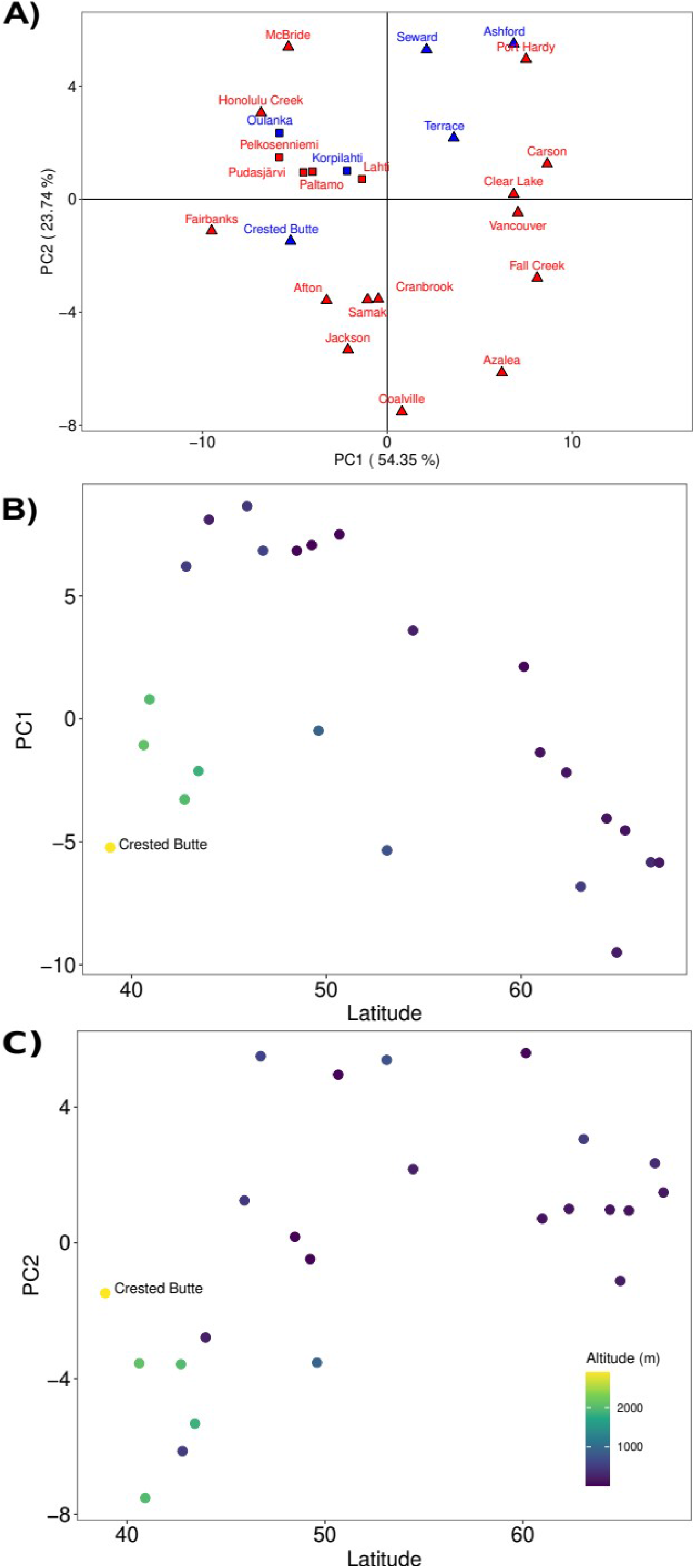
Principle Components Analysis. **A)** The distribution of all populations along the two first PC axes. The loadings of variables on each axis can be found in table S3. Blue circles give the populations that have been pool-sequenced for this study, red squares and triangles give the other Finnish and North American populations, respectively. **B)** and **C)** give PCs 1 and 2 as a function of latitude and altitude, respectively. The legend for altitude in both **B)** and **C)** is given in **C)**.

### Genomic analyses

The number of SNPs with a significant (q-value < 0.05) association between population-based FST, the two cold tolerance measures (CCRT and CTmin), and the two PCs of the bioclimatic data varied from 312 (chromosome 3, CCRT) to 2,480 (chromosome 4, PC2) across the chromosomes (figure 4). Using PC1 as an environmental variable with BayeScEnv gave a total of 2,976 and 1,528 SNPs with a q-value < 0.05 on the autosomes and on the X chromosome, respectively. Interestingly, the distribution across the chromosomes was not random. Using the distribution of all SNPs to obtain expected counts, there was a significant deviation from expectation (X^2^ = 2,906.4, d.f. = 4, p < 0.001). There were many more SNPs than expected on chromosome 4 (1,432 vs. 954) and on the X chromosome (1,528 vs. 526; figure 4). Results were similar for PC2 with 6,607 and 1,861 SNPs with a q-value < 0.05 on the autosomes and X chromosomes, respectively. Again, there was a significant deviation from the expected distribution of SNPs across the chromosomes (X^2^ = 1,681.9, d.f. = 4, p < 0.001) with an overrepresentation on the 4^th^ (2,480 vs. 1,794) and the X chromosomes (1,861 vs. 989; figure 4).

**Figure 4.**
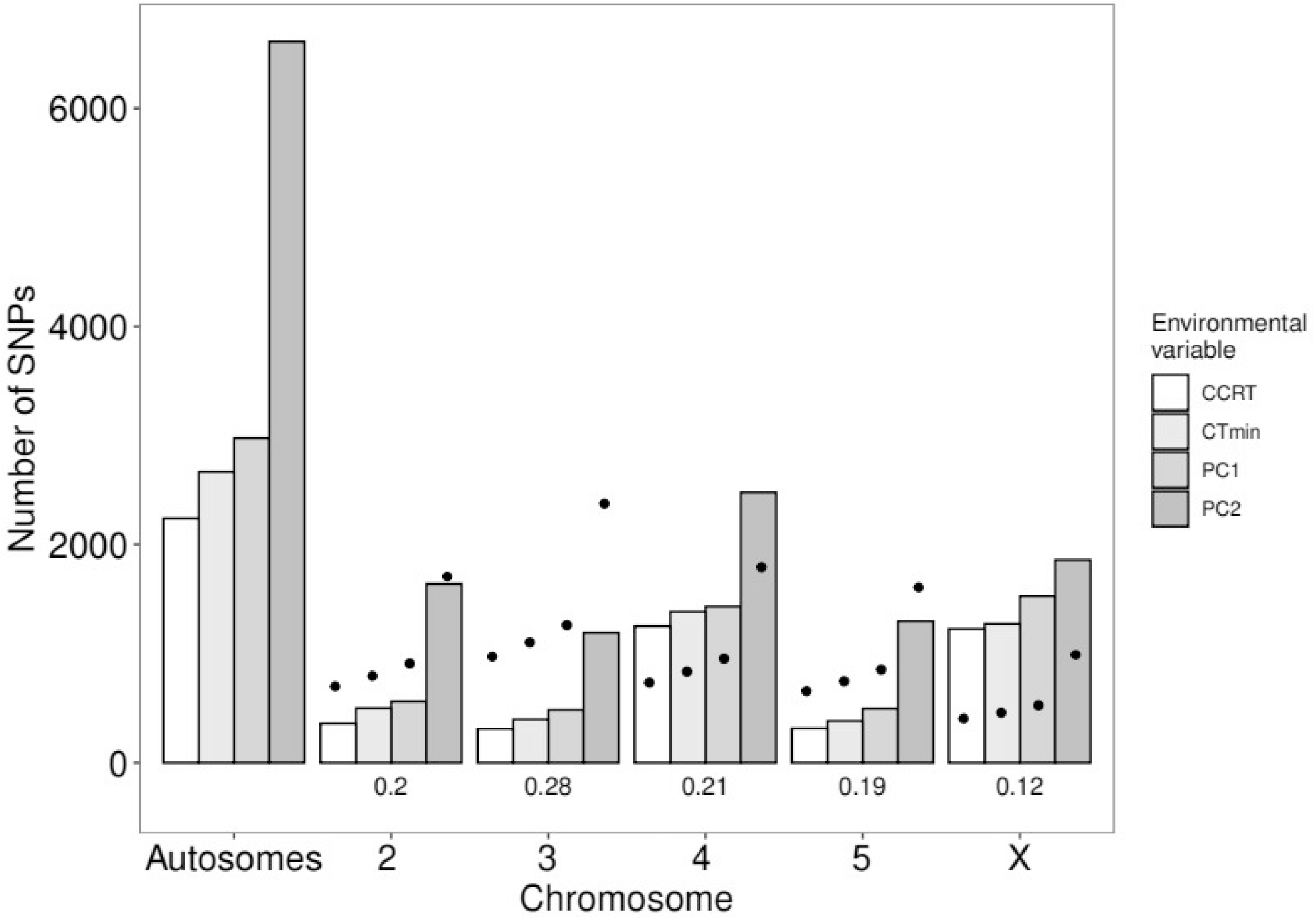
The observed counts of candidate SNPs (i.e. SNPs with a q-value < 0.05) across chromosomes for each environmental variable (see Methods). The total number of SNPs, and the proportion of all SNPs, on each chromosome are given below each set of bars. The expected counts on each chromosome, obtained from the proportions of all SNPs across chromosomes, are shown as points aligned with each bar.

We also used average CTmin values per population in similar analyses and found a total of 2,668 and 1,272 SNPs with a q-value < 0.05 on the autosomes and on the X chromosomes, respectively. The pattern of significant deviations from expected distributions (X^2^ = 2,526.2, d.f. = 4, p < 0.001) was also due to an excess on the 4th (1,383 vs. 835) and the X chromosomes (1,272 vs. 460; figure 4). Similar results were found for CCRT with a total of 2,240 and 1,228 SNPs with a q-value < 0.05, respectively. Once again, there was a significant deviation from the expected distribution of SNPs (X^2^ = 2,825.6, d.f. = 3, p < 0.001) with and excess on the 4^th^ (1,252 vs. 735) and the X chromosomes (1,228 vs. 405; figure 4). Manhattan plots of the distribution of SNPs across chromosomes are given in figures S11-S14.

To more closely examine the loci implicated in the four BayScEnv analyses, we identified genes within 10kb of, or containing the candidate SNPs (table S4). Overall, there is quite a large overlap among these genes with ~39% (1,102 in total) of them being shared by all the four analyses (table S4, figure S15). This is far more than would be expected (mean±SE: 3.6±0.06) by randomly re-sampling (without replacement) a number of genes equivalent to that associated with top SNPs for each BayScEnv analysis from the *D. montana* annotation, then computing the four-way intersection. This large observed overlap therefore probably reflects a genuine overlap among the underlying top SNPs (data not shown). Some (147, ~13%) of these common genes are novel to *D. montana* (i.e. not annotated in *D. virilis* or in other *Drosophila* spp.) and therefore have no annotation, but 955 have an identifiable *D. virilis* ortholog (table S4). The second largest set are those genes unique to the analysis of PC2 as an environmental covariate (figure S15). We performed functional enrichment analyses with DAVID keeping all annotation clusters with an enrichment score > 1.3 (corresponding to an average corrected p-value of 0.05). This revealed several common categories of genes associated with the climatic variables and population phenotypes (table S5). For example, terms associated with membrane and transmembrane structures, immunoglobulins, HAD hydrolase and nucleotide binding were enriched in most of the variables (table S5). Interestingly, there were also several gene ontology categories that were only enriched in one of the variables, such as glycoside and ATPase hydrolase in CCRT and ion channels and transport, as well as metal binding in PC1 (table S5). GOwinda analyses revealed significant enrichment of GO terms (after accounting for variation in gene lengths and correcting for multiple testing) only for genes near SNPs associated with variation in CCRT across populations. Interestingly, the term “carbohydrate derivative binding” was identified among the most enriched terms (table S6), which agrees closely with some terms identified using DAVID for the same variable (table S5). Similarly, for PC2, “nucleotide binding” was among the top scoring enriched terms in both GOwinda and DAVID analyses, though this was not significant after controlling for multiple testing in GOwinda (table S5, table S6)

We then compared loci near candidate SNPs with genes implicated in previous studies of climatic adaptation in *D*. *montana* including gene expression studies of traits connected to diapause and cold-tolerance (Kankare et al., 2010; 2016; Parker et al., 2015; 2016). Additionally, several candidate genes have been identified near the most significantly differentiated SNPs among *D. montana* populations from Oulanka (Finland), and from North American populations in Colorado and Vancouver (Parker et al., 2018). Finally, quasi-natural selection experiments identified several genes within 10kb of SNPs responding to selection for a shorter CDL for diapause induction (Kauranen et al., 2019). We tested for an overlap between the total set of genes within 10kb of outlier SNPs from all of the BayScEnv runs (N = 2,694) and the candidate gene sets identified in earlier studies (see table S7 for the gene sets and studies used). We computed a bootstrap distribution of overlaps by sampling 2,694 random genes from the *D. montana* annotation (N = 13,683). For each of the gene sets from previous studies this was done 100 times and the distribution compared to the empirical overlap. Results are given in table 2. In all the cases the empirical overlap was greater than expected by chance with empirical p-values < 0.05 (ranging from < 0.0001 to 0.01; table 2). The only gene that was found in all five of the previous studies used and in the comparison here is called *sidestep II* (*side-II*; table S8). Unfortunately, there is no information available about the biological processes or molecular functions connected to it. Moreover, from 44 other genes that were common to four of our previous studies and this study (table S8) most (27) have an ortholog in *D. melanogaster*. These genes have molecular functions such as transmembrane signaling or transporter, acetylcholinesterase, ATP binding, protein serine/threonine kinase, carboxylic ester hydrolase, or Rho guanyl-nucleotide exchange factor activity (Thurmond et al., 2019). Many of the genes are also connected to metal ion, nucleid acid or zinc ion binding (table S8). After identifying information on molecular or biological function and Interpro domains, eventually only five genes remained for which there was no information available (table S8).

**Table 2.** The overlap of the union of genes within 10kb of SNPs associated with PC1, PC2, CTmin, and CCRT and previous candidate gene sets (see table S7). Shown are the citation for the original study, the number of genes identified from the original study, the number of overlapping genes with the candidate gene set from this study, and the mean and 95% CI from 10,000 re-sampled sets of candidate genes.

| Reference | N genes | N overlaps | Mean [95%CI]<br>re-sampled N overlaps | Empirical p-value |
| --- | --- | --- | --- | --- |
| Kankare et al.,<br>2010 | 14 | 8 | 2.77 [0, 6] | 0.0004 |
| Parker et al.,<br>2015 | 42 | 13 | 6.28 [2, 11] | 0.001 |
| Kankare et al.,<br>2016 | 3 929 | 946 | 773.14 [732, 814] | < 0.0001 |
| Parker et al.,<br>2016 | 130 | 27 | 18.72 [11, 26] | 0.01 |
| Parker et al.,<br>2018 | 2 114 | 629 | 414.7 [382, 448] | < 0.0001 |
| Kauranen et al.,<br>2019 | 1 751 | 402 | 344.64 [314, 375] | < 0.0001 |

Examination of population genetic parameters identified the Crested Butte population as anomalous. The distribution of Tajima’s D is centered close to zero in most populations, being slightly more negative in North American populations (figure S16A). However, Crested Butte is an outlier with a greatly reduced genome-wide Tajima’s D (figure S16A). Similarly, diversity (π) is also lower in this population than in other populations. There is no overall relationship between latitude and π (Spearman’s rho = 0.14, N = 6, p = 0.8; figure S16B) but there is a strong correlation between latitude and Tajima’s D which is influenced by this population (with Crested Butte: rho = 0.88, N = 6, p = 0.03; without Crested Butte: rho = 0.8, N = 5, p = 0.13). Although Crested Butte occurs at a much higher altitude (>2800 meters) than other populations neither Tajima’s D nor π correlated significantly with altitude (Tajima’s D: rho = −0.6, N = 6, p = 0.24, π: rho = −0.6, N = 6, p = 0.24). Furthermore, F_ST_ was similar across all populations and chromosomes with the exception of Crested Butte which remained an outlier with high F_ST_ (figure 5). Bootstrapped 95% confidence intervals are presented to give a guide to statistical significance of differences among populations and confirm that Crested Butte has high F_ST_ (figure 5). Finally, F_ST_ was always highest on chromosome 4 and the X chromosome, complementing the results seen in BayeScEnv analyses (figure 4 *c.f.* figure 5) and as expected if ecological selection is influencing genomic divergence.

**Figure 5.**
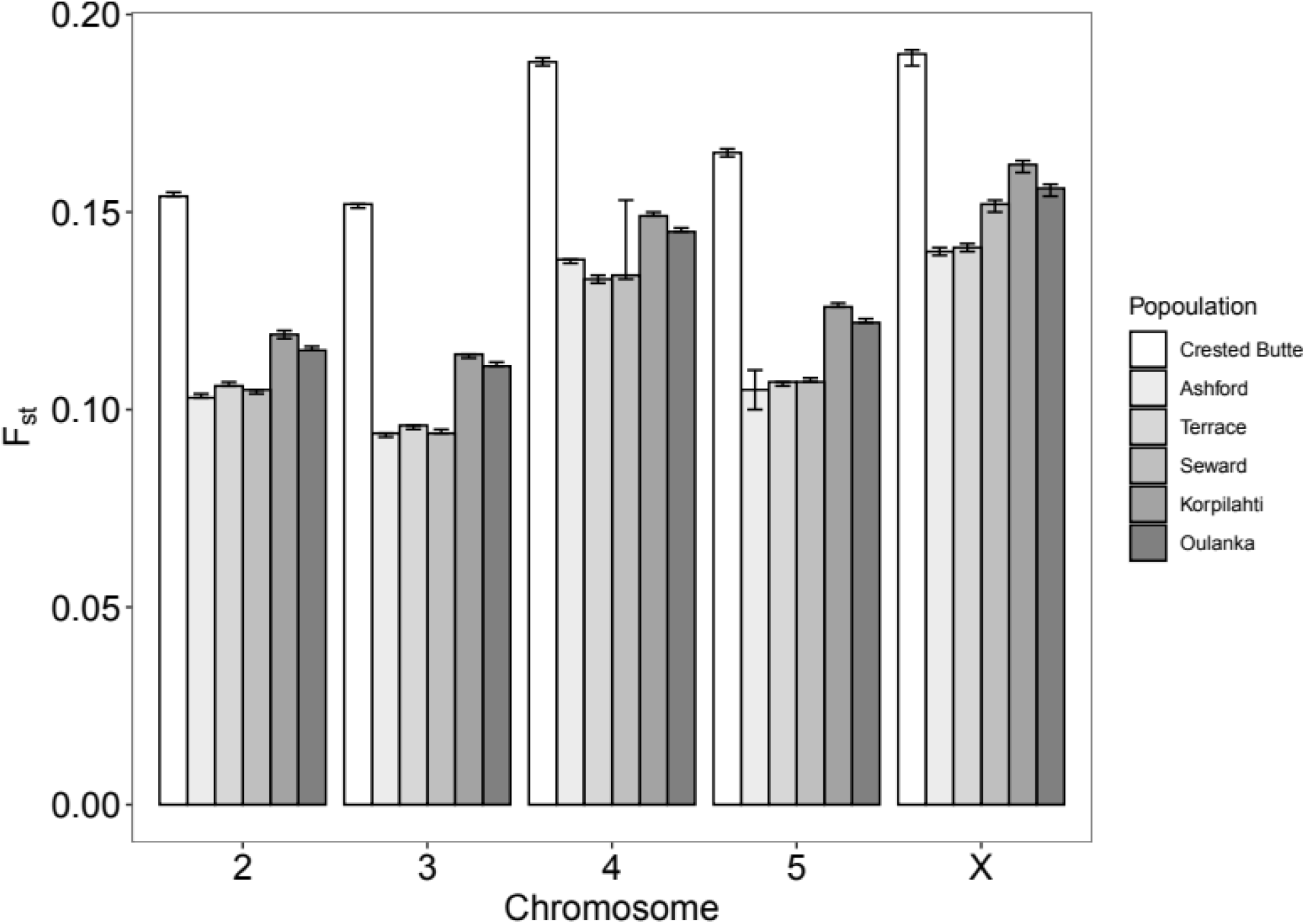
Mean, SD, and confidence intervals of the F_ST_ values across SNPs on each chromosome and population. F_ST_ was summarised for each SNP by computing all pairwise values then deriving population specific F_ST_ values by averaging across all pairwise values containing that population. 95% confidence intervals (given by the error bars) are derived from 100 bootstrap samples of SNPs on each chromosome.

## DISCUSSION

Detecting genomic signatures of climatic adaptation is an important, but challenging, task. Here we use multiple sources of evidence to study ecological adaptation and population divergence in a highly cold tolerant species of *Drosophila*, *D. montana*. This species is characterised by a wide circumpolar distribution extending to high latitudes both in North America and Europe, and to high altitudes in the southern part of its range in the Rocky Mountains of North America. These habitats impose extreme seasonal and climatic selective pressure. We collected bio-climatic data from 24 populations along a latitudinal gradient of about 2,900 km in North America, and six populations from a gradient of 720 km in Finland. We characterised population level cold-tolerance for six populations from these ranges using two methods, critical thermal minimum (CTmin) and chill coma recovery time (CCRT) and show that CTmin were lower and CCRT shorter in higher latitude populations, as one would expect. Thus, northern populations are more cold-tolerant. Finally, we performed pool-seq of these six populations to investigate the association between genomic and environmental differentiation.

The two methods examining cold tolerance gave somewhat different results, as CTmin, but not CCRT, differed significantly between sexes, with females having lower CTmin than males. In an earlier study investigating seasonal changes in *D. montana* CCRTs, only one out of six comparisons showed a significant difference between sexes (Vesala, Salminen, Kostal, Zahradnickova, & Hoikkala, 2012) and Gibert, Moreteau, Petavy, Karan & David (2001) did not detect sex-specific differences in CCRT in any of 84 *Drosophila* species they studied. However, several studies of *D. melanogaster* have detected shorter CCRT in females than in males, suggesting that females are more cold tolerant than males (David et al., 1998; Andersen et al., 2015; Bauerfeind, Kellermann, Moghadam, Loeschcke, & Fischer, 2014), perhaps related to their greater body mass (e.g. Wilder et al., 2010). Consequently, the extent and adaptive significance of sex-specific differences in CCRT in *Drosophila* remains unclear.

We derived principal components to summarise WorldClim climatic variables using data from all the 24 populations. The first principal component (PC1) separated populations roughly by a measure of “distance inland” and loaded heavily on climate and variables associated with precipitation and temperature. These results follow the geographic distribution of the populations, for example, the population with highest values for PC1 is Ashford, which is on the Pacific coast and receives most rain, but also experiences warm summers and mild winters. Principal component 2 (PC2), loaded heavily on bioclimatic variables associated with latitudinal clinality, which also mapped onto the populations intuitively as those with higher values on PC2 also occurred at higher latitudes. Interestingly, CTmin values were positively correlated with PC1, but not with PC2, while CCRT showed no relationship with either of these components. This suggests that CTmin and CCRT measure at least slightly different biochemical or physiological mechanisms (see e.g. MacMillan, Williams, Staples, & Sinclair, 2012; Findsen, Pedersen, Petersen, Nielsen & Overgaard 2013), and could hence be correlated with different climatic variables and also show relatively weak correlation to each other (Andersen et al., 2015). Indeed our results found no significant correlation in CTmin and CCRT across populations. It is also interesting to note that high altitude populations have very similar values on PC1 and PC2 to high latitude populations, most likely reflecting the similar climates in these populations. These similarities in climate are likely also underlying the similarities in CTmin between the high altitude population (Crested Butte) and the high latitude populations from Finland.

The Bayesian analysis identified SNPs showing an association of genetic differentiation with climatic and phenotypic variation. The extent to which the loci associated with the phenotypes and adaptations to different climatic conditions are shared indicates that these are closely associated in influencing genome evolution. We found that genes near SNPs showing a significant association between genetic and climatic differentiation overlapped to a large extent with genetic and phenotypic differentiation among populations. The largest intersection set, containing 1,102 genes, was the one containing genes near SNPs associated with all the four variables examined (PC1, PC2, CTmin and CCRT). However, PC2 loads heavily on bioclimatic variables relating to latitude, and our analysis using PC2 as a covariate has a large number of private genes (see figure 4), suggesting that there is also a substantial amount of genetic variation underlying adaptations to latitude unrelated to the phenotypic measures we have quantified. Moreover, this should be viewed with caution because the range of values for PC2 represented by the sequenced samples involve only 50% of the total variation in this axis. Nevertheless, our study is an excellent example of how strong ecological selection may be detected in genomic studies. In particular, because the Bayesian methods examining both ecological variables and relevant phenotypes gave significant overlap amongst the associated loci, and that these are further associated with more broad genomic differentiation between populations, gives confidence that we are consistently identifying genes associated with ecological selection.

Analyses of the functional annotation of these genes strengthens our conclusions that climate driven adaptation is important. Regions near SNPs associated with climatic variables were enriched for genes previously identified as candidates related to cold tolerance, diapause and responses to changes in day length in *D. montana* (Kankare et al., 2010; 2016; Parker et al., 2015; 2016). Thus, genetic variation across populations of these flies may be largely shaped by differences in ecological and climatic variation. This ecological specialisation may have also contributed to the divergence of *D. montana* from its relatives. Parker et al., (2018) surveyed the rates of molecular evolution in eleven cold tolerant and non-cold tolerant species of *Drosophila*. The genes found to be evolving at faster rates in cold-tolerant species were enriched for many of the same functional categories as in our current study, including e.g. membrane and transmembrane proteins and immunoglobulins (Parker et al., 2018).

The fact that many of the same annotation terms are enriched in clusters for all environmental variables in DAVID analyses, suggests that similar genes or biochemical pathways are involved in these adaptations. Membrane proteins and lipids are an important determinant of membrane and cuticular permeability at different temperatures, which in turn has an effect on the resistance to desiccation stress in insects (Gibbs, 2002; Stanziano, Sové, Rundle, & Sinclair, 2015). Importantly, there is evidence for a close link between the desiccation stress response and cold tolerance across species and in *Drosophila* in particular, suggesting an overlap in some of the mechanisms involved (Sinclair, Nelson, Nilson, Roberts, & Gibbs, 2007). Cold-hardy lines of *D. melanogaster* are known to exhibit elevated lipid metabolism, perhaps in order to allow rapid lipid membrane modification (Williams et al., 2016) in different environmental conditions. Pleckstrin homology (PH) domain was one of the functional clusters found in both the current study and in the species comparison of Parker et al., (2018). This domain is a flexible module of 100-120 amino acids which interacts with a variety of different ligands, composing a protein-protein interaction platform (Scheffzek & Welti, 2012). As changes in the membrane lipid biochemistry form an integral part of the cold tolerance response, genes associated with PH may assist in homeoviscous adaptation i.e. alteration of membrane phospholipid composition to maintain fluidity at low temperatures (Sinensky, 1974). Interestingly, the only gene found in all five of our previous studies of cold tolerance in *D. montana* was *sidestep II* (*side-II*). Unfortunately, there is no information available for the biological processes or molecular functions associated with this gene, but *side-II* has protein features including immunoglobulin and immunoglobulin-like domain superfamily (Thurmond et al., 2019) and could be involved in immunological processes during cold response of the flies (Parker et al., 2021). Insects are known to produce a diverse range of antimicrobial peptides and proteins as part of their immune activity against viruses, bacteria, fungi and parasites (Mylonakis, Podsiadlowski, Muhammed, & Vilcinskas, 2016) and hence immune responses could be part of a general stress response in cold tolerance (Sinclair, Ferguson, Salehipour-shirazi, & MacMillan, 2013; Ferguson, Heinrichs, & Sinclair, 2016). We also find confirmation of some enriched GO terms in GOwinda analyses that control for variation in gene length, although overlap is not substantial. In sum, our results indicate that some of the same biochemical processes that are targeted by selection on larger evolutionary scales (i.e. across species), are also involved in local adaptation for different populations within a species, providing a rare bridge between the processes of population differentiation and speciation.

At the chromosomal level, we found an over-representation of loci associated with ecological selection on chromosomes X (which corresponds to chromosome 2L in *D. melanogaster*) and 4. It is well-known that X chromosomes can generally evolve quickly due to selection on semi-recessive advantageous loci in the hemizygous sex, and smaller effective population size (Charlesworth, Coyne, & Barton, 1987) and are often most divergent between closely related species (Abbott, Norden, & Hansson, 2017; Ellegren et al., 2012). However, there is no obvious reason to expect faster divergence in chromosome 4. *D. montana* populations have considerable chromosomal polymorphism (e.g. Patterson and Stone, 1952; Stone, Guest, & Wilson, 1960; Morales-Hojas, Päällysaho, Vieira, Hoikkala, & Vieira, 2007) and American populations have been classified into several chromosomal forms on the basis of their geographical origin, size and chromosome structure (Throckmorton, 1982). Moreover, both the X chromosome and chromosome 4 of *D. montana* are known to harbour polymorphic inversions (Stone et al., 1960; Morales-Hojas et al., 2007) and such inversions have often been found to vary clinally in other systems (e.g. Kolaczkowski et al., 2011; Cheng et al., 2012; Kapun et al., 2016). Recent genomic data confirms that there are several inversions in both chromosomes X and 4, but we do not yet have any detailed information on their distributions within populations (Poikela et al., unpublished). The reduced recombination within inversions can independently capture advantageous alleles under selection (Kirkpatrick & Barton, 2006) and divergence is often greater in chromosomes carrying inverted regions and non-colinear regions (Lohse, Clarke, Ritchie, & Etges, 2015) and such regions may divergence more quickly during speciation with gene flow. Indeed, here we found that the X chromosome and chromosome 4 always have the highest levels of F_ST_ across all populations, as expected if the ecological selection on loci on these chromosomes influenced overall patterns of genomic diversion, perhaps due to hitchhiking.

Finally, our findings include intriguing result regarding the Crested Butte population which shows reduced genetic variability and a substantial reduction of Tajima’s D relative to the other populations. This population occurs at a very high altitude (>2800 m) and also shows reproductive incompatibilities with other populations (Jennings et al., 2014). It may have also been bottlenecked during its adaptation to this high altitude, so population expansion or selected sweeps could be prevalent within this population (Garlovsky et al., 2020). Whether ecological specialisation is associated with the spread of incompatibilities is an intriguing possibility.

## CONCLUSION

Identifying the genetic variation that underlies population divergence and ultimately speciation remains a challenge. Detecting associations between genetic and environmental differentiation at loci across populations has become a popular approach. Here we apply Bayesian methods to detect such loci across populations of *Drosophila montana*, which is an extremely cold-tolerant *Drosophila* species where we expect strong ecological selection. We identify many genes that are associated with both climate variables and population-level cold-tolerance phenotypes. These genes overlap with candidate genes from other studies of variation in cold-tolerance in *D. montana* and were also over-represented on chromosomes X and 4 which may be associated with inversion polymorphisms known to be present in these chromosomes. Our study thus provides a clear example of how strong ecological selection can be detected in genome studies, using Bayesian methods to detect local adaptation in combination with studies of ecologically important variables and phenotypes.

## Data Availability

Raw reads have been deposited with NCBI (https://www.ncbi.nlm.nih.gov) under the BioProject accession PRJNA588720.

## Acknowledgments

This work was supported by a combined Natural Environment Research Council and St Andrews 600th Anniversary PhD Studentship grant (NE/L501852/1) to RAWW, The Ella and Georg Ehrnrooth Fellowship to VT, NERC grant (NE/P000592/1) to MGR, and Academy of Finland project 267244 to AH and projects 268214 and 322980 to MK. Bioinformatics and Computational Biology analyses were supported by the University of St Andrews Bioinformatics Unit which is funded by a Wellcome Trust ISSF award (grant 105621/Z/14/Z). We would also like to thank R. Tapanainen and J. Kinnunen for assistance in lab work and Oscar E Gaggiotti for advice on BayeScEnv analyses. We also thank four anonymous reviewers for their valuable comments that improved the manuscript.

## Author contributions

AH, MGR, and MK conceived the study. VT performed the phenotyping experiments. RAWW performed all analyses. RAWW, MK and MGR led the writing of the MS. All authors contributed to the final form of the manuscript.

## REFERENCES

[dataset] Wiberg, R. A. W., Tyukmaeva, V., Hoikkala, A., Ritchie, M. G. & Kankare, M.; 2020; Pooled sequencing of *Drosophila montana* wild population samples; NCBI BioProject; PRJNA588720

Abbott, J. K., Norden, A. K., & Hansson, B. (2017). Sex chromosome evolution: Historical insights and future perspectives. Proceedings of the Royal Society B: Biological Sciences, 284. doi: 10.1098/rspb.2016.2806

Andersen, J. L., Manenti, T., Sorensen, J. G., MacMillan, H. A., Loeschcke, V., & Overgaard, J. (2015). How to assess Drosophila cold tolerance: Chill coma temperature and lower lethal temperature are the best predictors of cold distribution limits. Functional Ecology, 29, 55–65. doi: 10.1111/1365-2435.12310

Andrews, S. (2015). FastQC: A quality control tool for high throughput sequence data. Retrieved from http://www.bioinformatics.bbsrc.ac.uk/projects/fastqc/

Arnosti, D.N. (2003). Analysis and function of transcriptional regulatory elements: Insights from Drosophila. Annual Review of Entomology. 48, 579–602. doi: 10.1146/annurev.ento.48.091801.112749

Barnett, D. W., Garrison, E. K., Quinlan, A. R., Strömberg, M. P., & Marth, G. T. (2011). Bamtools: A C++ API and toolkit for analyzing and managing BAM files. Bioinformatics, 27, 1691–1692. doi: 10.1093/bioinformatics/btr174

Barton, N. H., & Gale, K. S. (1993). Genetic analysis of hybrid zones. In R. G. Harrison (Ed.), Hybrid Zones and the Evolutionary Process. (pp. 13–45). Oxford, UK: Oxford University Press.

Bauerfeind, S. S., Kellermann, V., Moghadam, N. N., Loeschcke, V., & Fischer, K. (2014). Temperature and photoperiod affect stress resistance traits in *Drosophila melanogaster*. Physiological Entomology, 39, 237–246. doi: 10.1111/phen.12068

Bergland, A. O., Behrman, E. L., O’Brien, K. R., Schmidt, P. S., & Petrov, D. (2014). Genomic evidence of rapid and stable adaptive oscillations over seasonal time scales in Drosophila. PLoS Genetics, 10, e1004775. doi: 10.1371/journal.pgen.1004775

Berriz, G.F., Beaver, J.E., Cenik, C., Tasan, M., & Roth, F.P. (2009). Next generation software for functional trend analysis. Bioinformatics. 25, 3043–3044. doi:10.1093/bioinformatics/btp498

Bolger, A. M., Lohse, M., & Usadel, B. (2014). Trimmomatic: A flexible trimmer for Illumina sequence data. Bioinformatics, 30, 2114–2120. doi: 10.1093/bioinformatics/btu170

Bradbury, D., Smithson, A., & Krauss, S. L. (2013). Signatures of diversifying selection at EST-SSR loci and association with climate in natural Eucalyptus populations. Molecular Ecology, 22, 5112–5129. doi: 10.1111/mec.12463

Broad Institute. Picard. Retrieved from http://broadinstitute.github.io/picard/

Carneiro, M., Baird, S. J. E., Afonso, S., Ramirez, E., Tarroso, P., Teotónio, H., … Ferrand, N. (2013). Steep clines within a highly permeable genome across a hybrid zone between two subspecies of the European rabbit. Molecular Ecology, 22, 2511–2525. doi: 10.1111/mec.12272

Chan, Y.F., Marks, M.E., Jones, F.C., Villarreal, G., Shapiro, M.D., Brady, S.D., Southwick, A.M., Absher, D.M., Grimwood, J., Schmutz, J., Myers, R.M., Petrov, D., Jonsson, B., Schluter, D., Bell, M.A. & Kingsley, D.M. (2010). Adaptive evolution of pelvic reduction in sticklebacks by recurrent deletion of a Pitx1 enhancer. Science. 327 302–305. doi: 10.1126/science.1182213

Charlesworth, B., Coyne, J. A., & Barton, N. H. (1987). The relative rates of evolution of sex chromosomes and autosomes. The American Naturalist, 130, 113–146. https://www.jstor.org/stable/2461884

Chen, J., Källman, T., Ma, X., Gyllenstrand, N., Zaina, G., Morgante, M., … Lascoux, M. (2012). Disentangling the roles of history and local selection in shaping clinal variation of allele frequencies and gene expression in Norway spruce (*Picea abies*). Genetics, 191, 865–881. doi: 10.1534/genetics.112.140749

Cruickshank, T. E., & Hahn, M. W. (2014). Reanalysis suggests that genomic islands of speciation are due to reduced diversity, not reduced gene flow. Molecular Ecology, 23, 3133–3157. doi: 10.1111/mec.12796

David, R., Gibert, P., Pla, E., Petavy, G., Karan, D., & Moreteau, B. (1998). Cold stress tolerance in *Drosophila*: Analysis of chill coma recovery in *D. melanogaster*. Journal of Thermal Biology, 23, 291–299. doi: 10.1016/S0306-4565(98)00020-5

Ellegren, H., Smeds, L., Burri, R., Olason, P. I., Backstrom, N., Kawakami, T., … Wolf, J. B. W. (2012). The genomic landscape of species divergence in *Ficedula* flycatchers. Nature, 491, 756–760. doi: 10.1038/nature11584.

Endler, J. A. (1973). Gene flow and differentiation. Science. 179, 243–250. doi: 10.1126/science.179.4070.243

Endler, J. A. (1977). Geographic Variation, Speciation, and Clines. Princeton, NJ: Princeton University Press.

Feder, A. F., Petrov, D. A., & Bergland, A. O. (2012). LDx: Estimation of linkage disequilibrium from high-throughput pooled resequencing data. PLoS ONE, 7. doi: 10.1371/journal.pone.0048588

Ferguson, L. V, Heinrichs, D. E., & Sinclair, B. J. (2016). Paradoxical acclimation responses in the thermal performance of insect immunity. Oecologia, 181, 77–85. doi: 10.1007/s00442-015-3529-3526.

Findsen, A., Pedersen, T. H., Petersen, A. G., Nielsen, O. B., & Overgaard, J. (2014). Why do insects enter and recover from chill coma? Low temperature and high extracellular potassium compromise muscle function in Locusta migratoria. The Journal of experimental biology, 217, 1297–1306. doi: 10.1242/jeb.098442

Foll, M., & Gaggiotti, O. E. (2008). A genome-scan method to identify selected loci appropriate for both dominant and codominant markers: A Bayesian perspective. Genetics, 180, 977–993. doi: 10.1534/genetics.108.092221

Foll, M., Gaggiotti, O. E., Daub, J. T., Vatsiou, A., & Excoffier, L. (2014). Widespread signals of convergent adaptation to high altitude in Asia and America. American Journal of Human Genetics, 95, 394–407. doi: 10.1016/j.ajhg.2014.09.002.

Garlovsky, M. D., Yusuf, L. H., Ritchie, M. G. & Snook, R. R. (2020). Within-population sperm competition intensity does not predict asymmetry in conpopulation sperm precedence. Philosophical Transactions of the Royal Society. 375, 20200071. doi: 10.1098/rstb.2020.0071

Gautier, M. (2015). Genome-wide scan for adaptive divergence and association with population-specific covariates. Genetics, 201, 1555–1579. doi: 10.1534/genetics.115.181453

Gibbs, A. G. (2002). Lipid melting points and cuticular permeability: New insights into an old problem. Journal of Insect Physiology, 48, 391–400. doi: 10.1016/s0022-1910(02)00059-8

Gibert, P., Moreteau, B., Petavy, G., Karan, D., & David, J. (2001). Chill-coma tolerance, a major climatic adaptation among *Drosophila* species. Evolution, 55,1063–1068. doi: 10.1554/0014-3820(2001)055[1063:cctamc]2.0.co;2

Hijmans, R. J., Cameron, S. E., Parra, J. L., Jones, P. G., & Jarvis, A. (2005). Very high resolution interpolated climate surfaces for global land areas. International Journal of Climatology, 25, 1965–1978. doi: 10.1002/joc.1276

Hijmans, R. E., van Etten, J., Cheng, J., Mattiuzzi, M., Sumner, M., Greenberg, J. A., … Wueest, R. (2016). raster: Geographic Data Analysis and Modeling. Retrieved from: https://CRAN.R-project.org/package=raster

Hivert, V., Leblois, R., Petit, E. J., Gautier, M., & Vitalis, R. (2018). Measuring genetic differentiation from pool-seq data. Genetics, 210, 315–330. doi: 10.1534/genetics.118.300900

Hoekstra, H., Drumm, K., & Nachman, M. (2004). Ecological genetics of adaptive color polymorphism in pocket mice: Geographic variation in selected and neutral genes. Evolution, 58, 1329–1341. doi: 10.1111/j.0014-3820.2004.tb01711.x

Huang, D. W., Sherman, B. T. and Lempicki, R. A. (2009a). Systematic and integrative analysis of large gene lists using DAVID bioinformatics resources. Nature Protocols. 4, 44–57. doi: 10.1038/nprot.2008.211

Huang, D. W., Sherman, B. T. and Lempicki, R. A. (2009b). Bioinformatics enrichment tools: paths toward the comprehensive functional analysis of large gene lists. Nucleic Acids Research 37, 1–13. doi: 10.1093/nar/gkn923

Jennings, J. H., Mazzi, D., Ritchie, M. G., & Hoikkala, A. (2011). Sexual and postmating reproductive isolation between allopatric *Drosophila montana* populations suggest speciation potential. BMC Evolutionary Biology, 11. doi: 10.1186/1471-2148-11-68

Jennings, J. H., Snook, R. R., & Hoikkala, A. (2014). Reproductive isolation among allopatric *Drosophila montana* populations. Evolution, 68, 3095–3108. doi: 10.1111/evo.12535

Kankare, M., Salminen, T., Laiho, A., Vesala, L., & Hoikkala, A. (2010). Changes in gene expression linked with adult reproductive diapause in a northern malt fly species: A candidate gene microarray study. BMC Ecology, 10. doi: 10.1186/1472-6785-10-3

Kankare, M., Parker, D. J., Merisalo, M., Salminen, T. S., & Hoikkala, A. (2016). Transcriptional differences between diapausing and non-diapausing *D. montana* females reared under the same photoperiod and temperature. PLoS ONE, 11, e0161852. doi: 10.1371/journal.pone.0161852

Kapun, M., Fabian, D.K., Goudet, J., & Flatt, T. (2016). Genomic evidence for adaptive inversion clines in *Drosophila melanogaster*. Molecular Biology and Evolution, 33, 1317–1336. doi: 10.1093/molbev/msw016

Kapun, M., Barrón, M. G., Staubach, F., Vieira, J., Obbard, D. J., Goubert, C., … González, J. (2019). Genomic analysis of European *Drosophila melanogaster* populations on a dense spatial scale reveals longitudinal population structure and continent-wide selection, and unknown DNA viruses. BioRxiv. doi: 10.1101/313759

Kauranen, H., Kinnunen, J., Hiillos, A-L., Lankinen, P., Hopkins, D., Wiberg, R. A. W., … Hoikkala, A. (2019). Selection for reproduction under short photoperiods changes diapause-associated traits and induces widespread genomic divergence. Journal of Experimental Biology, 222, jeb205831. doi:10.1242/jeb.205831

Kellermann, V., Overgaard, J., Hoffmann, A. A., Flojgaard, C., Svenning, J. C., & Loeschcke, V. (2012). Upper thermal limits of Drosophila are linked to species distributions and strongly constrained phylogenetically. Proceedings of the National Academy of Sciences of the United States of America, 109, 16228–16233. doi: 10.1073/pnas.1207553109

Kirkpatrick, M., & Barton, N. (2006). Chromosome inversions, local adaptation and speciation. Genetics, 173, 419–434. doi: 10.1534/genetics.105.047985

Klappert, K., Mazzi, D., Hoikkala, A., & Ritchie, M. G. (2007). Male courtship song and female preference variation between phylogeographically distinct populations of *drosophila montana*. Evolution, 61, 1481–1488. doi: 10.1111/j.1558-5646.2007.00125.x

Kofler R., & Schlötterer, C. (2012). GOwinda: Unbiased analysis of gene set enrichment for genome-wide association studies. Bioinformatics. 28, 2084–2085. doi: 10.1093/bioinformatics/bts315.

Kolaczkowski, B., Kern, A. D., Holloway, A. K., & Begun, D. J. (2011). Genomic differentiation between temperate and tropical Australian populations of *Drosophila melanogaster*. Genetics, 187, 245–260. doi: 10.1534/genetics.110.123059

Lakovaara, S. (1969). Malt as a culture medium for *Drosophila* species. Drosophila Information Service, 44, 128.

Lankinen, P., Tyukmaeva, V. I., & Hoikkala, A. (2013). Northern *Drosophila montana* flies show variation both within and between cline populations in the critical day length evoking reproductive diapause. Journal of Insect Physiology, 59, 745–751. doi: 10.1016/j.jinsphys.2013.05.006

Lê, S., Josse, J., & Husson, F. (2008). FactoMineR: An R package for multivariate analysis. Journal of Statistical Software, 25, 1–18. doi: 10.18637/jss.v025.i01

Li, H. (2013). Aligning sequence reads, clone sequences and assembly contigs with BWA-MEM. arXiv:1303.3997v1 [q-bio.GN] Retrieved from: https://arxiv.org/abs/1303.3997

Li, H., Handsaker, B., Wysoker, A., Fennell, T., Ruan, J., Homer, N., … The 1000 Genome Project Data Processing Subgroup. (2009). The Sequence Alignment/Map format and SAMtools. Bioinformatics, 25, 2078–2079. doi: 10.1093/bioinformatics/btp352

Lohse, K., Clarke, M., Ritchie, M. G., & Etges, W. J. (2015). Genome-wide tests for introgression between cactophilic Drosophila implicate a role of inversions during speciation. Evolution, 69, 1178–1190. doi: 10.1111/evo.12650

Machado, H. E., Bergland, A. O., O’Brien, K., Behrman, E. L., Schmidt, P. S., & Petrov, D. (2015). Comparative population genomics of latitudinal variation in *Drosophila simulans* and *Drosophila melanogaster*. Molecular Ecology, 25, 723–740. doi: 10.1111/mec.13446

MacMillan, H. A., Williams, C. M., Staples, J. F., & Sinclair, B. J. (2012). Reestablishment of ion homeostasis during chill-coma recovery in the cricket *Gryllus pennsylvanicus*. Proceedings of the National Academy of Sciences of the United States of America, 109, 20750–20755. doi: 10.1073/pnas.1212788109.

Maston, G.A., Evans, S.K. & Green, M.R. 2006. Transcriptional regulatory elements in the human genome. Annual Review of Genomics and Human Genetics 7, 29–59. doi: 10.1146/annurev.genom.7.080505.115623

McKenna, A., Hanna, M., Banks, E., Sivachenko, A., Cibulskis, K., Kernytsky, A., … DePristo, M. (2010). The Genome Analysis Toolkit: A MapReduce framework for analyzing next-generation DNA sequencing data. Genome Research, 20, 1297–1303. doi:10.1101/gr.107524.110.

Mirol, P.M., Schäfer, M.A., Orsing, L., Routtu, J., Schlötterer, C., Hoikkala, A. & Butlin, R.K. (2007). Phylogeographic patterns in Drosophila montana. Molecular Ecology 16, 1085–1097. doi: 10.1111/j.1365-294X.2006.03215.x

Morales-Hojas, R., Päällysaho, S., Vieira, C. P., Hoikkala, A., & Vieira, J. (2007). Comparative polytene chromosome maps of *D. montana* and *D. virilis*. Chromosoma, 116, 21–7.doi: 10.1007/s00412-006-0075-3

Mylonakis, E., Podsiadlowski, L., Muhammed, M., & Vilcinskas, A. (2016). Diversity, evolution and medical applications of insect antimicrobial peptides. Philosophical Transactions of the Royal Society B: Biological Sciences, 371, 20150290. doi: 10.1098/rstb.2015.0290

Noor, M. A. F., & Bennett, S. M. (2009). Islands of speciation or mirages in the desert? Examining the role of restricted recombination in maintaining species. Heredity, 103, 439–444. doi: 10.1038/hdy.2009.151

Parker, D. J., Vesala, L., Ritchie, M. G., Laiho, A., Hoikkala, A., & Kankare, M. (2015). How consistent are the transcriptome changes associated with cold acclimation in two species of the *Drosophila virilis* group? Heredity, 115, 13–21. doi: 10.1038/hdy.2015.6

Parker, D. J., Ritchie, M. G., & Kankare, M. (2016). Preparing for Winter: The transcriptomic response associated with different day lengths in *Drosophila montana*. G3: Genes, Genomes, Genetics, 6, 1373–1381. doi: 10.1534/g3.116.027870

Parker, D. J., Wiberg, R. A. W., Trivedi, U., Tyukmaeva, V. I., Gharbi, K., Butlin, R. K., … Ritchie, M. G. (2018). Inter- and intra-specific genomic divergence in *Drosophila montana* shows evidence for cold adaptation. Genome Biology and Evolution, 10, 2086–2101. doi:10.1093/gbe/evy147

Parker D.J., Envall T., Ritchie M.G. & Kankare, M. (2021). Sex-specific responses to cold in a very cold-tolerant, northern *Drosophila* species. Heredty. https://doi.org/10.1038/s41437-020-00398-2

Patterson J.T. & Stone W.S. (1952). Evolution in the Genus Drosophila. Macmillan, N.Y

Pennacchio, L.A., Bickmore, W., Dean, A., Nobrega, M.A. & Bejerano, G. 2013. Enhancers: five essential questions. Nature Reviews Genetics. 14 288–295. doi: 10.1038/nrg3458

Plummer, M., Best, N., Cowles, K., & Vines, K. (2006). CODA: Convergence diagnosis and output analysis for MCMC. R News, 6, 7–11.

Paolucci, S., Salis, L., Vermeulen, C. J., Beukeboom, L. W., & van de Zande, L. (2016). QTL analysis of the photoperiodic response and clinal distribution of period alleles in *Nasonia vitripennis*. Molecular Ecology, 25, 4805–4817. doi: 10.1111/mec.13802

R Development Core Team. (2020). R: A language and environment for statistical computing. Retrieved from http://www.r-project.org

Ravinet, M., Faria, R., Butlin, R. K., Galindo, J., Bierne, N., Rafajlovic, M., …Westram, A. M. (2017). Interpreting the genomic landscape of speciation: A road map for finding barriers to gene flow. Journal of Evolutionary Biology, 30, 1450–1477. doi: 10.1111/jeb.13047

Routtu, J., Mazzi, D., van der Linde, K., Mirol, P., Butlin, R., & Ritchie, M. G. (2007). The extent of variation in male song, wing and genital characters among allopatric *Drosophila montana* populations. Journal of Evolutionary Biology, 20, 1591–1601. doi: 10.1111/j.1420-9101.2007.01323.x

Scheffzek, K., & Welti, S. (2012). Pleckstrin homology (PH) like domains – versatile modules in protein–protein interaction platforms. FEBS Letters, 586, 2662–2673. doi: 10.1016/j.febslet.2012.06.006

Schlötterer, C., Tobler, R., Kofler, R., & Nolte, V. (2014). Sequencing pools of individuals — mining genome-wide polymorphism data without big funding. Nature Reviews Genetics, 15, 749–763. doi: 10.1038/nrg3803

Sinclair, B.J., Ferguson, L. V., Salehipour-shirazi, G., & MacMillan, H. A. (2013). Cross-tolerance and cross-talk in the cold: Relating low temperatures to desiccation and immune stress in insects. Integrative and Comparative Biology, 53, 545–556. doi: 10.1093/icb/ict004

Sinclair, B. J., Nelson, S., Nilson, T. L., Roberts, S. P., & Gibbs, A. G. (2007). The effect of selection for desiccation resistance on cold tolerance of *Drosophila melanogaster*. Physiological Entomology, 32, 322–327. doi: 10.1111/j.1365-3032.2007.00585.x

Sinensky, M. (1974). Homeoviscous adaptation–A homeostatic process that regulates the viscosity of membrane lipids in Escherichia coli. Proceedings of the National Academy of Sciences of the United States of America, 71, 522–525. doi: 10.1073/pnas.71.2.522

Stanziano, J. R., Sové, R. J., Rundle, H. D., & Sinclair, B. J. (2015). Rapid desiccation hardening changes the cuticular hydrocarbon profile of *Drosophila melanogaster*. Comparative Biochemistry and Physiology, 180, 38–42. doi: 10.1016/j.cbpa.2014

Stone, W. S., Guest, W. C., & Wilson, F. D. (1960). The evolutionary implications of the cytological polymorphism and phylogeny of the *virilis* group of Drosophila. Proceedings of the National Academy of Sciences of the United States of America, 46, 350–361. doi: 10.1073/pnas.46.3.350

Takahashi, Y. (2015). Mechanisms and tests for geographic clines in genetic polymorphisms. Population Ecology. 57, 355–362. https://doi.org/10.1007/s10144-014-0474-x

Throckmorton, L. H. (1969). The *virilis* Species Group. The Genetics and Biology of Drosophila, 3, (pp. 227–296).

Thurmond, J., Goodman, J. L., Strelets, V. B., Attrill, H., Gramates, L. S., Marygold, S. J., … The FlyBase Consortium. (2019). FlyBase 2.0: The next generation. Nucleic Acids Research, 47, 759–765. doi: 10.1093/nar/gky1003

Tyukmaeva, V. I., Salminen, T. S., Kankare, M., Knott, E. & Hoikkala, A. (2011). Adaptation to a seasonally varying environment: A strong latitudinal cline in reproductive diapause combined with high gene flow in *Drosophila montana*. Ecology and Evolution, 1, 160–168. doi: 10.1002/ece3.14

Tyukmaeva, V. I., Lankinen, P., Kinnunen, J., Kauranen, H., & Hoikkala, A. (2020). Latitudinal clines in the timing and temperature-sensitivity of photoperiodic reproductive diapause in *Drosophila montana*. Ecography, 43, 1–10. doi: 10.1111/ecog.04892

Vesala, L., & Hoikkala, A. (2011). Effects of photoperiodically induced reproductive diapause and cold hardening on the cold tolerance of Drosophila montana. Journal of Insect Physiology, 57, 46–51. doi: 10.1016/j.jinsphys.2010.09.007

Vesala, L., Salminen, T. S., Kostal, V., Zahradnickova, H., & Hoikkala, A. (2012). *Myo*-inositol as a main metabolite in overwintering flies: Seasonal metabolomic profiles and cold stress tolerance in a northern drosophilid fly. Journal of Experimental Biology, 215, 2891–2897. doi:10.1242/jeb.069948

Vesala, L., Salminen, T. S., Laiho, A., Hoikkala, A., & Kankare M. (2012). Cold tolerance and cold-induced modulation of gene expression in two Drosophila virilis group species with different distributions. Insect Molecular Biology, 21, 107–118. doi: 10.1111/j.1365-2583.2011.01119.x

Vigoder, F. M., Parker, D. J., Cook, N., Tournière, O., Sneddon, T., & Ritchie, M. G. (2016). Inducing Cold-Sensitivity in the Frigophilic Fly *Drosophila montana* by RNAi. PLoS ONE, 11, 1–9. doi: 10.1371/journal.pone.0165724

Vines, T. H., Dalziel, A. C., Albert, A.Y. K., Veen, T., Schulte, P. M., & Schluter, D. (2016). Cline coupling and uncoupling in a stickleback hybrid zone. Evolution. 70, 1023–1038. doi: 10.1111/evo.12917

de Villemereuil, P., & Gaggiotti, O. E. (2015). A new FST-based method to uncover local adaptation using environmental variables. Methods in Ecology and Evolution, 6, 1248–1258. doi: 10.1111/2041-210X.12418.

Werner, T., Koshikawa, S., Williams, T.M. & Carroll, S.B. (2010). Generation of a novel wing colour pattern by the Wingless morphogen. Nature. 464, 1143–1148. doi: 10.1038/nature08896

Wilder, S. M., Mayntz, D., Toft, S., Rypstra, A. L., Pilati, A., & Vanni, M. J. (2010). Intraspecific variation in prey quality: A comparison of nutrient presence in prey and nutrient extraction by predators. Oikos, 119, 350–358. doi: 10.1111/j.1600-0706.2009.17819.x

Williams, C. M., McCue, M. D., Sunny, N. E., Szejner-Sigal, A., Morgan, T. J., Allison, D. B., & Hahn, D. A. (2016). Cold adaptation increases rates of nutrient flow and metabolic plasticity during cold exposure in *Drosophila melanogaster*. Proceedings of the Royal Society B: Biological Sciences, 283, 20161317. doi: 10.1098/rspb.2016.1317

Wolf, J. B. W., & Ellegren, H. (2016). Making sense of genomic islands of differentiation in light of speciation. Nature Reviews Genetics, 18, 87–100. doi: 10.1038/nrg.2016.133

Wood, S.N. (2004) Stable and efficient multiple smoothing parameter estimation for generalized additive models. Journal of the American Statistical Association. 99, 673–686. doi: 10.1198/016214504000000980

