## Supplementary Materials for "Cold adaptation drives population genomic divergence in the ecological specialist, *Drosophila montana*"

Wiberg et al.

**Pseudocode bioinformatic pipeline with model commands:**

### 1. Trimming

```
java -jar Trimmomatic.jar PE -phred33 *_R1_001.fastq.gz *_R2_001.fastq.gz *_tqc_R1_pe.fq.gz *_ftqc_R1_se.fq.gz *_tqc_R2_pe.fq.gz *_ftqc_R2_se.fq.gz ILLUMINACLIP:TruSeq3-PE.fa:2:30:10:8:TRUE CROP:145 LEADING:20 TRAILING:20 SLIDINGWINDOW:5:20 MINLEN:100;
```

### 2. Mapping

```
bwa mem -t 5  D_montana_genome_freeze_v1.4.fa *_R1_pe.fq.gz *_R2_pe.fq.gz | samtools view -Sb -q 30 -f 0x02 - > *.bam

samtools rmdup *_srt.bam *_srt_rmdup.bam

java -Xmx2g -jar $gatk -T RealignerTargetCreator -R D_montana_genome_freeze_v1.4.fa -I *_srt_rmdup_srt_rdgrp.bam -o *_srt_rmdup_srt_rdgrp.intervals

java -Xmx4g -jar $gatk -T IndelRealigner -R D_montana_genome_freeze_v1.4.fa -I *_srt_rmdup_srt_rdgrp.bam -targetIntervals *_srt_rmdup_srt_rdgrp.intervals -o *_srt_rmdup_srt_rdgrp_indraln.bam
```

### 3. Subsample Seward sample

```
XX=94100000

cat <(samtools view -H Seward_srt.bam) <(samtools view Seward_srt.bam | shuf -n ${XX}) > Seward_subs.sam

samtools view -bS Seward_subs.sam -o Seward_subs.bam
```

### 4. SNP calling

```
samtools mpileup --max-depth 1000000 --skip-indels --min-MQ 20 --min-BQ 15 --fasta-ref
D_montana_genome_freeze_v1.4.fa Oulanka.bam Korpilahti.bam Ashford.bam Seward.bam
Crested_Butte.bamTerrace.bam > montana_clines.mpileup
```

```
PoolSNP.sh mpileup=montana_clines.mpileup output=montana_clines
reference=D_montana_genome_freeze_v1.4.fa
names=Oulanka,Korpilahti,Ashford,Seward,Crested_Butte,Terrace min-cov=37 max-cov=0.95
min-count=5 min-freq=0.001 miss-frac=0.01 base-quality=15 jobs=10
```

**5. BayeScEnv**

```
bayescenv *_all_bscanin.txt -burn 2000 -pilot 1000 -npb 5 -n 2000 -env *.txt -od *_bscanout
```

**6. Gowinda**

```
gowinda --output-file *_topsnps_gowinda.tab --gene-definition updownstream10000 --simulations
1000000 --mode gene --gene-set-file funcassociate_go_associations.txt  --snp-file  all_snps.tab --
candidate-snp-file *_candidate_snps_file.tab --annotation-file
D_montana_genome_freeze_v1.4_dmel.gtf
```

**Final models of population phenotypes**

```
gamm(
        Ctmin ~ s(latitude, bs="cr", k=5) + s(altitude, bs="cr", k=5) + sex ,
        random = list(exp_batch=~1),
        data=cold_pop_dat
)
```

```
gamm(
        CCRT ~ s(latitude, bs="cr", k=5),
        random = list(exp_batch=~1),
        data=cold_pop_dat
)
```

where the term *exp_batch* was the experimental batch random effect.

**Supplementary Tables, Figures and Legends**

**Table S1.** See supplementary .xls file

**Table S2.** Mapping statistics from merged .bam files. Statistics for Seward samples are given after downsampling of reads.

| **Sample** | **Number of reads** | **Median coverage** |
|---|---|---|
| Oulanka | 78,261,294 | 62x |
| Korpilahti | 88,285,498 | 71x |
| Ashford | 82,318,578 | 66x |
| Crested Butte | 97,443,044 | 80x |
| Seward | 92,743,585 | 76x |
| Terrace | 109,266,003 | 88x |

**Table S3.** Loadings (correlation coefficients) of the 55 bioclimatic variables on principal components (PCs) 1-4. Correlation coefficients > 0.9 are highlighted in bold.

| Variable | Variable Name | Dim.1 | Dim.2 | Dim.3 | Dim.4 |
|---|---|---|---|---|---|
| bio1 | Annual Mean Temperature | **0.95** | -0.26 | 0.15 | -0.12 |
| bio2 | Mean Diurnal Range<br>(Mean of monthly (max temp - min temp)) | -0.11 | -0.81 | -0.56 | 0 |
| bio3 | Isothermality<br>(BIO2/BIO7) (* 100) | 0.62 | -0.49 | -0.56 | -0.05 |
| bio4 | Temperature Seasonality<br>(standard deviation *100) | -0.9 | -0.17 | 0.2 | 0.29 |
| bio5 | Max Temperature of Warmest Month | 0.33 | **-0.92** | 0.01 | 0.13 |
| bio6 | Min Temperature of Coldest Month | **0.97** | 0.08 | 0.1 | -0.13 |
| bio7 | Temperature Annual Range<br>(BIO5-BIO6) | -0.8 | -0.53 | -0.1 | 0.19 |
| bio8 | Mean Temperature of Wettest Quarter | -0.33 | -0.07 | 0.71 | 0.21 |
| bio9 | Mean Temperature of Driest Quarter | 0.85 | -0.24 | -0.26 | -0.14 |
| bio10 | Mean Temperature of Warmest Quarter | 0.55 | -0.62 | 0.51 | 0.14 |
| bio11 | Mean Temperature of Coldest Quarter | **0.97** | -0.06 | 0 | -0.19 |
| bio12 | Annual Precipitation | 0.81 | 0.49 | -0.21 | 0.25 |
| bio13 | Precipitation of Wettest Month | 0.81 | 0.44 | -0.14 | 0.35 |
| bio14 | Precipitation of Driest Month | 0.29 | 0.66 | -0.2 | -0.56 |
| bio15 | Precipitation Seasonality<br>(Coefficient of Variation) | 0.55 | 0.22 | 0.28 | 0.63 |
| bio16 | Precipitation of Wettest Quarter | 0.81 | 0.43 | -0.16 | 0.36 |
| bio17 | Precipitation of Driest Quarter | 0.51 | 0.66 | -0.35 | -0.33 |
| bio18 | Precipitation of Warmest Quarter | -0.23 | **0.92** | 0.14 | -0.12 |
| bio19 | Precipitation of Coldest Quarter | 0.84 | 0.35 | -0.25 | 0.33 |
| tmin1 | Minimum Temperature – January | **0.97** | 0.08 | 0.11 | -0.13 |
| tmin2 | Minimum Temperature – February | **0.97** | 0.06 | 0.07 | -0.12 |
| tmin3 | Minimum Temperature – March | **0.97** | -0.01 | 0.18 | -0.13 |
| tmin4 | Minimum Temperature – April | 0.91 | -0.02 | 0.37 | -0.1 |
| tmin5 | Minimum Temperature – May | 0.71 | 0.02 | 0.68 | 0.02 |
| tmin6 | Minimum Temperature – June | 0.39 | 0.1 | 0.9 | 0.12 |
| tmin7 | Minimum Temperature – July | 0.36 | 0.08 | 0.91 | 0.06 |
| tmin8 | Minimum Temperature – August | 0.61 | 0.13 | 0.77 | 0 |
| tmin9 | Minimum Temperature – September | 0.81 | 0.19 | 0.55 | -0.04 |
| tmin10 | Minimum Temperature – October | 0.88 | 0.15 | 0.35 | -0.18 |
| tmin11 | Minimum Temperature – November | **0.93** | 0.05 | 0.23 | -0.2 |

| | | | | | |
|---|---|---|---|---|---|
| tmin12 | Minimum Temperature – December | **0.96** | 0.11 | 0.18 | -0.15 |
| tmax1 | Maximum Temperature – January | **0.91** | -0.21 | -0.15 | -0.25 |
| tmax2 | Maximum Temperature – February | **0.92** | -0.27 | -0.18 | -0.19 |
| tmax3 | Maximum Temperature – March | **0.91** | -0.38 | -0.04 | -0.12 |
| tmax4 | Maximum Temperature – April | 0.82 | -0.54 | 0.08 | -0.05 |
| tmax5 | Maximum Temperature – May | 0.58 | -0.73 | 0.25 | 0.08 |
| tmax6 | Maximum Temperature – June | 0.25 | -0.89 | 0.26 | 0.19 |
| tmax7 | Maximum Temperature – July | 0.31 | **-0.92** | 0.01 | 0.12 |
| tmax8 | Maximum Temperature – August | 0.5 | -0.84 | -0.09 | 0.07 |
| tmax9 | Maximum Temperature – September | 0.68 | -0.7 | -0.18 | 0.02 |
| tmax10 | Maximum Temperature – October | 0.77 | -0.57 | -0.23 | -0.14 |
| tmax11 | Maximum Temperature – November | 0.86 | -0.36 | -0.12 | -0.26 |
| tmax12 | Maximum Temperature – December | 0.91 | -0.19 | -0.09 | -0.28 |
| prec1 | Precipitation – January | 0.83 | 0.33 | -0.26 | 0.35 |
| prec2 | Precipitation – February | 0.83 | 0.35 | -0.27 | 0.32 |
| prec3 | Precipitation – March | 0.83 | 0.29 | -0.31 | 0.34 |
| prec4 | Precipitation – April | 0.85 | 0.33 | -0.3 | 0.22 |
| prec5 | Precipitation – May | 0.78 | 0.36 | -0.35 | 0.06 |
| prec6 | Precipitation – June | 0.22 | 0.78 | -0.14 | 0.11 |
| prec7 | Precipitation – July | -0.6 | 0.7 | 0.21 | -0.18 |
| prec8 | Precipitation – August | -0.35 | 0.82 | 0.25 | -0.15 |
| prec9 | Precipitation – September | 0.37 | 0.74 | 0 | -0.18 |
| prec10 | Precipitation – October | 0.72 | 0.57 | -0.02 | 0.01 |
| prec11 | Precipitation – November | 0.86 | 0.36 | -0.18 | 0.3 |
| prec12 | Precipitation – December | 0.85 | 0.35 | -0.22 | 0.31 |

**Table S4.** See supplementary .xls format spreadsheet (bscan top SNPs)

**Table S5.** See supplementary .xls format spreadsheet (DAVID results)

**Table S6.** See supplementary .xls format spreadsheet (GOwinda results)

**Table S7.** Previous studies to which the results from the current study are compared.

| Reference | Study | Geneset being compared |
|---|---|---|
| Kankare et al., 2010 | Gene expression change during diapause | Genes that significantly changed their expression in response to diapause |
| Parker et al., 2015 | Gene expression response to cold acclimation | Genes that were DE in either *D. montana* or in both *D. montana* and *D. virilis* in response to cold acclimation |
| Parker et al., 2016 | Gene expression response to different day lengths | Genes that responded in expression to day length in either diapausing or non-diapausing flies. |
| Kankare et al., 2016 | Gene expression differences in diapuasing and non-diapausing females | Genes that were either upregulated or downregulated in diapausing compared to non-diapausing females. |
| Parker et al., 2018 | *D. montana* genome: population differentiation among three populations. | Genes that were within 1kb of outlier SNPs in any of three pairwise Fst outlier scans. |
| Kauranen et al., 2019 | Quasinatural selection under shorter day lengths. | Genes within 10kb of a SNP that changed in allele frequency |

**Table S8.** See supplementary .xls format spreadsheet (genes/overlap between studies)

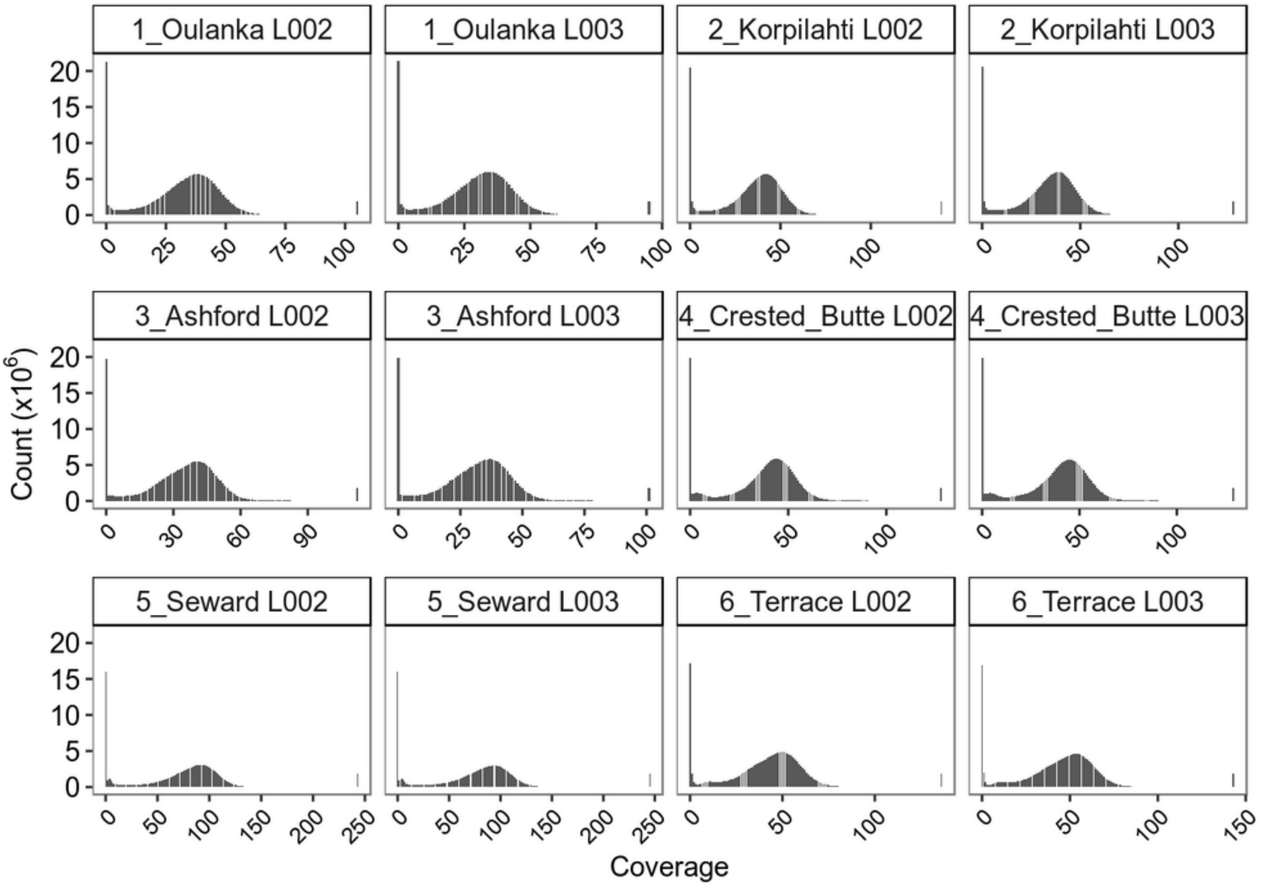


**Figure S1.** Coverage distributions of all samples before downsampling of Seward data. Note the different scales of the X-axes.

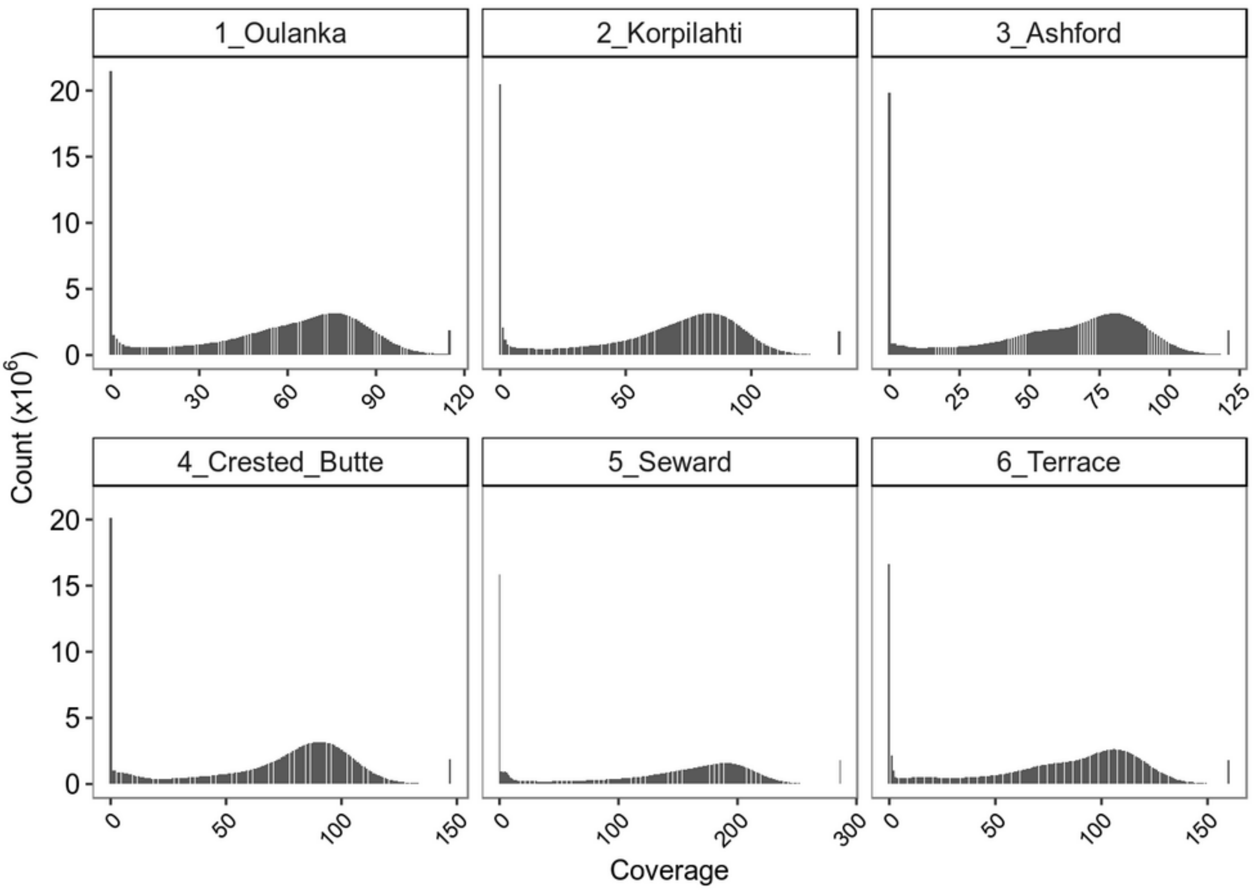


**Figure S2.** Coverage distributions of merged samples before downsampling of Seward data. Note the different scales of the X-axes.

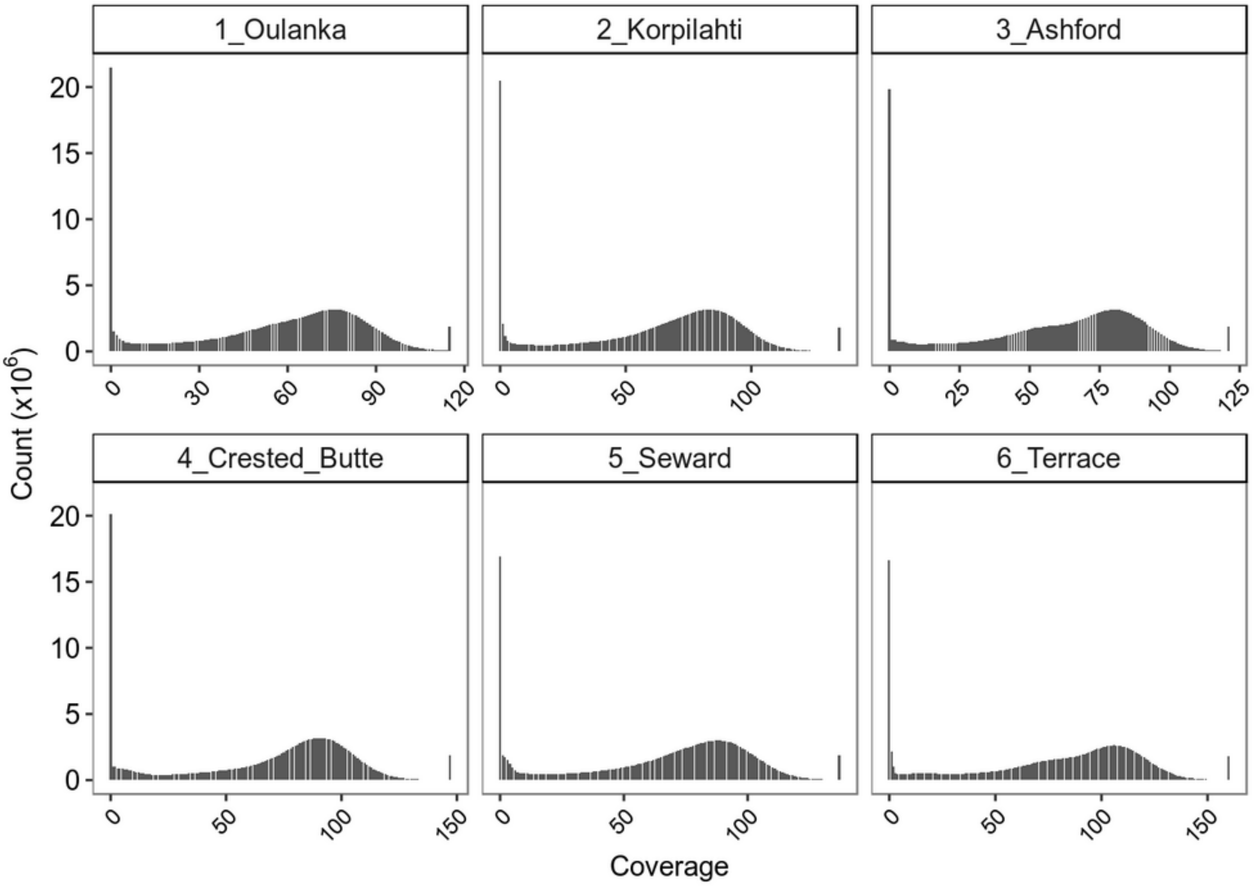


**Figure S3.** Coverage distributions of merged samples after downsampling of Seward data. Note the different scales of the X-axes.

**Figure S4.** Diagnostic gelman and trace plots for BayeScEnv MCMC chains with CTmin as the environmental variable. Analyses were split into **A)** autosomal SNPs and **B)** X chromosome SNPs. Gelman plots show the Gelman-Rubin shrink factors that summarise variation in parameter estimates across chains. Only data from the three chosen MCMC chains are included in these plots. Trace plots show parameter values across iterations of MCMC chains. These plots show data from all four chains that were run. See supplementary .pdf files.

**Figure S5.** Diagnostic plots for BayeScEnv MCMC chains with CCRT as the environmental variable. Analyses were split into **A)** autosomal SNPs and **B)** X chromosome SNPs. Gelman plots show the Gelman-Rubin shrink factors that summarise variation in parameter estimates across chains. Only data from the three chosen MCMC chains are included in these plots. Trace plots show parameter values across iterations of MCMC chains. These plots show data from all four chains that were run. See supplementary .pdf files.

**Figure S6.** Diagnostic plots for BayeScEnv MCMC chains with PC1 of the bioclimatic PCA as the environmental variable. Analyses were split into **A)** autosomal SNPs and **B)** X chromosome SNPs. Gelman plots show the Gelman-Rubin shrink factors that summarise variation in parameter estimates across chains. Only data from the three chosen MCMC chains are included in these plots. Trace plots show parameter values across iterations of MCMC chains. These plots show data from all four chains that were run. See supplementary .pdf files.

**Figure S7.** Diagnostic plots for BayeScEnv MCMC chains with PC2 of the bioclimatic PCA as the environmental variable. Analyses were split into **A)** autosomal SNPs and **B)** X chromosome SNPs. Gelman plots show the Gelman-Rubin shrink factors that summarise variation in parameter estimates across chains. Only data from the three chosen MCMC chains are included in these plots. Trace plots show parameter values across iterations of MCMC chains. These plots show data from all four chains that were run. See supplementary .pdf files.

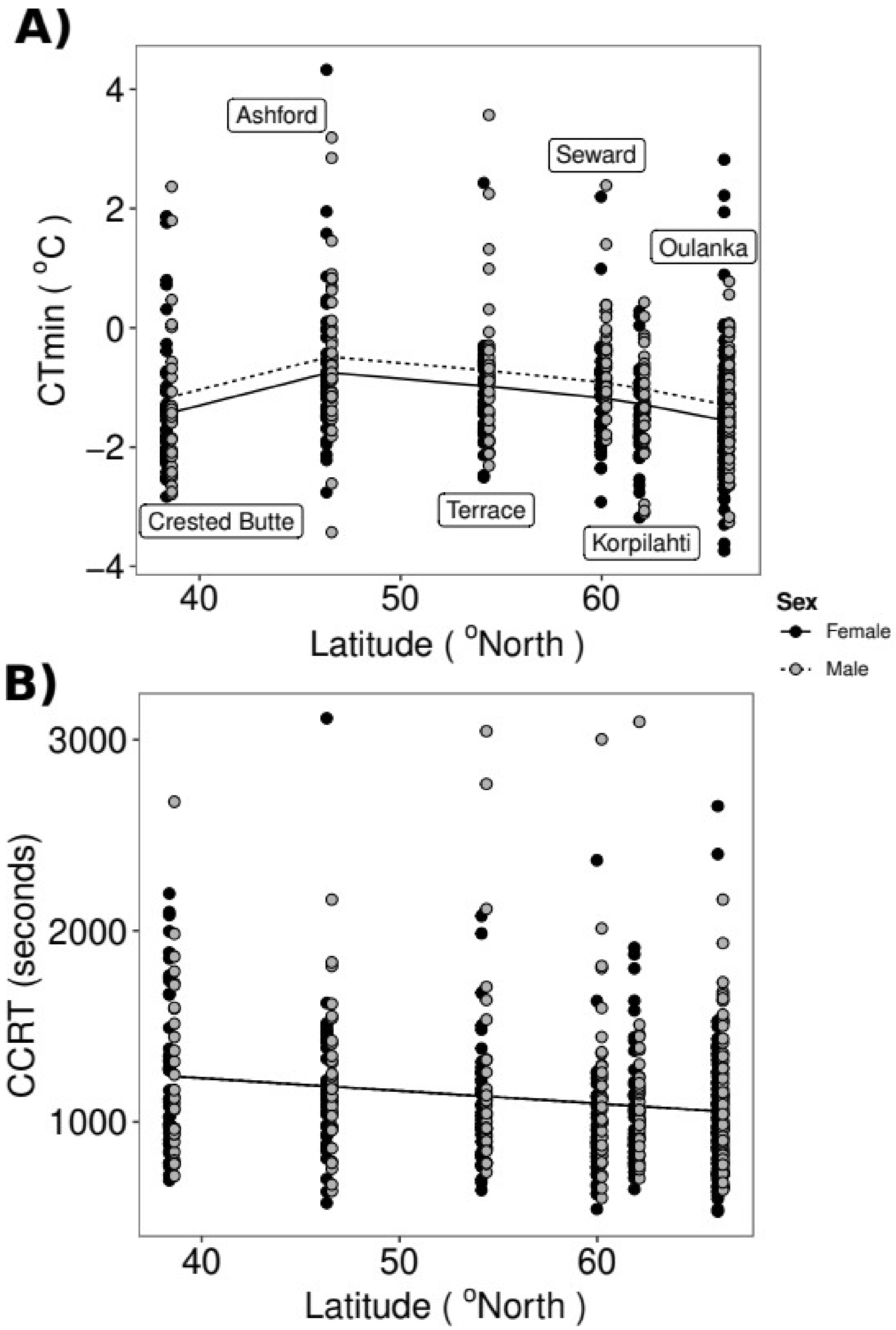


**Figure S8.** Full data for population phenotypes. **A)** and **B)** show the variation in CTmin and CCRT across populations and latitude, respectively. Solid and dashed lines show the predicted values from the partial effect of latitude from the best model (see Results) for males and females, respectively. In **B**, although the points are plotted separately for males and females, the best model only included latitude as a covariate.

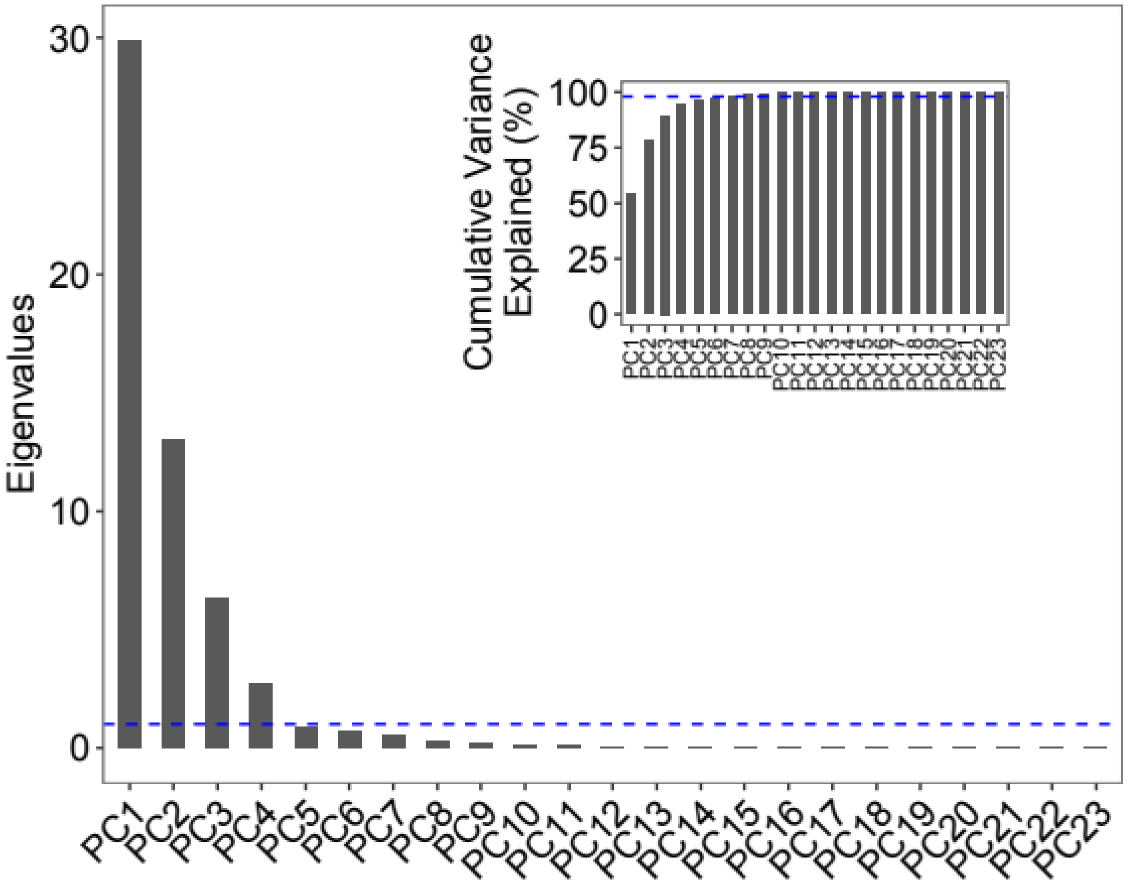


**Figure S9.** "Scree plot" of principal components 1-23 from a Principle Components Analysis of the 55 bioclimatic variables. Main plot shows the eigenvalues of the principal components, the horizontal dashed line shows the eigenvalue threshold of 1. The inset figure shows the cumulative amount of variance explained by the principal components. The horizontal dashed line gives 98% of the cumulative variance.

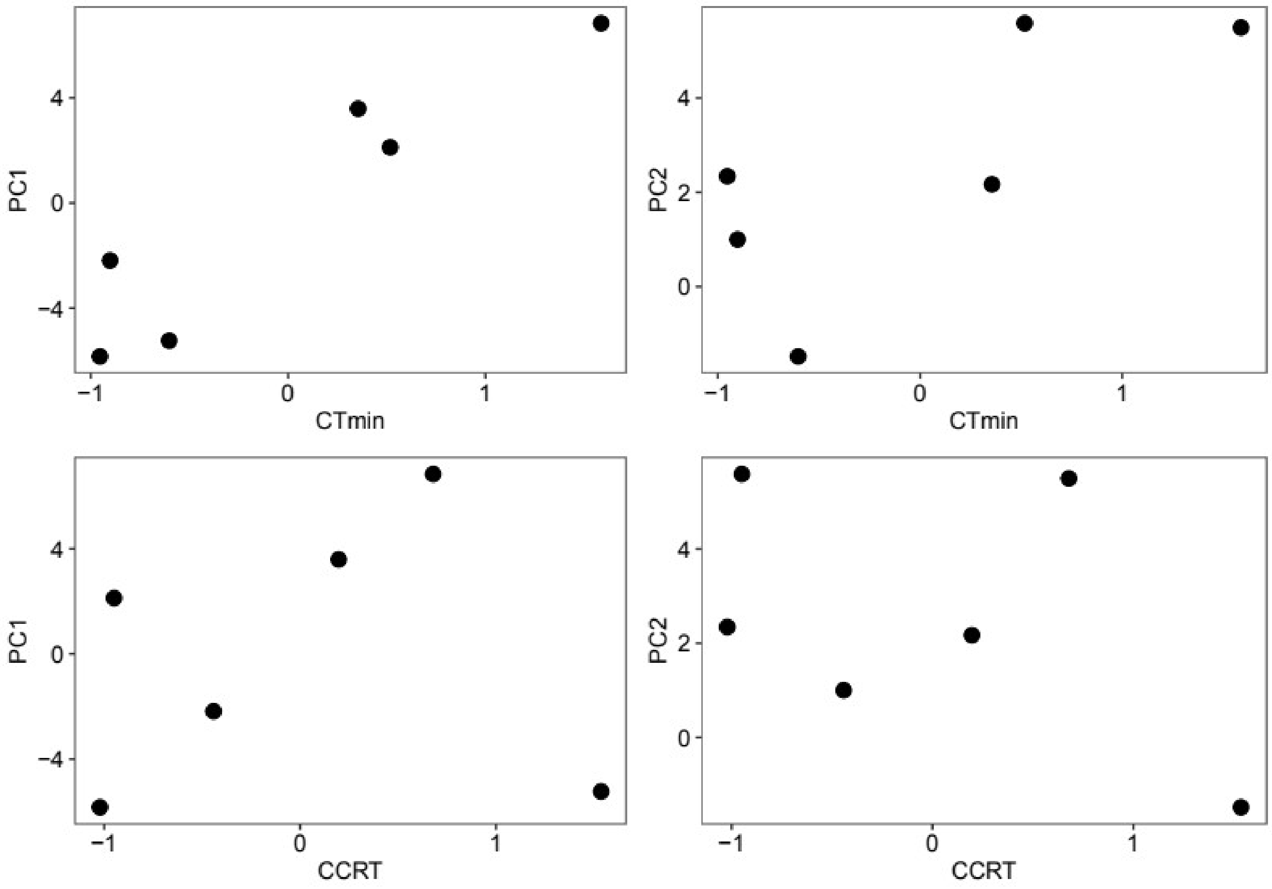


**Figure S10.** Scatter plots of the relationship between the population phenotypes (CTmin and CCRT) and the two chosen principal components (PC1 and PC2). PC values and phenotype values have been z-transformed.

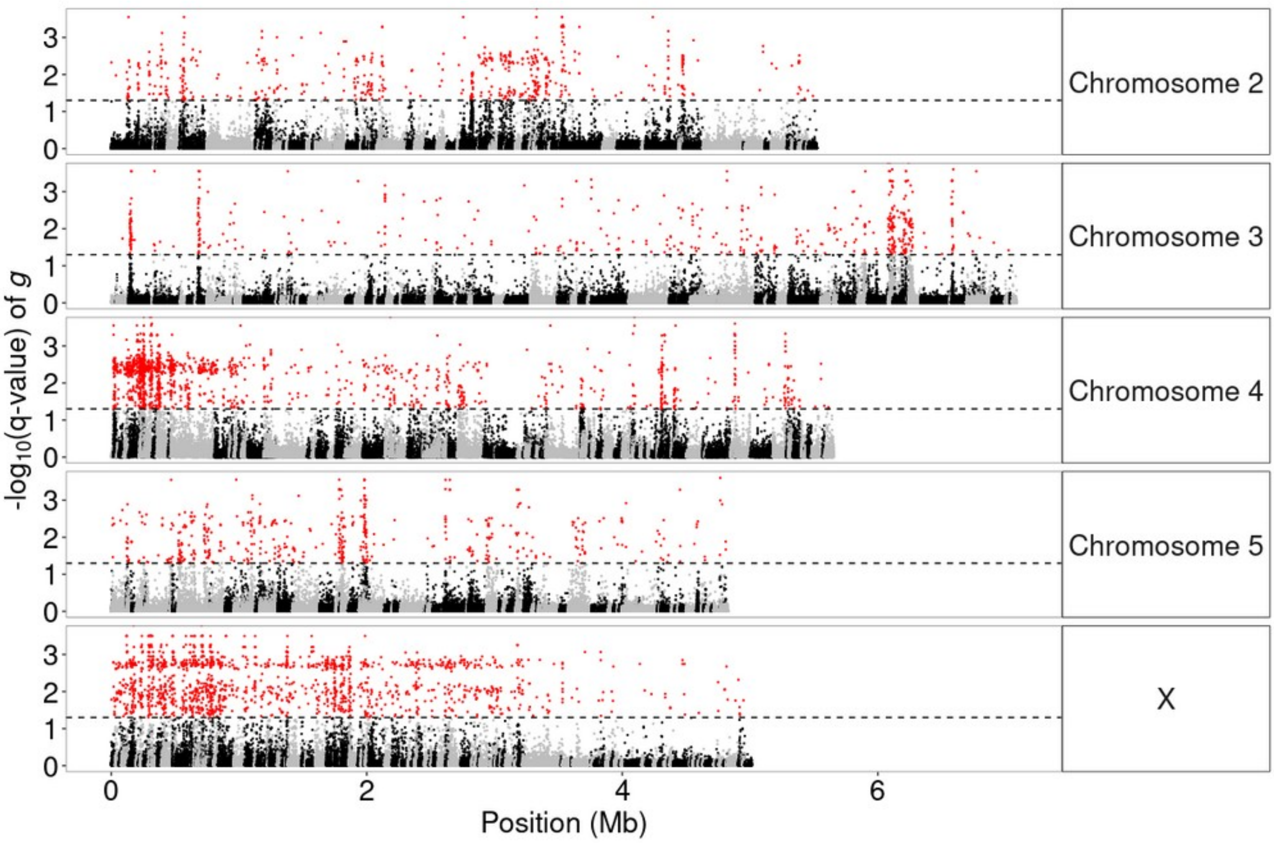


**Figure S11.** Manhattan plot of the q-values from BayeScEnv for a test of genetic differentiation with PC1. Red points show those SNPs with q-values < 0.05. The horizontal dashed line gives the FDR threshold used. Alternating black and grey points along chromosomes denote different scaffolds. Note that scaffolds can be place on chromosomes in this genome assembly but are not yet ordered.

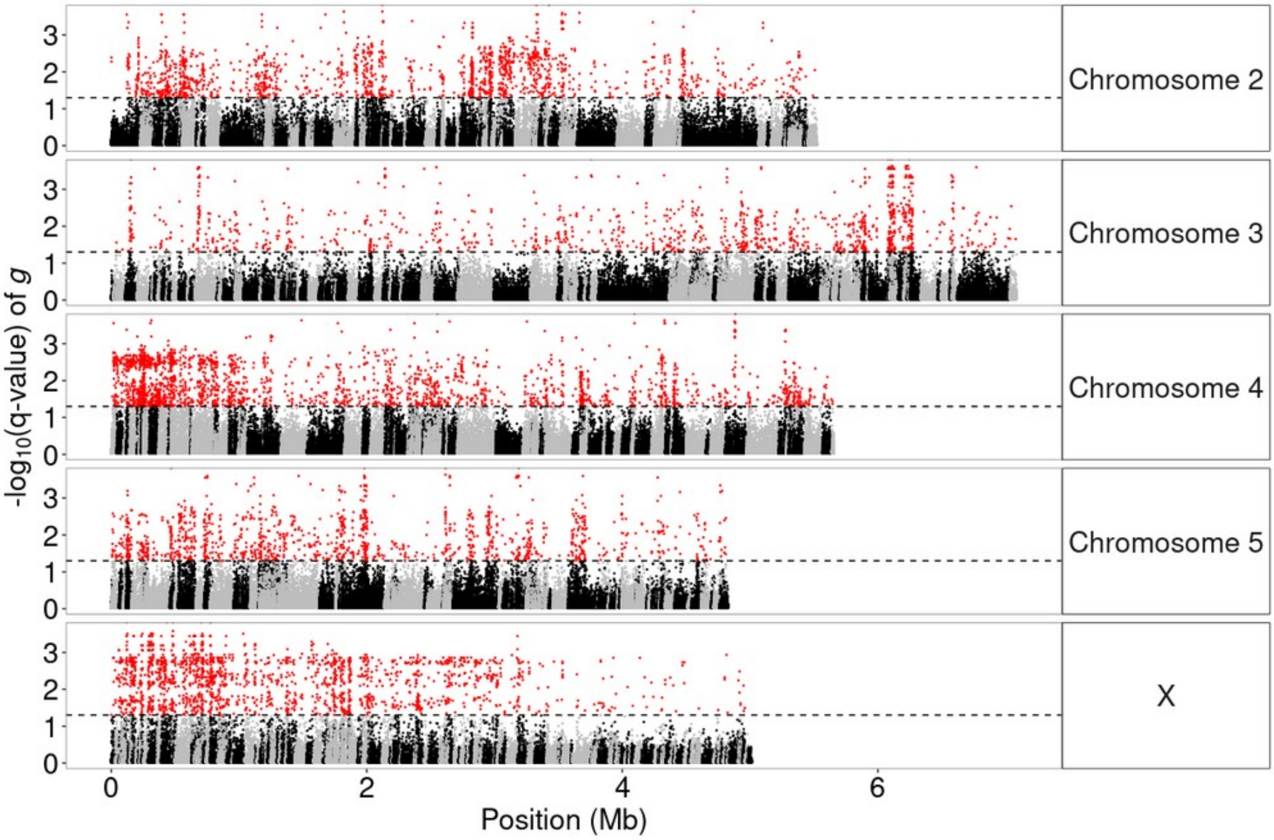


**Figure S12.** Manhattan plot of the q-values from BayeScEnv for a test of genetic differentiation with PC2. Red points show those SNPs with q-values < 0.05. The horizontal dashed line gives the FDR threshold used. Alternating black and grey points along chromosomes denote different scaffolds. Note that scaffolds can be place on chromosomes in this genome assembly but are not yet ordered.

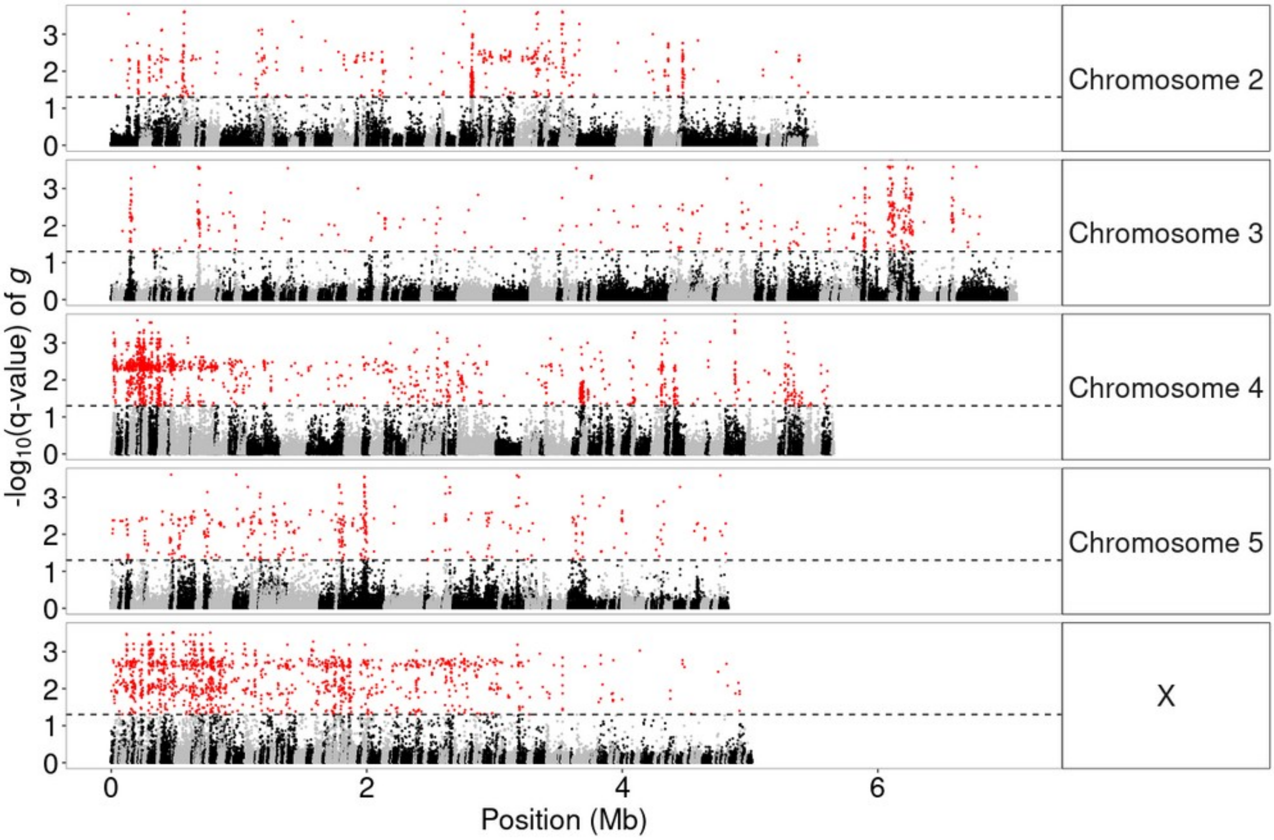


**Figure S13.** Manhattan plot of the q-values from BayeScEnv for a test of genetic differentiation with CTmin. Red points show those SNPs with q-values < 0.05. The horizontal dashed line gives the FDR threshold used. Alternating black and grey points along chromosomes denote different scaffolds. Note that scaffolds can be place on chromosomes in this genome assembly but are not yet ordered.

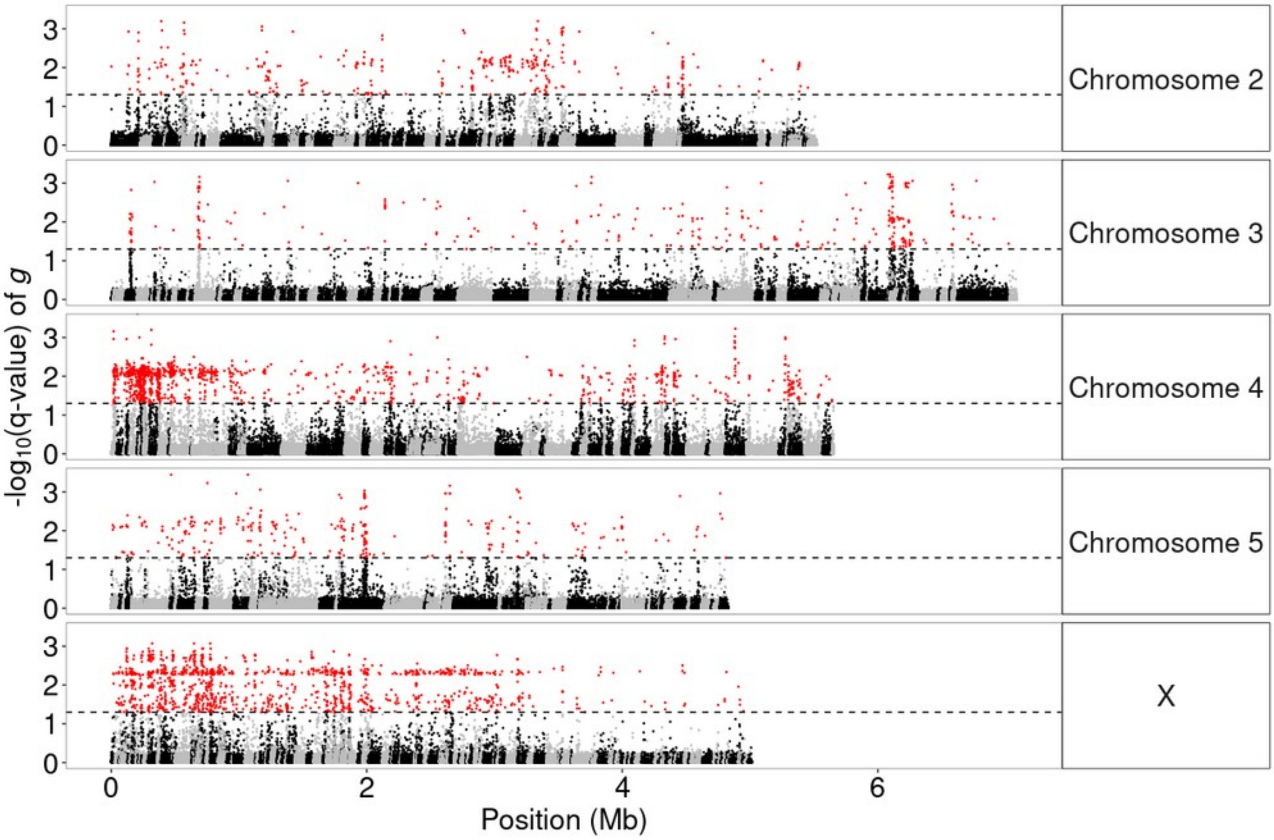


**Figure S14.** Manhattan plot of the q-values from BayeScEnv for a test of genetic differentiation with CCRT. Red points show those SNPs with q-values < 0.05. The horizontal dashed line gives the FDR threshold used. Alternating black and grey points along chromosomes denote different scaffolds. Note that scaffolds can be place on chromosomes in this genome assembly but are not yet ordered.

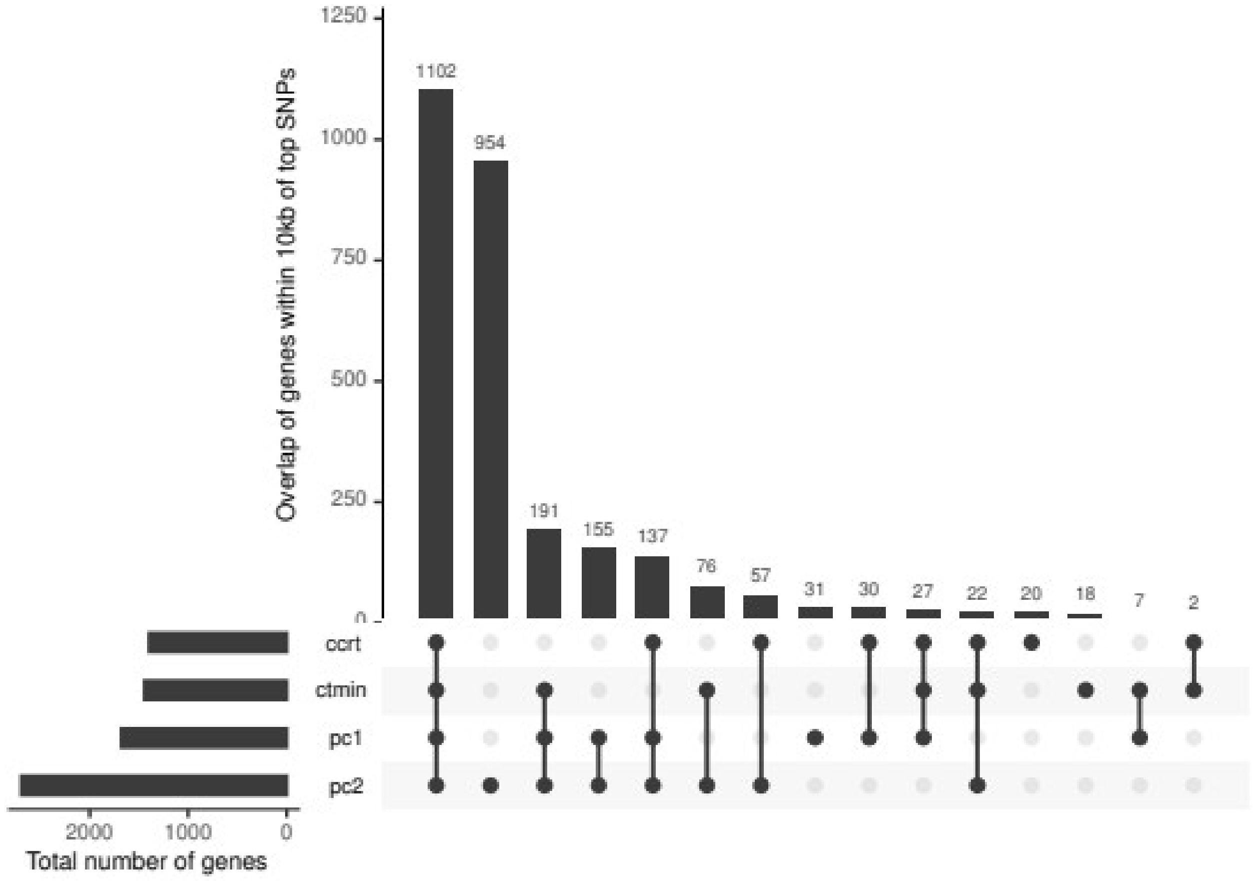


**Figure S15.** Overlap of genes within 10kb of top SNPs from BayeScEnv analyses for PC1, PC2 CTmin, and CCRT. The total number of genes near significant SNPs for each variable are given by the bars in the bottom left. The overlaps are depicted by points connected by lines along the *x*-axis and the height of the bars indicate the number of genes in common (intersection) to the variables connected by dots and lines.

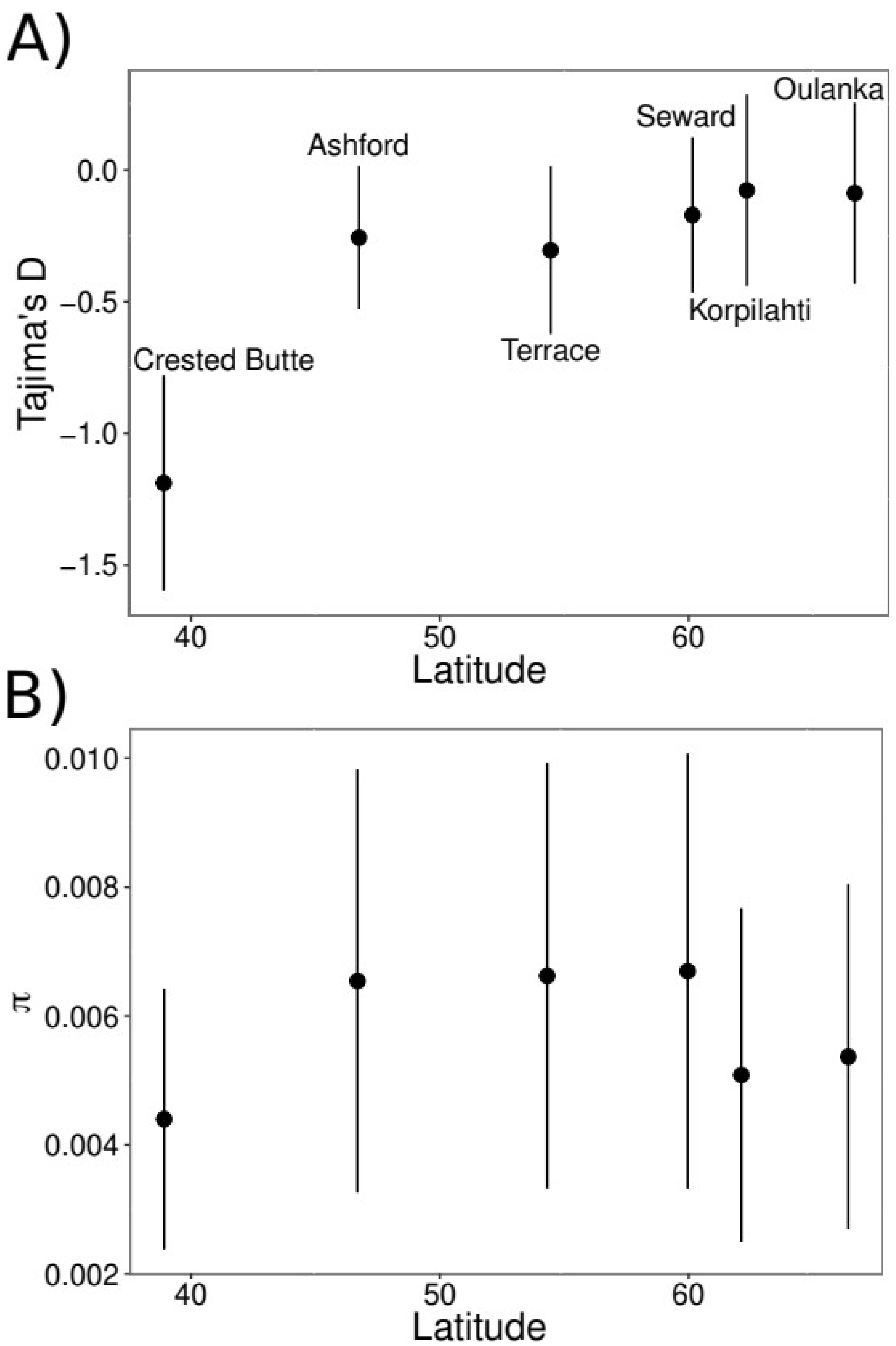


**Figure S16.** Mean and standard errors of **A)** nucleotide diversity (π) and **B)** Tajima's D as a function of latitude. Inset labels give the names of the corresponding populations. Statistics are computed from windows of 10kb with a step-size of 5kb.
