## Supplementary material for "Cold adaptation drives population genomic divergence in the ecological specialist, *Drosophila montana*": montana_clinalpops_analysis.R

```
##
# D. montana project clinal populations
# PCA/QBGLM data analysis
# Last Modified: Feb 2017
##

# Clean environment
rm(list = ls(all = TRUE))

#
setwd("~/Dropbox/st_andrews/montana_clines_MS/data/")
# Load libraries
require(raster)
library(FactoMineR)
library(ggplot2)
library(lme4)
library(nlme)
library(ggmap)
library(plyr)
library(cowplot)
source("~/projects/RData/RScripts/ggplot_theme.R")
se <- function(x) {sqrt(var(x, na.rm = TRUE))/sqrt(length(x))}
register_google(key = "AIzaSyAMRlZHArB72bBEvmSb5dqM0xwYRLc0eKU")
#----------------------------------------------#
# Get climate data for BayeScEnv and Quasi GLM
#----------------------------------------------#
# first load WC bio variables at the resolution of 2.5 deg
biod <- getData("worldclim", var="bio", res=2.5)
#BIO1 = Annual Mean Temperature
#BIO2 = Mean Diurnal Range (Mean of monthly (max temp - min temp)) ###
#BIO3 = Isothermality (BIO2/BIO7) (* 100) ###
#BIO4 = Temperature Seasonality (standard deviation *100) ###
#BIO5 = Max Temperature of Warmest Month
#BIO6 = Min Temperature of Coldest Month
#BIO7 = Temperature Annual Range (BIO5-BIO6)
#BIO8 = Mean Temperature of Wettest Quarter
#BIO9 = Mean Temperature of Driest Quarter
#BIO10 = Mean Temperature of Warmest Quarter
#BIO11 = Mean Temperature of Coldest Quarter ###
#BIO12 = Annual Precipitation
#BIO13 = Precipitation of Wettest Month
#BIO14 = Precipitation of Driest Month
#BIO15 = Precipitation Seasonality (Coefficient of Variation)
#BIO16 = Precipitation of Wettest Quarter
#BIO17 = Precipitation of Driest Quarter
#BIO18 = Precipitation of Warmest Quarter
#BIO19 = Precipitation of Coldest Quarter)
tmind <- getData("worldclim", var="tmin", res=2.5)
tmaxd <- getData("worldclim", var="tmax", res=2.5)
precd <- getData("worldclim", var="prec", res=2.5)


# read csv file with geographic coordinates
tableS1<-read.table("~/Dropbox/st_andrews/montana_clines_MS/Manuscript/V10_R2/supplemental_information/montana_pop_info_bioclimdat.tsv",
                    header=TRUE,sep="\t")

names(tableS1)
tableS1$Notes<-gsub("\n"," ",tableS1$Notes)
tableS1$Notes<-gsub("\r"," ",tableS1$Notes)
head(tableS1)
# extact for each coordinate bio clim variables
# extract(biovariable, c(long,lat))
bio<-extract(biod, tableS1[,c(5,4)])
tmin<-extract(tmind, tableS1[,c(5,4)])
tmax<-extract(tmaxd, tableS1[,c(5,4)])
precd<-extract(precd, tableS1[,c(5,4)])

# create a full dataset
tableS1<-cbind(tableS1,bio,tmin,tmax,precd)

# save into external file
#write.table(tableS1,
#            file="TableS1_montana_pop_info_bioclimdat.tsv",
#            sep="\t", row.names=FALSE ,quote=FALSE)

#######
# Run PCA for all populations
#######
#to get variance and mean for PCA
geo<-tableS1[,11:65]
names(geo)
rownames(geo)<-tableS1[,3]
# Run PCA
# Scale all variables
for(i in 1:ncol(geo)){
  geo[,i]<-scale(geo[,i],center=TRUE)
}
pca<-PCA(geo)
# Plots
pca$var$cor
pca_dat<-as.data.frame(pca$ind$coord)
pca_dat$loc<-rownames(pca_dat)
#Modify position of "Seward" label
pca_dat_labs<-pca_dat[,c(1,2,6)]
pca_dat_labs[1,2]<-pca_dat_labs[1,2]-0.3
pca_dat_labs$reg <- tableS1$Region
pca_dat_labs$PoolGen <- tableS1$PoolGen
head(pca_dat_labs)

ggplot()+
  # Other populations
  geom_point(data=pca_dat_labs[pca_dat_labs$PoolGen == 0 &
                                 pca_dat_labs$reg=="NoAm",],
             aes(Dim.1,Dim.2),shape = 24, fill = "red",size = 3)+
  geom_point(data=pca_dat_labs[pca_dat_labs$PoolGen == 0 &
                                 pca_dat_labs$reg=="Eur",],
             aes(Dim.1,Dim.2),shape = 22, fill = "red",size = 3)+
  
  # Sequenced populations
  geom_point(data=pca_dat_labs[pca_dat_labs$PoolGen == 1 &
                                 pca_dat_labs$reg=="NoAm",],
             aes(Dim.1,Dim.2),shape = 24, fill = "blue",size = 3)+
  geom_point(data=pca_dat_labs[pca_dat_labs$PoolGen == 1 &
                                 pca_dat_labs$reg=="Eur",],
             aes(Dim.1,Dim.2),shape = 22, fill = "blue",size = 3)+
  
  geom_text(data=pca_dat_labs[pca_dat_labs$PoolGen==1,],
             aes(Dim.1-0.5,Dim.2+0.45,label=loc),size = 4,colour="blue")+
  geom_text(data=pca_dat_labs[pca_dat_labs$PoolGen==0 &
                                pca_dat_labs$reg=="Eur",],
            aes(Dim.1-c(0,-0.5,-0.8,2.5),
                Dim.2+c(0.45,-0.45,0.45,0),
                label=loc),size = 4,colour="red")+
  geom_text(data=pca_dat_labs[pca_dat_labs$PoolGen==0 &
                                pca_dat_labs$reg=="NoAm",],
            aes(Dim.1-c(0,0,0,0,0,0,0,
                        0,0,-2.5,0,1,0,0),
                Dim.2+c(0.45,0.45,0.45,-0.45,0.45,0.45,0.45,
                        0.45,0.45,0.45,0.45,0.45,0.45,-0.45),
                label=loc),size = 4,colour="red")+
  
  scale_x_continuous(limits = c(-15,15))+
  xlab(paste("PC1 (",round(pca$eig[,2][1],2),"%)"))+
  ylab(paste("PC2 (",round(pca$eig[,2][2],2),"%)"))+
  geom_vline(xintercept = 0)+
  geom_hline(yintercept = 0)+
  my.theme+theme(axis.text = element_text(size = 14),
                 axis.title = element_text(size = 14))

# How much of PC1 and PC2 is covered by sequence populations, 

diff(range(pca_dat_labs$Dim.1[pca_dat_labs$PoolGen == 1]))/diff(range(pca_dat_labs$Dim.1))

diff(range(pca_dat_labs$Dim.2[pca_dat_labs$PoolGen == 1]))/diff(range(pca_dat_labs$Dim.2))

pca_dat_labs2<-pca_dat[,c(3,4,6)]
pca_dat_labs2[1,2]<-pca_dat_labs[1,2]-0.3
pca_dat_labs2$reg <- tableS1$Region
pca_dat_labs2$PoolGen <- tableS1$PoolGen

ggplot()+
  geom_point(data=pca_dat_labs2[pca_dat_labs2$PoolGen == 0 &
                                 pca_dat_labs$reg=="NoAm",],
             aes(Dim.3,Dim.4),shape = 24, fill = "red",size = 3)+
  geom_point(data=pca_dat_labs2[pca_dat_labs2$PoolGen == 0 &
                                 pca_dat_labs$reg=="Eur",],
             aes(Dim.3,Dim.4),shape = 22, fill = "red",size = 3)+
  
  geom_point(data=pca_dat_labs2[pca_dat_labs2$PoolGen == 1,],
             aes(Dim.3,Dim.4),shape = 21, fill = "blue",size = 3)+
  geom_label(data=pca_dat_labs2[pca_dat_labs2$PoolGen == 1,],
             aes(Dim.3+c(1,1.5,1,1,-1,1),
                 Dim.4+c(0.4,0.1,0.3,0.3,0.3,0.3),label=loc),
             size=4)+
  scale_x_continuous(limits = c(-10,10))+
  xlab(paste("PC3 (",round(pca$eig[,3][3],2),"%)"))+
  ylab(paste("PC4 (",round(pca$eig[,3][4],2),"%)"))+
  geom_vline(xintercept = 0)+
  geom_hline(yintercept = 0)+
  my.theme+theme(axis.text = element_text(size = 14),
                 axis.title = element_text(size = 14))

# Make a "Scree" plot
pca_eig_dat<-as.data.frame(pca$eig)
pca_eig_dat$comp<-rownames(pca_eig_dat)
pca_eig_dat$comp <- gsub("comp ","",pca_eig_dat$comp)
pca_eig_dat$comp <- as.numeric(pca_eig_dat$comp)
pca_eig_dat$comp<-factor(pca_eig_dat$comp,
                         labels=paste("PC",seq(1,23),sep=""))

plot1<-ggplot(data=pca_eig_dat)+
  geom_bar(aes(comp,eigenvalue),stat = "identity",width = 0.5)+
  geom_hline(yintercept = 1,linetype="dashed",colour="blue")+
  xlab("")+
  ylab("Eigenvalues")+
  my.theme+theme(axis.text = element_text(size = 14),
                 axis.text.x = element_text(angle=45,
                                            hjust=1,vjust=1),
                 axis.title = element_text(size = 14))

plot(plot1)
# First 4 vectors have eigenvalues > 1
plot1_ins<-ggplot(data=pca_eig_dat)+
  geom_bar(aes(comp,`cumulative percentage of variance`),
           stat = "identity",width = 0.5)+
  geom_hline(yintercept = 98,linetype="dashed",colour="blue")+
  xlab("")+
  ylab("Cumulative Variance\nExplained (%)")+
  my.theme+theme(axis.text = element_text(size = 14),
                 axis.text.x = element_text(size=8,angle=90,
                                            hjust=1,vjust=0.5),
                 axis.title = element_text(size = 14))
plot(plot1_ins)

ggdraw() +
  draw_plot(plot1, 0, 0, 1, 1) +
  draw_plot(plot1_ins, 0.4, 0.52, 0.5, 0.4)


# First 4 Principal Components have eigenvalues > 1
# Run PCA again keeping only the first 4 dimensions.
pca<-PCA(geo,ncp = 4)
pca$var$cor
cat(capture.output(pca$var$coord),
    file="~/Dropbox/st_andrews/montana_clines_MS/Manuscript/V10_R2/montana_clinalpops-GIS_PCA.rot",
    sep="")
pca$ind
pca$var$cor

# Describe the first 4 PCA components
dimdesc(pca,axes = c(1,2,3,4))
# pca$ind$coord
# Ztransform the PCA results
pca$ind$coord[,1]<-scale(x = pca$ind$coord[,1],center = TRUE,scale=TRUE)
pca$ind$coord[,2]<-scale(x = pca$ind$coord[,2],center = TRUE,scale=TRUE)
pca$ind$coord[,3]<-scale(x = pca$ind$coord[,3],center = TRUE,scale=TRUE)
pca$ind$coord[,4]<-scale(x = pca$ind$coord[,4],center = TRUE,scale=TRUE)
pca$ind$coord
plot(pca$ind$coord[,1],pca$ind$coord[,2])
# Bind the PCA results to the main data
tableS1[,c(1,3)]
colnames(pca$ind$coord)<-gsub("Dim.","PC",colnames(pca$ind$coord))
tableS1$PC1_scaled<-scale(x = pca$ind$coord[,1],center = TRUE,scale=TRUE)
tableS1$PC2_scaled<-scale(x = pca$ind$coord[,2],center = TRUE,scale=TRUE)
tableS1$PC1<-pca$ind$coord[,1]
tableS1$PC2<-pca$ind$coord[,2]

#write.table(TableS1,
#            file="~/Desktop/Data/montana_project/clinal_population_genomics/TableS1_montana_pop_info_bioclimdat.csv",
#            sep="\t", row.names=FALSE ,quote=FALSE)

# Read in table S1
#tableS1<-read.table("~/Projects/montana/clinal_populations/pop_data/TableS1_montana_pop_info_bioclimdat.csv",
#                     header=TRUE,sep=",")
head(tableS1)
pca$var
####
# Correlation between altitude and PC1 and PC2
####
tableS1[tableS1$LocationName=="Crested Butte",c(6,66,67)]
cor.test(tableS1$Alt_m,tableS1$PC1)
ggplot()+
  geom_point(data=tableS1,aes(Alt_m,PC1))+
  xlab("Altitude (m)")+
  ylab("PC1")+
  geom_label(data=tableS1[tableS1$LocationName=="Crested Butte",],
             aes(Alt_m-200,PC1+0.1,label=LocationName))+
  geom_label(aes(x=2500,y=1,label="cor = -0.398\np = 0.092"))+
  my.theme+theme(axis.title = element_text(size=10),
                 axis.text = element_text(size=10))

cor.test(tableS1$Alt_m,tableS1$PC2)
ggplot()+
  geom_point(data=tableS1,aes(Alt_m,PC2))+
  xlab("Altitude (m)")+
  ylab("PC2")+
  geom_label(data=tableS1[tableS1$LocationName=="Crested Butte",],
             aes(Alt_m-200,PC2+0.1,label=LocationName))+
  geom_label(aes(x=2500,y=1,label="cor = -0.578\np = 0.0096"))+
  my.theme+theme(axis.title = element_text(size=10),
                 axis.text = element_text(size=10))

# Plot latitude and longitude as a function of PC1 and PC2
head(tableS1)
cor.test(tableS1$Lat,tableS1$PC1,method = "spearman")
cor.test(tableS1$Alt_m,tableS1$PC1,method = "spearman")
pc1<-ggplot()+
  geom_point(data=tableS1,aes(Lat,PC1,colour=Alt_m),size=3)+
  xlab("Latitude")+
  ylab("PC1")+
  scale_colour_viridis_c("Altitude (m)",)+
  geom_label(data=tableS1[tableS1$LocationName=="Crested Butte",],
             aes(Lat+2.5,PC1+0.15,label=LocationName))+
#  geom_label(aes(x=65,y=1,label="cor = -0.517\np = 0.0097"))+
  my.theme+theme(axis.title = element_text(size=16),
                 axis.text = element_text(size=16))
pc1
cor.test(tableS1$Lat,tableS1$PC2,method = "spearman")
cor.test(tableS1$Alt_m,tableS1$PC2,method = "spearman")
pc2<-ggplot()+
  geom_point(data=tableS1,aes(Lat,PC2,colour=Alt_m),size=3)+
  xlab("Latitude")+
  ylab("PC2")+
  scale_color_viridis_c("Altitude (m)")+
  geom_label(data=tableS1[tableS1$LocationName=="Crested Butte",],
             aes(Lat+2.5,PC2+0.15,label=LocationName))+
#  geom_label(aes(x=65,y=1,label="cor = -0.517\np = 0.0097"))+
  my.theme+theme(axis.title = element_text(size=16),
                 axis.text = element_text(size=16))
pc2

plot_grid(pc1,pc2)
#######
# Run PCA analysis on only the 6 populations for pool seq
#######
tableS1_pool <- tableS1[tableS1$PoolGen == 1,] 

#to get variance and mean for PCA
geo<-tableS1_pool[,11:65]
rownames(geo)<-tableS1_pool[,3]
# Run PCA
pca<-PCA(geo)
# Plots
pca_dat<-as.data.frame(pca$ind$coord)
pca_dat$loc<-rownames(pca_dat)
#Modify position of "Seward" label
pca_dat_labs<-pca_dat[,c(1,2,6)]
pca_dat_labs[1,2]<-pca_dat_labs[1,2]-0.3
pca_dat_labs$reg <- tableS1_pool$Region
pca_dat_labs$PoolGen <- tableS1_pool$PoolGen

ggplot()+
  geom_point(data=pca_dat_labs,
             aes(Dim.1,Dim.2),size = 3)+
  geom_label(data=pca_dat_labs,
             aes(Dim.1-0.5,Dim.2+0.45,label=loc),size = 4)+
  scale_x_continuous(limits = c(-10,10))+
  xlab(paste("PC1 (",round(pca$eig$`percentage of variance`[1],2),"%)"))+
  ylab(paste("PC2 (",round(pca$eig$`percentage of variance`[2],2),"%)"))+
  geom_vline(xintercept = 0)+
  geom_hline(yintercept = 0)+
  my.theme+theme(axis.text = element_text(size = 14),
                 axis.title = element_text(size = 14))

pca_dat_labs2<-pca_dat[,c(3,4,6)]
pca_dat_labs2[1,2]<-pca_dat_labs[1,2]-0.3
pca_dat_labs2$reg <- tableS1_pool$Region
pca_dat_labs2$PoolGen <- tableS1_pool$PoolGen

ggplot()+
  geom_point(data=pca_dat_labs2,
             aes(Dim.3,Dim.4),size = 3)+
  geom_label(data=pca_dat_labs2,
             aes(Dim.3+0.1,Dim.4+0.15,label=loc),
             size=4)+
  scale_x_continuous(limits = c(-4,4))+
  xlab(paste("PC3 (",round(pca$eig$`percentage of variance`[3],2),"%)"))+
  ylab(paste("PC4 (",round(pca$eig$`percentage of variance`[4],2),"%)"))+
  geom_vline(xintercept = 0)+
  geom_hline(yintercept = 0)+
  my.theme+theme(axis.text = element_text(size = 14),
                 axis.title = element_text(size = 14))

colnames(pca$ind$coord)<-gsub("Dim.","PC",colnames(pca$ind$coord))
tableS1_pool$PC1_scaled<-scale(x = pca$ind$coord[,1],center = TRUE,scale=TRUE)
tableS1_pool$PC2_scaled<-scale(x = pca$ind$coord[,2],center = TRUE,scale=TRUE)
tableS1_pool$PC1<-pca$ind$coord[,1]
tableS1_pool$PC2<-pca$ind$coord[,2]


cor.test(tableS1_pool$Alt_m,tableS1_pool$PC1,method = "spearman")
cor.test(tableS1_pool$Lat,tableS1_pool$PC1,method = "spearman")
cor.test(tableS1_pool$Alt_m,tableS1_pool$PC2,method = "spearman")
cor.test(tableS1_pool$Lat,tableS1_pool$PC2,method = "spearman")

# Make a "Scree" plot
pca_eig_dat<-as.data.frame(pca$eig)
pca_eig_dat$comp<-rownames(pca_eig_dat)
pca_eig_dat$comp <- gsub("comp ","",pca_eig_dat$comp)
pca_eig_dat$comp <- as.numeric(pca_eig_dat$comp)

plot1<-ggplot(data=pca_eig_dat)+
  geom_bar(aes(comp,eigenvalue),stat = "identity",width = 0.5)+
  geom_hline(yintercept = 1,linetype="dashed",colour="blue")+
  xlab("")+
  ylab("Eigenvalues")+
  my.theme+theme(axis.text = element_text(size = 10),
                 axis.title = element_text(size = 10))
plot(plot1)
pca_eig_dat$comp<-factor(pca_eig_dat$comp,
                         labels=c("PC1","PC2","PC3","PC4","PC5"))
# First 4 vectors have eigenvalues > 1
plot1_ins<-ggplot(data=pca_eig_dat)+
  geom_bar(aes(comp,`cumulative percentage of variance`),
           stat = "identity",width = 0.5)+
#  geom_hline(yintercept = 80,linetype="dashed",colour="blue")+
  xlab("")+
  ylab("Cumulative Variance\nExplained (%)")+
  my.theme+theme(axis.text = element_text(size = 10),
                 axis.title = element_text(size = 10))
plot(plot1_ins)

# make plot1_ins an "inset" in plot1
# Specify position of plot2 (in percentages of plot1)
# This is in the top left and 25% width and 25% height
xleft   = 0.4
xright  = 0.95
ybottom = 0.55
ytop    = 0.95 

# Calculate position in plot1 coordinates
# Extract x and y values from plot1
l1 = ggplot_build(plot1)
x1 = l1$layout$panel_ranges[[1]]$x.range[1]
x2 = l1$layout$panel_ranges[[1]]$x.range[2]
y1 = l1$layout$panel_ranges[[1]]$y.range[1]
y2 = l1$layout$panel_ranges[[1]]$y.range[2]
xdif = x2-x1
ydif = y2-y1
xmin  = x1 + (xleft*xdif)
xmax  = x1 + (xright*xdif)
ymin  = y1 + (ybottom*ydif)
ymax  = y1 + (ytop*ydif) 

# Get plot2 and make grob
g2 = ggplotGrob(plot1_ins)
plot3 = plot1 + annotation_custom(grob = g2, 
                                  xmin=xmin, xmax=xmax, 
                                  ymin=ymin, ymax=ymax)
plot(plot3)


# First 4 Principal Components have eigenvalues > 1
# Run PCA again keeping only the first 4 dimensions.
pca<-PCA(geo,ncp = 4)
pca$var$cor
cat(capture.output(pca$var$coord),
    file="~/Dropbox/st_andrews/montana_clines_MS/Manuscript/V10_R2/montana_clinalpops-GIS_PCA_popsub.rot",
    sep="")
pca$ind
pca$var$cor
# Describe the first 4 PCA components
dimdesc(pca,axes = c(1,2,3,4))
# Save TableS1

# Load TableS1
tableS1<-read.table("~/Dropbox/st_andrews/montana_clines_MS/Manuscript/V10_R2/supplemental_information/montana_pop_info_bioclimdat.tsv",
                    header=TRUE,sep="\t")
head(tableS1)

# Load Veneras cold tolerance data
order<-c("Oulanka","Korpilahti",
         "Ashford","Seward","Crested Butte","Terrace")
cold_pop_dat<-read.table("~/Dropbox/st_andrews/montana_clines_MS/Manuscript/V10_R2/supplemental_information/cold_pop_cage_all_data_Axel.csv",header=TRUE,sep=" ")
cold_pop_dat$population<-as.character(cold_pop_dat$population)
cold_pop_dat$population[
  cold_pop_dat$population=="Colorado"]<-"Crested Butte"
cold_pop_dat$exp_batch<-factor(cold_pop_dat$exp_batch)
head(cold_pop_dat,n=20)

summary(cold_pop_dat)
# Remove incomplete records
cold_pop_dat<-na.omit(cold_pop_dat)

levels(cold_pop_dat$exp_batch)
levels(cold_pop_dat$strain)

tapply(cold_pop_dat$number,INDEX = list(cold_pop_dat$sex,cold_pop_dat$strain),length)

tapply(cold_pop_dat$number,INDEX = list(cold_pop_dat$exp_batch),length)
tapply(cold_pop_dat$exp_batch,INDEX = list(cold_pop_dat$population),function(x){length(unique(x))})
tapply(cold_pop_dat$number,INDEX = list(cold_pop_dat$sex,cold_pop_dat$exp_batch),length)
tapply(cold_pop_dat$population,INDEX = list(cold_pop_dat$sex,cold_pop_dat$exp_batch),function(x){length(unique(x))})

plot_dat<-ddply(cold_pop_dat,.variables = .(population,sex),summarise,
                mean_ctmin=mean(Ctmin,na.rm=TRUE),
                se_ctmin=se(Ctmin),
                lat=mean(latitude),
                alt=mean(altitude))
plot_dat
head(cold_pop_dat)
# Does batch have an effect?
anova(lm(Ctmin~population*sex+exp_batch,data=cold_pop_dat))
anova(lm(CCRT_totalsec~population*sex+exp_batch,data=cold_pop_dat))

# With latitude as the predictor instead of population
m<-lmer(Ctmin~altitude+latitude+sex+latitude:sex+(1|exp_batch),data=cold_pop_dat)
m1<-lm(Ctmin~altitude+latitude+sex+latitude:sex,data=cold_pop_dat)
anova(m,m1)
# Removing the random effect is not fine.
m1<-lmer(Ctmin~altitude+latitude+sex+(1|exp_batch),data=cold_pop_dat)
m2<-lmer(Ctmin~altitude+sex+(1|exp_batch),data=cold_pop_dat)
m3<-lmer(Ctmin~latitude+sex+(1|exp_batch),data=cold_pop_dat)
plot(m)
hist(resid(m))
anova(m,m1)
anova(m1,m2)
anova(m1,m3)
# m1 is the best
plot(m1)
hist(resid(m1))
str(cold_pop_dat)
summary(m1)

# Fit GAM (cubic spline). With a GAMM I can include the batch random effect
library(mgcv)
gam0<-gam(Ctmin~s(latitude,bs="cr",k=5)+s(altitude,bs="cr",k=5)+sex,data=cold_pop_dat)
summary(gam0)
gam1<-gamm(Ctmin~s(latitude,bs="cr",k=5)+s(altitude,bs="cr",k=5)+sex,data=cold_pop_dat,random=list(exp_batch=~1))
summary(gam1$lme)
sd(unlist(ranef(gam1$lme)[[3]]))
summary(gam1$gam)
plot(gam1$gam)

cold_pop_dat$fit<-predict(gam1$gam,se.fit=TRUE)$fit
cold_pop_dat$sefit<-predict(gam1$gam,se.fit=TRUE)$se.fit

pred_dat<-expand.grid(
  exp_batch=levels(cold_pop_dat$exp_batch),
  sex=levels(cold_pop_dat$sex),
  latitude=seq(min(cold_pop_dat$latitude),max(cold_pop_dat$latitude)),
  altitude=seq(min(cold_pop_dat$altitude),max(cold_pop_dat$altitude),500))

pred_dat$fit<-predict(object = gam1$gam,newdata=pred_dat,type="response")
head(pred_dat)
pred_dat2<-ddply(pred_dat,.variables = .(sex,latitude,altitude),summarize,
                 "latitude"=mean(latitude),
                 "altitude"=mean(altitude),
                 "fit"=mean(fit))
pred_dat2

ggplot()+
  geom_errorbar(data=plot_dat,aes(lat,
                                  ymax=mean_ctmin+se_ctmin,
                                  ymin=mean_ctmin-se_ctmin,group=sex),
                width=0,position=position_dodge(0.5))+
#  geom_point(data=cold_pop_dat,aes(x=latitude,Ctmin,fill=sex),
#             shape=21,size=3,position=position_dodge(0.5))+
  geom_line(data=cold_pop_dat,aes(x=latitude,y=fit,linetype=sex))+
  geom_point(data=plot_dat,aes(x=lat,mean_ctmin,fill=sex),
             shape=21,size=5,position=position_dodge(0.5))+
  geom_label(data=plot_dat[plot_dat$sex=="female",],aes(x=lat,y=mean_ctmin+0.5,label=population))+
  scale_fill_manual("Sex",values=c("black","grey70"),labels=c("Female","Male"))+
  scale_linetype_discrete("Sex",labels=c("Female","Male"))+
  xlab(expression(paste("Latitude ( "^o,"North )")))+
  ylab(expression(paste("CTmin ( "^o,C," )")))+
  my.theme+theme(axis.text = element_text(size=16),axis.title = element_text(size=16))

# All data plotted
head(cold_pop_dat)
ggplot()+
  geom_line(data=cold_pop_dat,aes(x=latitude,y=fit,linetype=sex))+
  geom_point(data=cold_pop_dat,aes(x=latitude,Ctmin,fill=sex),
             shape=21,size=2,position=position_dodge(0.5))+
  geom_label(data=plot_dat[plot_dat$sex=="female",],aes(x=lat,y=mean_ctmin+0.5,label=population))+
  scale_fill_manual("Sex",values=c("black","grey70"),labels=c("Female","Male"))+
  scale_linetype_discrete("Sex",labels=c("Female","Male"))+
  xlab(expression(paste("Latitude ( "^o,"North )")))+
  ylab(expression(paste("CTmin ( "^o,C," )")))+
  my.theme+theme(axis.text = element_text(size=16),axis.title = element_text(size=16))

# Is CCRT affected by the ambient room temperature during the experiment?
ggplot()+
  geom_point(data=cold_pop_dat,aes(x=meanT,y=CCRT_totalsec))+
  ylab("CCRT (seconds)")+
  xlab("Mean Ambient Temperature (C)")+
  facet_wrap(population~.)+
  my.theme

anova(lm(CCRT_totalsec~population+meanT,data=cold_pop_dat))
#It appears not
tapply(cold_pop_dat$CCRT_totalsec,INDEX = list(cold_pop_dat$population),mean)
tapply(cold_pop_dat$CCRT_totalsec,INDEX = list(cold_pop_dat$population),median)
ggplot()+
  geom_histogram(data=cold_pop_dat,aes(x=CCRT_totalsec))+
  xlab("CCRT (seconds)")+
  ylab("Count")+
  facet_wrap(population~.)+
  my.theme

plot_dat<-ddply(cold_pop_dat,.variables = .(population,sex),summarise,
                mean_ccrt=mean(CCRT_totalsec,na.rm=TRUE),
                se_ccrt=se(CCRT_totalsec),
                lat=mean(latitude))
plot_dat

# With latitude as the predictor instead of population
m<-lmer(CCRT_totalsec~latitude*sex+(1|exp_batch),data=cold_pop_dat)
m1<-lm(CCRT_totalsec~latitude*sex,data=cold_pop_dat)
anova(m,m1)
# Removing the random effect is NOT fine
m<-lmer(CCRT_totalsec~latitude*sex+(1|exp_batch),data=cold_pop_dat)
m1<-lmer(CCRT_totalsec~latitude+sex+(1|exp_batch),data=cold_pop_dat)
m2<-lmer(CCRT_totalsec~latitude+(1|exp_batch),data=cold_pop_dat)
m3<-lmer(CCRT_totalsec~(1|exp_batch),data=cold_pop_dat)
summary(m)
anova(m,m1)
anova(m1,m2)
anova(m2,m3)
# m2 is the best 
plot(m2)
hist(resid(m))

# Fit GAM (cubic spline). With a GAMM I can include the batch random effect
gam0<-gam(CCRT_totalsec~s(latitude,bs="cr",k=5)+s(altitude,bs="cr",k=5)+sex,data=cold_pop_dat)
summary(gam0)
gam1<-gamm(CCRT_totalsec~s(latitude,bs="cr",k=5)+s(altitude,bs="cr",k=5)+sex,data=cold_pop_dat,random=list(exp_batch=~1))
gam1<-gamm(CCRT_totalsec~s(latitude,bs="cr",k=5),data=cold_pop_dat,random=list(exp_batch=~1))
summary(gam1$lme)
summary(gam1$gam)
plot(gam1$gam)

cold_pop_dat$fit<-predict(gam1$gam,se.fit=TRUE)$fit
cold_pop_dat$sefit<-predict(gam1$gam,se.fit=TRUE)$se.fit

pred_dat<-expand.grid(
  exp_batch=levels(cold_pop_dat$exp_batch),
  sex=levels(cold_pop_dat$sex),
  latitude=seq(min(cold_pop_dat$latitude),max(cold_pop_dat$latitude)),
  altitude=seq(min(cold_pop_dat$altitude),max(cold_pop_dat$altitude),500))

pred_dat$fit<-predict(object = gam1$gam,newdata=pred_dat,type="response")
head(pred_dat)
pred_dat2<-ddply(pred_dat,.variables = .(sex,latitude,altitude),summarize,
                 "latitude"=mean(latitude),
                 "altitude"=mean(altitude),
                 "fit"=mean(fit))
pred_dat2

pred_dat<-expand.grid(
                      exp_batch=levels(cold_pop_dat$exp_batch),
                      latitude=seq(min(cold_pop_dat$latitude),max(cold_pop_dat$latitude)))
pred_dat$fit<-predict(object = m2,newdata=pred_dat,type="response")
head(pred_dat)

ggplot()+
  geom_errorbar(data=plot_dat,aes(lat,
                                  ymax=mean_ccrt+se_ccrt,
                                  ymin=mean_ccrt-se_ccrt,group=sex),
                width=0,position=position_dodge(0.25))+
  geom_line(data=cold_pop_dat,aes(x=latitude,y=fit))+
  geom_point(data=plot_dat,aes(lat,mean_ccrt,fill=sex),
             shape=21,size=3,position=position_dodge(0.25))+
  scale_fill_manual("Sex",values=c("black","grey70"),labels=c("Female","Male"))+
  xlab(expression(paste("Latitude ( "^o,"North )")))+
  ylab(expression(paste("CCRT (seconds)")))+
  my.theme+theme(axis.text = element_text(size=16),axis.title = element_text(size=16))

# Plot all data
head(cold_pop_dat)
ggplot()+
  geom_line(data=cold_pop_dat,aes(x=latitude,y=fit,linetype=sex))+
  geom_point(data=cold_pop_dat,aes(x=latitude,CCRT_totalsec,fill=sex),
             shape=21,size=2,position=position_dodge(0.5))+
  scale_fill_manual("Sex",values=c("black","grey70"),labels=c("Female","Male"))+
  scale_linetype_discrete("Sex",labels=c("Female","Male"))+
  xlab(expression(paste("Latitude ( "^o,"North )")))+
  ylab("CCRT (seconds)")+
  my.theme+theme(axis.text = element_text(size=16),axis.title = element_text(size=16))

# Summarise ctmin and ccrt to means for each population
ctmin<-tapply(cold_pop_dat$Ctmin,
       INDEX = list(cold_pop_dat$population),
       mean, na.rm = TRUE)
ctmin<-tapply(cold_pop_dat$Ctmin,
              INDEX = list(cold_pop_dat$population,cold_pop_dat$sex),
              mean, na.rm = TRUE)

ctmin_med<-tapply(cold_pop_dat$Ctmin,
              INDEX = list(cold_pop_dat$population),
              median, na.rm = TRUE)
plot(x=ctmin,y=ctmin_med)
cor.test(x=ctmin,y=ctmin_med)
# Male-female
plot(x=ctmin[,1],y=ctmin[,2])
cor.test(x=ctmin[,1],y=ctmin[,2])


tableS1$ctmin<-vector(length=nrow(tableS1))
tableS1$ctmin[
  match(order,
        tableS1$LocationName[
          tableS1$PoolGen=="1"])]<-scale(ctmin[
            match(order,names(ctmin))],center = TRUE,scale=TRUE)

ccrt<-tapply(cold_pop_dat$CCRT_totalsec,
       INDEX = list(cold_pop_dat$population),
       mean, na.rm = TRUE)
ccrt_med<-tapply(cold_pop_dat$CCRT_totalsec,
             INDEX = list(cold_pop_dat$population),
             median, na.rm = TRUE)

plot(x=ccrt,y=ccrt_med)
cor.test(x=ccrt,y=ccrt_med)

tableS1$ccrt<-vector(length=nrow(tableS1))
tableS1$ccrt[
  match(order,
        tableS1$LocationName[
          tableS1$PoolGen=="1"])]<-scale(ccrt[
            match(order,names(ccrt))],center = TRUE,scale=TRUE)

env_vars_tab<-tableS1[
  match(order,tableS1$LocationName[tableS1$PoolGen=="1"]),c(3,66,67)]


ctmin<-scale(ctmin[match(order,names(ctmin))])
ccrt<-scale(ccrt[match(order,names(ccrt))])

env_vars_tab$ctmin<-ctmin
env_vars_tab$ccrt<-ccrt
env_vars_tab$LocationName<-gsub(" ","_",env_vars_tab$LocationName)
env_vars_tab

# Write env data table
write.table(env_vars_tab,
            "~/Dropbox/st_andrews/montana_clines_MS/Manuscript/V10_R2/supplemental_information/env_vars_bscan.tab",
            col.names = FALSE,row.names=FALSE,quote=FALSE,sep=" ")
# Write Table S1
write.table(tableS1,
            "~/Dropbox/st_andrews/montana_clines_MS/Manuscript/V10_R2/supplemental_informationTable/montana_pop_info_bioclimdat.tsv",
            col.names = TRUE,row.names=FALSE,quote=FALSE,sep="\t")


# Load data (table S1)
tableS1<-read.table("~/Dropbox/st_andrews/montana_clines_MS/Manuscript/V10_R2/supplemental_information/montana_pop_info_bioclimdat.tsv",
                 header=TRUE, stringsAsFactors=FALSE,sep="\t")

# Are ctmin or ccrt correlated to either PC1 or PC2
cor.test(tableS1$ctmin[tableS1$PoolGen == "1"],tableS1$PC1[tableS1$PoolGen == "1"])
ct_pc1<-ggplot()+geom_point(data=tableS1[which(tableS1$PoolGen == "1"),],aes(ctmin,PC1),size=3)+xlab("CTmin")+my.theme
cor.test(tableS1$ctmin[tableS1$PoolGen == "1"],tableS1$PC2[tableS1$PoolGen == "1"])
ct_pc2<-ggplot()+geom_point(data=tableS1[which(tableS1$PoolGen == "1"),],aes(ctmin,PC2),size=3)+xlab("CTmin")+my.theme
cor.test(tableS1$ccrt[tableS1$PoolGen == "1"],tableS1$PC1[tableS1$PoolGen == "1"])
ccrt_pc1<-ggplot()+geom_point(data=tableS1[which(tableS1$PoolGen == "1"),],aes(ccrt,PC1),size=3)+xlab("CCRT")+my.theme
cor.test(tableS1$ccrt[tableS1$PoolGen == "1"],tableS1$PC2[tableS1$PoolGen == "1"])
ccrt_pc2<-ggplot()+geom_point(data=tableS1[which(tableS1$PoolGen == "1"),],aes(ccrt,PC2),size=3)+xlab("CCRT")+my.theme
plot_grid(ct_pc1,ct_pc2,ccrt_pc1,ccrt_pc2)
# Draw maps
# Are ctmin or ccrt correlated to each other
ct_ccrt<-ggplot()+geom_point(data=tableS1[which(tableS1$PoolGen == "1"),],aes(x=ctmin,y=ccrt),size=3)+xlab("CTmin")+ylab("CCRT")+my.theme
cor.test(tableS1$ctmin[tableS1$PoolGen == "1"],tableS1$ccrt[tableS1$PoolGen == "1"])


tableS1[tableS1$Region=="NoAm",]
lat<-c(min(tableS1$Lat[tableS1$Region=="NoAm"]),max(tableS1$Lat[tableS1$Region=="NoAm"]))
lon<-c(min(tableS1$Long[tableS1$Region=="NoAm"]),max(tableS1$Long[tableS1$Region=="NoAm"]))

NA_W_map<-get_map(source = "stamen",maptype = "terrain",location = c(lon=mean(lon),lat=mean(lat)),zoom = 4)

ggmap(NA_W_map)+
  geom_point(data=tableS1[tableS1$Region=="NoAm" & tableS1$PoolGen == 1,],
             aes(x=Long,y=Lat),fill="Blue",size=5,shape=21,alpha=1/2)+
  geom_point(data=tableS1[tableS1$Region=="NoAm" & tableS1$PoolGen == 0,],
             aes(x=Long,y=Lat),fill="Red",size=5,shape=24,alpha=1/2)+
  geom_label(data=tableS1[tableS1$Region=="NoAm"& tableS1$PoolGen == 1,],
             aes(x = Long+c(-10,-10,-10,-14),
                 y=Lat+c(0,0,0,-2),label=LocationName),size = 7)+
  xlab("Longitude")+
  ylab("Latitude")+my.theme+theme(axis.text = element_text(size=14),
                                  text = element_text(size = 14))

lat<-c(min(tableS1$Lat[tableS1$Region=="Eur"]),max(tableS1$Lat[tableS1$Region=="Eur"]))
lon<-c(min(tableS1$Long[tableS1$Region=="Eur"]),max(tableS1$Long[tableS1$Region=="Eur"]))

Eur_N_map<-get_map(source = "stamen",maptype = "terrain",location = c(lon=mean(lon),lat=mean(lat)),zoom = 4)

ggmap(Eur_N_map)+
  geom_point(data=tableS1[tableS1$Region=="Eur" & tableS1$PoolGen == 1,],
             aes(x=Long,y=Lat),fill="Blue",size = 5,shape=21,alpha=1/2)+
  geom_point(data=tableS1[tableS1$Region=="Eur"& tableS1$PoolGen == 0,],
             aes(x=Long,y=Lat),fill="Red",size = 5,shape=22,alpha=1/2)+
  geom_label(data=tableS1[tableS1$Region=="Eur" & tableS1$PoolGen == 1,],
             aes(x = Long+c(3,3),y=Lat,label=LocationName),size = 7)+
  xlab("Longitude")+
  ylab("Latitude")+my.theme+theme(axis.text = element_text(size=14),
                                  text = element_text(size = 14))
```
