## Supplementary material for "Cold adaptation drives population genomic divergence in the ecological specialist, *Drosophila montana*": montana_clinalpops_BSCAN.R

```
##
# D. montana project clinal populations
# BSCAN data analysis
# Last Modified: Aug 2017
##
# Clean environment
rm(list = ls(all = TRUE))

#
# Load libraries
#install.packages("coda")
#source("http://bioconductor.org/biocLite.R")
#biocLite("qvalue")
library(ggplot2)
library(qvalue)
library(reshape)
library(coda)
library(UpSetR)
#citation("ggmap")
source("~/projects/RData/RScripts/ggplot_theme.R")
options(scipen=10)
#Define Functions
l_uniq<-function(x){length(unique(x))}

#------------------------------#
# Check BAYESCENV convergence
#------------------------------#
var<-"ccrt"
# X chromosomes
# read in the .sel files to check convergence
chain1<-read.table(
  paste("~/projects/D_montana/clinal_population_genomics/data/SNPs/clim_",
        var,"_bscanout/X_minc1_cov0-1000_neff_all_bscanin.sel",
        sep=""),
  header=TRUE)

chain2<-read.table(
  paste("~/projects/D_montana/clinal_population_genomics/data/SNPs/clim_",
        var,"_bscanout_chain2/X_minc1_cov0-1000_neff_all_bscanin.sel",
        sep=""),
  header=TRUE)

chain3<-read.table(
  paste("~/projects/D_montana/clinal_population_genomics/data/SNPs/clim_",
        var,"_bscanout_chain3/X_minc1_cov0-1000_neff_all_bscanin.sel",
        sep=""),
  header=TRUE)

chain4<-read.table(
  paste("~/projects/D_montana/clinal_population_genomics/data/SNPs/clim_",
        var,"_bscanout_chain4/X_minc1_cov0-1000_neff_all_bscanin.sel",
        sep=""),
  header=TRUE)

chain1<-chain1[-c(1)]
chain1<-mcmc(chain1,thin=10)
head(chain1,n=50)
rejectionRate(chain1)
plot(chain1)
# Check autocorrelation between sampled values and check effective sample size
autocorr.diag(chain1)
effectiveSize(chain1)
# Check convergence
geweke.diag(chain1, frac1=0.3, frac2=0.3)
heidel.diag(chain1, eps=0.1, pvalue=0.05)

chain2<-chain2[-c(1)]
chain2<-mcmc(chain2,thin=10)
head(chain2,n=50)
plot(chain2)
# Check autocorrelation between sampled values and check effective sample size
autocorr.diag(chain2)
effectiveSize(chain2)
# Check convergence
geweke.diag(chain2, frac1=0.3, frac2=0.3)
heidel.diag(chain2, eps=0.1, pvalue=0.05)

chain3<-chain3[-c(1)]
chain3<-mcmc(chain3,thin=10)
head(chain3,n=50)
plot(chain3)
# Check autocorrelation between sampled values and check effective sample size
autocorr.diag(chain3)
effectiveSize(chain3)
# Check convergence
geweke.diag(chain3, frac1=0.3, frac2=0.3)
heidel.diag(chain3, eps=0.1, pvalue=0.05)

chain4<-chain4[-c(1)]
chain4<-mcmc(chain4,thin=10)
head(chain4,n=50)
plot(chain4)
# Check autocorrelation between sampled values and check effective sample size
autocorr.diag(chain4)
effectiveSize(chain4)
# Check convergence
geweke.diag(chain4, frac1=0.3, frac2=0.3)
heidel.diag(chain4, eps=0.1, pvalue=0.05)

combined_chains<-mcmc.list(chain2,chain3,chain4)
pdf(
  paste("~/projects/D_montana/clinal_population_genomics/data/SNPs/X_clim_",
        var,"_BayScEnvChainsPlot.pdf",sep=""),width = 32,height = 32,
  pointsize = 30)
plot(combined_chains)
dev.off()
gelman.diag(combined_chains)
gelman.plot(combined_chains,ylim=c(0,4))

# Autosomes
# read in the .sel files to check convergence
var="ccrt"
chain1<-read.table(
  paste("~/projects/D_montana/clinal_population_genomics/data/SNPs/clim_",
        var,"_bscanout/","Aut_minc1_cov0-1000_raw_all_bscanin.sel",
        sep=""),
  header=TRUE)
chain2<-read.table(
  paste("~/projects/D_montana/clinal_population_genomics/data/SNPs/clim_",
        var,"_bscanout_chain2/Aut_minc1_cov0-1000_raw_all_bscanin.sel",
        sep=""),
  header=TRUE)
chain3<-read.table(
  paste("~/projects/D_montana/clinal_population_genomics/data/SNPs/clim_",
        var,"_bscanout_chain3/Aut_minc1_cov0-1000_raw_all_bscanin.sel",
        sep=""),
  header=TRUE)
chain4<-read.table(
  paste("~/projects/D_montana/clinal_population_genomics/data/SNPs/clim_",
        var,"_bscanout_chain4/Aut_minc1_cov0-1000_raw_all_bscanin.sel",
        sep=""),
  header=TRUE)

chain1<-chain1[-c(1)]
chain1<-mcmc(chain1,thin=10)
head(chain1,n=50)
plot(chain1)
# Check autocorrelation between sampled values and check effective sample size
autocorr.diag(chain1)
effectiveSize(chain1)
# Check convergence
geweke.diag(chain1, frac1=0.3, frac2=0.3)
heidel.diag(chain1, eps=0.1, pvalue=0.05)

chain2<-chain2[-c(1)]
chain2<-mcmc(chain2,thin=10)
head(chain2,n=50)
plot(chain2)
# Check autocorrelation between sampled values and check effective sample size
autocorr.diag(chain2)
effectiveSize(chain2)
# Check convergence
geweke.diag(chain2, frac1=0.3, frac2=0.3)
heidel.diag(chain2, eps=0.1, pvalue=0.05)

chain3<-chain3[-c(1)]
chain3<-mcmc(chain3,thin=10)
head(chain3,n=50)
plot(chain3)
# Check autocorrelation between sampled values and check effective sample size
autocorr.diag(chain3)
effectiveSize(chain3)
# Check convergence
geweke.diag(chain3, frac1=0.3, frac2=0.3)
heidel.diag(chain3, eps=0.1, pvalue=0.05)

chain4<-chain4[-c(1)]
chain4<-mcmc(chain4,thin=10)
head(chain4,n=50)
plot(chain4)
# Check autocorrelation between sampled values and check effective sample size
autocorr.diag(chain4)
effectiveSize(chain4)
# Check convergence
geweke.diag(chain4, frac1=0.3, frac2=0.3)
heidel.diag(chain4, eps=0.1, pvalue=0.05)

combined_chains<-mcmc.list(chain2,chain3,chain4)
pdf(
  paste("~/projects/D_montana/clinal_population_genomics/data/SNPs/Aut_clim_",
        var,"_BayScEnvChainsPlot.pdf",sep=""),width = 32,height = 32,
  pointsize = 30)
plot(combined_chains)
dev.off()
gelman.diag(combined_chains)
gelman.plot(combined_chains,ylim=c(0,4))

#------------------------------#
# Load BAYESCENV data
#------------------------------#
# Take the max (should be conservative) of p-values across chains

# Take the top SNPs as the set of SNPs where the max q-value across chains < 0.05

chrom="X"
var<-"pc1"
bscansnps_dat_chains<-data.frame()
dir<-"~/projects/D_montana/clinal_population_genomics/data/SNPs"
if(chrom == "Aut"){
  bscansnps_dat<-read.table(
    paste(dir,"/",chrom,"_minc1_cov0-1000_raw_all_snps.txt",
                                  sep=""),header=FALSE)
  colnames(bscansnps_dat)<-c("chr","pos","snp")
  
  chains<-c("_chain2","_chain3","_chain4")
  top_snps<-list()
  for(chain in chains){
    top_snps[[chain]]<-scan(
      paste(dir,"/clim_",var,"_bscanout",
            chain,"/",chrom,"_",var,"_sig_snps.txt",sep=""),
                          what = "character")
  }
  # Get the common top snps
  common_top_snps<-Reduce(intersect,top_snps)
  top_snps_g<-list()
  top_snps_g_qval<-list()
  top_snps_fst<-list()
  top_snps_snp<-list()
  bkr_snps_g<-list()
  bkr_snps_g_qval<-list()
  bkr_snps_fst<-list()
  bkr_snps_snp<-list()
  for(chain in chains){
    bscanout_dat<-read.table(
      paste(dir,"/clim_",var,"_bscanout",
            chain,"/",chrom,
            "_minc1_cov0-1000_raw_all_bscanin_fst_sub.txt",sep=""),
      header = FALSE)
    colnames(bscanout_dat)<-c("snp","g_qval","g","fst")
    top_snps_g[[chain]]<-bscanout_dat$g[as.numeric(common_top_snps)]
    top_snps_g_qval[[chain]]<-bscanout_dat$g_qval[
      as.numeric(common_top_snps)]
    top_snps_fst[[chain]]<-bscanout_dat$fst[as.numeric(common_top_snps)]
    top_snps_snp[[chain]]<-as.numeric(common_top_snps)
    
    bkr_snps_g[[chain]]<-bscanout_dat$g[-as.numeric(common_top_snps)]
    bkr_snps_g_qval[[chain]]<-bscanout_dat$g_qval[
      -as.numeric(common_top_snps)]
    bkr_snps_fst[[chain]]<-bscanout_dat$fst[-as.numeric(common_top_snps)]
    bkr_snps_snp[[chain]]<-bscanout_dat$snp[-as.numeric(common_top_snps)]
  }
  top_snps_g<-apply(
    matrix(unlist(top_snps_g),
           ncol=length(top_snps_g),byrow=FALSE),1,mean)
  top_snps_g_qval<-apply(
    matrix(unlist(top_snps_g_qval),
           ncol=length(top_snps_g_qval),byrow=FALSE),1,max)
  top_snps_fst<-apply(
    matrix(unlist(top_snps_fst),
           ncol=length(top_snps_fst),byrow=FALSE),1,mean)
  top_snps_snp<-apply(
    matrix(unlist(top_snps_snp),
           ncol=length(top_snps_snp),byrow=FALSE),1,mean)
  top_snps_bscanout_dat<-data.frame("g"=top_snps_g,
                                    "g_qval"=top_snps_g_qval,
                                    "fst"=top_snps_fst,
                                    "snp"=top_snps_snp)
  bkr_snps_g<-apply(
    matrix(unlist(bkr_snps_g),
           ncol=length(bkr_snps_g),byrow=FALSE),1,mean)
  bkr_snps_g_qval<-apply(
    matrix(unlist(bkr_snps_g_qval),
           ncol=length(bkr_snps_g_qval),byrow=FALSE),1,max)
  bkr_snps_fst<-apply(
    matrix(unlist(bkr_snps_fst),
           ncol=length(bkr_snps_fst),byrow=FALSE),1,mean)
  bkr_snps_snp<-apply(
    matrix(unlist(bkr_snps_snp),
           ncol=length(bkr_snps_snp),byrow=FALSE),1,mean)
  bkr_snps_bscanout_dat<-data.frame("g"=bkr_snps_g,
                                    "g_qval"=bkr_snps_g_qval,
                                    "fst"=bkr_snps_fst,
                                    "snp"=bkr_snps_snp)
  bscanout_dat<-rbind(bkr_snps_bscanout_dat,top_snps_bscanout_dat)
  bscanout_dat<-bscanout_dat[order(bscanout_dat$snp),]
}else if(chrom == "X"){
  bscansnps_dat<-read.table(
    paste(dir,"/",chrom,"_minc1_cov0-1000_neff_all_snps.txt",sep=""),
    header=FALSE)
  colnames(bscansnps_dat)<-c("chr","pos","snp")
  
  # Take the top SNPs as the union of top SNPs across chains
  # For all other SNPs, take the average g and q-value across chains
  chains<-c("_chain2","_chain3","_chain4")
  top_snps<-list()
  for(chain in chains){
    top_snps[[chain]]<-scan(paste(dir,"/clim_",var,"_bscanout",
                                chain,"/",chrom,"_",
                                var,"_sig_snps.txt",sep=""),
                          what = "character")
  }
  # Get the common top snps
  common_top_snps<-Reduce(intersect,top_snps)
  top_snps_g<-list()
  top_snps_g_qval<-list()
  top_snps_fst<-list()
  top_snps_snp<-list()
  bkr_snps_g<-list()
  bkr_snps_g_qval<-list()
  bkr_snps_fst<-list()
  bkr_snps_snp<-list()
  for(chain in chains){
    bscanout_dat<-read.table(
      paste(dir,"/clim_",var,"_bscanout",
            chain,"/",chrom,
            "_minc1_cov0-1000_neff_all_bscanin_fst_sub.txt",sep=""),
      header = FALSE)
    colnames(bscanout_dat)<-c("snp","g_qval","g","fst")
    top_snps_g[[chain]]<-bscanout_dat$g[as.numeric(common_top_snps)]
    top_snps_g_qval[[chain]]<-bscanout_dat$g_qval[
      as.numeric(common_top_snps)]
    top_snps_fst[[chain]]<-bscanout_dat$fst[as.numeric(common_top_snps)]
    top_snps_snp[[chain]]<-as.numeric(common_top_snps)
    
    bkr_snps_g[[chain]]<-bscanout_dat$g[-as.numeric(common_top_snps)]
    bkr_snps_g_qval[[chain]]<-bscanout_dat$g_qval[
      -as.numeric(common_top_snps)]
    bkr_snps_fst[[chain]]<-bscanout_dat$fst[-as.numeric(common_top_snps)]
    bkr_snps_snp[[chain]]<-bscanout_dat$snp[-as.numeric(common_top_snps)]
  }
  top_snps_g<-apply(
    matrix(unlist(top_snps_g),
           ncol=length(top_snps_g),byrow=FALSE),1,mean)
  top_snps_g_qval<-apply(
    matrix(unlist(top_snps_g_qval),
           ncol=length(top_snps_g_qval),byrow=FALSE),1,max)
  top_snps_fst<-apply(
    matrix(unlist(top_snps_fst),
           ncol=length(top_snps_fst),byrow=FALSE),1,mean)
  top_snps_snp<-apply(
    matrix(unlist(top_snps_snp),
           ncol=length(top_snps_snp),byrow=FALSE),1,mean)
  top_snps_bscanout_dat<-data.frame("g"=top_snps_g,
                                    "g_qval"=top_snps_g_qval,
                                    "fst"=top_snps_fst,
                                    "snp"=top_snps_snp)
#  length(top_snps_g)
#  length(top_snps_g_qval)
#  length(top_snps_fst)
#  length(top_snps_snp)
  bkr_snps_g<-apply(
    matrix(unlist(bkr_snps_g),
           ncol=length(bkr_snps_g),byrow=FALSE),1,mean)
  bkr_snps_g_qval<-apply(
    matrix(unlist(bkr_snps_g_qval),
           ncol=length(bkr_snps_g_qval),byrow=FALSE),1,max)
  bkr_snps_fst<-apply(
    matrix(unlist(bkr_snps_fst),
           ncol=length(bkr_snps_fst),byrow=FALSE),1,mean)
  bkr_snps_snp<-apply(
    matrix(unlist(bkr_snps_snp),
           ncol=length(bkr_snps_snp),byrow=FALSE),1,mean)
  bkr_snps_bscanout_dat<-data.frame("g"=bkr_snps_g,
                                    "g_qval"=bkr_snps_g_qval,
                                    "fst"=bkr_snps_fst,
                                    "snp"=bkr_snps_snp)
  bscanout_dat<-rbind(bkr_snps_bscanout_dat,top_snps_bscanout_dat)
  bscanout_dat<-bscanout_dat[order(bscanout_dat$snp),]
}

head(bscanout_dat)
length(which(bscanout_dat$g_qval<0.05))
head(bscansnps_dat)
hist(bscanout_dat$g_qval,xlim=c(0,1))
# Combine the data
bscandat<-cbind(bscansnps_dat[,c(1,2)],bscanout_dat[,c(1,2,3)])
bscandat$scaff<-gsub("-size.*","",bscandat$chr)
head(bscandat)
#-----------------------------------#
# Write the top SNPs to a .gtf file #
#-----------------------------------#
#####
chr_info<-read.table("~/projects/D_montana/montana_genome/montana_scaffsizes.tab",
                     header=FALSE,sep=",")
colnames(chr_info)<-c("chr","length")
head(chr_info)

top_bed_dat<-cbind(as.character(bscandat$chr[bscandat$g_qval<0.05]),
                   bscandat$pos[bscandat$g_qval<0.05],
                   bscandat$pos[bscandat$g_qval<0.05],
                   paste(bscandat$chr[bscandat$g_qval<0.05],
                         "_",bscandat$pos[bscandat$g_qval<0.05],
                         sep=""))
top_bed_dat<-data.frame(top_bed_dat)
top_bed_dat$X2<-as.numeric(as.character(top_bed_dat$X2))
top_bed_dat$X3<-as.numeric(as.character(top_bed_dat$X3))
head(top_bed_dat)

write.table(top_bed_dat,
            paste(dir,"/",chrom,"_",
                  var,"_minc1_cov0-1000_raw_top_snps.bed",sep=""),
            row.names=FALSE,col.names=FALSE,quote=FALSE,sep="\t")
X<-10000
top_bed_dat$X2<-top_bed_dat$X2-X
top_bed_dat$X3<-top_bed_dat$X3+X

# Remove coordinates < 0
top_bed_dat[which(top_bed_dat[,2] < 0),2] <- 0
# Remove coordinates > max_chrom_len
for(c in unique(top_bed_dat$chr)){
  chrom_max_length<-chr_info$length[chr_info$chr==c]
  top_bed_dat[
    which(top_bed_dat[,1]==c & top_bed_dat[,3] > chrom_max_length),3]<-chrom_max_length
}
head(top_bed_dat)
write.table(top_bed_dat,
            paste(dir,"/",chrom,"_",
                  var,"_minc1_cov0-1000_raw_top_snps_10kb.bed",sep=""),
            row.names=FALSE,col.names=FALSE,quote=FALSE,sep="\t")
#####

#------------------------------#
# Load the scaffold size data  #
# and the linkage group data   #
#------------------------------#
# Scaffold sizes from genome freeze v.1.4
scaffdat<-read.table(
  "~/projects/D_montana/montana_genome/montana_scaffsizes.tab",
                     header=FALSE,sep=",")
colnames(scaffdat)<-c("scaff","size")
head(scaffdat)
# Load the v1 genome linkage groups
LGdat1<-read.table(
  "~/projects/D_montana/montana_genome/markers_and_positions_v2.1.csv",
                   header=TRUE,sep="\t")
colnames(LGdat1)<-c("scaff","len","lkg_pos","lg","lgmrk")
head(LGdat1)
# Load the v1.4 genome linkage groups
LGdat1.4<-read.table(
  "~/projects/D_montana/montana_genome/vennys_map_info_v1.4.txt",
                     header=TRUE,sep="\t")
LGdat1.4$scaf_name<-gsub("-size.*","",LGdat1.4$scaf_name)
colnames(LGdat1.4)<-c("scaff","linkage_gr","size")
LGdat1.4$linkage_gr<-paste("Chr ",LGdat1.4$linkage_gr,sep="")
LGdat1.4$lkg_pos<-vector(length=nrow(LGdat1.4))
# Add scaffold positions on LGs to LGdat1.4
for(row in 1:nrow(LGdat1.4)){
  scf<-LGdat1.4$scaff[row]
  LGdat1.4$lkg_pos[row]<-LGdat1$lkg_pos[LGdat1$scaff==scf]
}
LGdat1.4<-LGdat1.4[order(LGdat1.4$linkage_gr,LGdat1.4$lkg_pos),]
head(LGdat1.4)

#---------------------------------#
# BAYSCENV SUMMARIES AND ANALYSIS #
#---------------------------------#
## Add linkage groups and positions to the data set
# Add scaffold and position information to bscan_dat
bscandat$lg<-vector(length=nrow(bscandat))
bscandat$lg_pos<-vector(length=nrow(bscandat))
bscandat$len<-vector(length=nrow(bscandat))
for(scaff in unique(bscandat$chr)){
  if(scaff %in% unique(LGdat1$scaff)){
    bscandat$lg[
      bscandat$chr==as.character(scaff)]<-as.character(LGdat1$lg[
        LGdat1$scaff==as.character(scaff)])
    bscandat$lg_pos[
      bscandat$chr==as.character(scaff)]<-LGdat1$lkg_pos[
        LGdat1$scaff==as.character(scaff)]
    bscandat$len[
      bscandat$chr==as.character(scaff)]<-LGdat1$len[
        LGdat1$scaff==as.character(scaff)]
  }else{
    bscandat$lg[bscandat$chr==as.character(scaff)]<-"unplaced"
    bscandat$lg_pos[bscandat$chr==as.character(scaff)]<-0
  }
}
head(bscandat)

# Subset to "placed" scaffolds. 
# I should have no unplaced SNPs because 
# I limit my analysis to SNPs we can reliably place
nrow(bscandat[bscandat$lg=="unplaced",])
bscan_dat_pl<-bscandat[bscandat$lg!="unplaced",]
head(bscan_dat_pl)
# Sort first by linkage group and then by scaffold position on linkage group
bscan_dat_pl<-bscan_dat_pl[order(bscan_dat_pl$lg,bscan_dat_pl$lg_pos),]
# Add new absolute positions along lgs for the SNPs.
bscan_dat_pl$abspos<-vector(length=nrow(bscan_dat_pl))
bscan_dat_pl_abspos<-data.frame()
bscan_dat_pl$chr<-as.character(bscan_dat_pl$chr)
for(lg in unique(bscan_dat_pl$lg)){
  lg_dat<-bscan_dat_pl[bscan_dat_pl$lg==lg,]
  tot_l<-0
  last_scaff<-unique(lg_dat$chr)[1]
  last_scaff_l<-LGdat1$len[LGdat1$scaff==last_scaff]
  for(scaff in unique(lg_dat$chr)){
    curr_scaff_l<-LGdat1$len[LGdat1$scaff==scaff]
    if(scaff == last_scaff){
      lg_dat$abspos[lg_dat$chr==scaff]<- lg_dat$pos[
        lg_dat$chr==scaff]+tot_l
    }else{
      tot_l<-tot_l+last_scaff_l
      lg_dat$abspos[lg_dat$chr==scaff] <- lg_dat$pos[
        lg_dat$chr==scaff]+tot_l
      last_scaff_l<-curr_scaff_l
    }
  }
  bscan_dat_pl_abspos<-rbind(bscan_dat_pl_abspos,lg_dat)
}
head(bscan_dat_pl_abspos)
# Write as table: PLACED SNPS
write.table(bscan_dat_pl_abspos,
            paste(dir,"/",chrom,"_",var,"_all_bscanin_fst_sub_pl.txt",
                  sep=""),
            quote=FALSE,col.names=TRUE,row.names=FALSE,sep = "\t")


#----------------#
# PLOTTING       #
#----------------#
####
# Read tables
var<-"pc1"
dir<-"~/projects/D_montana/clinal_population_genomics/data/SNPs"
X_bscan_dat_pl_abspos<-read.table(
  paste(dir,"/X_",var,
        "_all_bscanin_fst_sub_pl.txt",sep=""),
  header=TRUE,sep="\t")
Aut_bscan_dat_pl_abspos<-read.table(
  paste(dir,"/Aut_",var,
        "_all_bscanin_fst_sub_pl.txt",sep=""),
                     header=TRUE,sep="\t")
head(Aut_bscan_dat_pl_abspos)
head(X_bscan_dat_pl_abspos)
bscan_dat_pl_abspos<-rbind(
  Aut_bscan_dat_pl_abspos[,c(1,2,3,4,5,7,8,9,10)],
  X_bscan_dat_pl_abspos[,c(1,2,3,4,5,7,8,9,10)])
rm(Aut_bscan_dat_pl_abspos)
rm(X_bscan_dat_pl_abspos)
head(bscan_dat_pl_abspos)
nrow(bscan_dat_pl_abspos)

# Plotting: PLACED SNPS
head(bscan_dat_pl_abspos)
bscan_dat_pl_abspos$chrom<-gsub("LG","Chromosome ",bscan_dat_pl_abspos$lg)
manhplot<-ggplot()+
  geom_point(data=bscan_dat_pl_abspos[
    which(bscan_dat_pl_abspos$g_qval >= 0.05),],
    aes(abspos/1000000,-log10(g_qval),colour=chr),
    size=0.2)+
  geom_point(data=bscan_dat_pl_abspos[
    which(bscan_dat_pl_abspos$g_qval < 0.05),],
    aes(abspos/1000000,-log10(g_qval)),
    size=0.2,colour="red")+
  geom_hline(yintercept = -log10(0.05),linetype="dashed")+
  scale_colour_manual("",
                      values=rep(c("black","grey"),
                                 length.out=length(
                                   unique(bscan_dat_pl_abspos$chr))),
                      guide=FALSE)+
  xlab("Position (Mb)")+
  ylab(expression(paste("-log"[10],"(q-value) of ",italic(g),sep="")))+
  facet_grid(chrom~.)+
  #ggtitle(paste("Results for ",var,sep=""))+
  my.theme+
  theme(axis.text.y = element_text(size=20),
        axis.text.x = element_text(size=20),
        axis.title = element_text(size=20),
        strip.text.y = element_text(size=20,angle=0))
#png(paste("manhattan_",var,sep=""))
manhplot # w = 1192, h = 788


ggplot()+
  geom_histogram(data=bscan_dat_pl_abspos,aes(fst))+
  facet_grid(lg~.)+
  my.theme+theme(axis.text=element_text(size=10),
                 axis.title = element_text(size=10),
                 strip.text.x = element_text(size=10))
  
# Subset X snps from background
X<-100000
bscan_dat_pl_abspos_sub<-bscan_dat_pl_abspos[
  which(bscan_dat_pl_abspos$g_qval >= 0.05),]
bscan_dat_pl_abspos_sub<-bscan_dat_pl_abspos[
  sample(seq(1,nrow(bscan_dat_pl_abspos_sub)),
         size=X,replace=FALSE),]

fst_v_qval_plot<-ggplot() +
  geom_point(data = bscan_dat_pl_abspos_sub,
             aes(-log10(g_qval),fst),size=0.4,colour="grey30",alpha=1/3) +
  geom_point(data = bscan_dat_pl_abspos[
    bscan_dat_pl_abspos$g_qval < 0.05,],
             aes(-log10(g_qval),fst),colour="firebrick",alpha=1/3) +
  geom_vline(xintercept = -log10(0.05),linetype="dashed")+
  xlab(expression(paste("-log"[10],"(q-value) of ",italic(g))))+
  ylab(expression(paste("F"[st])))+
  ggtitle(var) + 
  my.theme+theme(axis.text=element_text(size=10),
                 axis.title = element_text(size=10),
                 strip.text.x = element_text(size=10))
fst_v_qval_plot 


# How many "top" SNPs are there 
qval_thr<-0.05
nrow(bscan_dat_pl_abspos[bscan_dat_pl_abspos$g_qval < qval_thr,]) # Mapped
var
# How many SNPs on each lg
lg_snps<-tapply(bscan_dat_pl_abspos$len,
                INDEX = list(bscan_dat_pl_abspos$lg),FUN=length)
lg_snps
# How many top SNPs on each lg
lg_top_snps<-tapply(bscan_dat_pl_abspos$len[
  bscan_dat_pl_abspos$g_qval < qval_thr],
                    INDEX = list(
                      bscan_dat_pl_abspos$lg[
                        bscan_dat_pl_abspos$g_qval < qval_thr]),
                    FUN=length)
lg_top_snps
lg_top_snps<-lg_top_snps[!(is.na(lg_top_snps))]
lg_snps<-lg_snps[names(lg_top_snps)]
chisq.test(lg_snps)
chisq.test(lg_top_snps)
chisq.test(lg_top_snps,p = lg_snps/sum(lg_snps))
str(chisq.test(lg_top_snps,p = lg_snps/sum(lg_snps)))

# How long is each lg
lg_ls<-tapply(LGdat1.4$size,
              INDEX = list(LGdat1.4$linkage_gr),sum)/1000000
# Correlations?
cor.test(lg_snps,lg_ls,method="spearman")
cor.test(lg_top_snps,lg_ls,method="spearman")
plot(lg_ls,lg_snps)
plot(lg_ls,lg_top_snps)

# Write the top SNPs as a table
#PLACED
head(bscan_dat_pl_abspos)
write.table(bscan_dat_pl_abspos[
  bscan_dat_pl_abspos$g_qval < qval_thr,c(1,2,5,3,4)],
            paste(chrom,"_minc5_cov37-106_neff_all_bscanin_",
                  var,"_fst_topSNPs.txt",
                  sep=""),sep="\t",
            quote=FALSE,col.names=FALSE,row.names=FALSE)


#----------------#
# CLOSEST GENES  #
#----------------#

# How many bootstrap samples to do
nboots<-10000

# Load the lists of closest genes
Autpc1<-read.table("~/projects/D_montana/clinal_population_genomics/data/SNPs/Aut_pc1_minc1_cov0-1000_raw_top_snps_10kb_genesoverlap.list",header=FALSE,sep="\t")
Xpc1<-read.table("~/projects/D_montana/clinal_population_genomics/data/SNPs/X_pc1_minc1_cov0-1000_raw_top_snps_10kb_genesoverlap.list",header=FALSE,sep="\t")
pc1<-rbind(Autpc1,Xpc1)
rm(Autpc1,Xpc1)

Autpc2<-read.table("~/projects/D_montana/clinal_population_genomics/data/SNPs/Aut_pc2_minc1_cov0-1000_raw_top_snps_10kb_genesoverlap.list",header=FALSE,sep="\t")
Xpc2<-read.table("~/projects/D_montana/clinal_population_genomics/data/SNPs/X_pc2_minc1_cov0-1000_raw_top_snps_10kb_genesoverlap.list",header=FALSE,sep="\t")
pc2<-rbind(Autpc2,Xpc2)
rm(Autpc2,Xpc2)

Autctmin<-read.table("~/projects/D_montana/clinal_population_genomics/data/SNPs/Aut_ctmin_minc1_cov0-1000_raw_top_snps_10kb_genesoverlap.list",header=FALSE,sep="\t")
Xctmin<-read.table("~/projects/D_montana/clinal_population_genomics/data/SNPs/X_ctmin_minc1_cov0-1000_raw_top_snps_10kb_genesoverlap.list",header=FALSE,sep="\t")
ctmin<-rbind(Autctmin,Xctmin)
rm(Autctmin,Xctmin)

Autccrt<-read.table("~/projects/D_montana/clinal_population_genomics/data/SNPs/Aut_ccrt_minc1_cov0-1000_raw_top_snps_10kb_genesoverlap.list",header=FALSE,sep="\t")
Xccrt<-read.table("~/projects/D_montana/clinal_population_genomics/data/SNPs/X_ccrt_minc1_cov0-1000_raw_top_snps_10kb_genesoverlap.list",header=FALSE,sep="\t")
ccrt<-rbind(Autccrt,Xccrt)
rm(Autccrt,Xccrt)

top_genes<-c(as.character(pc1$V1),as.character(pc2$V1),as.character(ctmin$V1),as.character(ccrt$V1))
top_genes<-unique(top_genes)
all_top_genes<-data.frame("top_genes"=top_genes)

dmon_genes<-read.table("~/projects/D_montana/montana_genome/D_montana_genome_freeze_v1.4.tab",header=FALSE)
colnames(dmon_genes)<-c("gene","sym")
nrow(dmon_genes)
head(dmon_genes)

#--------------------------------------------#
# Load Parker et al., 2015 Table S1
#--------------------------------------------#
parker2015<-read.table("~/Dropbox/st_andrews/montana_clines_MS/data/gene_lists/Parker_et_al_2015/Parker_cold_acc_genes.csv",
                       header=TRUE,sep=",")

head(parker2015)
head(ccrt)
# How many are DE in both
unique(parker2015$Species)
nrow(parker2015[parker2015$sp=="both",])
N<-length(parker2015$FlybaseNumber[parker2015$Species %in% c("both","D. montana")])
# How many genes occur in the Parker et al., 2015 set of genes that are 
# DE in Dmon and in both
all_top_genes$parker2015<-rep("0",nrow(all_top_genes))
which_o<-which(as.character(all_top_genes$top_genes) %in% 
  as.character(parker2015$FlybaseNumber[parker2015$Species %in% c("both","D. montana")]))
all_top_genes$parker2015[which_o]<-"1"


all_o<-c(as.character(
  parker2015$FlybaseNumber[parker2015$Species %in% c("both","D. montana") &
                             as.character(parker2015$FlybaseNumber) %in% as.character(pc1$V1)]),
  as.character(
    parker2015$FlybaseNumber[parker2015$Species %in% c("both","D. montana") &
                               as.character(parker2015$FlybaseNumber) %in% as.character(pc2$V1)]),
  as.character(
    parker2015$FlybaseNumber[parker2015$Species %in% c("both","D. montana") &
                               as.character(parker2015$FlybaseNumber) %in% as.character(ctmin$V1)]),
  as.character(
    parker2015$FlybaseNumber[parker2015$Species %in% c("both","D. montana") &
                               as.character(parker2015$FlybaseNumber) %in% as.character(ccrt$V1)]))

n_overlaps<-length(unique(all_o))
# How many expected by chance:
# Bootstrap
n_genes<-length(unique(c(pc1$V1,pc2$V1,ccrt$V1,ctmin$V1)))
(n_overlaps/n_genes)*100

bootsamps<-vector(length=nboots)
for(boot in 1:nboots){
  genes<-sample(dmon_genes$gene,size = n_genes,replace = FALSE)
  bootsamps[boot]<-length(
      parker2015$FlybaseNumber[parker2015$Species %in% c("both","D. montana") & 
                           parker2015$FlybaseNumber %in% genes])
}
hist_dat<-data.frame("perc"=((bootsamps/n_genes)*100))
# 'p-value'
p<-(length(bootsamps[bootsamps>n_overlaps])/length(bootsamps))
if(p == 0){
  p <- "< 0.001"
} else {
  p<-paste("= ",p,sep="")
}
parker2015plot<-
  ggplot()+
  ggtitle(
    paste("Parker et al., 2015 (N = ",N,"): N overlaps = ",n_overlaps,
          "\nBootstrap Ovarlaps = ",round(mean(bootsamps),2),
          " [",paste(quantile(bootsamps,probs = c(0.025,0.975)),collapse=", "),"], p ",p,sep=""))+
  
  geom_histogram(data=hist_dat,aes(perc),bins=10)+
  geom_vline(xintercept = (n_overlaps/n_genes)*100,linetype="solid",colour="red",size=1)+
  xlab("Overlapping Genes (%)")+
  ylab("Count")+
  my.theme
parker2015plot

#--------------------------------------------#
# Load Kankare et al., 2016 diapause Table S1
#--------------------------------------------#
kankare2016<-read.table("~/Dropbox/st_andrews/montana_clines_MS/data/gene_lists/Kankare_et_al_2016/Kankare_et_al_2016_TableS1.csv",
                        header=TRUE,sep=",",quote = '"')
kankare2016$Gene.Name<-gsub("Dvir_","Dvir\\\\",kankare2016$Gene.Name)
nrow(kankare2016)
# Remove any genes that are not Dvir orthologs
kankare2016<-kankare2016[grep("Dvir",kankare2016$Gene.Name),]
# Add FBgn id column. 
kankare2016$FBgnID<-dmon_genes$gene[match(kankare2016$Gene.Name,dmon_genes$sym)]
N<-nrow(kankare2016)
kankare2016<-kankare2016[which(!is.na(kankare2016$FBgnID)),]
N<-length(unique(kankare2016$FBgnID))
nrow(kankare2016)
head(kankare2016)

all_top_genes$kankare2016<-rep("0",nrow(all_top_genes))
which_o<-which(as.character(all_top_genes$top_genes) %in% unique(kankare2016$FBgnID))
all_top_genes$kankare2016[which_o]<-"1"

all_o<-c(as.character(kankare2016$Gene.Name[kankare2016$FBgnID %in% as.character(pc1$V1)]),
         as.character(kankare2016$Gene.Name[kankare2016$FBgnID %in% as.character(pc2$V1)]),
         as.character(kankare2016$Gene.Name[kankare2016$FBgnID %in% as.character(ctmin$V1)]),
         as.character(kankare2016$Gene.Name[kankare2016$FBgnID %in% as.character(ccrt$V1)]))

n_overlaps<-length(unique(all_o))
n_genes<-length(unique(c(pc1$V1,pc2$V1,ccrt$V1,ctmin$V1)))
(n_overlaps/n_genes)*100

bootsamps<-vector(length=nboots)
for(boot in 1:nboots){
  genes<-sample(dmon_genes$gene,size = n_genes,replace = FALSE)
  bootsamps[boot]<-length(
    unique(kankare2016$Gene.Name[kankare2016$FBgnID %in% genes]))
}
hist_dat<-data.frame("perc"=((bootsamps/n_genes)*100))
# 'p-value'
p<-(length(bootsamps[bootsamps>n_overlaps])/length(bootsamps))
if(p == 0){
  p <- "< 0.001"
} else {
  p<-paste("= ",p,sep="")
}
kankare2016plot<-ggplot()+
  ggtitle(
    paste("Kankare et al., 2016 (N = ",N,"): N overlaps = ",n_overlaps,
          "\nBootstrap Ovarlaps = ",round(mean(bootsamps),2),
          " [",paste(quantile(bootsamps,probs = c(0.025,0.975)),collapse=", "),"], p ",p,sep=""))+
  geom_histogram(data=hist_dat,aes(perc),bins=10)+
  geom_vline(xintercept = (n_overlaps/n_genes)*100,linetype="solid",colour="red",size=1)+
  xlab("Overlapping Genes (%)")+
  ylab("Count")+
  my.theme
kankare2016plot
#--------------------------------------------#
# Load Kankare et al., 2010 montana genome paper
#--------------------------------------------#
kankare2010<-read.table("~/Dropbox/st_andrews/montana_clines_MS/data/gene_lists/Kankare_et_al_2010/DEgenes.csv",
                        header=TRUE,sep=",")
head(kankare2010)
nrow(kankare2010)
# Remove any genes that are not Dvir orthologs
kankare2010<-kankare2010[grep("Dvir",kankare2010$DROSOPHILA_ORTHOLOGS),]
# Add FBgn id column. 
kankare2010$FBgnID<-dmon_genes$gene[match(kankare2010$DROSOPHILA_ORTHOLOGS,dmon_genes$sym)]
# Remove any genes with NA as Dvir FBgnID orthologs
kankare2010<-kankare2010[which(!is.na(kankare2010$FBgnID)),]
N<-length(unique(kankare2010$FBgnID))

all_top_genes$kankare2010<-rep("0",nrow(all_top_genes))
which_o<-which(as.character(all_top_genes$top_genes) %in% unique(kankare2010$FBgnID))
all_top_genes$kankare2010[which_o]<-"1"

all_o<-c(as.character(kankare2010$DROSOPHILA_ORTHOLOGS[kankare2010$FBgnID %in% as.character(pc1$V1)]),
         as.character(kankare2010$DROSOPHILA_ORTHOLOGS[kankare2010$FBgnID %in% as.character(pc2$V1)]),
         as.character(kankare2010$DROSOPHILA_ORTHOLOGS[kankare2010$FBgnID %in% as.character(ctmin$V1)]),
         as.character(kankare2010$DROSOPHILA_ORTHOLOGS[kankare2010$FBgnID %in% as.character(ccrt$V1)]))


n_overlaps<-length(unique(all_o))
n_genes<-length(unique(c(pc1$V1,pc2$V1,ccrt$V1,ctmin$V1)))
(n_overlaps/n_genes)*100

bootsamps<-vector(length=nboots)
for(boot in 1:nboots){
  genes<-sample(dmon_genes$gene,size = n_genes,replace = FALSE)
  bootsamps[boot]<-length(
    unique(kankare2010$DROSOPHILA_ORTHOLOGS[as.character(kankare2010$FBgnID) %in% as.character(genes)]))
}
hist_dat<-data.frame("perc"=((bootsamps/n_genes)*100))
# 'p-value'
p<-(length(bootsamps[bootsamps>n_overlaps])/length(bootsamps))
if(p == 0){
  p <- "< 0.001"
} else {
  p<-paste("= ",p,sep="")
}
kankare2010plot<-ggplot()+
  ggtitle(
    paste("Kankare et al., 2010 (N = ",N,"): N overlaps = ",n_overlaps,
          "\nBootstrap Ovarlaps = ",round(mean(bootsamps),2),
          " [",paste(quantile(bootsamps,probs = c(0.025,0.975)),collapse=", "),"], p ",p,sep=""))+
  geom_histogram(data=hist_dat,aes(perc),bins=10)+
  geom_vline(xintercept = (n_overlaps/n_genes)*100,linetype="solid",colour="red",size=1)+
  xlab("Overlapping Genes (%)")+
  ylab("Count")+
  my.theme
kankare2010plot

#--------------------------------------------#
# Load Kauranen et al., 2019 diapause paper
#--------------------------------------------#
kauranen2019<-read.table("~/Dropbox/st_andrews/montana_clines_MS/data/gene_lists/Kauranen_et_al_2019/TableS2.csv",
                        header=TRUE,sep=",")
head(kauranen2019)
nrow(kauranen2019)
N<-nrow(kauranen2019)

all_top_genes$kauranen2019<-rep("0",nrow(all_top_genes))
which_o<-which(as.character(all_top_genes$top_genes) %in% unique(kauranen2019$SUBMITTED.ID))
all_top_genes$kauranen2019[which_o]<-"1"

all_o<-c(as.character(kauranen2019$SUBMITTED.ID[kauranen2019$SUBMITTED.ID %in% as.character(pc1$V1)]),
         as.character(kauranen2019$SUBMITTED.ID[kauranen2019$SUBMITTED.ID %in% as.character(pc2$V1)]),
         as.character(kauranen2019$SUBMITTED.ID[kauranen2019$SUBMITTED.ID %in% as.character(ctmin$V1)]),
         as.character(kauranen2019$SUBMITTED.ID[kauranen2019$SUBMITTED.ID %in% as.character(ccrt$V1)]))


n_overlaps<-length(unique(all_o))
n_genes<-length(unique(c(pc1$V1,pc2$V1,ccrt$V1,ctmin$V1)))
(n_overlaps/n_genes)*100

bootsamps<-vector(length=nboots)
for(boot in 1:nboots){
  genes<-sample(dmon_genes$gene,size = n_genes,replace = FALSE)
  bootsamps[boot]<-length(
    unique(kauranen2019$SUBMITTED.ID[as.character(kauranen2019$SUBMITTED.ID) %in% as.character(genes)]))
}
hist_dat<-data.frame("perc"=((bootsamps/n_genes)*100))
# 'p-value'
p<-(length(bootsamps[bootsamps>n_overlaps])/length(bootsamps))
if(p == 0){
  p <- "< 0.001"
} else {
  p<-paste("= ",p,sep="")
}
kauranen2019plot<-ggplot()+
  ggtitle(
    paste("Kauranen et al., 2019 (N = ",N,"): N overlaps = ",n_overlaps,
          "\nBootstrap Ovarlaps = ",round(mean(bootsamps),2),
          " [",paste(quantile(bootsamps,probs = c(0.025,0.975)),collapse=", "),"], p ",p,sep=""))+
  geom_histogram(data=hist_dat,aes(perc),bins=10)+
  geom_vline(xintercept = (n_overlaps/n_genes)*100,linetype="solid",colour="red",size=1)+
  xlab("Overlapping Genes (%)")+
  ylab("Count")+
  my.theme
kauranen2019plot

#--------------------------------------------#
# Load Parker et al., 2016 changing light Table S1
#--------------------------------------------#
parker2016<-read.table("~/Dropbox/st_andrews/montana_clines_MS/data/gene_lists/Parker_et_al_2016/light_and_dia_high_low_all_annot_3a_HIGH_AS_REF2.csv.txt",
                        header=TRUE,sep=",")
parker2016hits<-parker2016$FlybaseGeneNumber[which(parker2016$FDR_dia_light < 0.05 | parker2016$FDR_ND_light < 0.05)]
parker2016hits<-parker2016hits[which(!is.na(parker2016hits))]
N<-length(unique(parker2016hits))

all_top_genes$parker2016<-rep("0",nrow(all_top_genes))
which_o<-which(as.character(all_top_genes$top_genes) %in% unique(parker2016hits))
all_top_genes$parker2016[which_o]<-"1"

all_o<-c(parker2016hits[which(as.character(parker2016hits) %in% as.character(pc1$V1))],
         parker2016hits[which(as.character(parker2016hits) %in% as.character(pc2$V1))],
         parker2016hits[which(as.character(parker2016hits) %in% as.character(ctmin$V1))],
         parker2016hits[which(as.character(parker2016hits) %in% as.character(ccrt$V1))])

n_overlaps<-length(unique(all_o))
n_genes<-length(unique(c(pc1$V1,pc2$V1,ccrt$V1,ctmin$V1)))
(n_overlaps/n_genes)*100

bootsamps<-vector(length=nboots)
for(boot in 1:nboots){
  genes<-sample(dmon_genes$gene,size = n_genes,replace = FALSE)
  bootsamps[boot]<-length(
    unique(parker2016hits[as.character(parker2016hits) %in% as.character(genes)]))
}
hist_dat<-data.frame("perc"=((bootsamps/n_genes)*100))
# 'p-value'
p<-(length(bootsamps[bootsamps>n_overlaps])/length(bootsamps))
if(p == 0){
  p <- "< 0.001"
} else {
  p<-paste("= ",p,sep="")
}
parker2016plot<-ggplot()+
  ggtitle(
    paste("Parker et al., 2016 (N = ",N,"): N overlaps = ",n_overlaps,
          "\nBootstrap Ovarlaps = ",round(mean(bootsamps),2),
          " [",paste(quantile(bootsamps,probs = c(0.025,0.975)),collapse=", "),"], p ",p,sep=""))+
  geom_histogram(data=hist_dat,aes(perc),bins=10)+
  geom_vline(xintercept = (n_overlaps/n_genes)*100,linetype="solid",colour="red",size=1)+
  xlab("Overlapping Genes (%)")+
  ylab("Count")+
  my.theme
parker2016plot


#--------------------------------------------#
# Load Parker et al., 2018 montana genome paper
#--------------------------------------------#
parker2018<-read.table("~/Dropbox/st_andrews/montana_clines_MS/data/gene_lists/Parker_et_al_2018_montana_genome_paper_SOI/Supplemental_Table_2.csv",
                       sep=",",header=TRUE,comment.char = "#")
pw_tests<-c("CoOl","OlVa","CoVa")
parker2018_all_pw<-data.frame()
for(pw in pw_tests){
  pw_dat<-read.table(
    paste(
      "~/Dropbox/st_andrews/montana_clines_MS/data/gene_lists/Parker_et_al_2018_montana_genome_paper_SOI/montana_genome_reseqpops_",
      pw,
      "_sigsnps_closestgene1kb.tab",sep=""),
             sep="\t",header=FALSE,comment.char = "#")
  # Remove "." entries (no nearby genes)
  pw_dat<-pw_dat[pw_dat$V7 != ".",]
  pw_dat$pw<-rep(pw,nrow(pw_dat))
  parker2018_all_pw<-rbind(parker2018_all_pw,pw_dat)
}
colnames(parker2018_all_pw)<-c("snp_scaff","snp_pos1","snp_pos2","g_scaff","g_start","g_end","g_name","dist","pw")

head(parker2018)
parker2018_all_pw$FBgnID<-parker2018$FB_vir_gene_id[match(parker2018_all_pw$g_name,parker2018$DM_gene_name)]

head(parker2018_all_pw)
# Sanity check that these numbers match the paper
tapply(parker2018_all_pw$FBgnID,INDEX = list(parker2018_all_pw$pw),function(x){length(unique(x))})
tapply(parker2018_all_pw$g_name,INDEX = list(parker2018_all_pw$pw),function(x){length(unique(x))})
tapply(parker2018_all_pw$FBgnID,INDEX = list(parker2018_all_pw$pw),length)

# Subset to those that have D. virilis ortholog
parker2018_all_pw<-parker2018_all_pw[which(!is.na(parker2018_all_pw$FBgnID)),]
N<-length(unique(parker2018_all_pw$FBgnID))

all_top_genes$parker2018<-rep("0",nrow(all_top_genes))
which_o<-which(as.character(all_top_genes$top_genes) %in% unique(parker2018_all_pw$FBgnID))
all_top_genes$parker2018[which_o]<-"1"

all_o<-c(parker2018_all_pw$FBgnID[which(as.character(parker2018_all_pw$FBgnID) %in% as.character(pc1$V1))],
         parker2018_all_pw$FBgnID[which(as.character(parker2018_all_pw$FBgnID) %in% as.character(pc2$V1))],
         parker2018_all_pw$FBgnID[which(as.character(parker2018_all_pw$FBgnID) %in% as.character(ctmin$V1))],
         parker2018_all_pw$FBgnID[which(as.character(parker2018_all_pw$FBgnID) %in% as.character(ccrt$V1))])

n_overlaps<-length(unique(all_o))
n_genes<-length(unique(c(pc1$V1,pc2$V1,ccrt$V1,ctmin$V1)))
(n_overlaps/n_genes)*100

bootsamps<-vector(length=nboots)
for(boot in 1:nboots){
  genes<-sample(dmon_genes$gene,size = n_genes,replace = FALSE)
  bootsamps[boot]<-length(
    unique(parker2018_all_pw$FBgnID[which(as.character(parker2018_all_pw$FBgnID) %in% as.character(genes))]))
}
hist_dat<-data.frame("perc"=((bootsamps/n_genes)*100))
# 'p-value'
p<-(length(bootsamps[bootsamps>n_overlaps])/length(bootsamps))
if(p == 0){
  p <- "< 0.001"
} else {
  p<-paste("= ",p,sep="")
}
parker2018plot<-ggplot()+
  ggtitle(
    paste("Parker et al., 2018 (N = ",N,"): N overlaps = ",n_overlaps,
          "\nBootstrap Ovarlaps = ",round(mean(bootsamps),2),
          " [",paste(quantile(bootsamps,probs = c(0.025,0.975)),collapse=", "),"], p ",p,sep=""))+
  geom_histogram(data=hist_dat,aes(perc),bins=10)+
  geom_vline(xintercept = (n_overlaps/n_genes)*100,linetype="solid",colour="red",size=1)+
  xlab("Overlapping Genes (%)")+
  ylab("Count")+
  my.theme
parker2018plot


cowplot::plot_grid(kankare2010plot,
                   parker2015plot,
                   kankare2016plot,
                   parker2016plot,
                   parker2018plot,
                   kauranen2019plot,
                   ncol=1)

# Save table giving the top genes that overlap
all_top_genes$name<-dmon_genes$sym[match(all_top_genes$top_genes,dmon_genes$gene)]
write.table(all_top_genes,"~/Dropbox/st_andrews/montana_clines_MS/geneoverlaps.csv",
      sep = ",",col.names=TRUE,row.names=FALSE,quote=FALSE)

# ####


# Make an UpSet plot of the overlaps between genes associated with the different PCs and phenotypes

chroms<-c("Aut","X")
vars<-c("pc1","pc2","ctmin","ccrt")
genes<-list()
counts<-list()
for(var in vars){
  genes[[var]]<-vector()
  for(chrom in chroms){
    g<-scan(paste("~/projects/D_montana/clinal_population_genomics/data/SNPs/",
                  chrom,"_",var,"_minc1_cov0-1000_raw_top_snps_10kb_genesoverlap.list",sep=""),
            what = "character")
    genes[[var]]<-c(genes[[var]],g)
  }
  counts[[var]]<-length(genes[[var]])
}
length(genes$pc2)
head(genes$pc2)
upset(fromList(genes),order.by = "freq")


# Load the total set of genes
dmon_genes<-read.table("~/projects/D_montana/montana_genome/D_montana_genome_freeze_v1.4.tab",header=FALSE)
colnames(dmon_genes)<-c("gene","sym")
nrow(dmon_genes)
head(dmon_genes)

# Do some re-sampling
# for each iteration get 4 samples of genes (pc1, pc2, ctmin, ccrt) and check the size of the intersection.
n<-1000 # number of samples
intersection<-vector(length=n)
i<-1
for(i in 1:n){
  iter_samples<-list()
  for(var in vars){
    iter_samples[[var]]<-as.character(sample(dmon_genes$gene,counts[[var]],replace = FALSE))
  }
  intersection[i]<-length(Reduce(intersect,iter_samples))
}
hist(intersection)
mean(intersection)
se(intersection)
quantile(intersection,probs=c(0.025,0.975))


# Plot the locations of important genes on the manhattan plot
# Genes which are DE in at least two of the previous RNAseq studies 
# and also occur in this study.
#####
# Common genes
commongenes<-c("FBgn0203225",
               "FBgn0204964","FBgn0205258",
               "FBgn0205992","FBgn0207876")
geneids<-c("CG42313","Pex12",
           "CG9008","Yp3","Slik")
commongenedat<-data.frame(lg=vector(length=length(commongenes)),
                    geneid=geneids,
                    gene=commongenes,
                    g_spos=vector(length=length(commongenes)),
                    g_epos=vector(length=length(commongenes)),
                    gscaff=vector(length=length(commongenes)),
                    g_lgspos=vector(length=length(commongenes)),
                    g_lgepos=vector(length=length(commongenes)))


# PC1
pc1_uniquegenes<-c("FBgn0205243","FBgn0207689")
pc1_geneids<-c("GJ1807","Inos")
pc1_genedat<-data.frame(lg=vector(length=length(pc1_uniquegenes)),
                        geneid=pc1_geneids,
                        gene=pc1_uniquegenes,
                        g_spos=vector(length=length(pc1_uniquegenes)),
                        g_epos=vector(length=length(pc1_uniquegenes)),
                        gscaff=vector(length=length(pc1_uniquegenes)),
                        g_lgspos=vector(length=length(pc1_uniquegenes)),
                        g_lgepos=vector(length=length(pc1_uniquegenes)))

# PC2
pc2_uniquegenes<-c("FBgn0199132","FBgn0203500",
                   "FBgn0204709","FBgn0210867")
pc2_geneids<-c("Ltn1","GJ16316","vri","Sap47")
pc2_genedat<-data.frame(lg=vector(length=length(pc2_uniquegenes)),
                        geneid=pc2_geneids,
                        gene=pc2_uniquegenes,
                        g_spos=vector(length=length(pc2_uniquegenes)),
                        g_epos=vector(length=length(pc2_uniquegenes)),
                        gscaff=vector(length=length(pc2_uniquegenes)),
                        g_lgspos=vector(length=length(pc2_uniquegenes)),
                        g_lgepos=vector(length=length(pc2_uniquegenes)))

# Combine with common genes
genedat<-rbind(commongenedat,pc1_genedat,pc2_genedat)
genedat$dat<-c(rep("Common",nrow(commongenedat)),
               rep("PC1",nrow(pc1_genedat)),
               rep("PC2",nrow(pc2_genedat)))


# Gather plotting data for arrows: PC1
i<-1
for(gene in genedat$gene){
  # if PC1 or Common:
  if(genedat$dat[genedat$gene==gene] =="PC1" | 
     genedat$dat[genedat$gene==gene] =="Common"){
    # Get the positions of SNPs near gene
    # If there are more than one SNP, take the closest.
    s_spos<-min(clgenepc1_1Mb$s_spos[clgenepc1_1Mb$name == gene])
    s_scaff<-clgenepc1_1Mb$scaff[clgenepc1_1Mb$name == gene]
    
    # Get the distance of the gene from the SNP
    # If there are more than one SNP, take the closest.
    dist<-clgenepc1_1Mb$dist[clgenepc1_1Mb$name == gene]
    
    # Get the length of the gene
    g_l<-clgenepc1_1Mb$g_epos[
      clgenepc1_1Mb$name == gene]-clgenepc1_1Mb$g_spos[
        clgenepc1_1Mb$name == gene]
    
    # Get the scaffold and scaffold pos of the gene
    genedat$g_spos[i]<-clgenepc1_1Mb$g_spos[clgenepc1_1Mb$name == gene]
    genedat$g_epos[i]<-clgenepc1_1Mb$g_epos[clgenepc1_1Mb$name == gene]
    genedat$gscaff[i]<-as.character(
      clgenepc1_1Mb$gscaff[clgenepc1_1Mb$name == gene])
    
    # Get the linkage group and linkage group Pos of the SNP
    lg<-bscandatpc1$lg[bscandatpc1$chr == as.character(s_scaff) & 
                      bscandat$pos == s_spos]
    lgabspos<-bscandatpc1$lg_snp_pos[bscandatpc1$chr == as.character(s_scaff) & 
                                       bscandatpc1$pos == s_spos]
    genedat$lg[i]<-as.character(lg)
    genedat$g_lgspos[i]<-lgabspos+dist
    genedat$g_lgepos[i]<-lgabspos+dist+g_l
    i<-i+1
  }else if(genedat$dat[genedat$gene==gene] == "PC2"){
    # Get the positions of SNPs near gene
    # If there are more than one SNP, take the closest.
    s_spos<-min(clgenepc2_1Mb$s_spos[clgenepc2_1Mb$name == gene])
    s_scaff<-clgenepc2_1Mb$scaff[clgenepc2_1Mb$name == gene]
    
    # Get the distance of the gene from the SNP
    # If there are more than one SNP, take the closest.
    dist<-clgenepc2_1Mb$dist[clgenepc2_1Mb$name == gene]
    
    # Get the length of the gene
    g_l<-clgenepc2_1Mb$g_epos[
      clgenepc2_1Mb$name == gene]-clgenepc2_1Mb$g_spos[
        clgenepc2_1Mb$name == gene]
    
    # Get the scaffold and scaffold pos of the gene
    genedat$g_spos[i]<-clgenepc2_1Mb$g_spos[clgenepc2_1Mb$name == gene]
    genedat$g_epos[i]<-clgenepc2_1Mb$g_epos[clgenepc2_1Mb$name == gene]
    genedat$gscaff[i]<-as.character(
      clgenepc2_1Mb$gscaff[clgenepc2_1Mb$name == gene])
    
    # Get the linkage group and linkage group Pos of the SNP
    lg<-bscandatpc2$lg[bscandatpc2$chr == as.character(s_scaff) & 
                      bscandatpc2$pos == s_spos]
    lgabspos<-bscandatpc2$lg_snp_pos[bscandatpc2$chr == as.character(s_scaff) & 
                                    bscandatpc2$pos == s_spos]
    genedat$lg[i]<-as.character(lg)
    genedat$g_lgspos[i]<-lgabspos+dist
    genedat$g_lgepos[i]<-lgabspos+dist+g_l
    i<-i+1
  }
}
genedat


# Plot manhattan plot with arrows and labels for interesting genes: PC1
manhplot + ggtitle("") +
  geom_segment(data=genedat[genedat$dat == "PC1" & genedat$dat == "Common",],
               aes(yend=3,y=3.5,
                   xend=(g_lgspos/1000000),
                   x=(g_lgspos/1000000)),
               arrow = arrow(length = unit(0.1,"cm")))+
  geom_text(data=genedat,aes(y=3.6,x=(g_lgepos/1000000),label=geneid),size=2)

#####


###
# Other interesting genes from this study
#####
commongenes<-c("FBgn0202755","FBgn0204659")
geneids<-c("Hsp60C","mio")
commongenedat<-data.frame(lg=vector(length=length(commongenes)),
                          geneid=geneids,
                          gene=commongenes,
                          g_spos=vector(length=length(commongenes)),
                          g_epos=vector(length=length(commongenes)),
                          gscaff=vector(length=length(commongenes)),
                          g_lgspos=vector(length=length(commongenes)),
                          g_lgepos=vector(length=length(commongenes)))

# PC1
pc1_uniquegenes<-c("FBgn0022834","FBgn0015211","FBgn0199202",
                   "FBgn0198702","FBgn0208901")
pc1_geneids<-c("tim","gl","DnaJ-1","bora","so")
pc1_genedat<-data.frame(lg=vector(length=length(pc1_uniquegenes)),
                        geneid=pc1_geneids,
                        gene=pc1_uniquegenes,
                        g_spos=vector(length=length(pc1_uniquegenes)),
                        g_epos=vector(length=length(pc1_uniquegenes)),
                        gscaff=vector(length=length(pc1_uniquegenes)),
                        g_lgspos=vector(length=length(pc1_uniquegenes)),
                        g_lgepos=vector(length=length(pc1_uniquegenes)))

# PC2
pc2_uniquegenes<-c("FBgn0198685","FBgn0201989","FBgn0208150","FBgn0210387")
pc2_geneids<-c("Clk","nocte","inaC","slmb")
pc2_genedat<-data.frame(lg=vector(length=length(pc2_uniquegenes)),
                        geneid=pc2_geneids,
                        gene=pc2_uniquegenes,
                        g_spos=vector(length=length(pc2_uniquegenes)),
                        g_epos=vector(length=length(pc2_uniquegenes)),
                        gscaff=vector(length=length(pc2_uniquegenes)),
                        g_lgspos=vector(length=length(pc2_uniquegenes)),
                        g_lgepos=vector(length=length(pc2_uniquegenes)))

# Combine with common genes
genedat<-rbind(commongenedat,pc1_genedat,pc2_genedat)
genedat$dat<-c(rep("Common",nrow(commongenedat)),
               rep("PC1",nrow(pc1_genedat)),
               rep("PC2",nrow(pc2_genedat)))


# Gather plotting data
i<-1
for(gene in genedat$gene){
  # if PC1 or Common:
  if(genedat$dat[genedat$gene==gene] =="PC1" | 
     genedat$dat[genedat$gene==gene] =="Common"){
    # Get the positions of SNPs near gene
    # If there are more than one SNP, take the closest.
    s_spos<-min(clgenepc1_1Mb$s_spos[clgenepc1_1Mb$name == gene])
    s_scaff<-clgenepc1_1Mb$scaff[clgenepc1_1Mb$name == gene]
    
    # Get the distance of the gene from the SNP
    # If there are more than one SNP, take the closest.
    dist<-clgenepc1_1Mb$dist[clgenepc1_1Mb$name == gene]
    
    # Get the length of the gene
    g_l<-clgenepc1_1Mb$g_epos[
      clgenepc1_1Mb$name == gene]-clgenepc1_1Mb$g_spos[
        clgenepc1_1Mb$name == gene]
    
    # Get the scaffold and scaffold pos of the gene
    genedat$g_spos[i]<-clgenepc1_1Mb$g_spos[clgenepc1_1Mb$name == gene]
    genedat$g_epos[i]<-clgenepc1_1Mb$g_epos[clgenepc1_1Mb$name == gene]
    genedat$gscaff[i]<-as.character(
      clgenepc1_1Mb$gscaff[clgenepc1_1Mb$name == gene])
    
    # Get the linkage group and linkage group Pos of the SNP
    lg<-bscandatpc1$lg[bscandatpc1$chr == as.character(s_scaff) & 
                         bscandat$pos == s_spos]
    lgabspos<-bscandatpc1$lg_snp_pos[bscandatpc1$chr == as.character(s_scaff) & 
                                       bscandatpc1$pos == s_spos]
    genedat$lg[i]<-as.character(lg)
    genedat$g_lgspos[i]<-lgabspos+dist
    genedat$g_lgepos[i]<-lgabspos+dist+g_l
    i<-i+1
  }else if(genedat$dat[genedat$gene==gene] == "PC2"){
    # Get the positions of SNPs near gene
    # If there are more than one SNP, take the closest.
    s_spos<-min(clgenepc2_1Mb$s_spos[clgenepc2_1Mb$name == gene])
    s_scaff<-clgenepc2_1Mb$scaff[clgenepc2_1Mb$name == gene]
    
    # Get the distance of the gene from the SNP
    # If there are more than one SNP, take the closest.
    dist<-clgenepc2_1Mb$dist[clgenepc2_1Mb$name == gene]
    
    # Get the length of the gene
    g_l<-clgenepc2_1Mb$g_epos[
      clgenepc2_1Mb$name == gene]-clgenepc2_1Mb$g_spos[
        clgenepc2_1Mb$name == gene]
    
    # Get the scaffold and scaffold pos of the gene
    genedat$g_spos[i]<-clgenepc2_1Mb$g_spos[clgenepc2_1Mb$name == gene]
    genedat$g_epos[i]<-clgenepc2_1Mb$g_epos[clgenepc2_1Mb$name == gene]
    genedat$gscaff[i]<-as.character(
      clgenepc2_1Mb$gscaff[clgenepc2_1Mb$name == gene])
    
    # Get the linkage group and linkage group Pos of the SNP
    lg<-bscandatpc2$lg[bscandatpc2$chr == as.character(s_scaff) & 
                         bscandatpc2$pos == s_spos]
    lgabspos<-bscandatpc2$lg_snp_pos[bscandatpc2$chr == as.character(s_scaff) & 
                                       bscandatpc2$pos == s_spos]
    genedat$lg[i]<-as.character(lg)
    genedat$g_lgspos[i]<-lgabspos+dist
    genedat$g_lgepos[i]<-lgabspos+dist+g_l
    i<-i+1
  }
}
genedat
#####


# Run models with altitude and latitude for top snps.
# quasibinomial glms
# Load full .sync file
full_sync<-read.table("~/projects/D_montana/clinal_population_genomics/data/SNPs/bscanin/montana_clines_PoolSNP_AutXMins10000.sync",sep="\t")
head(full_sync)
# pop_order: Oulanka,Korpilahti,Ashford,Seward,Crested_Butte,Terrace
pops<-c("Oulanka","Korpilahti","Ashford","Seward","Crested_Butte","Terrace")
colnames(full_sync)<-c("chr","pos","ref",pops)
full_sync$chr<-gsub("-size.*","",full_sync$chr)
full_sync$snp<-paste(full_sync$chr,"_",full_sync$pos,sep="")

# Subset to "placed" SNPs
var<-"pc1" # Just use this data set, they are all the same
dir<-"~/projects/D_montana/clinal_population_genomics/data/SNPs"
X_bscan_dat_pl_abspos<-read.table(
  paste(dir,"/X_",var,
        "_all_bscanin_fst_sub_pl.txt",sep=""),
  header=TRUE,sep="\t")
Aut_bscan_dat_pl_abspos<-read.table(
  paste(dir,"/Aut_",var,
        "_all_bscanin_fst_sub_pl.txt",sep=""),
  header=TRUE,sep="\t")
bscan_dat_pl_abspos<-rbind(
  Aut_bscan_dat_pl_abspos[,c(1,2,3,4,5,7,8,9,10)],
  X_bscan_dat_pl_abspos[,c(1,2,3,4,5,7,8,9,10)])
rm(Aut_bscan_dat_pl_abspos)
rm(X_bscan_dat_pl_abspos)
head(bscan_dat_pl_abspos)
nrow(bscan_dat_pl_abspos)

bscan_dat_pl_abspos$chrom<-gsub("LG","Chromosome ",bscan_dat_pl_abspos$lg)
bscan_dat_pl_abspos$snp<-paste(bscan_dat_pl_abspos$chr,"_",bscan_dat_pl_abspos$pos,sep="")

nrow(full_sync)
length(bscan_dat_pl_abspos$snp)
length(which(full_sync$snp %in% bscan_dat_pl_abspos$snp))
placed_sync<-full_sync[which(full_sync$snp %in% bscan_dat_pl_abspos$snp),]
nrow(placed_sync)

# Save this file for (e.g.) supplement
write.table(placed_sync,"~/Dropbox/st_andrews/montana_clines_MS/Manuscript/V7_R1/supplemental_information/PoolSNP_placedSNPs.sync",
            col.names = TRUE,row.names = FALSE,quote=FALSE,sep="\t")

# Subset to top SNPs
placed_sync<-read.table("~/Dropbox/st_andrews/montana_clines_MS/Manuscript/V7_R1/supplemental_information/PoolSNP_placedSNPs.sync",
                        header=TRUE,sep="\t")
vars<-c("pc1","pc2","ctmin","ccrt")
top_snps<-list()
for(var in vars){
  top_snps[[var]]<-scan(paste("~/projects/D_montana/clinal_population_genomics/data/SNPs/",var,"_top_snps.list",sep=""),
                        what = "character",quiet = TRUE)
}
lapply(top_snps, length)

top_snps_list<-Reduce(intersect,x = top_snps)
length(top_snps_list)

top_sync<-placed_sync[which(placed_sync$snp %in% top_snps_list),]
nrow(top_sync)

# Reduce "top_sync" to alt and ref count columns for each snp
head(top_sync)
top_sync_counts<-top_sync[,c(1,2,3,10)]
top_sync_counts<-cbind(top_sync_counts,matrix(ncol=(length(pops)*2),nrow = nrow(top_sync_counts)))
colnames(top_sync_counts)<-c(colnames(top_sync[,c(1,2,3,10)]),paste(rep(pops,each=2),c("_maj_c","_alt_c"),sep=""))
head(top_sync_counts)
snp<-160
for(snp in 1:nrow(top_sync)){
  counts_m<-matrix(
    as.numeric(
      do.call(
        rbind,
        strsplit(
          as.character(unlist(as.vector(top_sync[snp,c(4,5,6,7,8,9)]))),split = ":"))),
    ncol = 6)
  # Remove all 0 columns (i.e. no counts in any population)
  counts_m<-counts_m[,which(apply(counts_m, 2, sum) != 0)]
  # Remove all columns with sum < 5 (i.e. no counts in any population)
  counts_m<-counts_m[,which(apply(counts_m, 2, sum) > 5)]
  if(ncol(counts_m)==2){
    # SNP biallelic, continue
    # Which allele is major (overall highest counts)
    maj_al<-which.max(apply(counts_m,2,sum))
    maj_al_c<-counts_m[,maj_al]
    alt_al_c<-counts_m[,-maj_al]
    top_sync_counts[snp,5:ncol(top_sync_counts)]<-c(t(matrix(cbind(maj_al_c,alt_al_c),ncol=2)))
  }
}
head(top_sync_counts)
head(top_sync)
#top_sync[which(top_sync$snp=="scaffold1332_18658"),]
# Save this file for (e.g.) supplement
write.table(top_sync_counts,"~/Dropbox/st_andrews/montana_clines_MS/Manuscript/V7_R1/supplemental_information/PoolSNP_topSNPs_counts.tab",
            col.names = TRUE,row.names = FALSE,quote=FALSE,sep="\t")


top_sync_counts<-read.table("~/Dropbox/st_andrews/montana_clines_MS/Manuscript/V7_R1/supplemental_information/PoolSNP_topSNPs_counts.tab",
                            header=TRUE,sep="\t")

# Melt the data
top_sync_counts_m<-melt(data=top_sync_counts,
                        id.vars=c("chr","pos","ref","snp"),
                        measure.vars=paste(pops,"_maj_c",sep=""))
top_sync_counts_m2<-melt(data=top_sync_counts,
                        id.vars=c("chr","pos","ref","snp"),
                        measure.vars=paste(pops,"_alt_c",sep=""))

colnames(top_sync_counts_m)<-c("chr","pos","ref","snp","pop","maj_c")
top_sync_counts_m$alt_c<-top_sync_counts_m2$value
top_sync_counts_m$pop<-gsub("_maj_c","",top_sync_counts_m$pop)
top_sync_counts_m$snp<-as.character(top_sync_counts_m$snp)

head(top_sync_counts_m)
top_sync_counts_m[which(top_sync_counts_m$snp == "scaffold1003_1161"),]

# Add lat, alt, pc1, pc2, ccrt, and ctmin data
tableS1<-read.table("~/Dropbox/st_andrews/montana_clines_MS/Manuscript/V7_R1/supplemental_information/TableS1_montana_pop_info_bioclimdat.tsv",
                    header=TRUE, stringsAsFactors=FALSE,sep="\t")
tableS1<-tableS1[tableS1$PoolGen==1,]
tableS1$LocationName<-gsub("Crested Butte","Crested_Butte",tableS1$LocationName)
head(tableS1)

snp<-"scaffold1003_1161"
snp_mod_res<-data.frame("snp"=unique(top_sync_counts_m$snp),
                        "Lat_e"=vector(length=length(unique(top_sync_counts_m$snp))),
                        "Lat_se"=vector(length=length(unique(top_sync_counts_m$snp))),
                        "Lat_p"=vector(length=length(unique(top_sync_counts_m$snp))),
                        "Alt_e"=vector(length=length(unique(top_sync_counts_m$snp))),
                        "Alt_se"=vector(length=length(unique(top_sync_counts_m$snp))),
                        "Alt_p"=vector(length=length(unique(top_sync_counts_m$snp))),
                        "var_e"=vector(length=length(unique(top_sync_counts_m$snp))),
                        "var_se"=vector(length=length(unique(top_sync_counts_m$snp))),
                        "var_p"=vector(length=length(unique(top_sync_counts_m$snp))))
snp_c_dat_all<-data.frame()
for(snp in unique(top_sync_counts_m$snp)){
  snp_c_dat<-top_sync_counts_m[which(top_sync_counts_m$snp==snp),]
  pop_dat<-tableS1[,c("LocationName","Lat","Alt_m","PC1_scaled","PC2_scaled","ctmin","ccrt")]
  snp_c_dat<-cbind(snp_c_dat,pop_dat[match(snp_c_dat$pop,pop_dat$LocationName),2:ncol(pop_dat)])
  snp_c_dat_all<-rbind(snp_c_dat_all,snp_c_dat)
  # Now fit a glm with quasibinomial error and lat + alt
  #summary(glm(cbind(maj_c,alt_c)~PC2_scaled,data=snp_c_dat,family="quasibinomial"))
  mod_s<-summary(glm(maj_c/(maj_c+alt_c)~Lat+Alt_m+PC2_scaled,data=snp_c_dat))
  # Save results from glm
  coefs<-mod_s$coefficients
  snp_mod_res[which(snp_mod_res$snp==snp),2:ncol(snp_mod_res)]<-c(coefs[2,c(1,2,4)],coefs[3,c(1,2,4)],coefs[4,c(1,2,4)])
}
nrow(snp_mod_res)
nrow(snp_mod_res[which(snp_mod_res$Lat_p<0.05),])
#snp_mod_res[which(snp_mod_res$Lat_p<0.05),]
nrow(snp_mod_res[which(snp_mod_res$Alt_p<0.05),])
#snp_mod_res[which(snp_mod_res$Lat_p<0.05),]

nrow(snp_mod_res[which(snp_mod_res$var_p<0.05),])
snp_mod_res[which(snp_mod_res$var_p<0.05),]

ggplot()+
  geom_point(data=snp_c_dat_all[which(snp_c_dat_all$snp %in% as.character(snp_mod_res$snp[which(snp_mod_res$Lat_p<0.05)])),],
             aes(x=Lat,y=maj_c/(maj_c+alt_c),fill=Alt_m),shape=21,size=4)+
  xlab("Latitude")+
  ylab("Major allele frequency")+
  scale_fill_continuous("Altitude")+
  facet_wrap(snp~.)+
  my.theme

ggplot()+
  geom_point(data=snp_c_dat_all[which(snp_c_dat_all$snp %in% as.character(snp_mod_res$snp[which(snp_mod_res$Lat_p<0.05)])),],
             aes(x=Lat,y=maj_c/(maj_c+alt_c),fill=Alt_m),shape=21,size=4)+
  xlab("Latitude")+
  ylab("Major allele frequency")+
  scale_fill_continuous("Altitude")+
  facet_wrap(snp~.)+
  my.theme


ggplot()+
  geom_point(data=snp_c_dat,aes(x=PC1_scaled,y=maj_c/(maj_c+alt_c)),fill="black",shape=21,size=4)


###
# Convert dmon annotation to dmelnames
dmongff<-read.table("~/projects/D_montana/montana_genome/D_montana_genome_freeze_v1.4_dvironlyred.gff",
                    header=FALSE,sep="\t")

dmongff<-cbind(dmongff[,c(1,2,3,4,5,6,7,8)],do.call(rbind,strsplit(as.character(dmongff$V9),";")))
dmongff$`1`<-gsub("ID=","",dmongff$`1`)
dmongff$`1`<-as.character(dmongff$`1`)
dmongff$dmelFB<-vector(length=nrow(dmongff))
head(dmongff)

dmelnames<-read.table("~/projects/D_montana/montana_genome/dvirgenes_FlyBase_dmel_IDs.txt",header=FALSE,sep="\t",fill=TRUE)
dmelnames<-dmelnames[which(!grepl("<newline>",dmelnames$V1)),]
head(dmelnames)
dmelnames$V1<-as.character(dmelnames$V1)
dmelnames$V2<-as.character(dmelnames$V2)

dmel2dvir<-read.table("~/projects/D_montana/montana_genome/dvirgenes_FlyBase_Fields_download.txt",fill=TRUE)
dmel2dvir$V1<-as.character(dmel2dvir$V1)
dmel2dvir$V3<-as.character(dmel2dvir$V3)
head(dmel2dvir)

dvirFB<-"FBgn0211248"
dmelFB<-"FBgn0031034"
for(dvirFB in unique(dmongff$`1`)){
  gene_name<-dmel2dvir$V3[which(dmel2dvir$V1==dvirFB)]
  dmelnames[which(dmelnames$V1==gene_name),]
  dmelFB<-dmelnames$V2[which(dmelnames$V1==gene_name)]
  # does it already exist
  if(length(unique(dmelFB))>1){
    # dvir gene has two different orthologs in dmel...what?!
    cat(dvirFB," - whaaaat!\n")
  }else if(length(unique(dmelFB))>0){
    dmelFB<-unique(dmelFB)
    if(dmelFB %in% dmongff$dmelFB){
      # How many
      i<-length(unique(dmongff$dmelFB[grep(dmelFB,dmongff$dmelFB)]))
      i<-i+1
      dmelFB<-paste(dmelFB,"_",i,sep="")
    }
    # If more than 2 scaffolds
    if(length(unique(as.character(dmongff$V1[which(dmongff$`1`==dvirFB)])))>1){
      i<-1
      for(scaff in as.character(unique(dmongff$V1[which(dmongff$`1`==dvirFB)]))){
        dmongff$dmelFB[which(dmongff$`1`==dvirFB & dmongff$V1 == scaff)]<-paste(unique(dmelFB),"_",i,sep="")
        i<-i+1
      }
    }else{
      dmongff$dmelFB[which(dmongff$`1`==dvirFB)]<-unique(dmelFB)
    }
  }else{
    dmongff$dmelFB[which(dmongff$`1`==dvirFB)]<-"NA"
  }
}

dmongff$dmelgene_id<-paste('gene_id "',dmongff$dmelFB,'";',sep="")
head(dmongff,n=100)
dmongff<-dmongff[which(!(dmongff$dmelFB=="NA")),]
dmongff[grepl("FBgn0031034",dmongff$dmelFB),]

dmongff$V1<-gsub("-size.*","",dmongff$V1)
head(dmongff[,c(1,2,3,4,5,6,7,8,12)])
write.table(dmongff[,c(1,2,3,4,5,6,7,8,12)],"~/projects/D_montana/montana_genome/D_montana_genome_freeze_v1.4_dmel.gtf",
            col.names = FALSE,row.names=FALSE,quote=FALSE,sep="\t")


snp_counts<-read.table("~/Dropbox/st_andrews/montana_clines_MS/Manuscript/V12_R2/chrom_snp_counts.csv",header=TRUE,sep=",")
head(snp_counts)

total_snps<-sum(snp_counts$n_snps[which(snp_counts$variable=="Total" & snp_counts$chromosome %in% c("Autosomes","X"))])
snp_counts_tot<-snp_counts[which(snp_counts$variable=="Total" & snp_counts$chromosome != "Autosomes"),]
snp_counts_tot$prop<-snp_counts_tot$n_snps/total_snps
snp_counts_tot

snp_counts$expected<-vector(length=nrow(snp_counts))
for(chrom in unique(snp_counts_tot$chromosome)){
  for(var in unique(snp_counts$variable)){
    total_top<-sum(snp_counts$n_snps[which(snp_counts$variable==var & snp_counts$chromosome != "Autosomes")])
    snp_counts$expected[which(snp_counts$variable==var & snp_counts$chromosome != "Autosomes")]<-round(snp_counts_tot$prop*total_top,2)
  }
}

snp_counts

ggplot()+
  geom_bar(data=snp_counts[which(snp_counts$variable!="Total"),],
           aes(x=chromosome,y=n_snps,fill=variable),stat="identity",position="dodge",colour="black")+
  geom_point(data=snp_counts[which(snp_counts$variable!="Total" & snp_counts$chromosome %in% c("LG2","LG3","LG4","LG5","X")),],
           aes(x=chromosome,y=expected,group=variable),position=position_dodge(0.95),colour="black")+
  scale_fill_manual("Environmental\nvariable",values=c("white","grey90","grey80","grey70"))+
  ylab("Number of SNPs")+
  xlab("Chromosome")+
  geom_text(data=snp_counts[which(snp_counts$variable=="Total"),],
            aes(x=chromosome,y=-100,label=n_snps))+
  geom_text(data=snp_counts[which(snp_counts$variable=="Total" & snp_counts$chromosome %in% c("LG2","LG3","LG4","LG5","X")),],
              aes(x=chromosome,y=-300,label=round((n_snps)/total_snps,2)))+
  my.theme+theme(axis.text=element_text(size=16),
                 axis.title=element_text(size=16))
```
