## Supplementary material for "Cold adaptation drives population genomic divergence in the ecological specialist, *Drosophila montana*": montana_clinalpops_D.R

```
##
# D. montana project clinal populations
# QBGLM data analysis
# Last Modified: Feb 2017
##
# Clean environment
rm(list = ls(all = TRUE))

#
dir<-"~/projects/D_montana/clinal_population_genomics/data/SNPs"
# Load libraries
library(ggplot2)
library(qvalue)
library(reshape)
library(reshape2)
se <- function(x) {sqrt(var(x, na.rm = TRUE))/sqrt(length(x))}
#citation("ggmap")
source("~/projects/RData/RScripts/ggplot_theme.R")
options(scipen=10)


#
# Read in the Tajima's D data.
# ####
tajD_dat<-read.table(paste(dir,"/montana_clines_PoolSNP_thetas_10000-5000.D",sep=""),header=FALSE)
colnames(tajD_dat)<-c("chr","winpos","Oulanka","Korpilahti","Ashford","Seward","Crested_Butte","Terrace")
tajD_dat<-tajD_dat[,1:8]
# Remove the "size" from scaffold names
tajD_dat$chr<-gsub("-size.*","",tajD_dat$chr)

head(tajD_dat)

# Melt the data
tajD_dat_m<-melt(data = tajD_dat, measure.vars = c("Oulanka","Korpilahti","Ashford","Seward","Crested_Butte","Terrace"))
colnames(tajD_dat_m)<-c("chr","pos","pop","TajD")
head(tajD_dat_m)


# D distribution
ggplot()+
  geom_histogram(data=tajD_dat_m,aes(TajD))+
  geom_vline(xintercept = 0,colour="grey50")+
  xlab("Tajima's D")+
  facet_grid(pop~.)+
  my.theme+theme(strip.text.y = element_text(angle=0))

ggplot()+
  geom_hline(yintercept = 0,colour="grey50")+
  geom_boxplot(data=tajD_dat_m,
               aes(pop,TajD))+
  ylab("Tajima's D")+
  xlab("")+
  my.theme+theme(axis.text = element_text(size=10),
                      axis.title.y = element_text(size=14),
                      plot.title = element_text(size=14))

tapply(tajD_dat_m$TajD,INDEX=list(tajD_dat_m$pop),mean,na.rm=TRUE)
tapply(tajD_dat_m$TajD,INDEX=list(tajD_dat_m$pop),sd,na.rm=TRUE)
# Oukanka, Latitude: 66.67 Alt: 337
# Korpilahti, Latitude: 62.33 Alt: 133
# Ashford, Latitude: 46.75 Alt: 573
# Seward, Latitude: 60.15 Alt: 35
# Crested Butte, Latitude: 38.9 Alt: 2900 
# Terrace, Latitude: 54.45 Alt: 217
lat<-c(66.67,62.33,46.75,60.15,38.9,54.45)
alt<-c(335,133,573,35,2900,217)
summary_dat<-data.frame("lat"=lat,
                        "alt"=alt,
                        "m_tajD"=tapply(tajD_dat_m$TajD,INDEX=list(tajD_dat_m$pop),mean,na.rm=TRUE),
                        "sd_tajD"=tapply(tajD_dat_m$TajD,INDEX=list(tajD_dat_m$pop),sd,na.rm=TRUE))
summary_dat$pop<-rownames(summary_dat)
head(summary_dat)
ggplot()+
  geom_point(data=summary_dat,
               aes(alt,m_tajD),size=3)+
  geom_errorbar(data=summary_dat,
             aes(alt,ymin=m_tajD-sd_tajD,ymax=m_tajD+sd_tajD),width=0)+
  ylab("Tajima's D")+
  xlab("")+
  my.theme+theme(axis.text = element_text(size=14),
                 axis.title.y = element_text(size=18),
                 plot.title = element_text(size=14),
                 plot.background = element_rect(colour="black"),
                 panel.background = element_rect(colour="black"))

cor.test(summary_dat$lat,summary_dat$m_tajD,method="spearman")
cor.test(summary_dat$lat[summary_dat$pop!="Crested_Butte"],summary_dat$m_tajD[summary_dat$pop!="Crested_Butte"],
         method="spearman")
cor.test(summary_dat$alt,summary_dat$m_tajD,method="spearman")
cor.test(summary_dat$alt[summary_dat$pop!="Crested_Butte"],summary_dat$m_tajD[summary_dat$pop!="Crested_Butte"],
         method="spearman")

# Convert the D data to linkage group data
# Scaffold sizes from genome freeze v.1.4
genome_dir<-"~/projects/D_montana/montana_genome/"

scaffdat<-read.table(paste(genome_dir,"/montana_scaffsizes.tab",sep=""),
                     header=FALSE,sep=",")
colnames(scaffdat)<-c("chr","size")
nrow(scaffdat)
head(scaffdat[scaffdat$chr == "scaffold3615",])

# Load the v1 genome linkage groups
LGdat1<-read.table(paste(genome_dir,"markers_and_positions_v2.1.csv",sep=""),header=TRUE,sep="\t")
colnames(LGdat1)<-c("scaff","linkage_gr","lkg_pos","lg","lgmrk")
head(LGdat1[LGdat1$scaff=="scaffold3615",])
head(LGdat1)
# Load the v1.4 genome linkage groups
LGdat1.4<-read.table(paste(genome_dir,"vennys_map_info_v1.4.txt",sep=""),header=TRUE,sep="\t")
LGdat1.4$scaf_name<-gsub("-size.*","",LGdat1.4$scaf_name)
colnames(LGdat1.4)<-c("scaff","linkage_gr","size")
LGdat1.4$linkage_gr<-paste("Chr ",LGdat1.4$linkage_gr,sep="")
LGdat1.4$lkg_pos<-vector(length=nrow(LGdat1.4))
# Add scaffold positions on LGs to LGdat1.4
for(row in 1:nrow(LGdat1.4)){
  scf<-LGdat1.4$scaff[row]
  LGdat1.4$lkg_pos[row]<-LGdat1$lkg_pos[LGdat1$scaff==scf]
}
head(LGdat1.4[LGdat1.4$scaff=="scaffold3615",])
LGdat1.4<-LGdat1.4[order(LGdat1.4$linkage_gr,LGdat1.4$lkg_pos),]

## Add linkage groups and positions to the data set
head(LGdat1.4)
head(scaffdat)
head(tajD_dat)
tajD_dat$lg<-vector(length=nrow(tajD_dat))
tajD_dat$lgpos<-vector(length=nrow(tajD_dat))
tajD_dat$scfsize<-vector(length=nrow(tajD_dat))
tajD_dat_lg<-data.frame()
for(scaff in unique(tajD_dat$chr)){
  scaff_data_sub<-tajD_dat[tajD_dat$chr==scaff,]
  lg<-as.character(LGdat1.4$linkage_gr[LGdat1.4$scaff==scaff])
  # If lg is null, the scaffold is not anchored to a linkage group.
  # Call it "Unplaced"
  if(identical(lg,character(0))){
    lg<-"Unplaced"
    #print(lg)
    scaff_data_sub$lg<-rep(lg,nrow(scaff_data_sub))
    scaff_data_sub$lgpos<-rep(NA,nrow(scaff_data_sub))
    scaff_data_sub$scfsize<-rep(scaffdat$size[scaffdat$chr==scaff],
                               nrow(scaff_data_sub))
    tajD_dat_lg<-rbind(tajD_dat_lg,scaff_data_sub)
  }else{
    scaff_data_sub$lg<-rep(lg,nrow(scaff_data_sub))
    scaff_data_sub$lgpos<-rep(LGdat1.4$lkg_pos[LGdat1.4$scaff==scaff],
                            nrow(scaff_data_sub))
    scaff_data_sub$scfsize<-rep(LGdat1.4$size[LGdat1.4$scaff==scaff],
                            nrow(scaff_data_sub))
    tajD_dat_lg<-rbind(tajD_dat_lg,scaff_data_sub)
  }
}
head(tajD_dat_lg)
# Subset to "placed" scaffolds
tajD_dat_lg_pl<-tajD_dat_lg[grep("Unplaced",tajD_dat_lg$lg,invert=TRUE),]
tajD_dat_lg_pl<-tajD_dat_lg_pl[
  order(tajD_dat_lg_pl$lg,tajD_dat_lg_pl$lgpos,tajD_dat_lg_pl$winpos),]
head(tajD_dat_lg_pl)
## Add an absoulte position for each linkage group
tajD_dat_lg_pl$lg_win_pos<-vector(length=nrow(tajD_dat_lg_pl))
tajD_dat_lg_pl_scaff_data<-data.frame()
for(LG in unique(tajD_dat_lg_pl$lg)){
  # Restart totpos at 0
  totpos<-0
  scaff_data_new_sub2<-tajD_dat_lg_pl[tajD_dat_lg_pl$lg == LG,]
  # Last scaffold length is 0 for the first scaffold on the LG
  last_scaff_l<-0
  # Looping through all the scaffolds on the LG ensures we can see the 
  # regions where we don't have enough SNPs
  for(scaff in LGdat1.4$scaff[LGdat1.4$linkage_gr==LG]){
    scaff_data_new_sub3<-scaff_data_new_sub2[scaff_data_new_sub2$chr == scaff,]
    # Get scaffold size from the scaffdat table
    curr_scaff_l<-scaffdat$size[scaffdat$chr == scaff]
    #print(scaff)
    #print(curr_scaff_l)
    # Get a new position for each SNP by:
    # lg_snp_pos <- startpos + last_scaff_l + scaff_snp_pos
    scaff_data_new_sub3$lg_win_pos<-scaff_data_new_sub3$winpos+totpos
    tajD_dat_lg_pl_scaff_data<-rbind(tajD_dat_lg_pl_scaff_data,scaff_data_new_sub3)
    last_scaff_l<-curr_scaff_l
    totpos <- totpos+last_scaff_l
  }
}
tajD_dat_lg_pl_scaff_data$lg<-factor(tajD_dat_lg_pl_scaff_data$lg)
head(tajD_dat_lg_pl_scaff_data)
tajD_dat_lg_pl_scaff_data_m<-melt(tajD_dat_lg_pl_scaff_data,
                                measure.vars =  c("Oulanka","Korpilahti","Ashford","Seward","Crested_Butte","Terrace"))
colnames(tajD_dat_lg_pl_scaff_data_m)<-c("chr","winpos","lg","lgpos","scfsize","lg_win_pos","pop","TajD")
head(tajD_dat_lg_pl_scaff_data_m)

# Plot TajD overall
# There are big gaps because I remove scaffolds that are very short, these are problematic.
D_plot<-ggplot()+
  geom_hline(yintercept = 0,colour="grey")+
  geom_line(data=tajD_dat_lg_pl_scaff_data_m,
            aes(lg_win_pos/1000,TajD,colour=chr,linetype=pop))+
  scale_color_manual("",values=rep(c("black","red"),
                                   length.out=length(unique(tajD_dat_lg_pl_scaff_data_m$chr))),
                     guide=FALSE)+
  scale_linetype_discrete("")+
  ylab("Tajima's D")+
  xlab("Position (kb)")+
  #ylim(-2,2)+
  facet_grid(lg~.)+
  my.theme+
  theme(axis.text.y = element_text(size=10),
        axis.text.x = element_text(size=10),
        axis.title = element_text(size=10),
        strip.text.y = element_text(size=10,angle=0),
        plot.title = element_text(hjust=0,size=10))
D_plot
```
