## Supplementary material for "Cold adaptation drives population genomic divergence in the ecological specialist, *Drosophila montana*": montana_clinalpops_fst.R

```
##
# D. montana project clinal populations
# FST data analysis
# Last Modified: Apr 2020
##
# Clean environment
rm(list = ls(all = TRUE))

#
dir<-"~/projects/D_montana/clinal_population_genomics/data/SNPs"
# Load libraries
library(ggplot2)
library(reshape)
se <- function(x) {sqrt(var(x, na.rm = TRUE))/sqrt(length(x))}
#citation("ggmap")
source("~/projects/RData/RScripts/ggplot_theme.R")
options(scipen=10)


#
# Read in the FST data.
# ####
fst_dat<-read.table(paste(dir,"/sync2fst.out",sep=""),header=TRUE)
head(fst_dat)

map_data<-data.frame("id"=c("Pool1","Pool2","Pool3","Pool4","Pool5","Pool6"),
                     "pop"=c("Oulanka","Korpilahti","Ashford","Seward","Crested_Butte","Terrace"))

# 

ids<-c("Pool1","Pool2","Pool3","Pool4","Pool5","Pool6")
id<-"Pool1"
fst_chr_dat<-fst_dat[,c(1,2,3)]
head(fst_chr_dat)
for(id in ids){
  fst_sub<-fst_dat[grepl(id,colnames(fst_dat))]
  mean_fst<-apply(fst_sub,MARGIN = 1,mean,na.rm=TRUE)
  fst_chr_dat[[as.character(map_data$pop[which(map_data$id==id)])]]<-mean_fst
}
head(fst_chr_dat)
write.table(fst_chr_dat,paste(dir,"/pop_fst.tab",sep=""),
            row.names = FALSE,col.names=TRUE,quote=FALSE,sep="\t")


#---------------------------------------------------------------------#
fst_chr_dat<-read.table(paste(dir,"/pop_fst.tab",sep=""),header=TRUE,sep="\t")
fst_chr_dat$chr<-gsub("-size.*","",fst_chr_dat$chr)

# Convert the fst data to linkage group data
# Scaffold sizes from genome freeze v.1.4
genome_dir<-"~/projects/D_montana/montana_genome/"

scaffdat<-read.table(paste(genome_dir,"/montana_scaffsizes.tab",sep=""),
                     header=FALSE,sep=",")
colnames(scaffdat)<-c("chr","size")
nrow(scaffdat)
head(scaffdat[scaffdat$chr == "scaffold3615",])

# Load the v1 genome linkage groups
LGdat1<-read.table(paste(genome_dir,"markers_and_positions_v2.1.csv",sep=""),header=TRUE,sep="\t")
colnames(LGdat1)<-c("scaff","linkage_gr","lkg_pos","lg","lgmrk")
head(LGdat1[LGdat1$scaff=="scaffold3615",])
head(LGdat1)
# Load the v1.4 genome linkage groups
LGdat1.4<-read.table(paste(genome_dir,"vennys_map_info_v1.4.txt",sep=""),header=TRUE,sep="\t")
LGdat1.4$scaf_name<-gsub("-size.*","",LGdat1.4$scaf_name)
colnames(LGdat1.4)<-c("scaff","linkage_gr","size")
LGdat1.4$linkage_gr<-paste("Chr ",LGdat1.4$linkage_gr,sep="")
LGdat1.4$lkg_pos<-vector(length=nrow(LGdat1.4))
# Add scaffold positions on LGs to LGdat1.4
for(row in 1:nrow(LGdat1.4)){
  scf<-LGdat1.4$scaff[row]
  LGdat1.4$lkg_pos[row]<-LGdat1$lkg_pos[LGdat1$scaff==scf]
}
head(LGdat1.4[LGdat1.4$scaff=="scaffold3615",])
LGdat1.4<-LGdat1.4[order(LGdat1.4$linkage_gr,LGdat1.4$lkg_pos),]

## Add linkage groups and positions to the data set
head(LGdat1.4)
head(scaffdat)
head(fst_chr_dat)
fst_chr_dat$lg<-vector(length=nrow(fst_chr_dat))
fst_chr_dat$lgpos<-vector(length=nrow(fst_chr_dat))
fst_chr_dat$scfsize<-vector(length=nrow(fst_chr_dat))
fst_chr_dat_lg<-data.frame()
for(scaff in unique(fst_chr_dat$chr)){
  scaff_data_sub<-fst_chr_dat[fst_chr_dat$chr==scaff,]
  lg<-as.character(LGdat1.4$linkage_gr[LGdat1.4$scaff==scaff])
  # If lg is null, the scaffold is not anchored to a linkage group.
  # Call it "Unplaced"
  if(identical(lg,character(0))){
    lg<-"Unplaced"
    #print(lg)
    scaff_data_sub$lg<-rep(lg,nrow(scaff_data_sub))
    scaff_data_sub$lgpos<-rep(NA,nrow(scaff_data_sub))
    scaff_data_sub$scfsize<-rep(scaffdat$size[scaffdat$chr==scaff],
                                nrow(scaff_data_sub))
    fst_chr_dat_lg<-rbind(fst_chr_dat_lg,scaff_data_sub)
  }else{
    scaff_data_sub$lg<-rep(lg,nrow(scaff_data_sub))
    scaff_data_sub$lgpos<-rep(LGdat1.4$lkg_pos[LGdat1.4$scaff==scaff],
                              nrow(scaff_data_sub))
    scaff_data_sub$scfsize<-rep(LGdat1.4$size[LGdat1.4$scaff==scaff],
                                nrow(scaff_data_sub))
    fst_chr_dat_lg<-rbind(fst_chr_dat_lg,scaff_data_sub)
  }
}

# Limit to placed SNPs
fst_chr_dat_lg<-fst_chr_dat_lg[which(fst_chr_dat_lg$lg!="Unplaced"),]

head(fst_chr_dat_lg)
nrow(fst_chr_dat_lg)

fst_chr_dat_lg_m<-melt(fst_chr_dat_lg,
                       measure.vars = c("Oulanka","Korpilahti","Ashford","Seward","Crested_Butte","Terrace"))
ggplot()+
  geom_histogram(data=fst_chr_dat_lg_m,aes(x=value))+
  facet_grid(lg~variable)

round(tapply(fst_chr_dat_lg_m$value,INDEX = list(fst_chr_dat_lg_m$lg,fst_chr_dat_lg_m$variable),mean),4)
round(tapply(fst_chr_dat_lg_m$value,INDEX = list(fst_chr_dat_lg_m$lg,fst_chr_dat_lg_m$variable),sd),4)
round(tapply(fst_chr_dat_lg_m$value,INDEX = list(fst_chr_dat_lg_m$lg,fst_chr_dat_lg_m$variable),length),2)

# Re-sampling SNPs (w/ replacement) to get 95% CIs
head(fst_chr_dat_lg)
n<-100
lg<-"Chr 2"
boot_ci<-matrix(ncol=7,nrow=5)
boot_ci[,1]<-unique(fst_chr_dat_lg$lg)
for(lg in unique(fst_chr_dat_lg$lg)){
  boot_vals<-matrix(nrow=n,ncol=6)
  lg_dat<-fst_chr_dat_lg[which(fst_chr_dat_lg$lg==lg),]
  n_snps<-nrow(lg_dat)
  rows<-1:nrow(lg_dat)
  for(boot in 1:n){
    boot_rows<-sample(rows,n_snps,replace = TRUE)
    boot_dat<-lg_dat[boot_rows,]
    boot_dat_m<-melt(boot_dat,measure.vars = c("Oulanka","Korpilahti","Ashford","Seward","Crested_Butte","Terrace"))
    boot_vals[boot,]<-tapply(boot_dat_m$value,INDEX = list(boot_dat_m$variable),mean)
  }
  ci_upr<-round(apply(boot_vals,MARGIN = 2,quantile,probs=0.975),4)
  ci_lwr<-round(apply(boot_vals,MARGIN = 2,quantile,probs=0.025),4)
  boot_ci[which(boot_ci[,1]==lg),c(2:7)]<-paste(ci_lwr,", ",ci_upr,sep="")
}
boot_ci
  

anova(lm(value~lg+variable,data=fst_chr_dat_lg_m))


# 

chrom_fst<-read.table("~/Dropbox/st_andrews/montana_clines_MS/Manuscript/V12_R2/chrom_fst.csv",header=TRUE,sep=",")
chrom_fst$population<-factor(chrom_fst$population,levels=c("Crested Butte","Ashford","Terrace","Seward","Korpilahti","Oulanka"))
head(chrom_fst)

ggplot()+
  geom_bar(data=chrom_fst,
           aes(x=chromosome,y=fst,fill=population),stat="identity",position=position_dodge(0.9),colour="black")+
  scale_fill_grey("Popoulation",start = 1,end = 0.5)+
  geom_errorbar(data=chrom_fst,
           aes(x=chromosome,ymin=ci_lwr,ymax=ci_upr,group=population),width=0.5,position=position_dodge(0.9),colour="black")+
  ylab(expression(paste(F[st])))+
  xlab("Chromosome")+
  my.theme+theme(axis.text=element_text(size=16),
                 axis.title=element_text(size=16))
```
