## Supplementary figures and images for "Cold adaptation drives population genomic divergence in the ecological specialist, *Drosophila montana*"

### figureS2.png

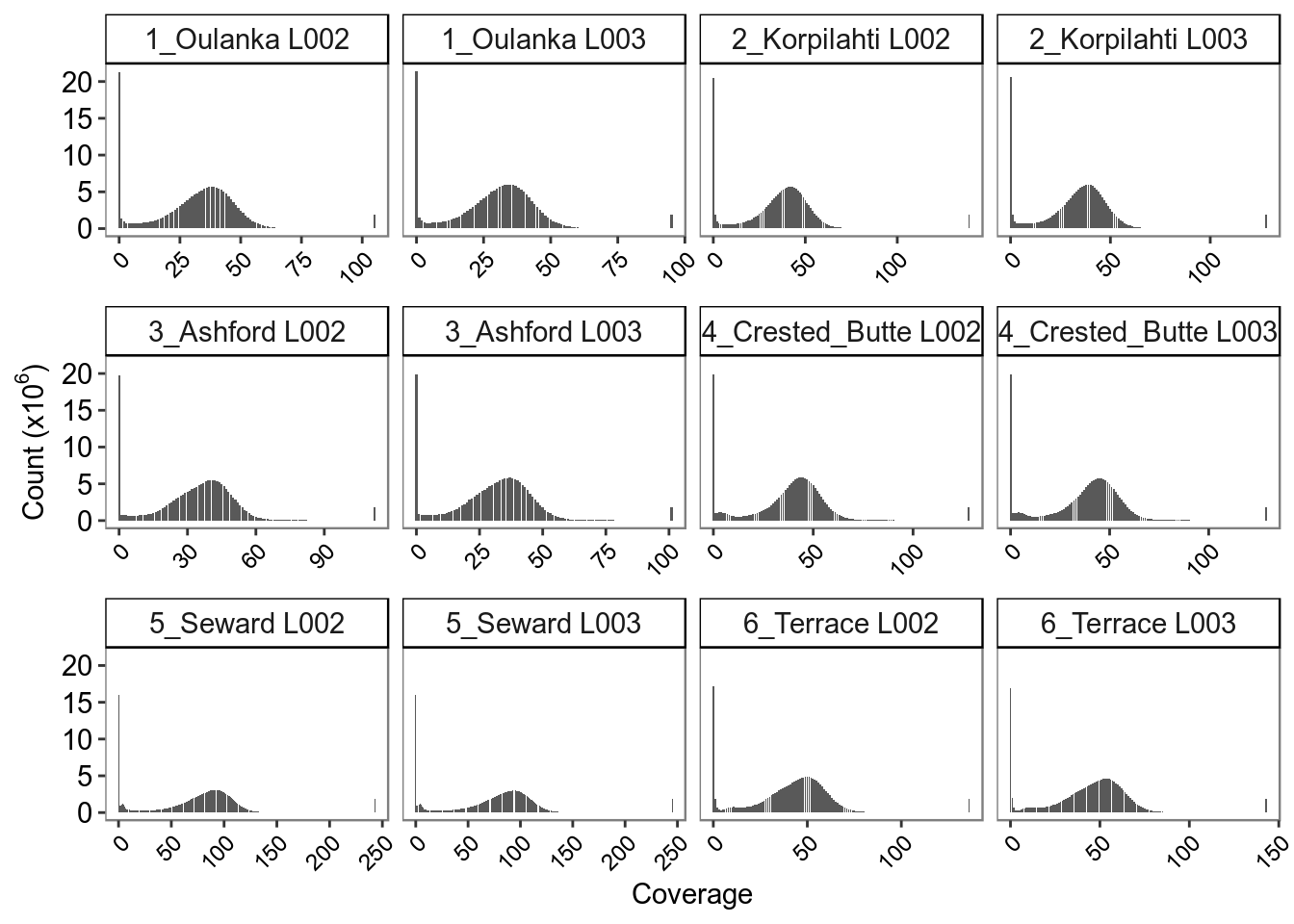

### figureS3.png

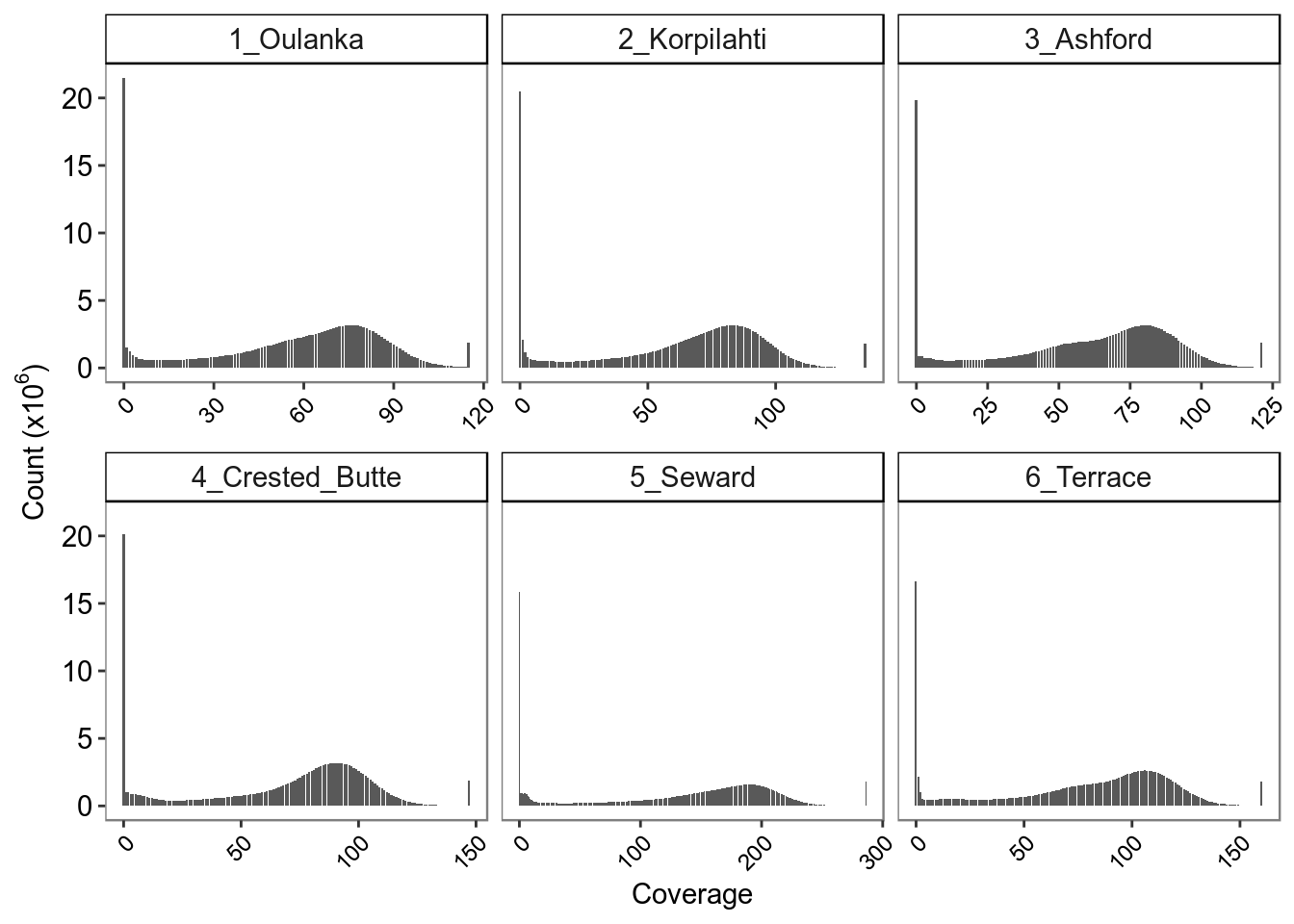

### figureS4.png

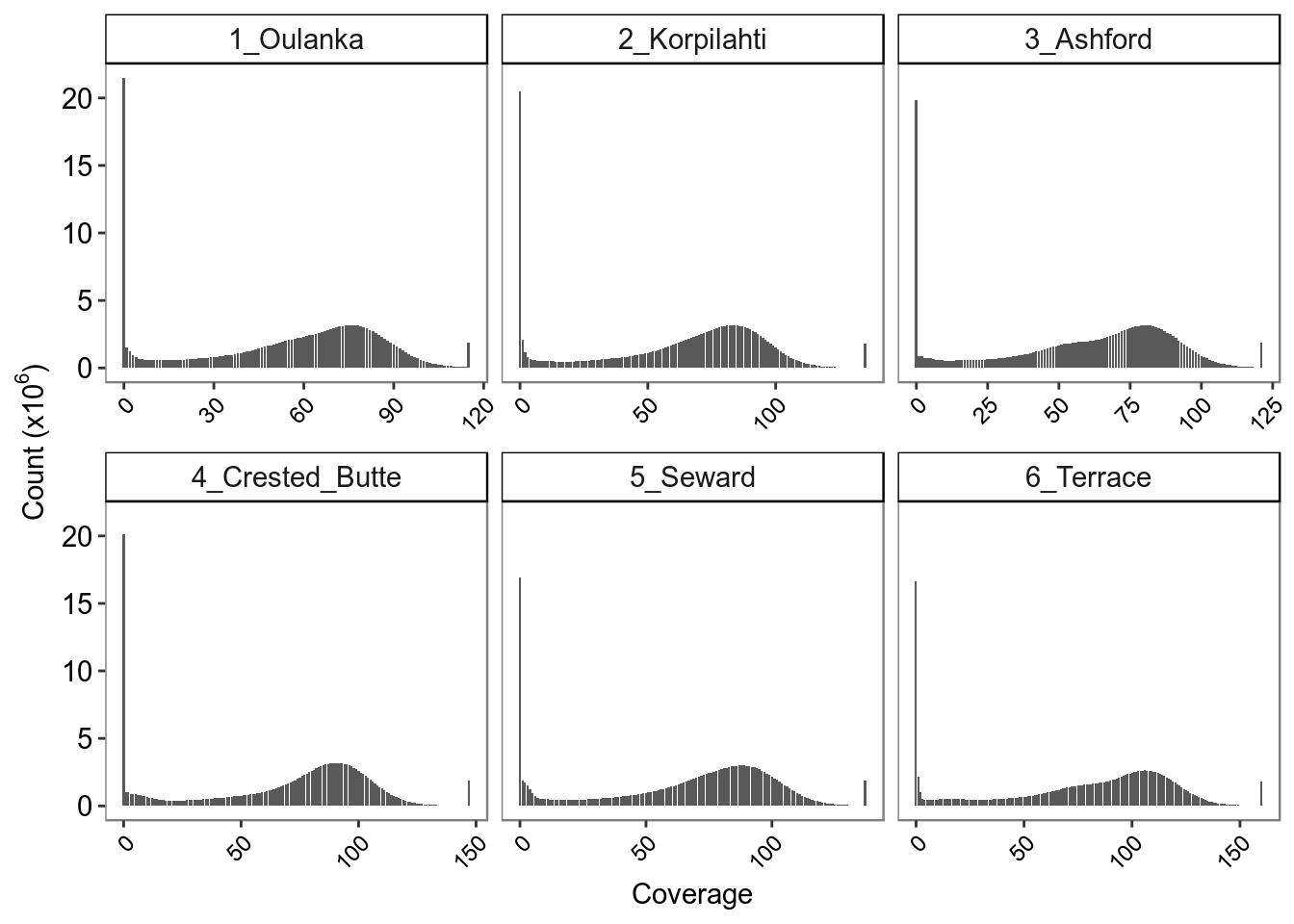

### figureS4A.pdf

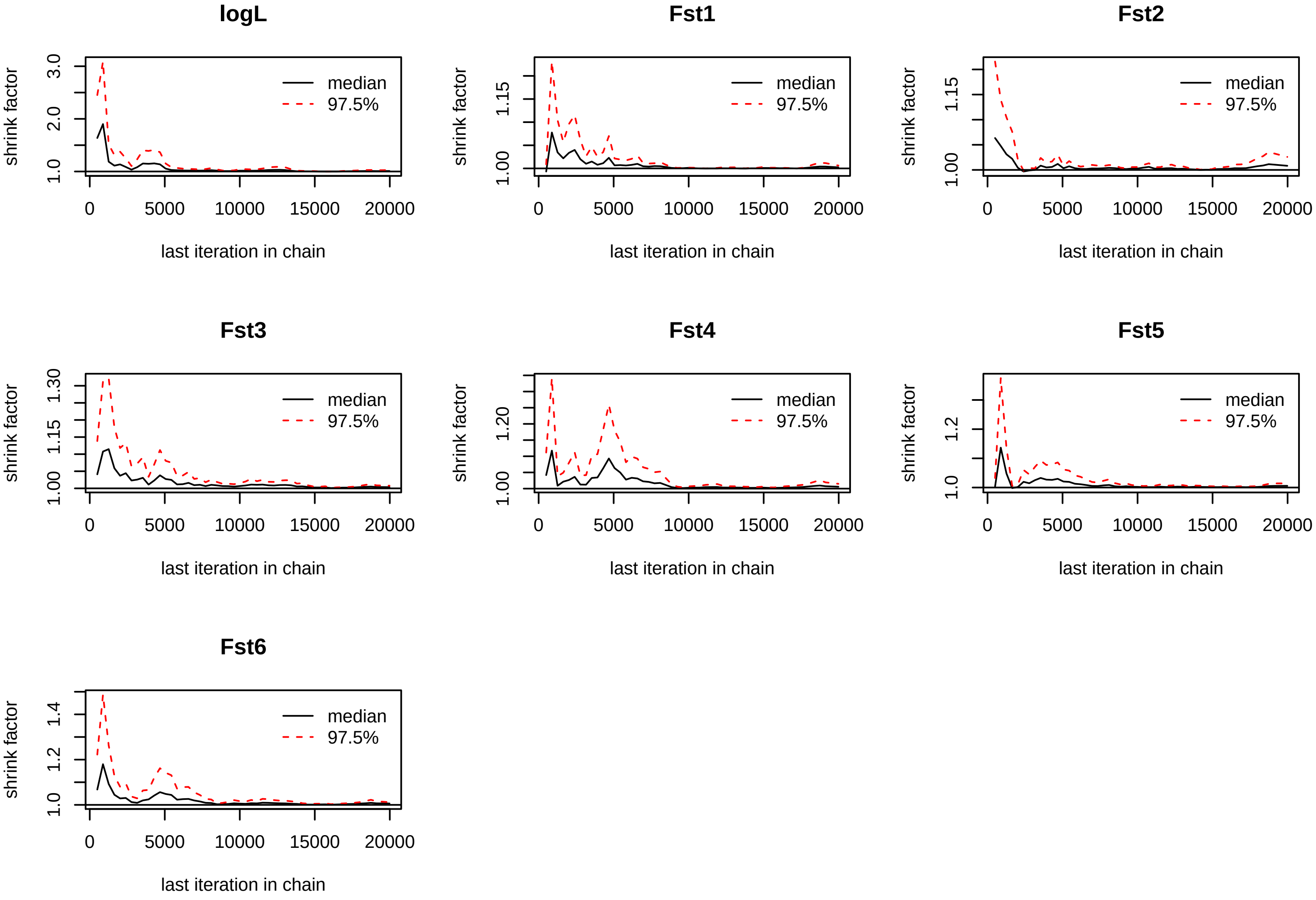

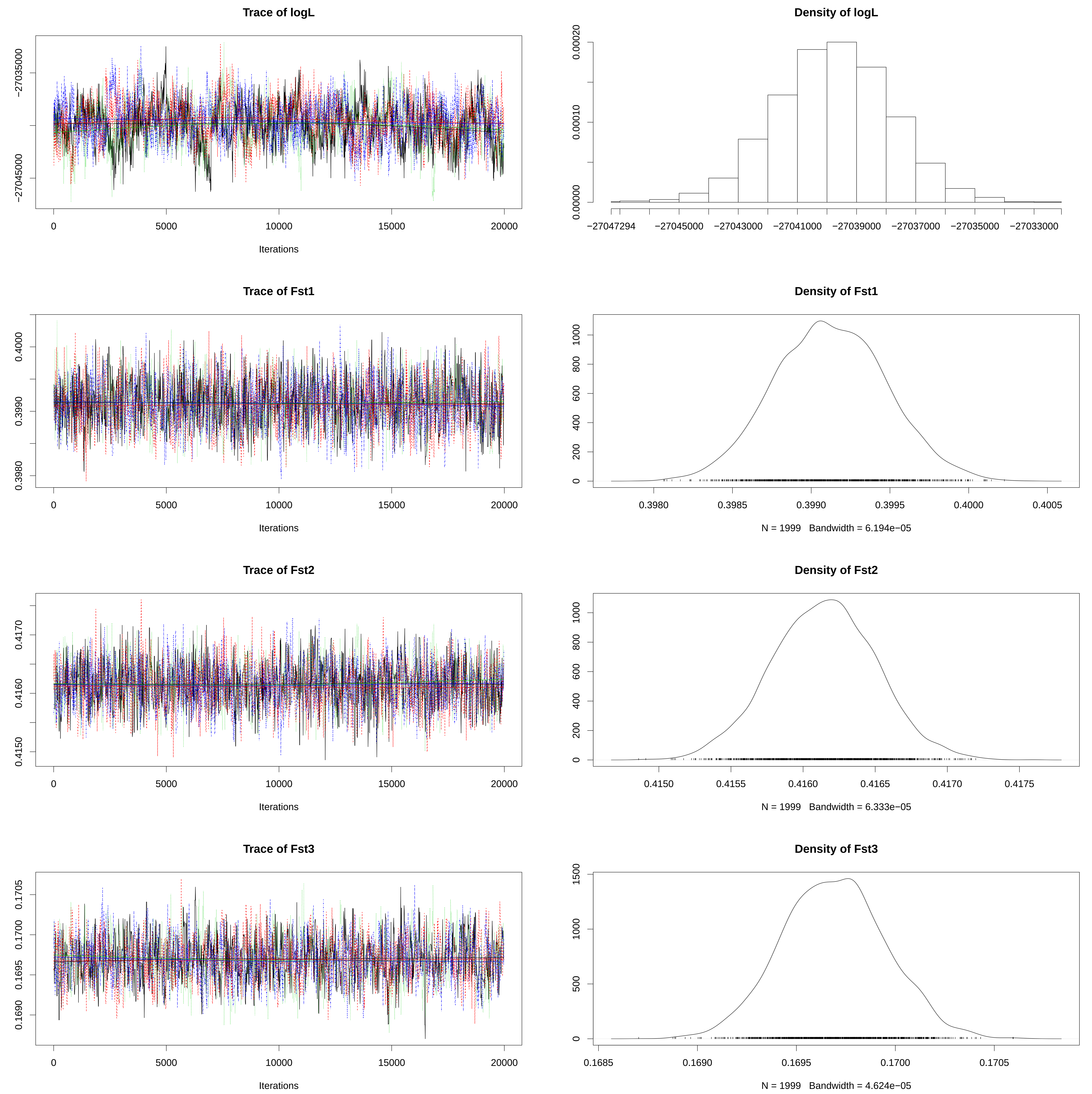

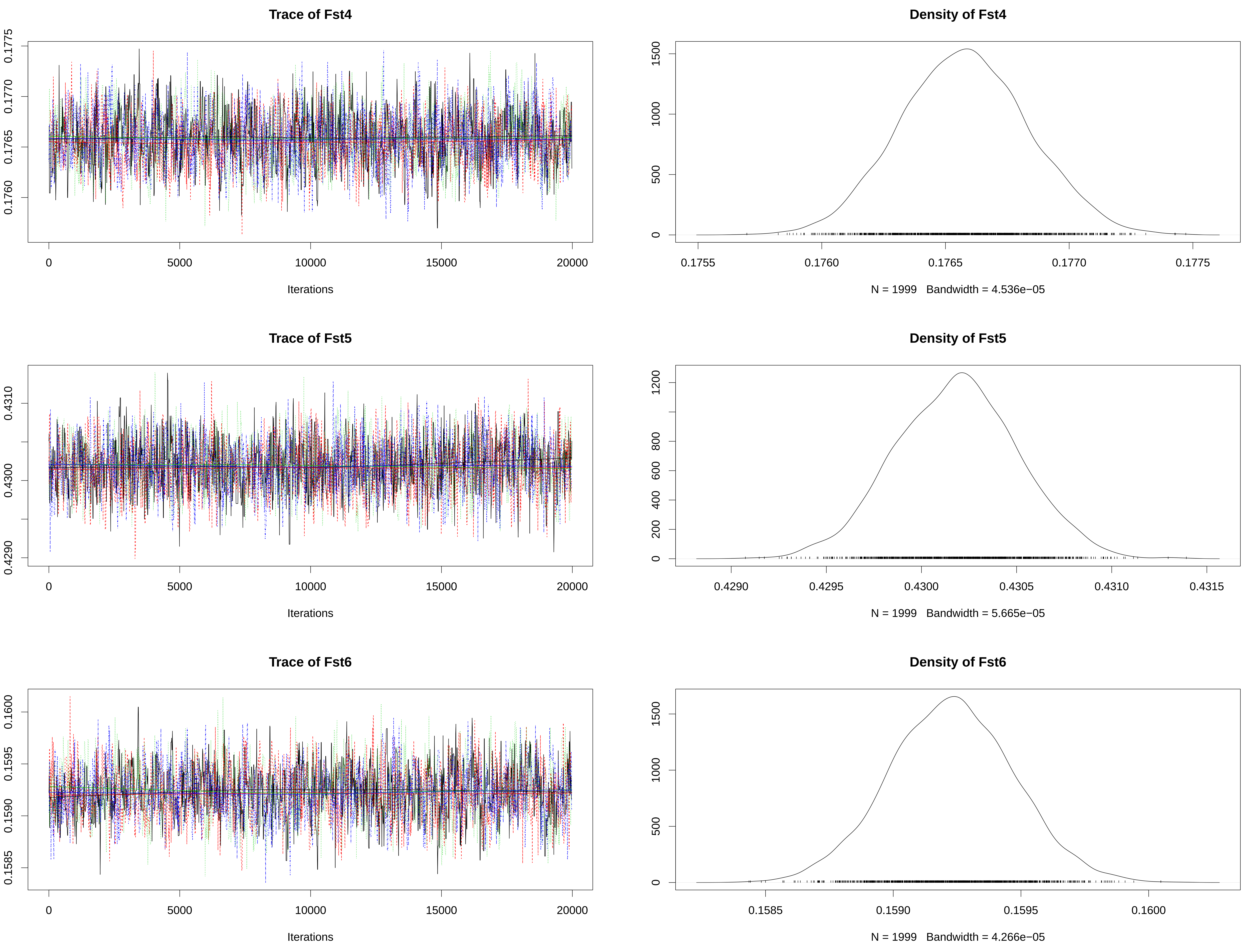

### figureS4B.pdf

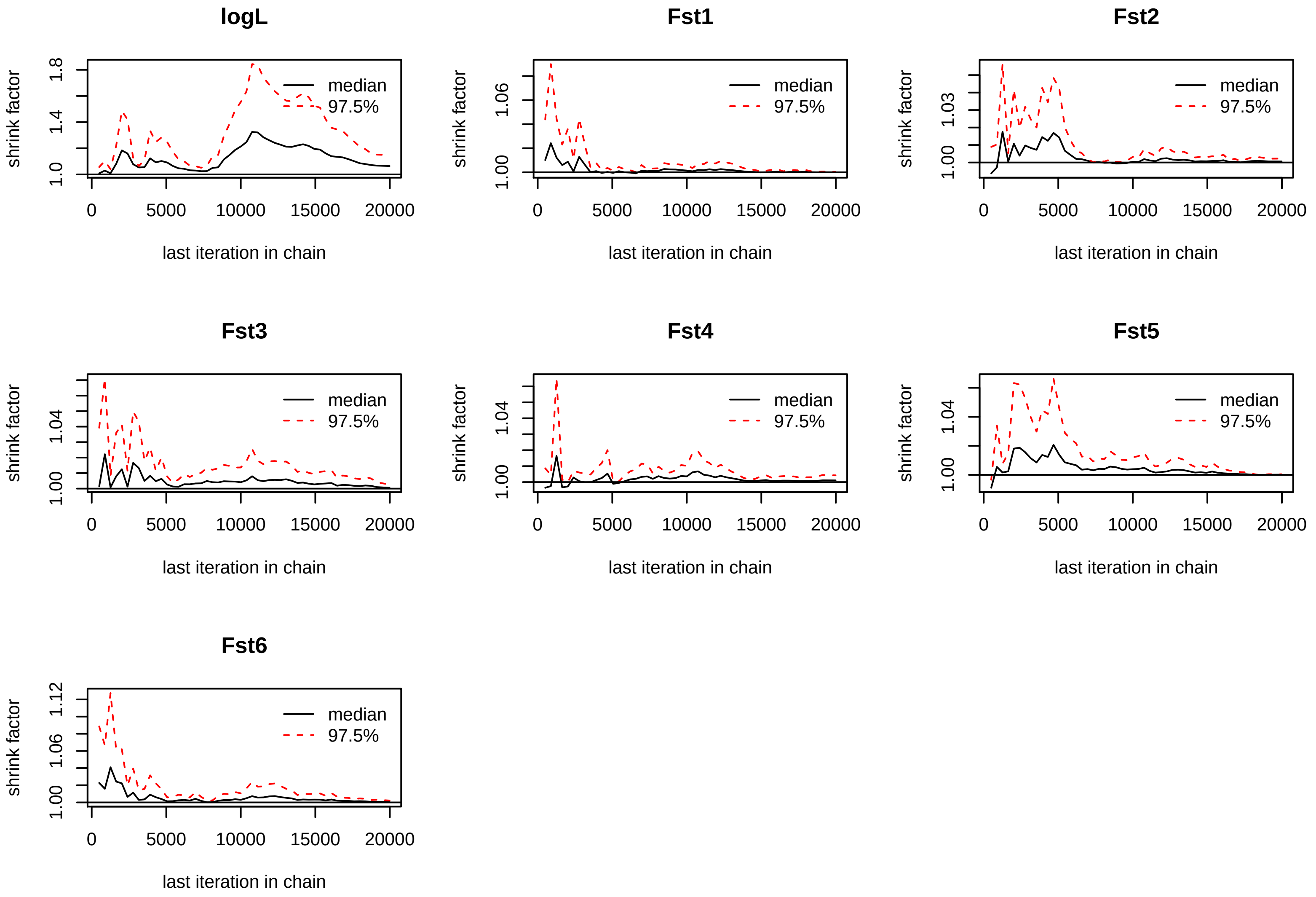

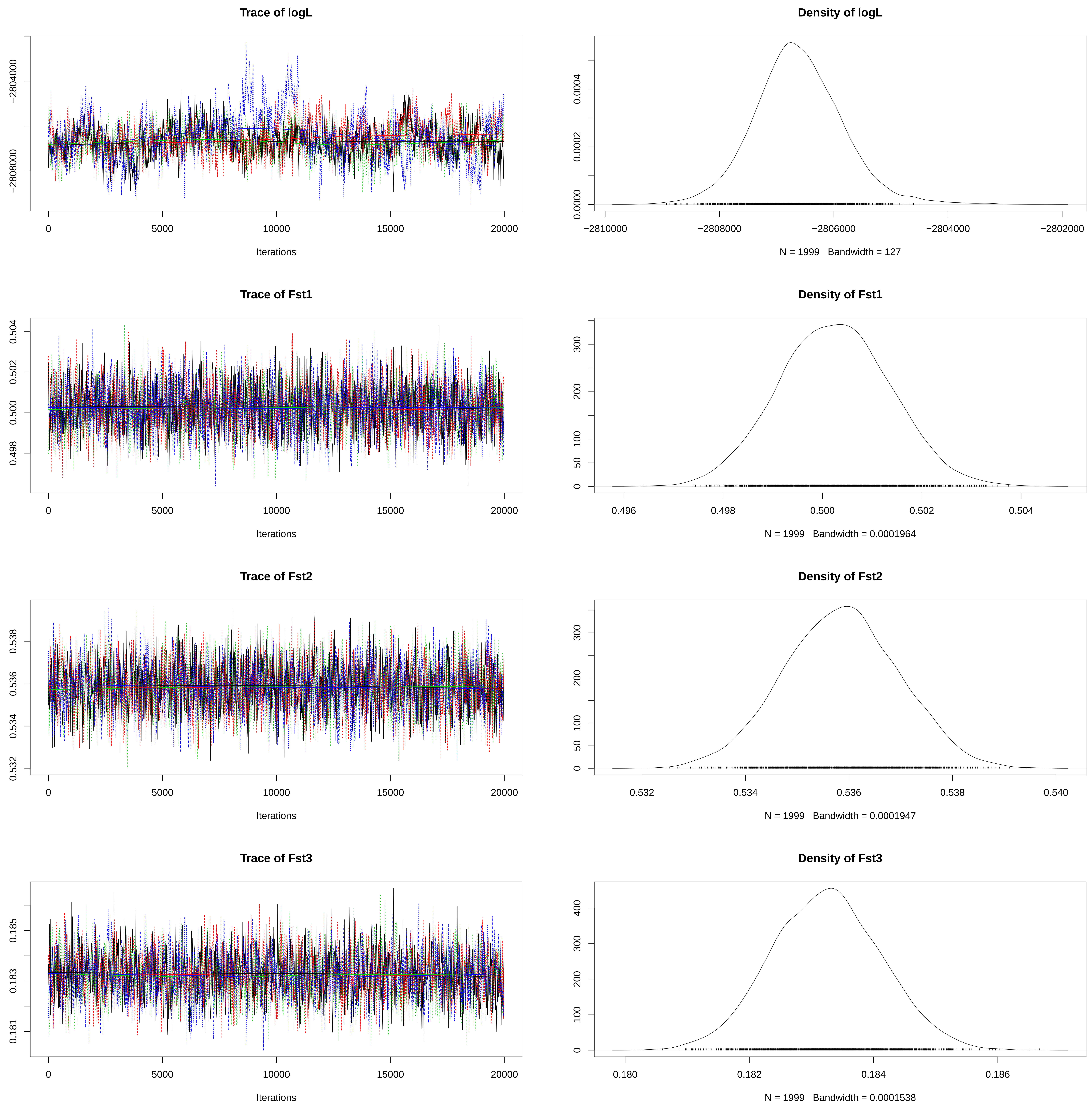

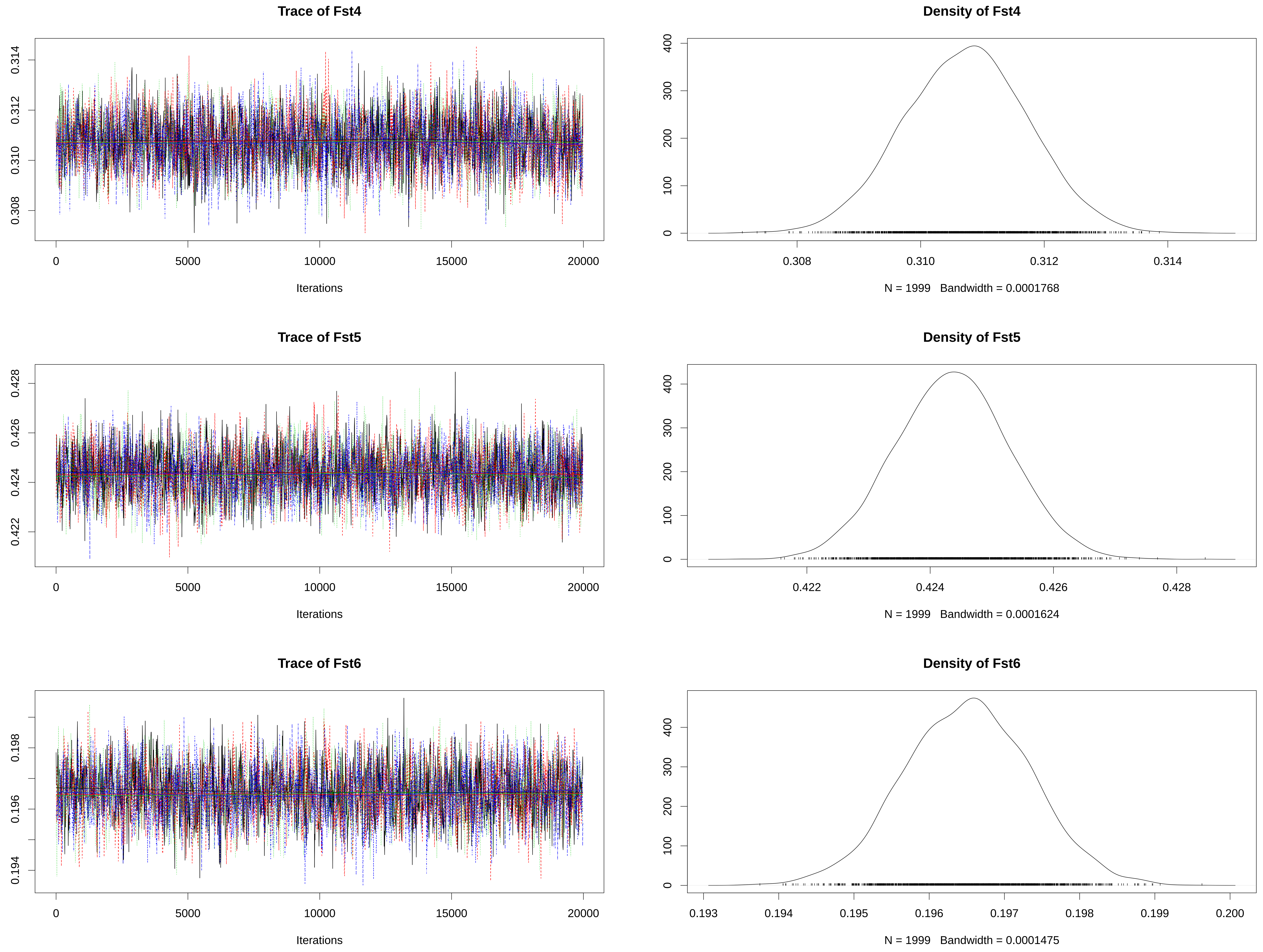

### figureS5A.pdf

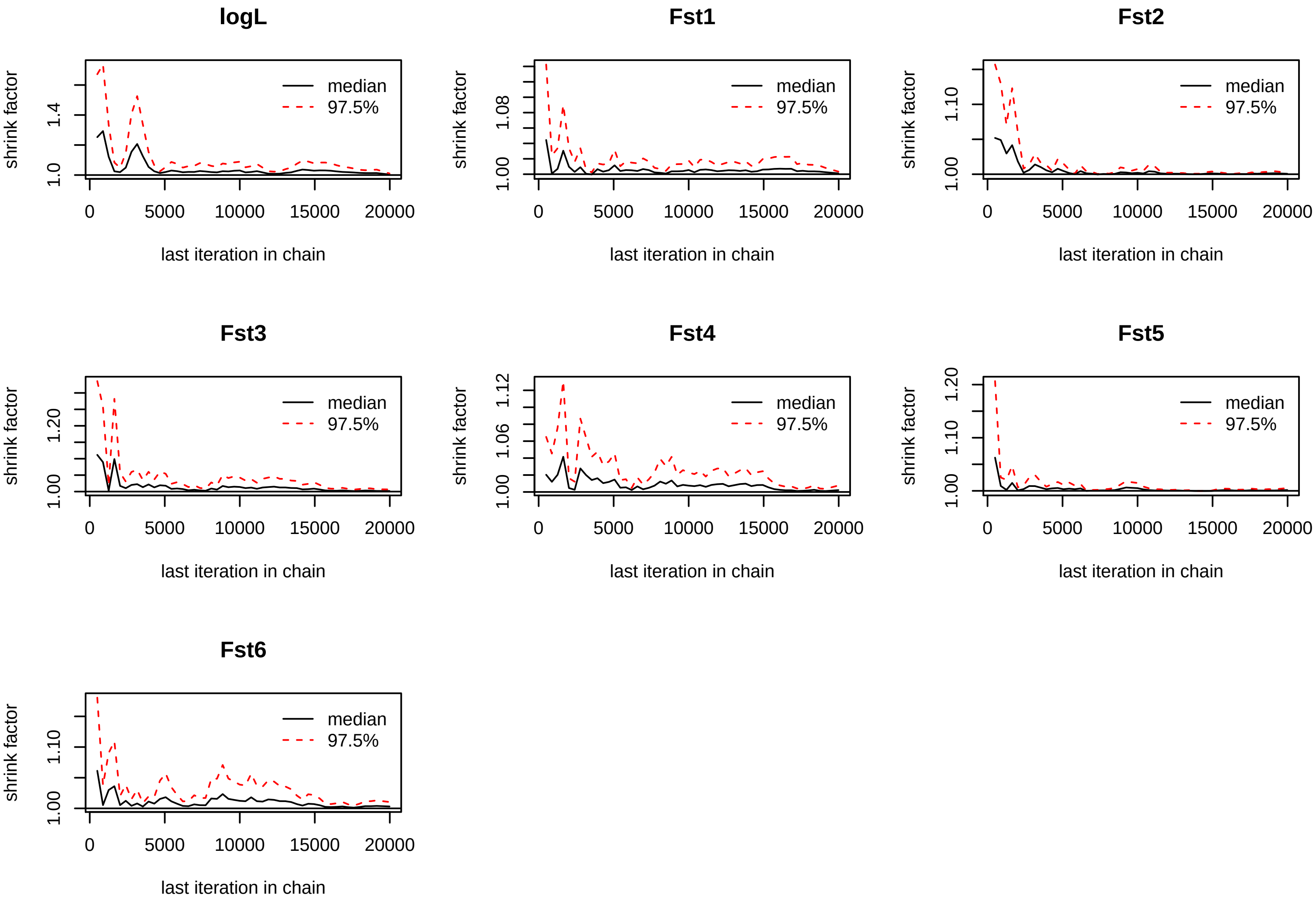

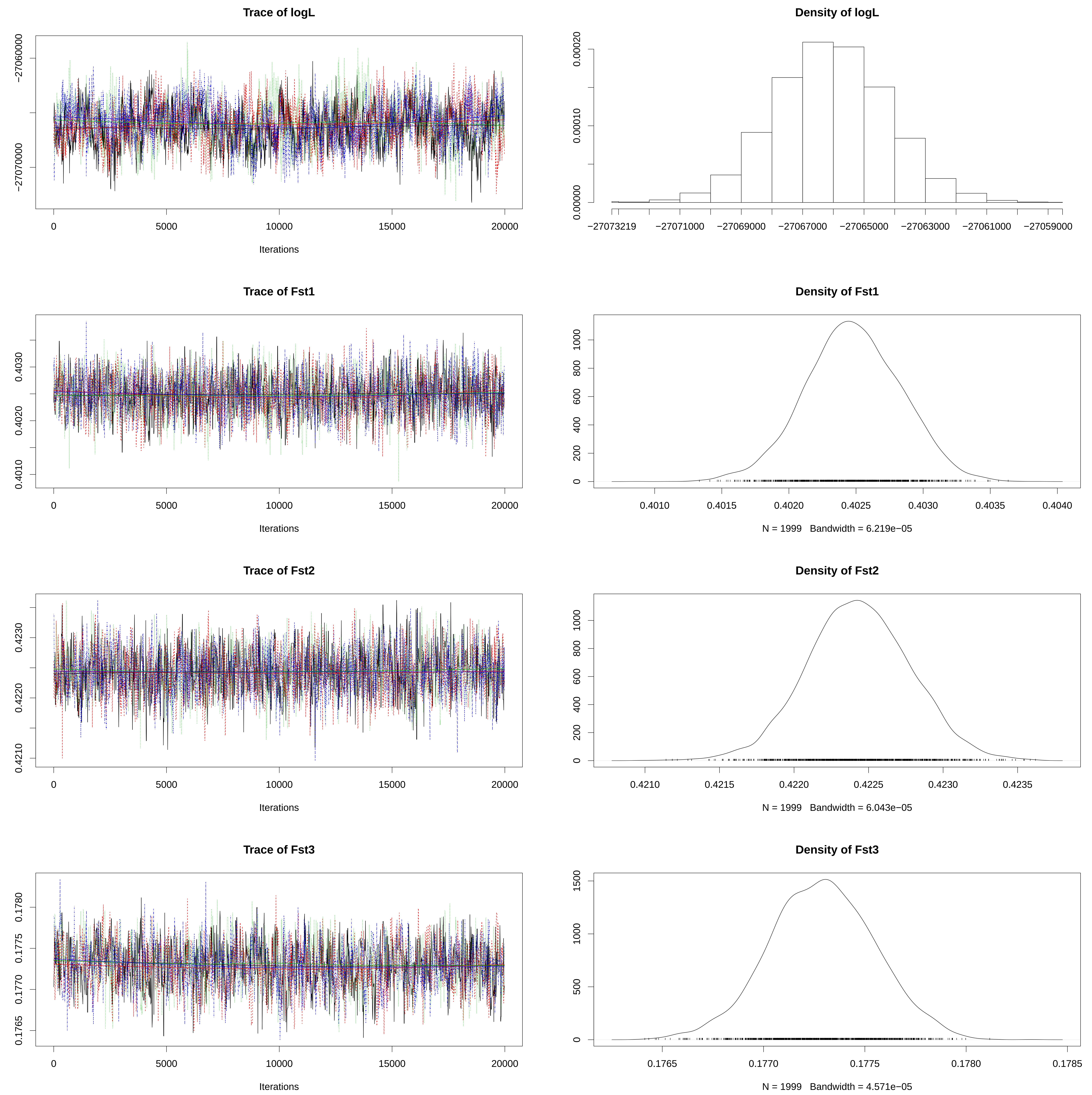

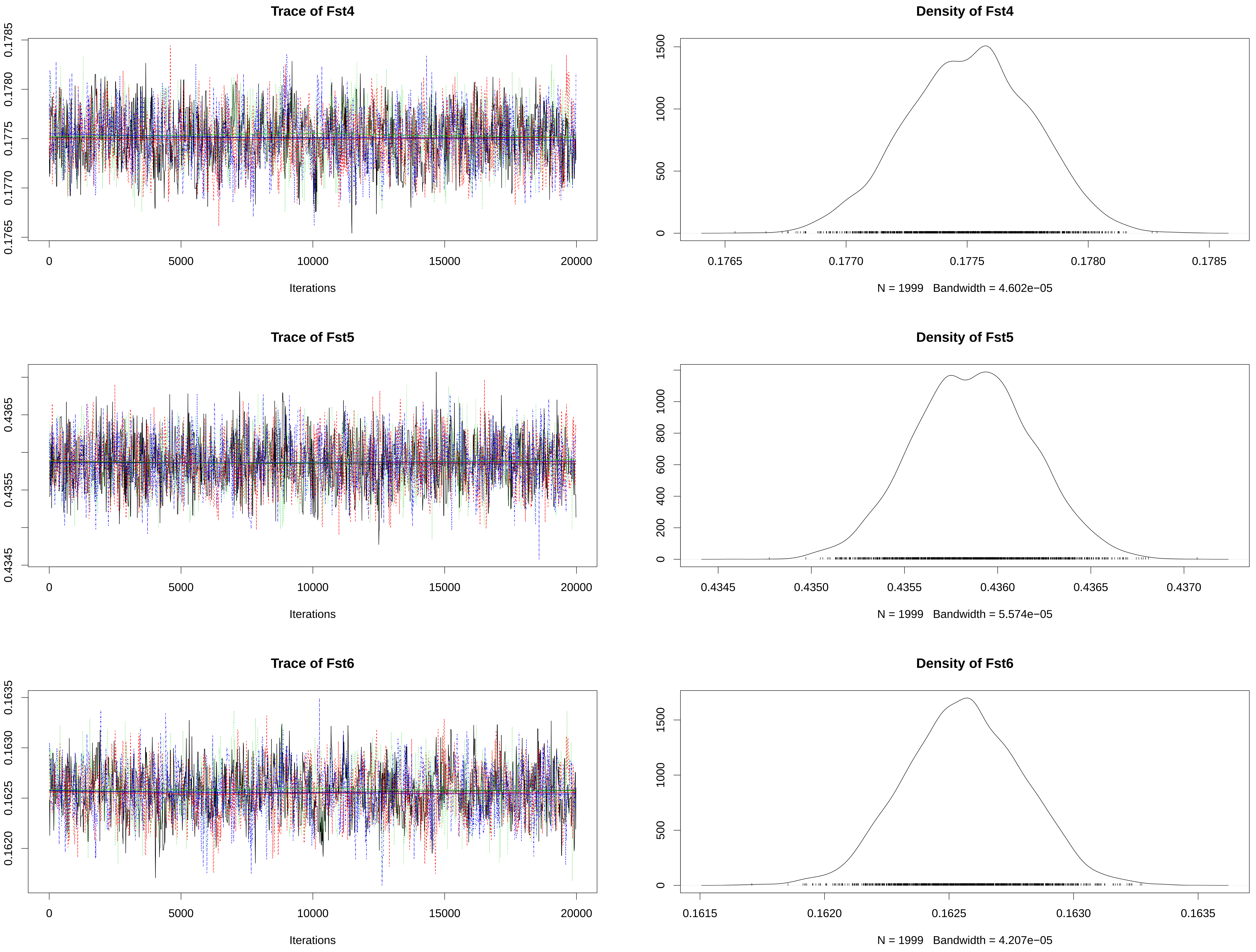

### figureS5B.pdf

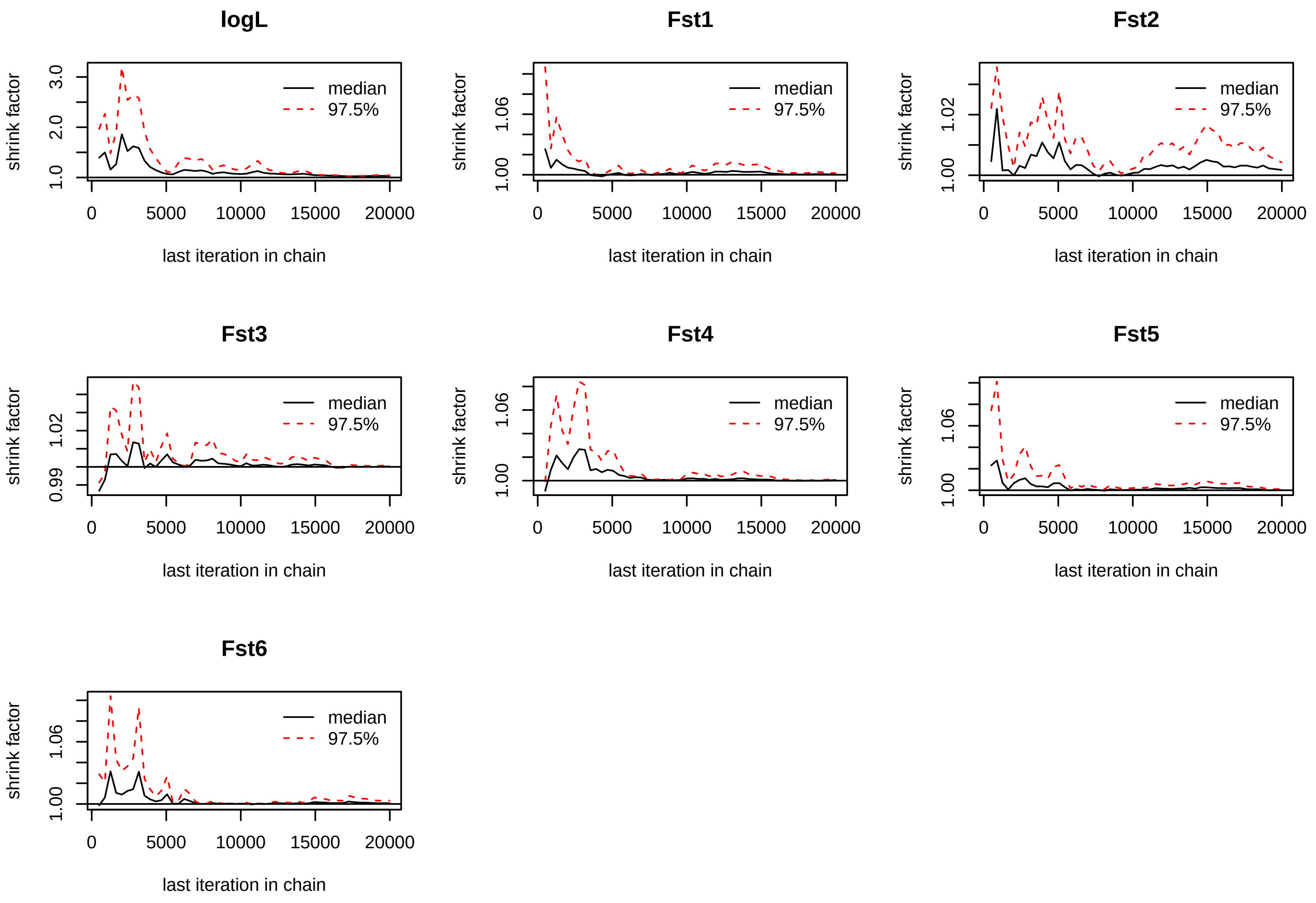

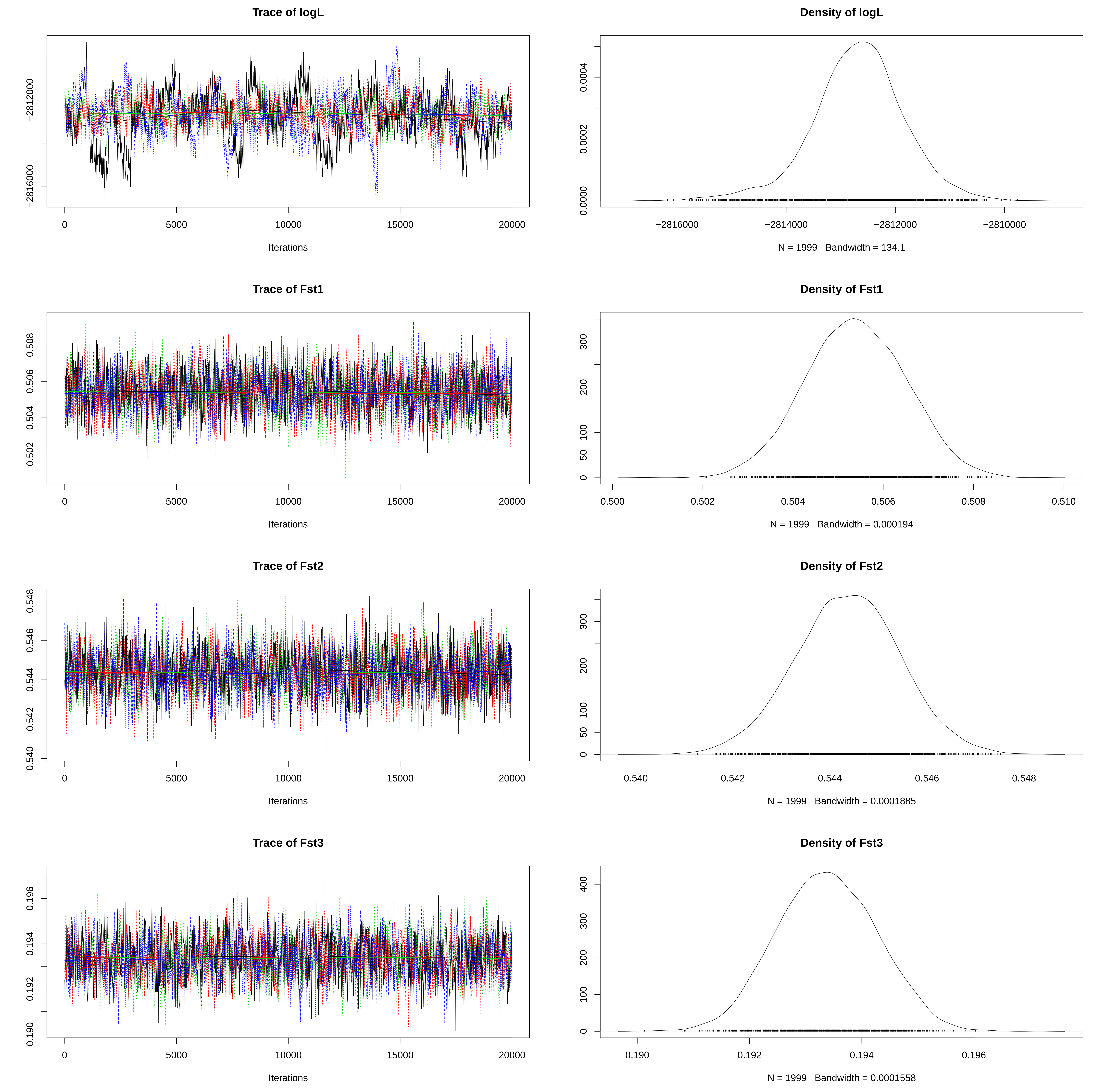

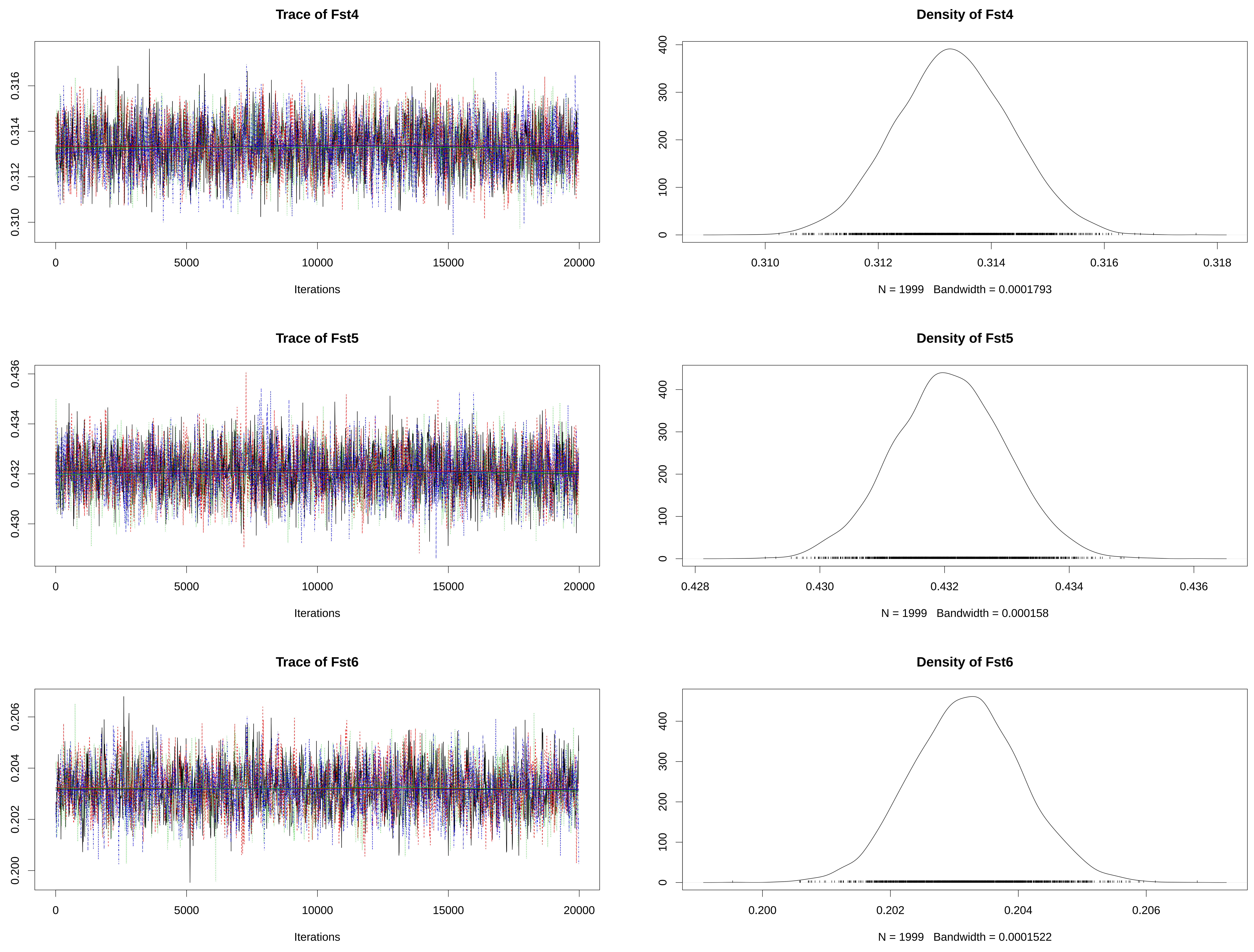

### figureS6A.pdf

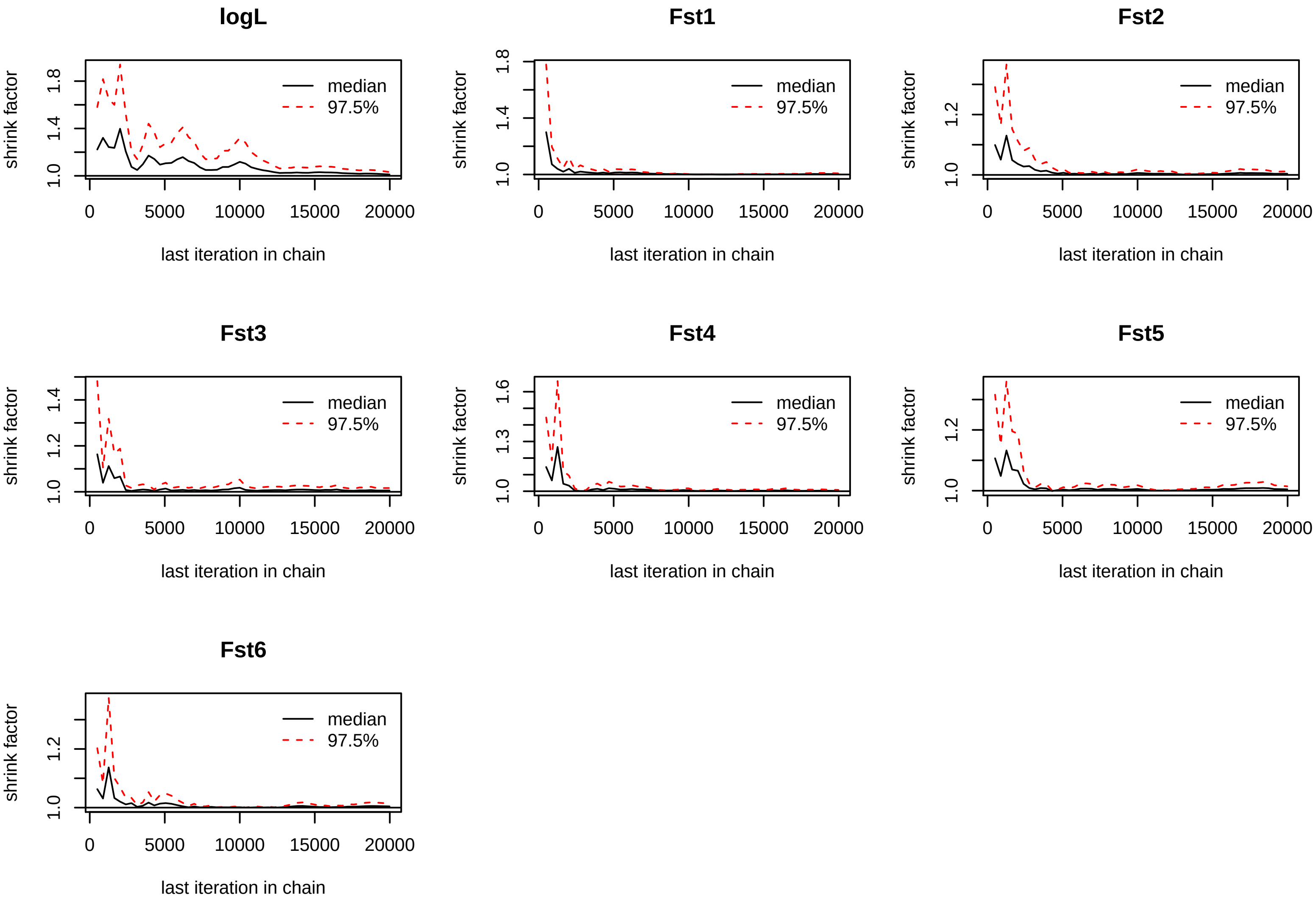

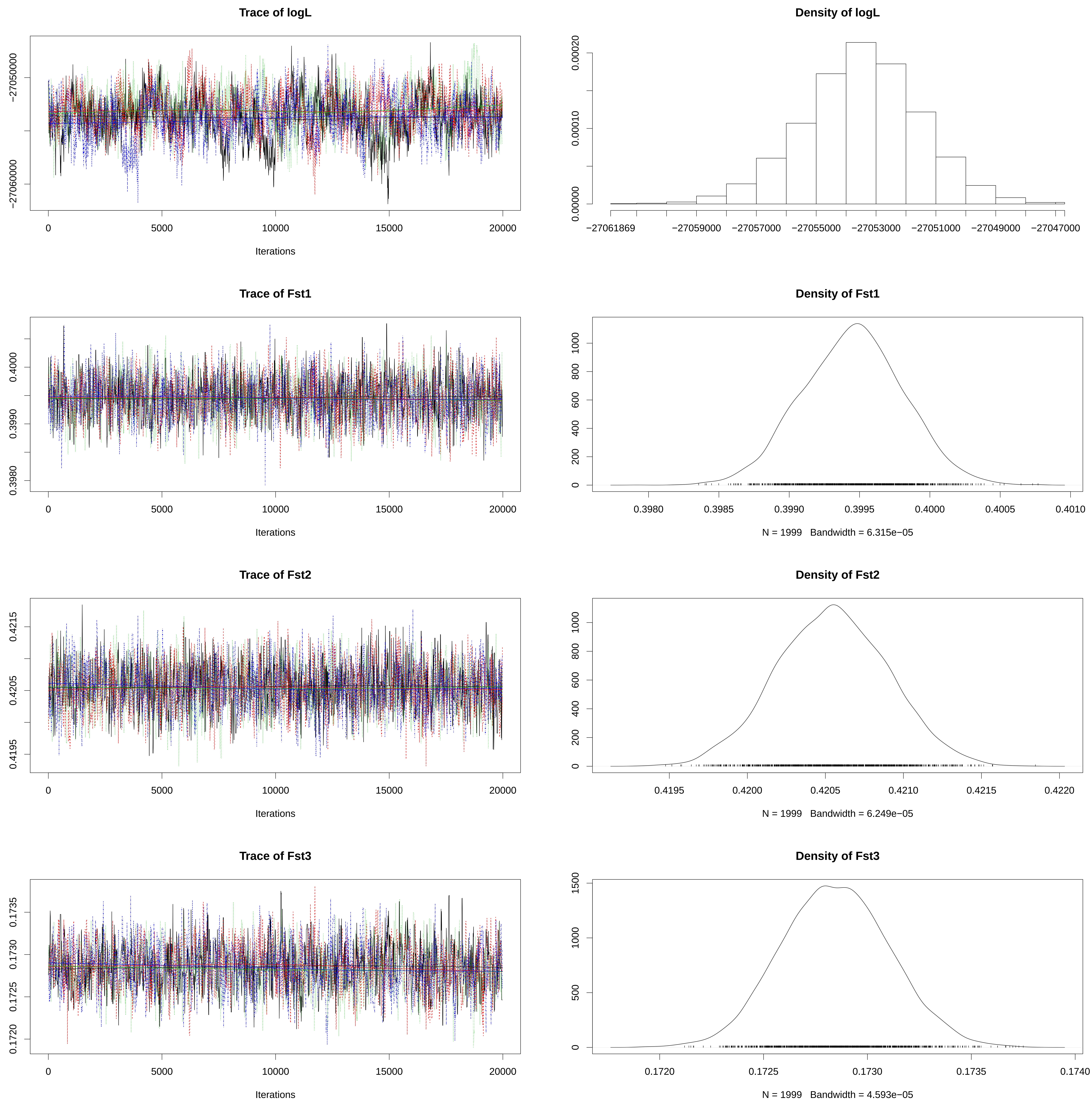

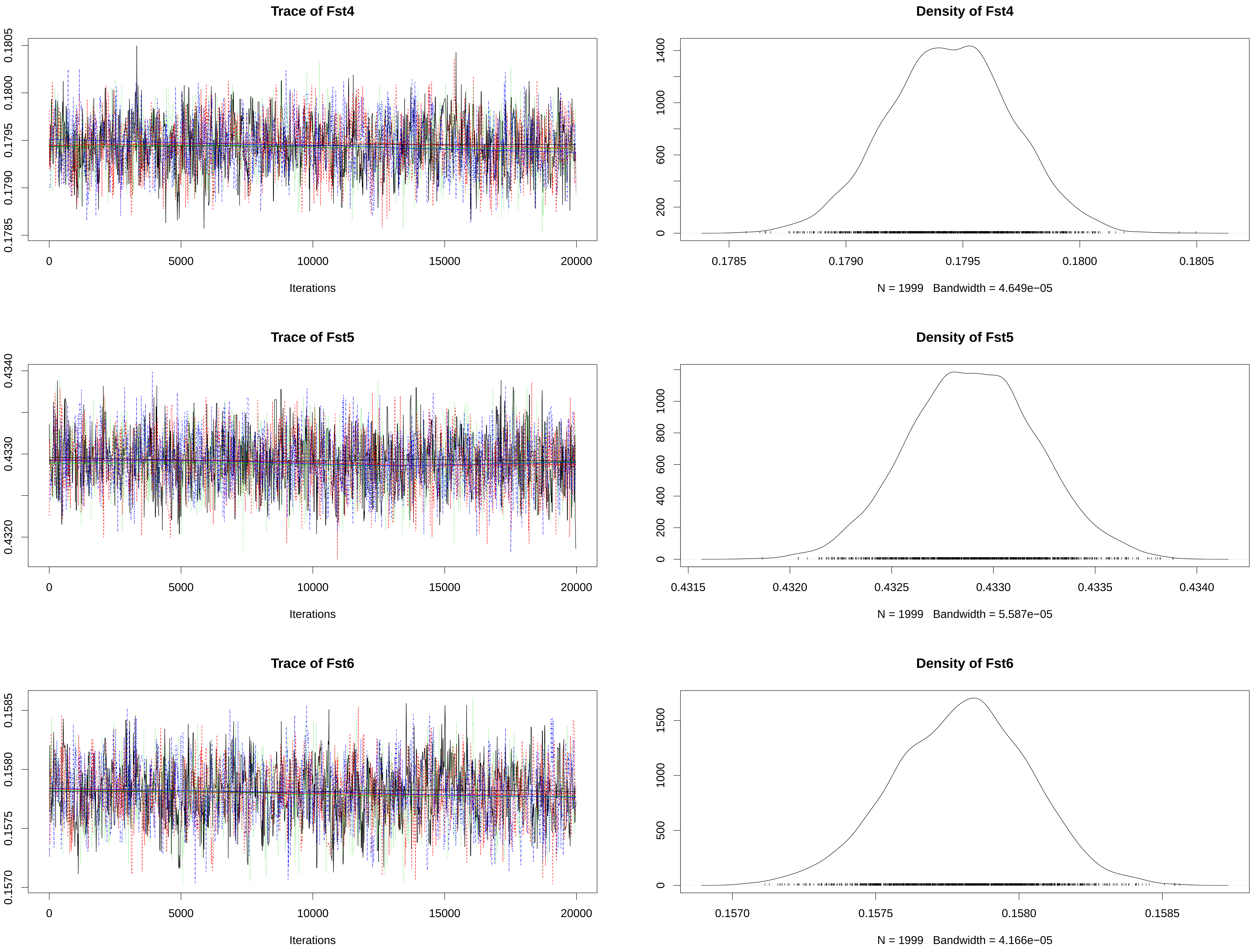

### figureS6B.pdf

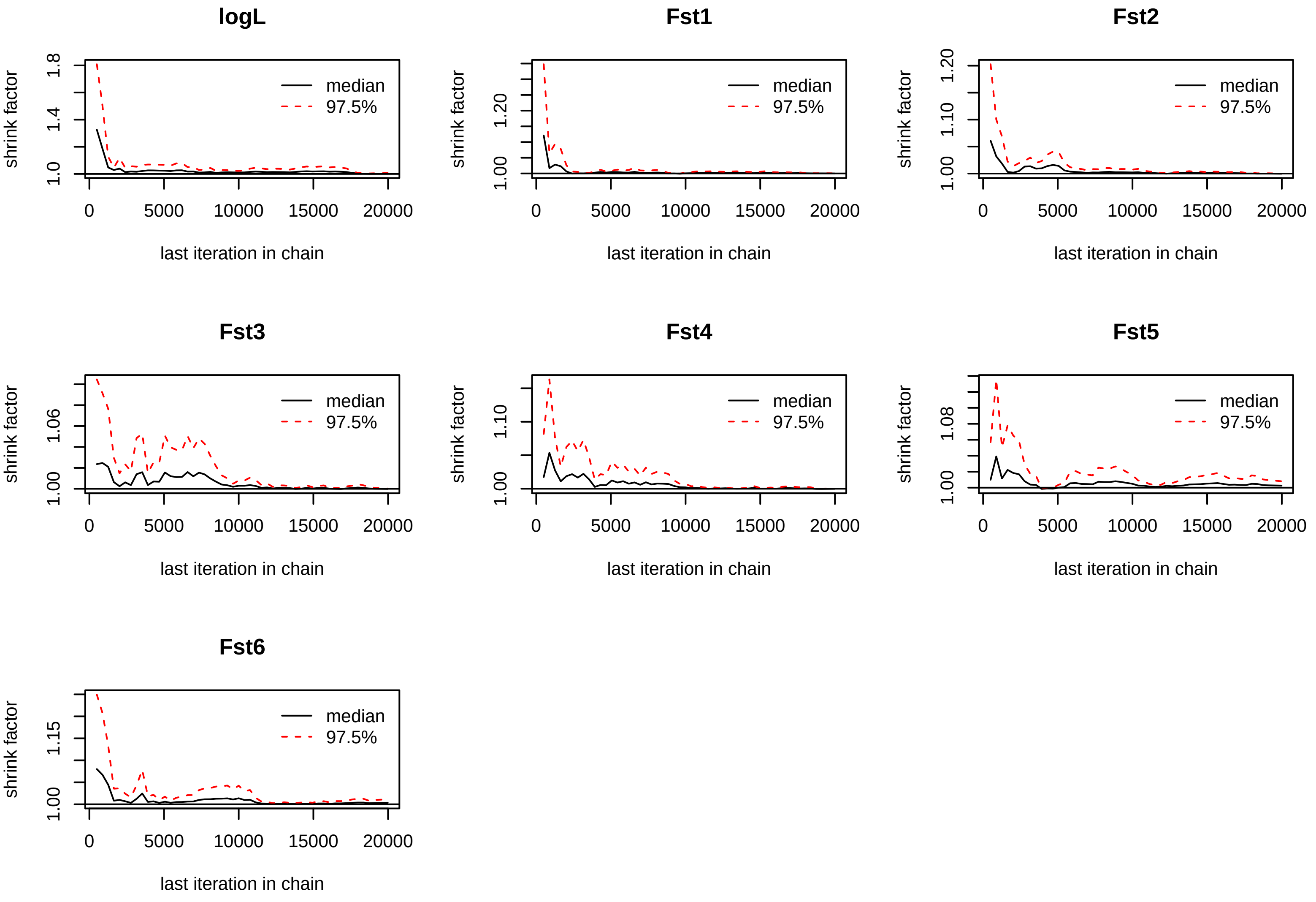

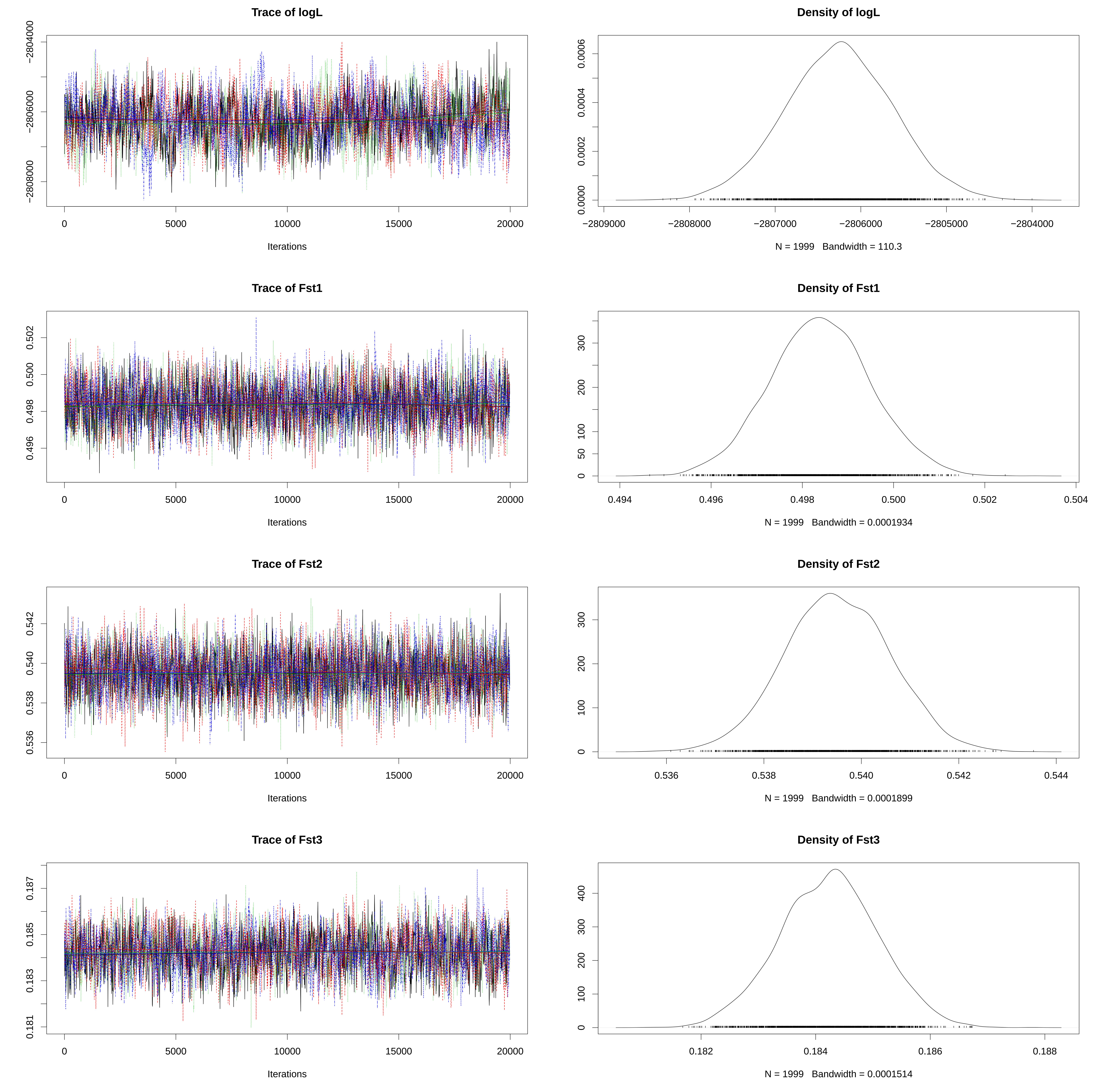

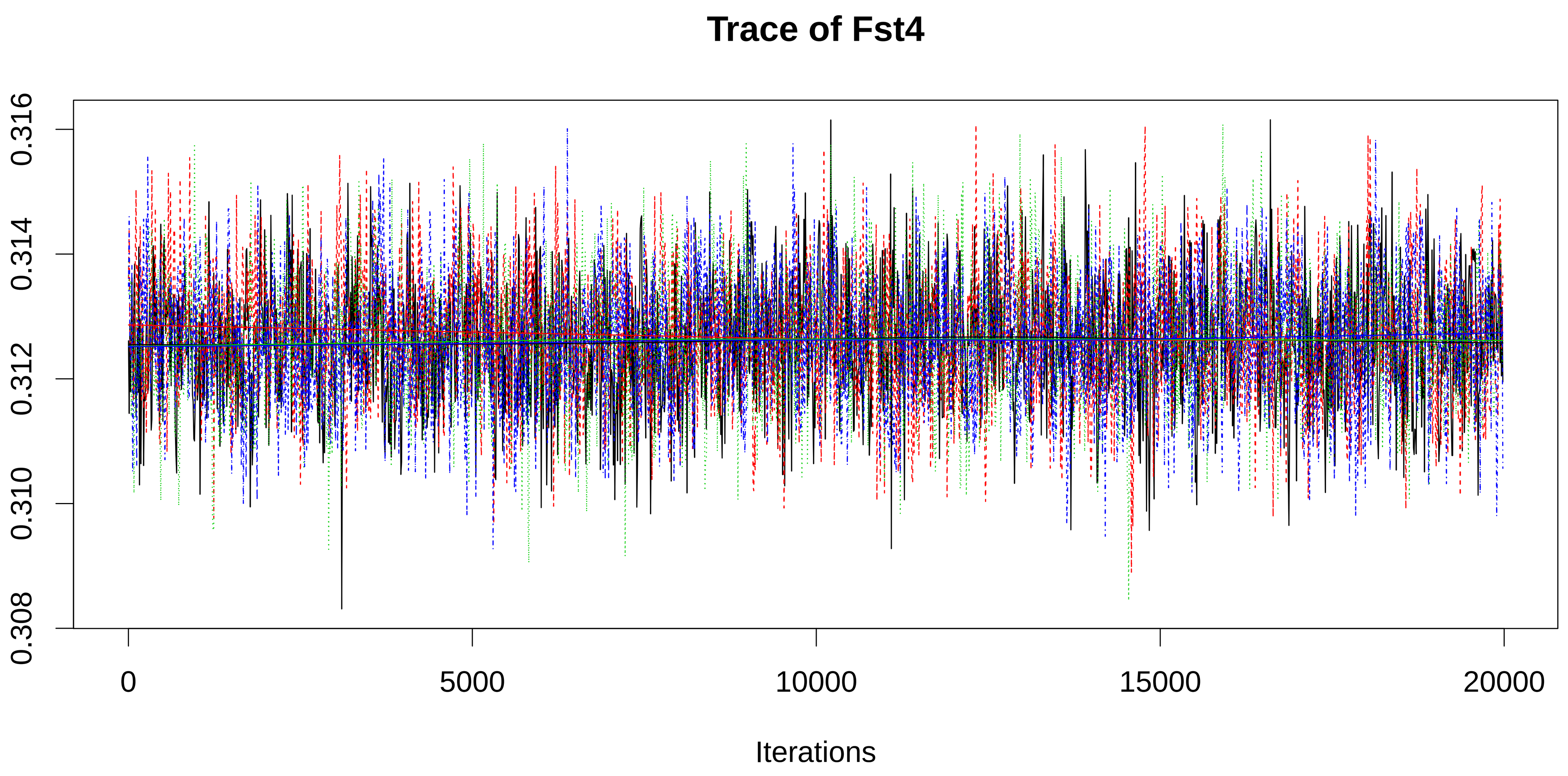

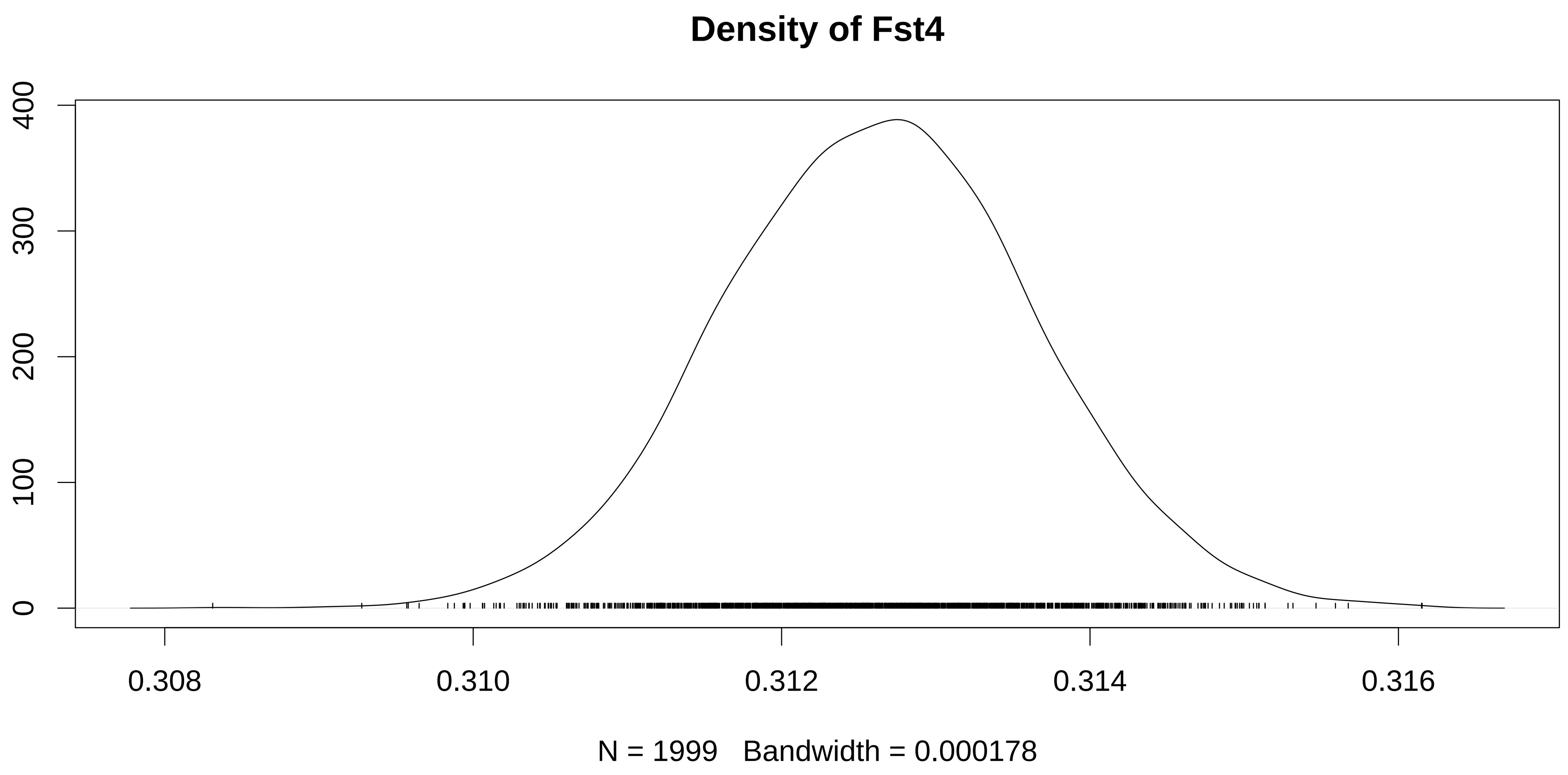

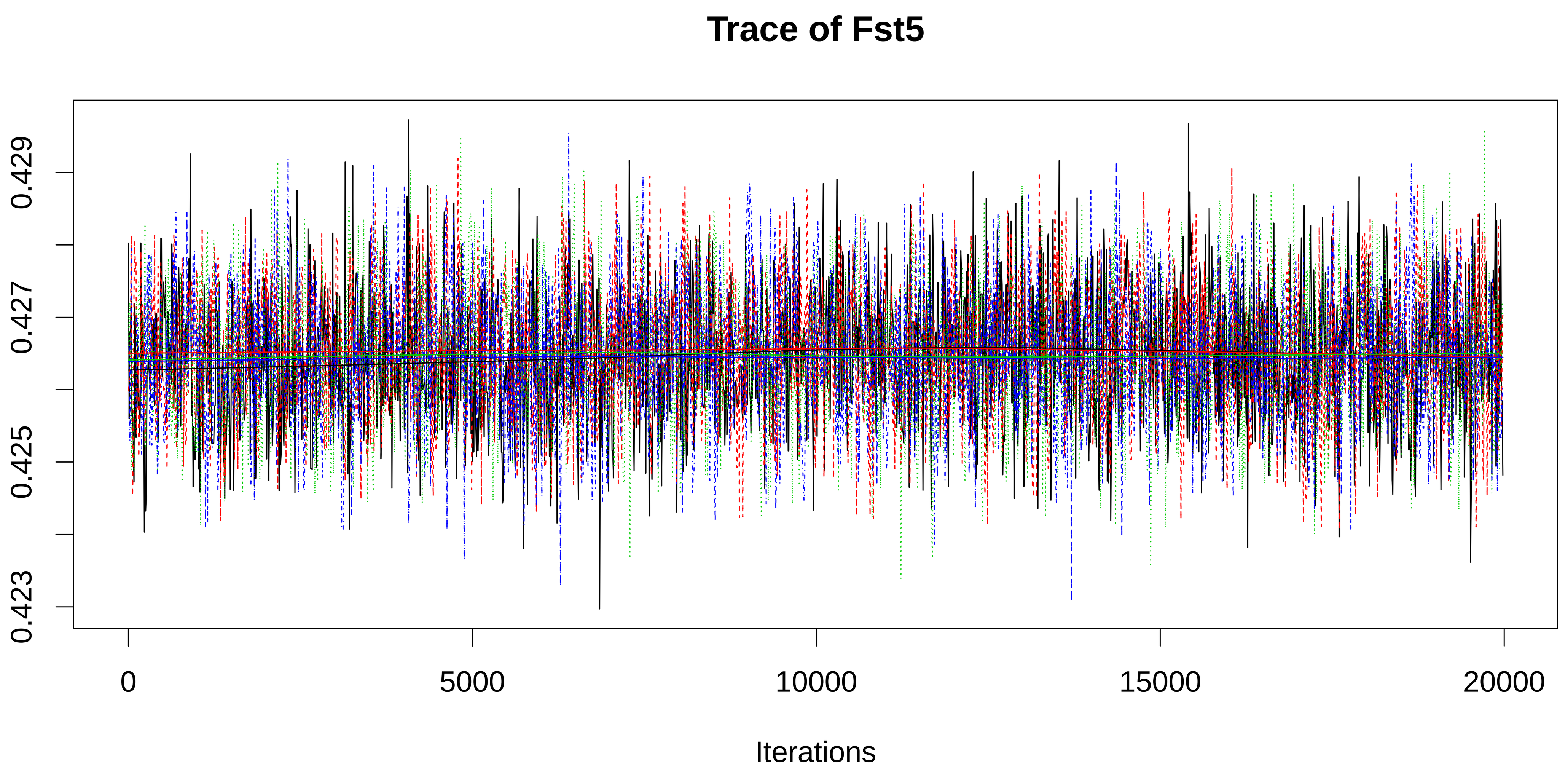

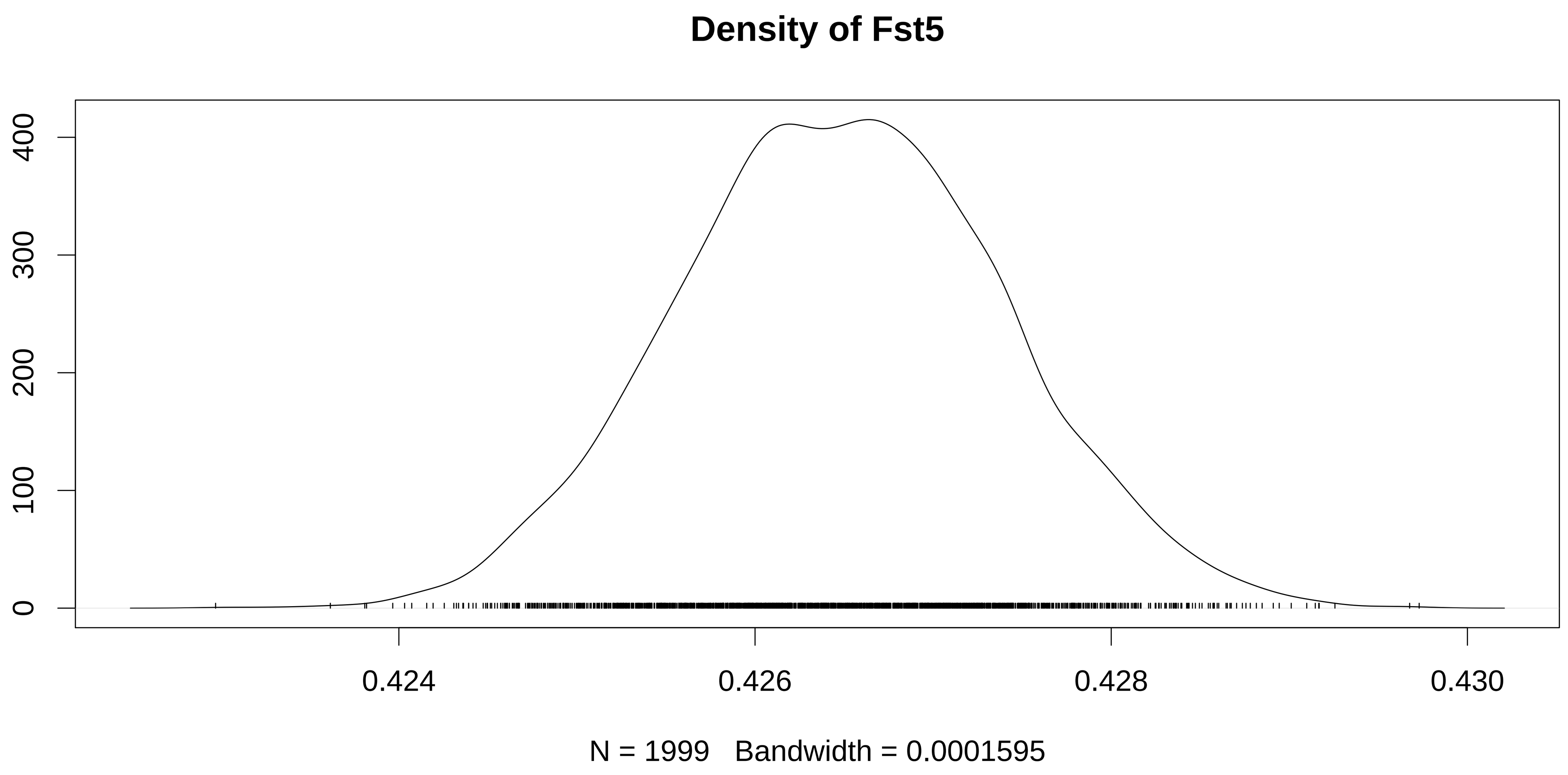

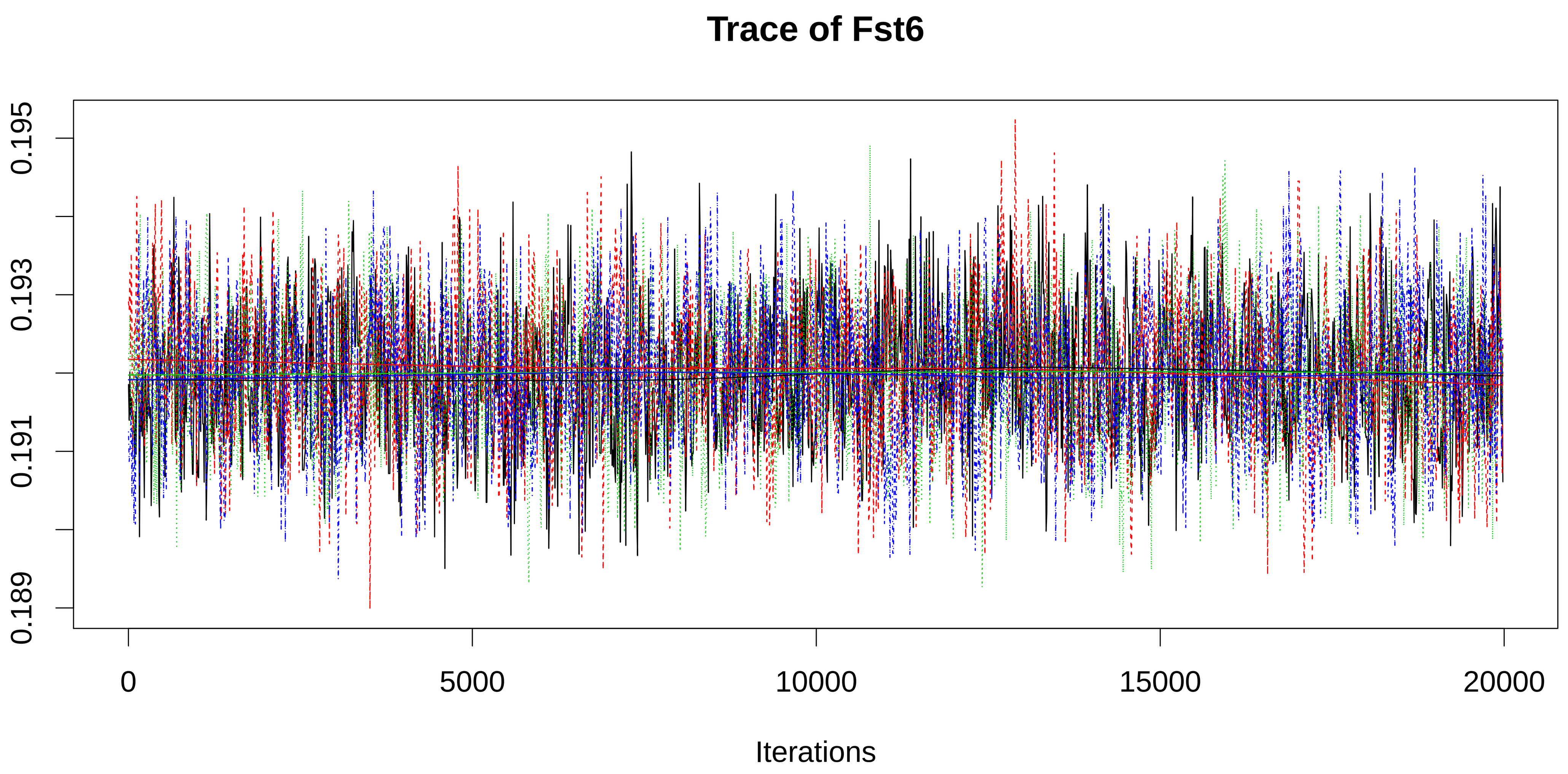

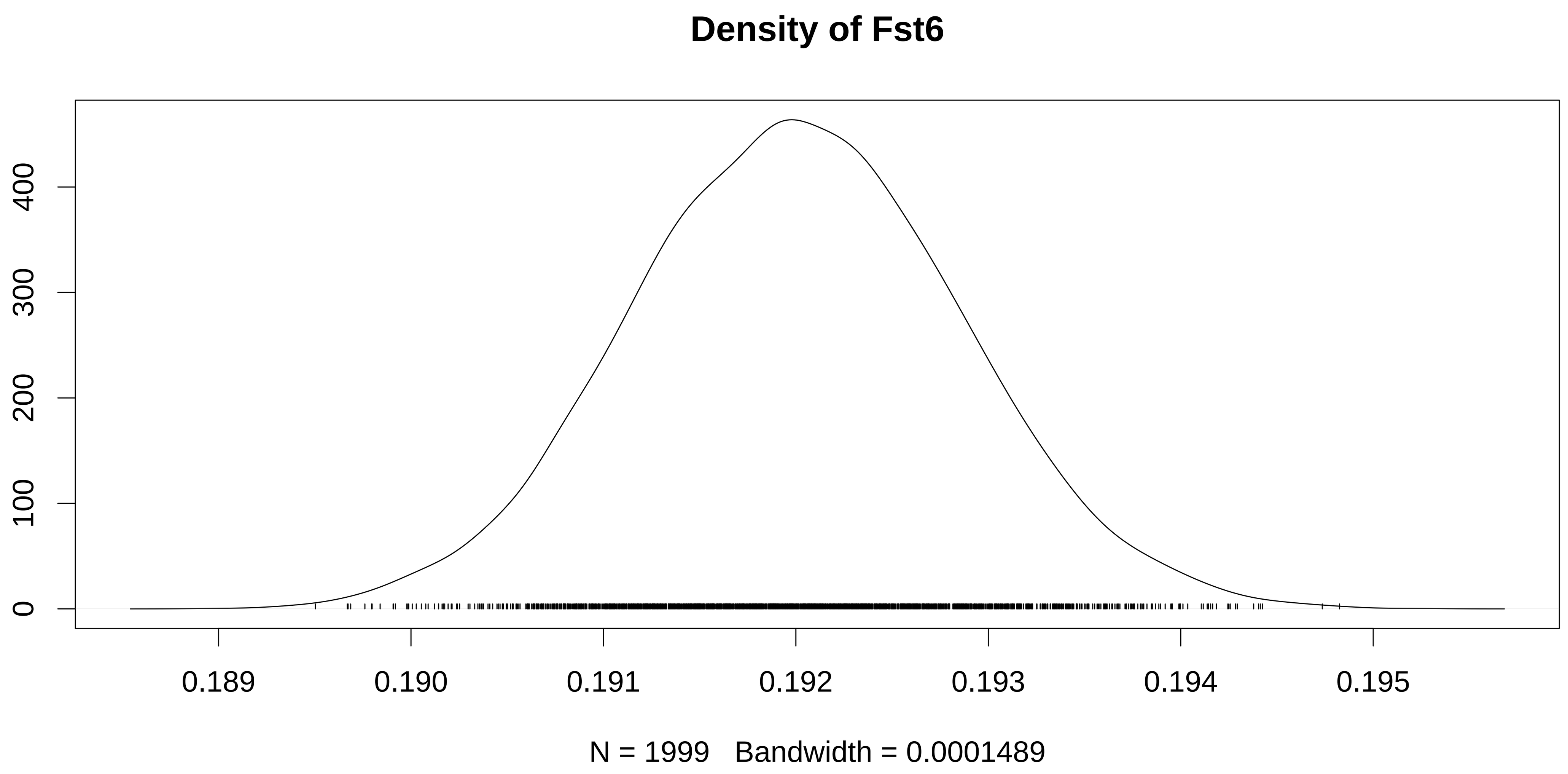

### figureS7A.pdf

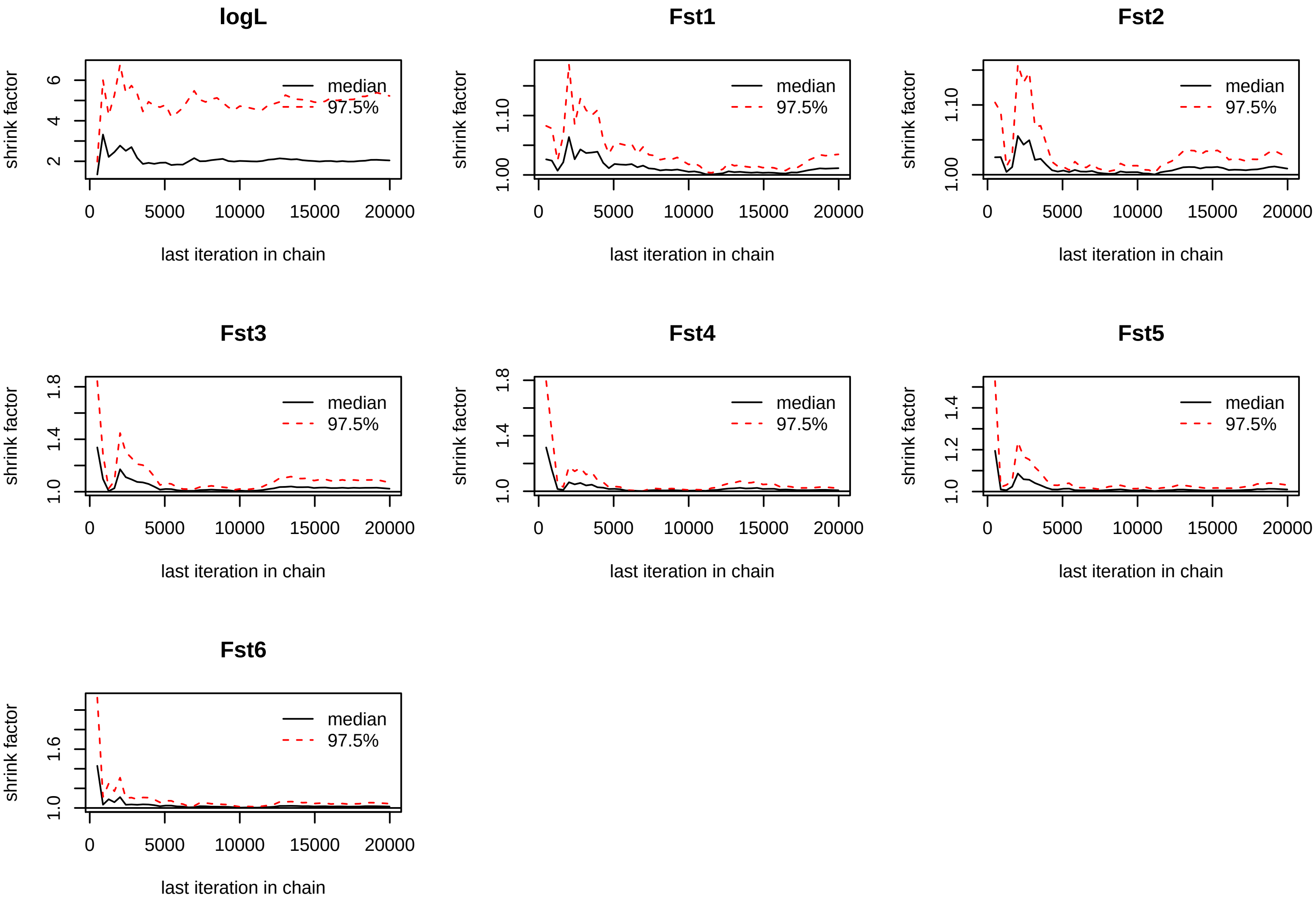

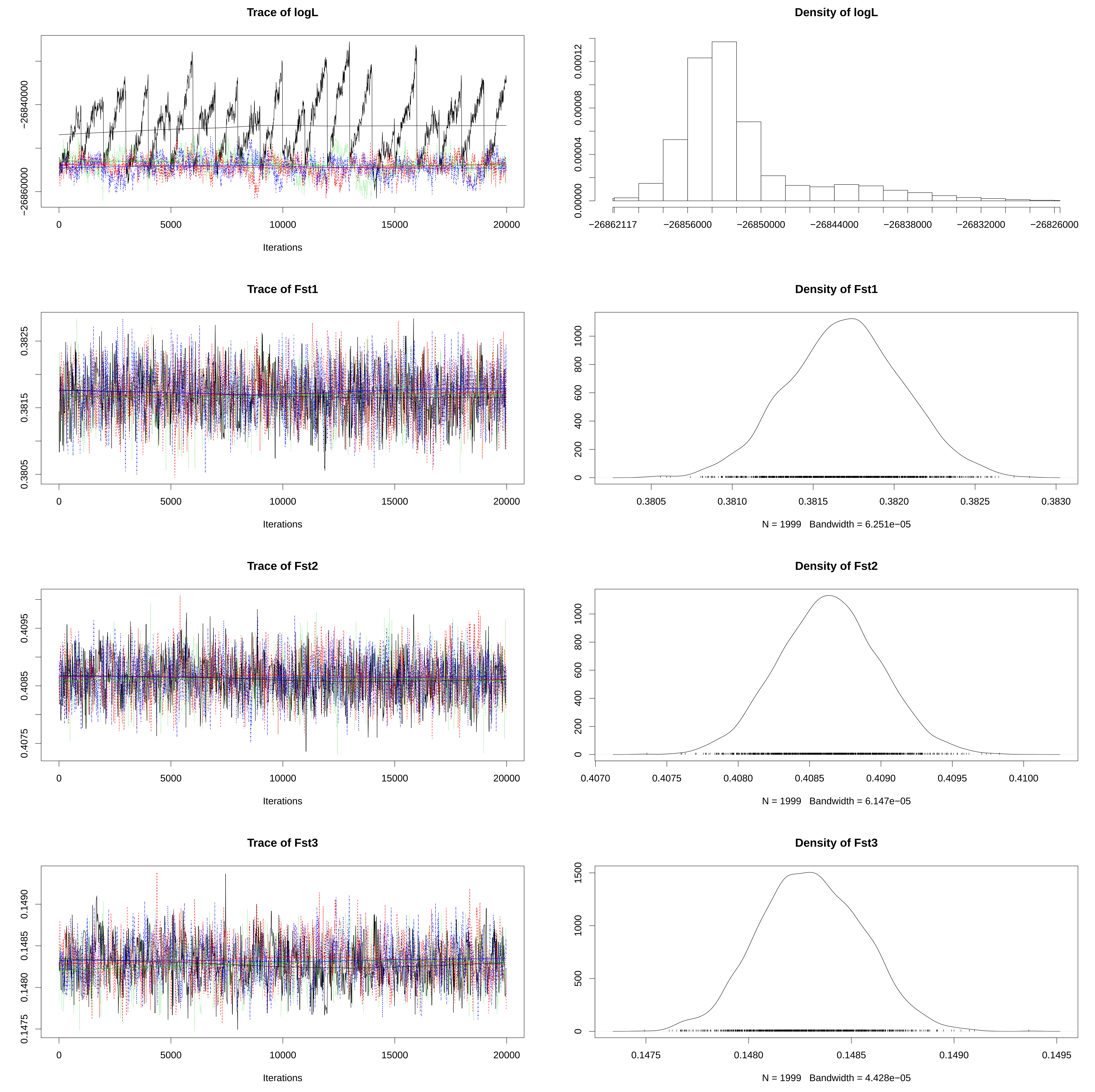

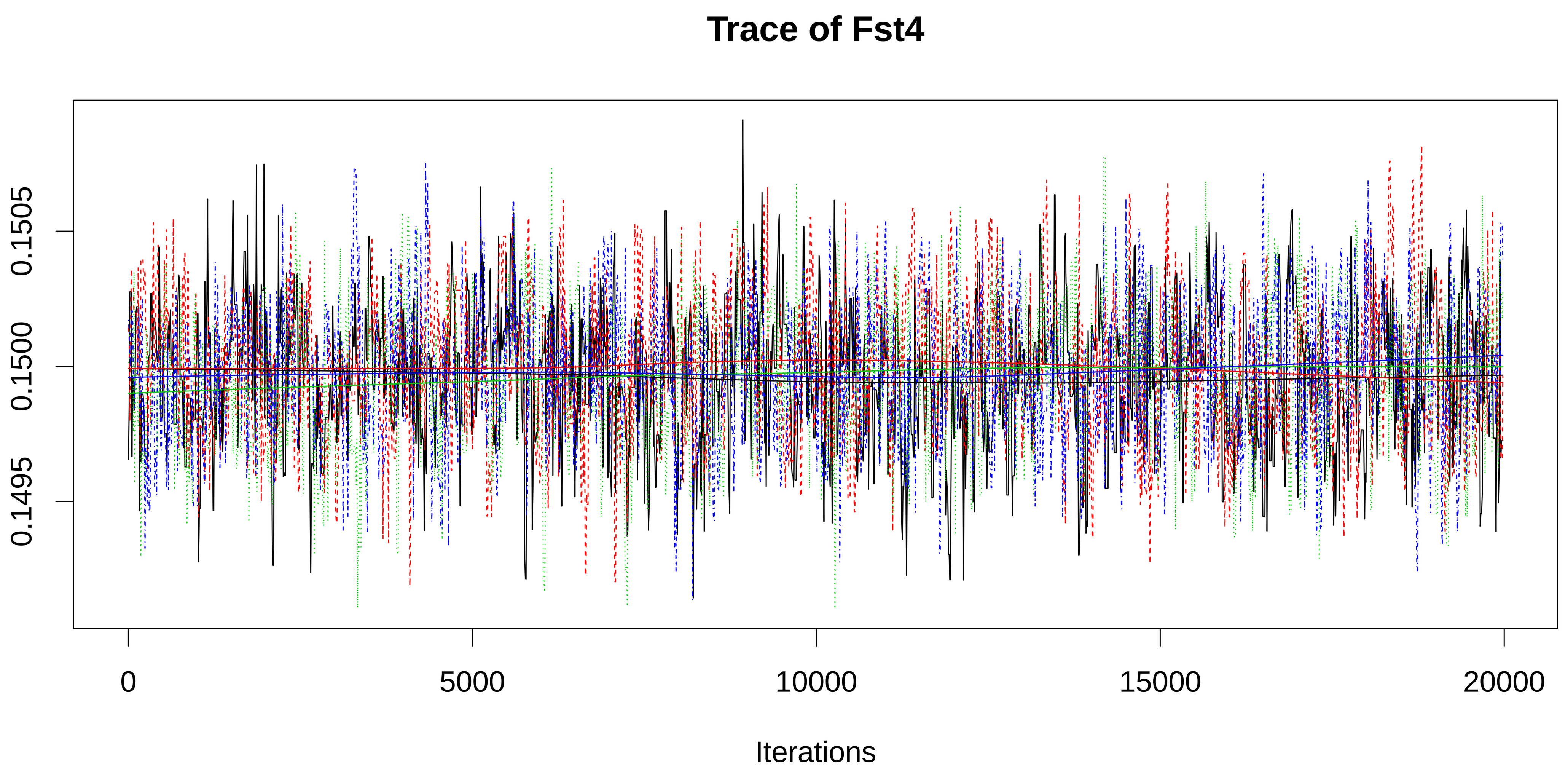

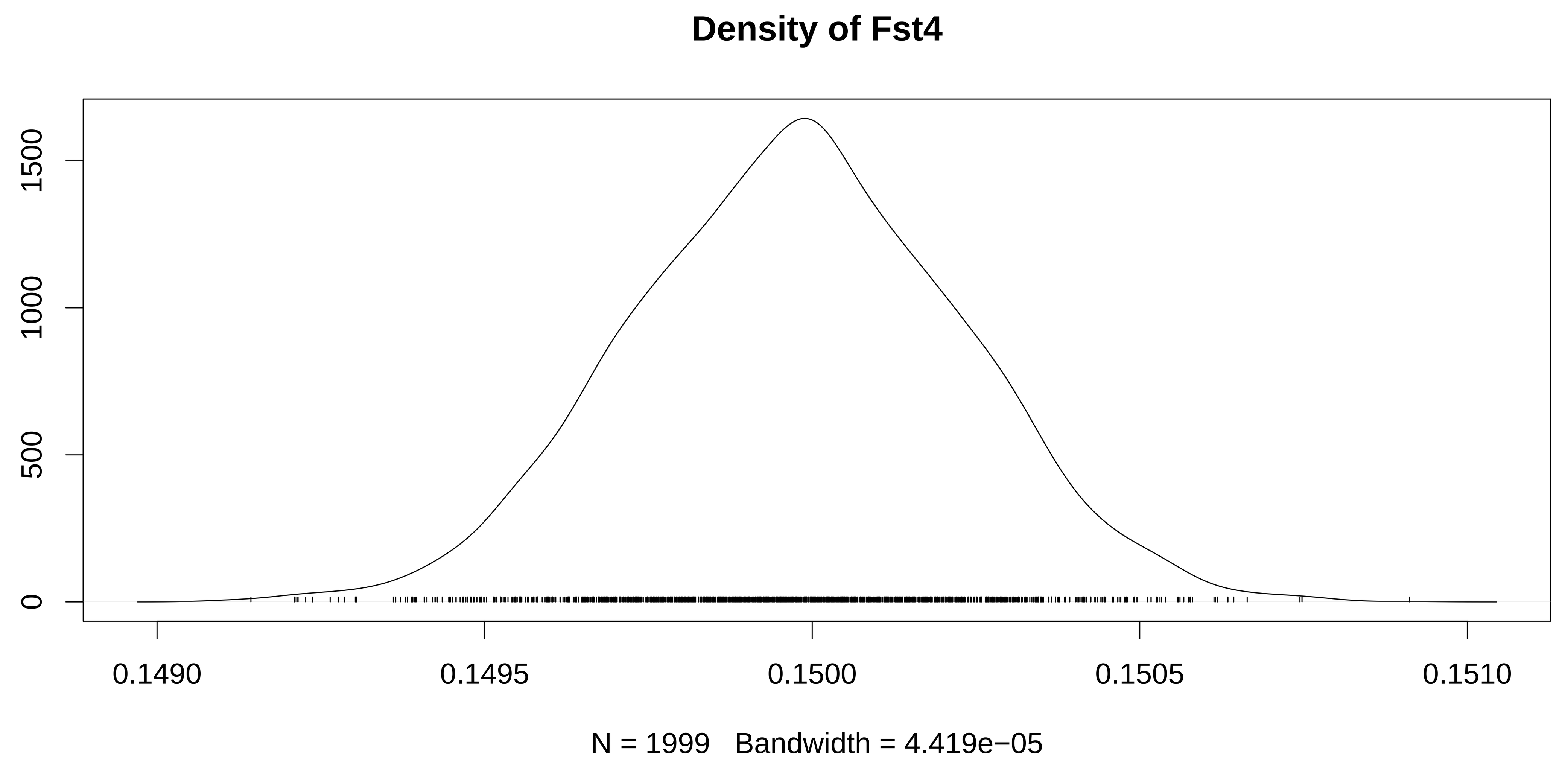

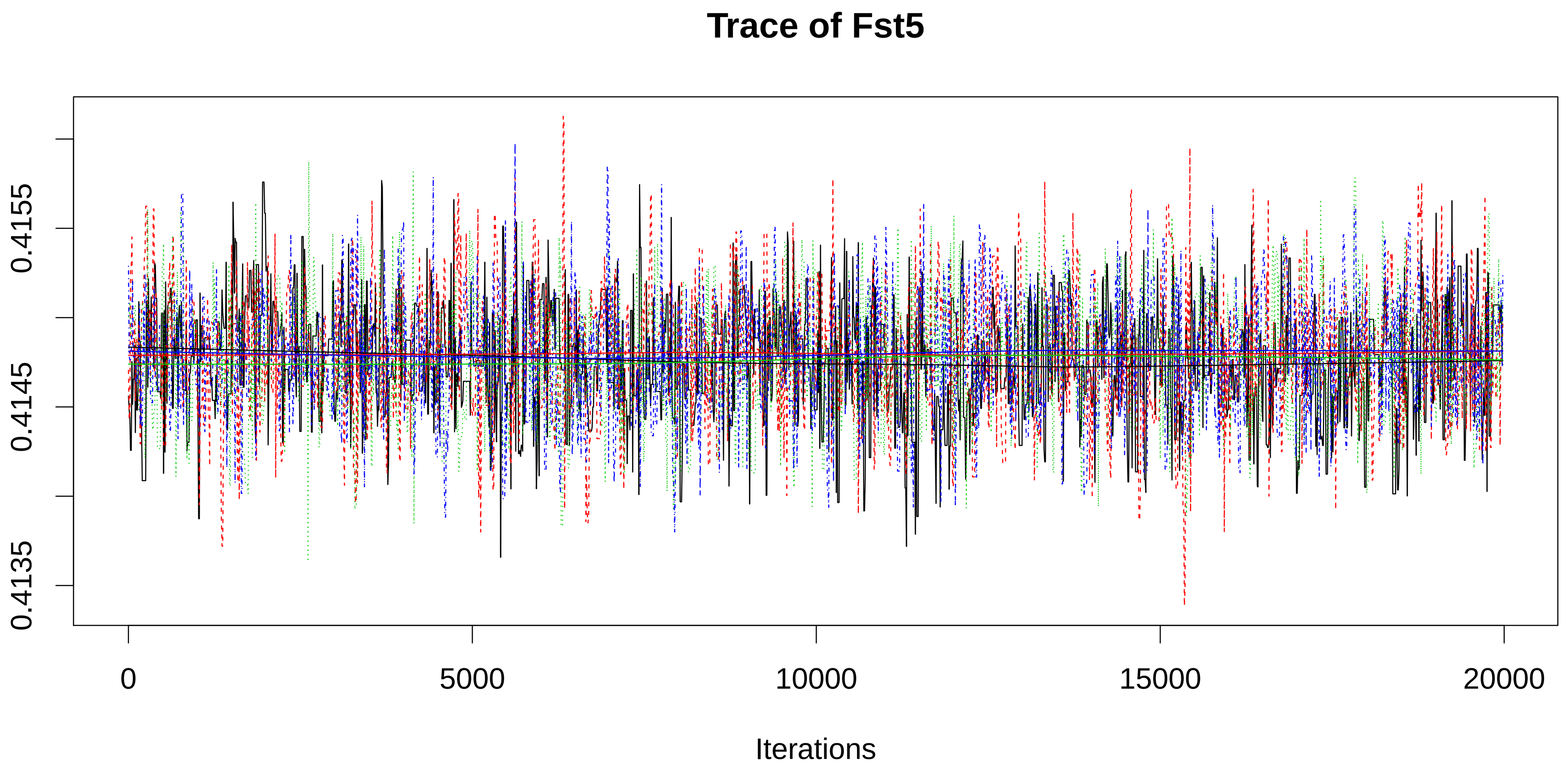

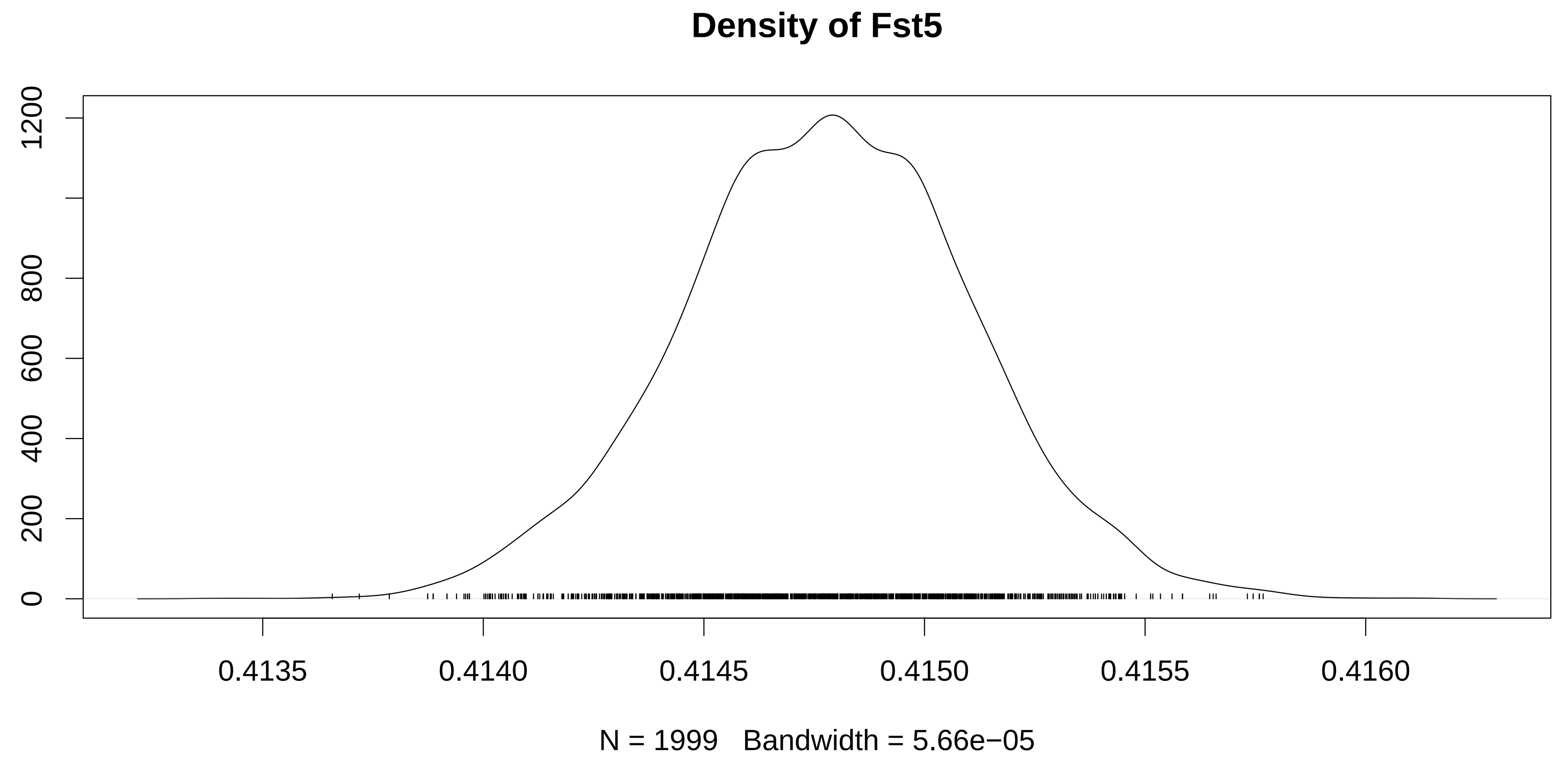

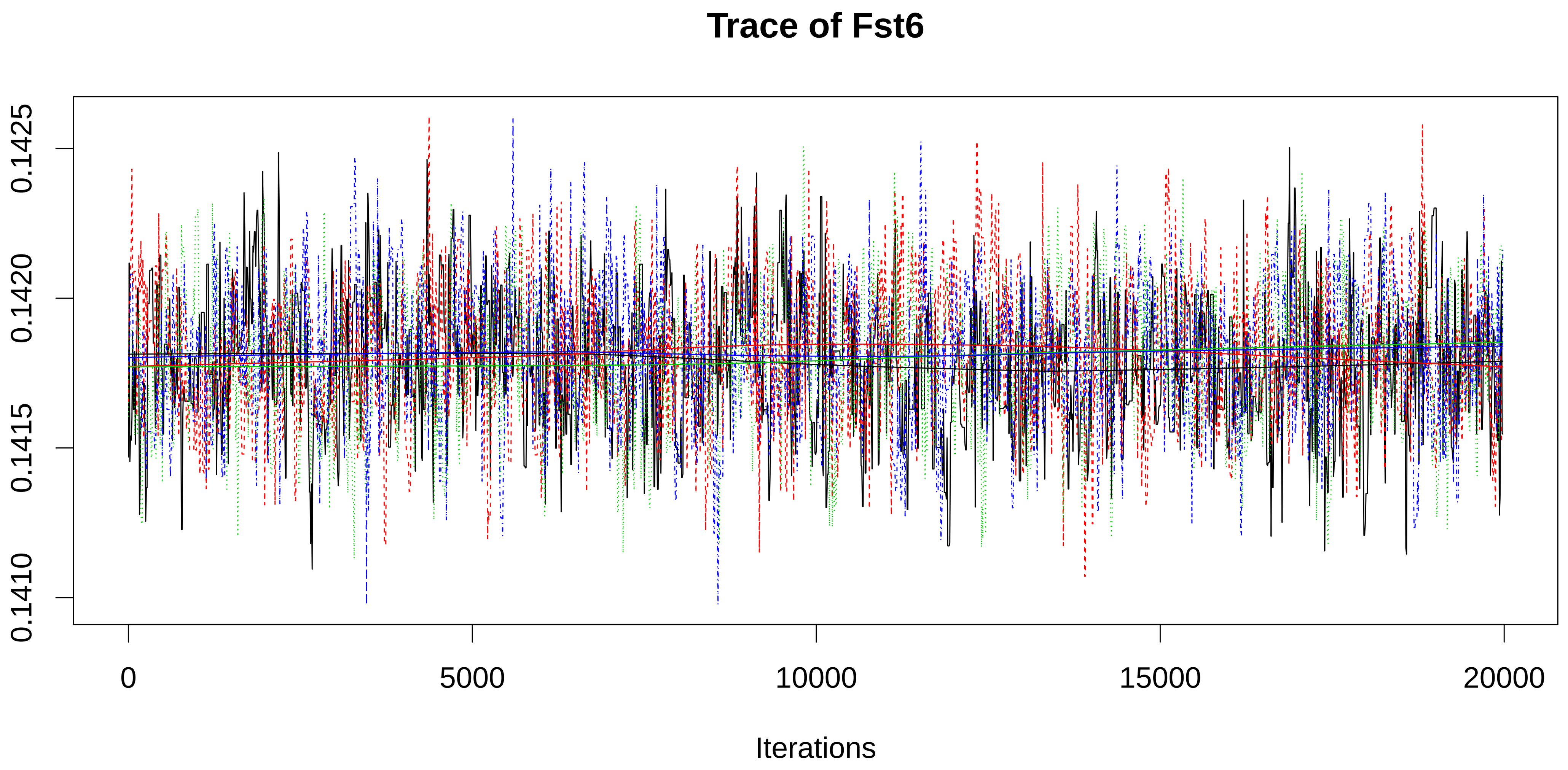

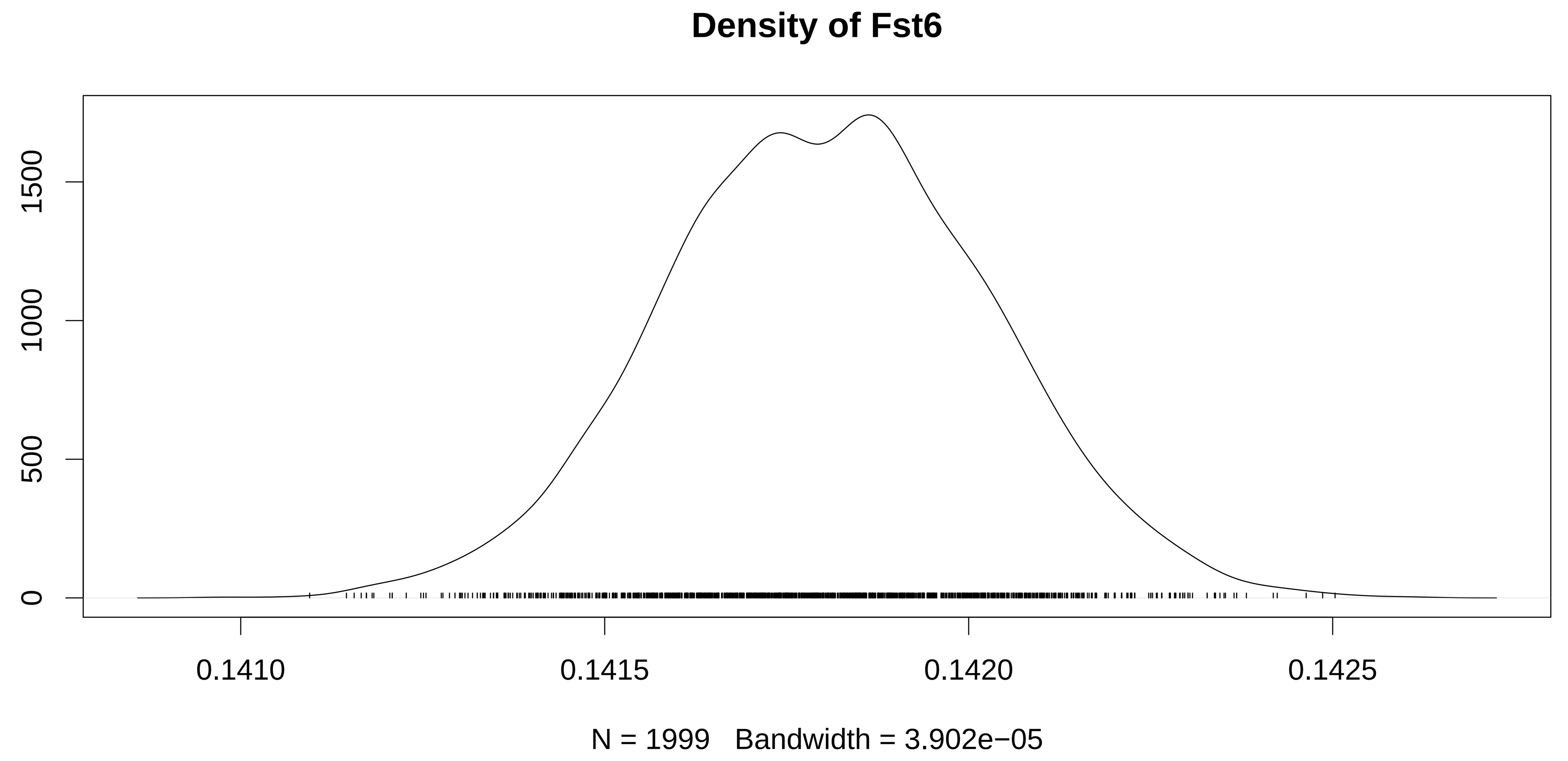

### figureS7B.pdf

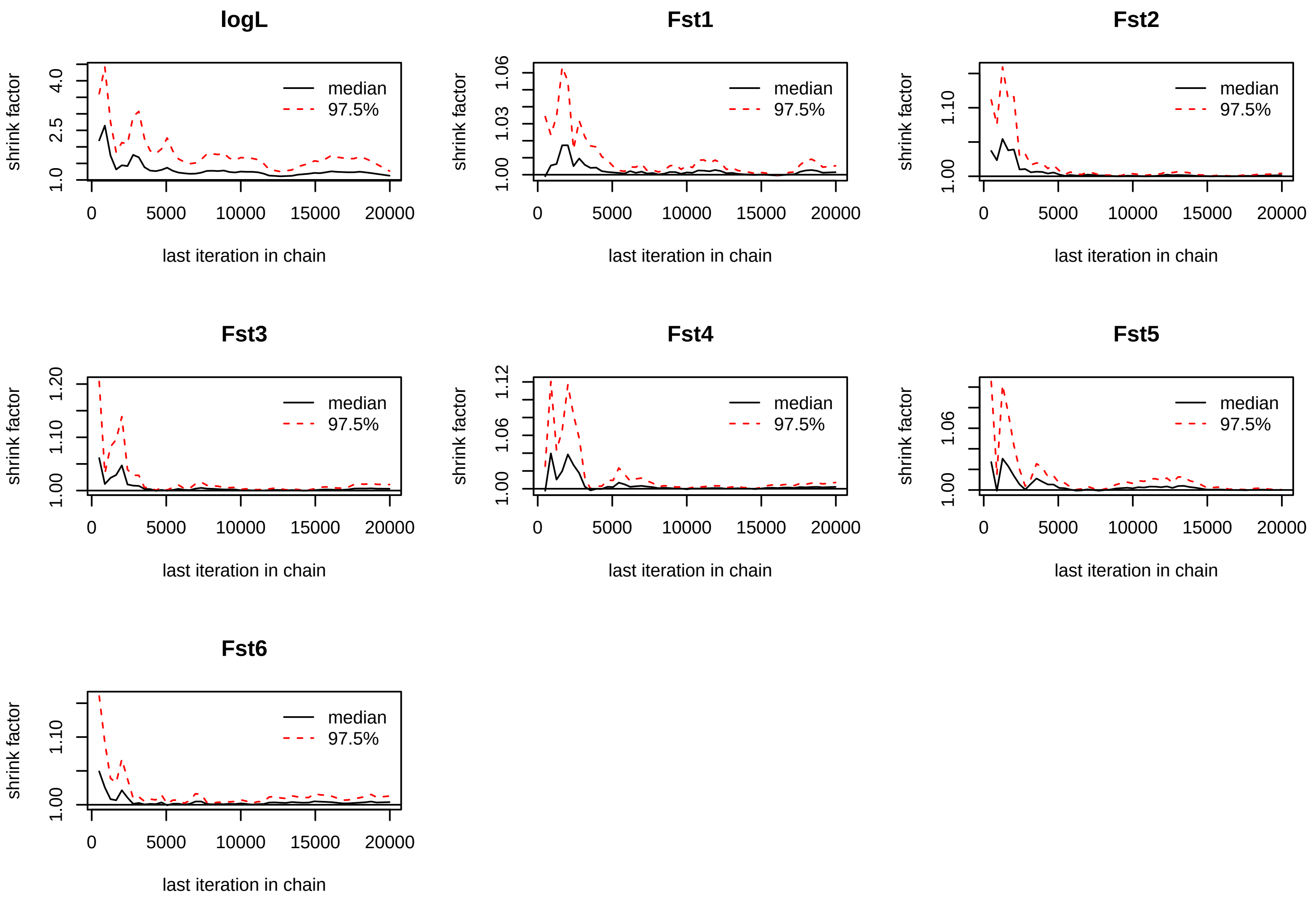

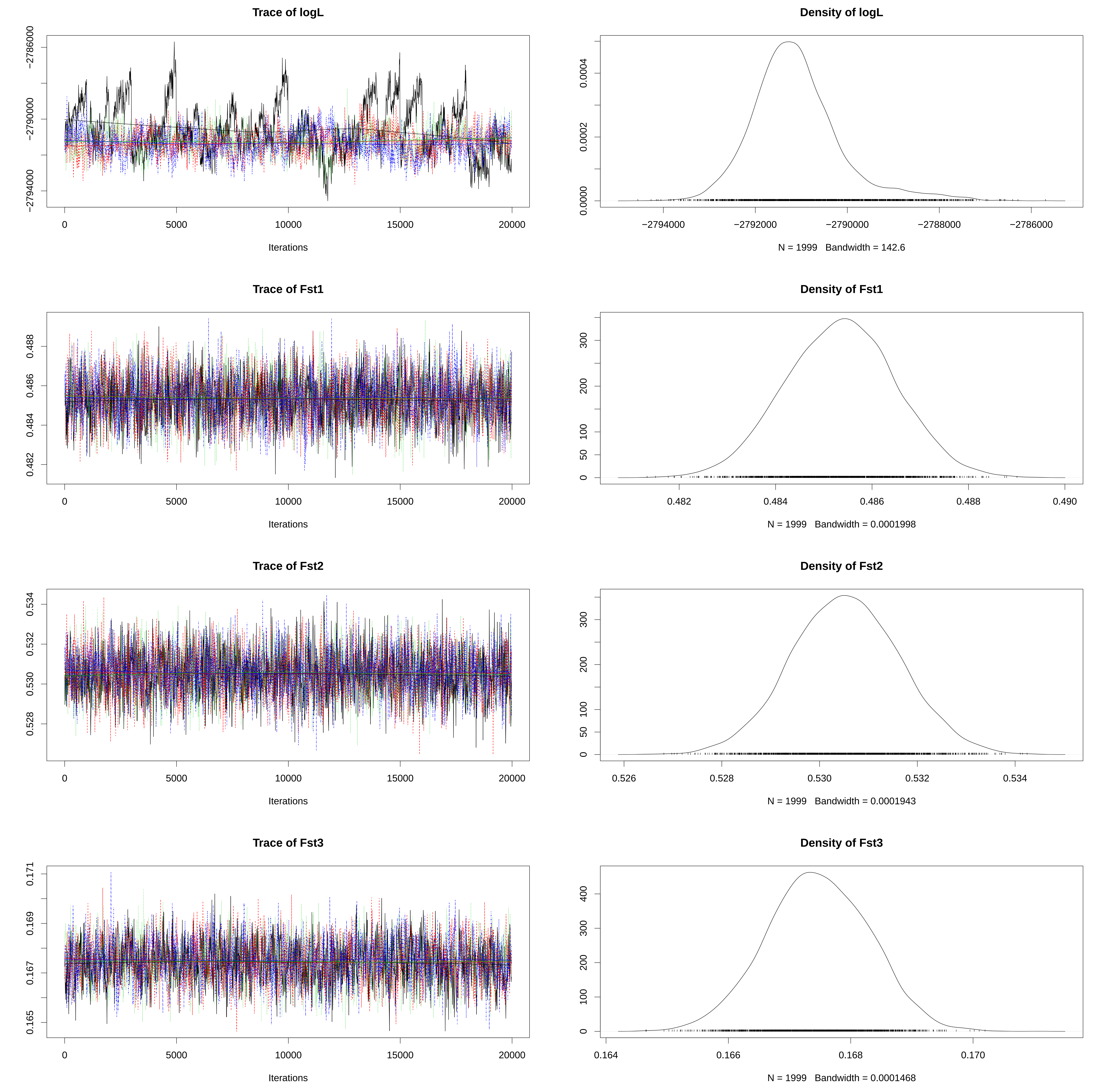

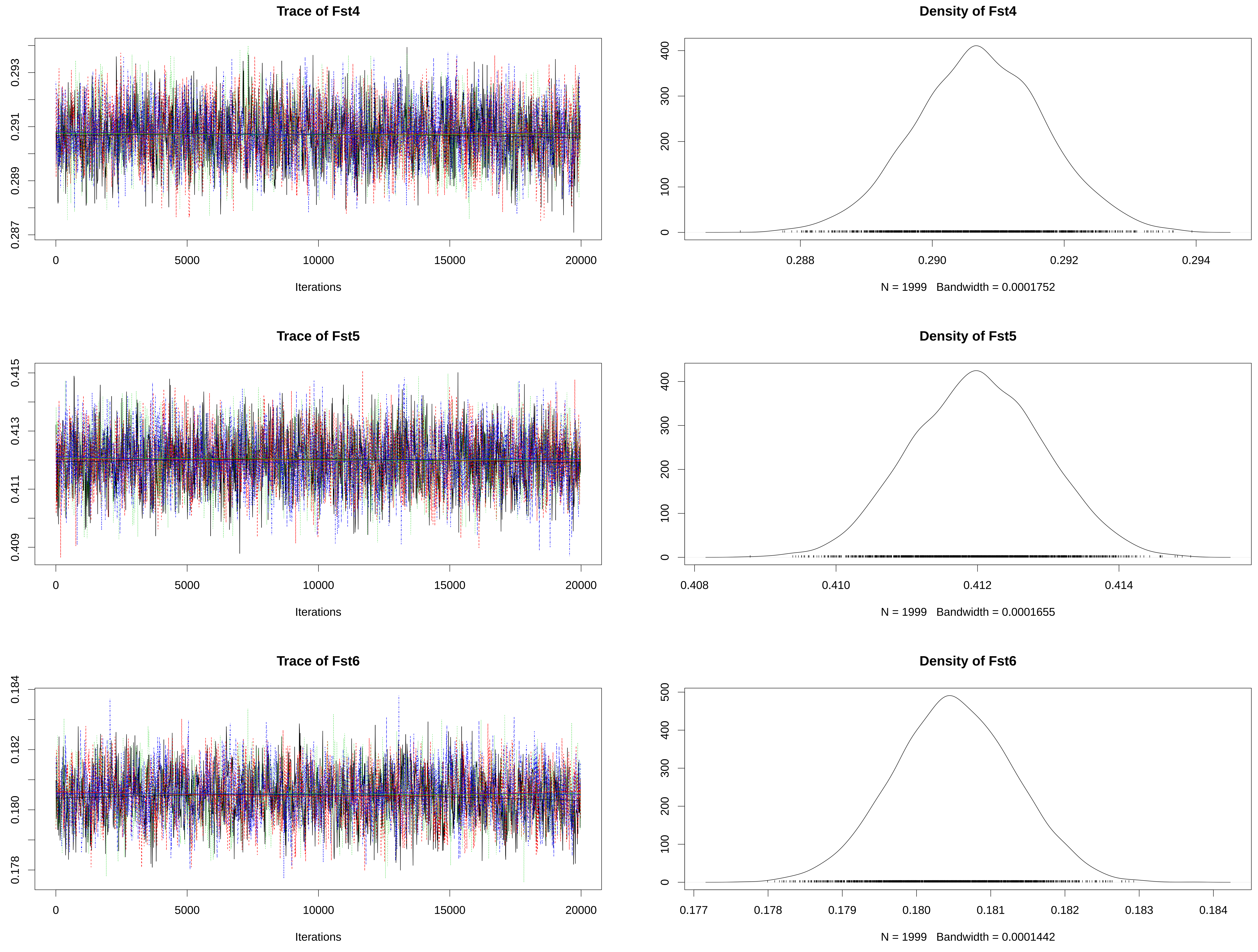

### figureS8.pdf

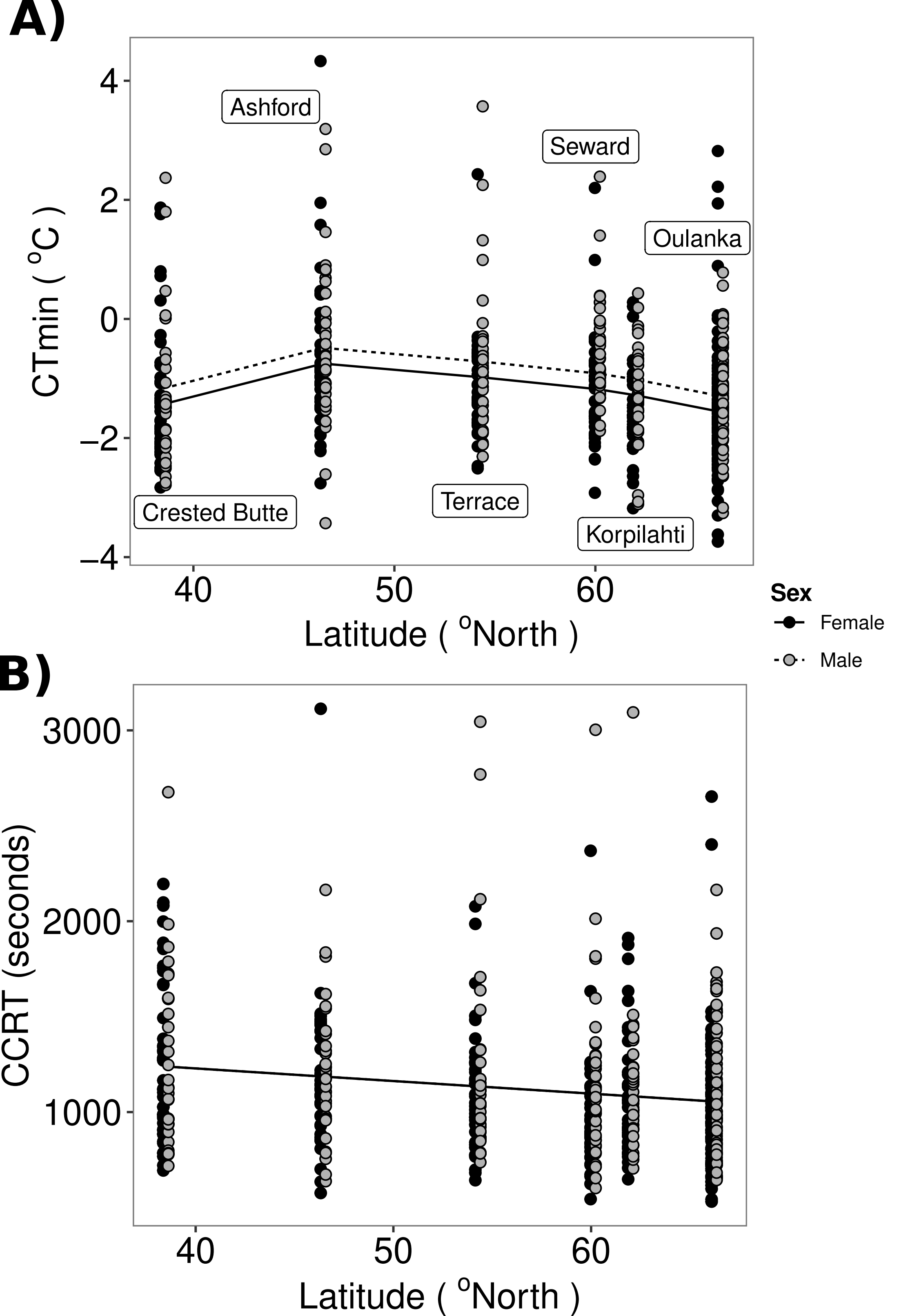

### figureS8A.pdf

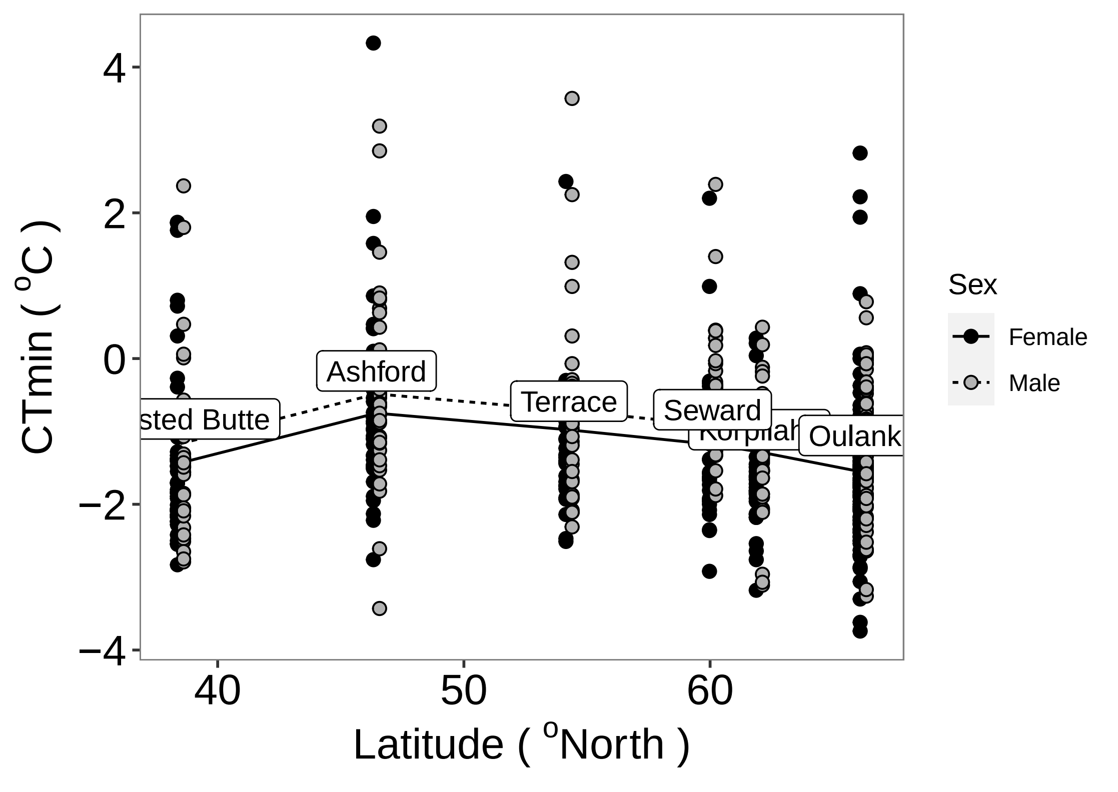

### figureS8B.pdf

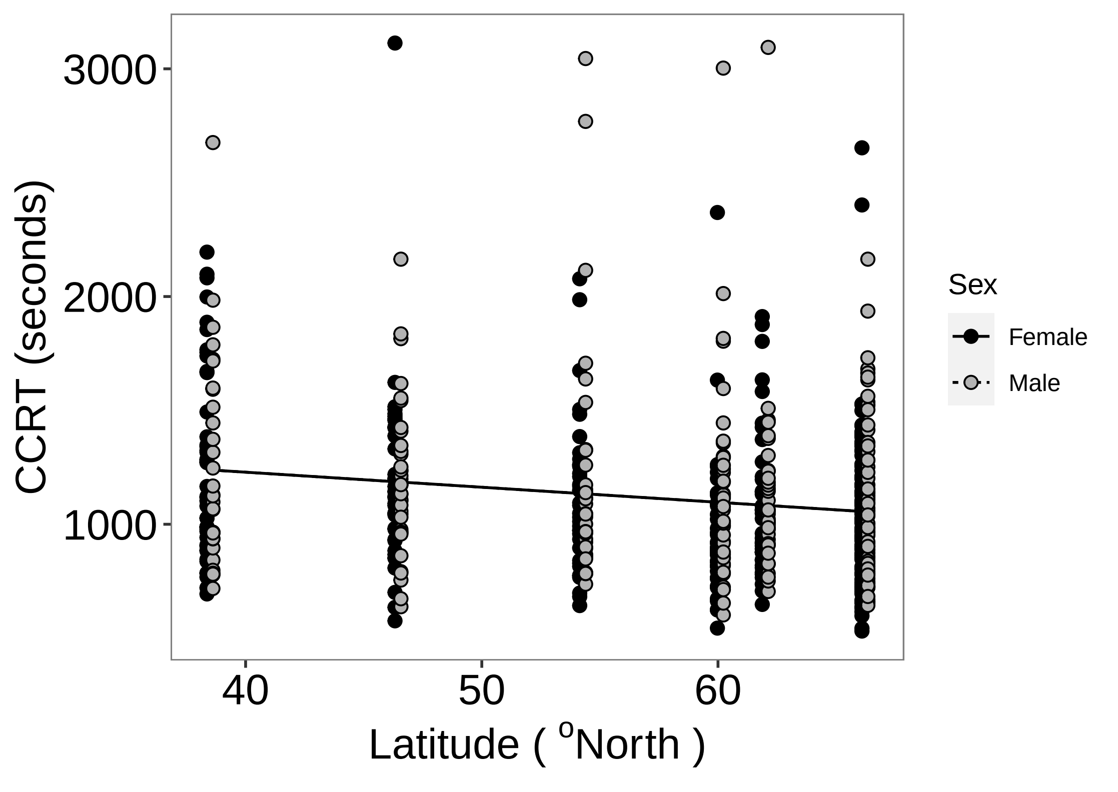

### figureS9_scree_plot.pdf

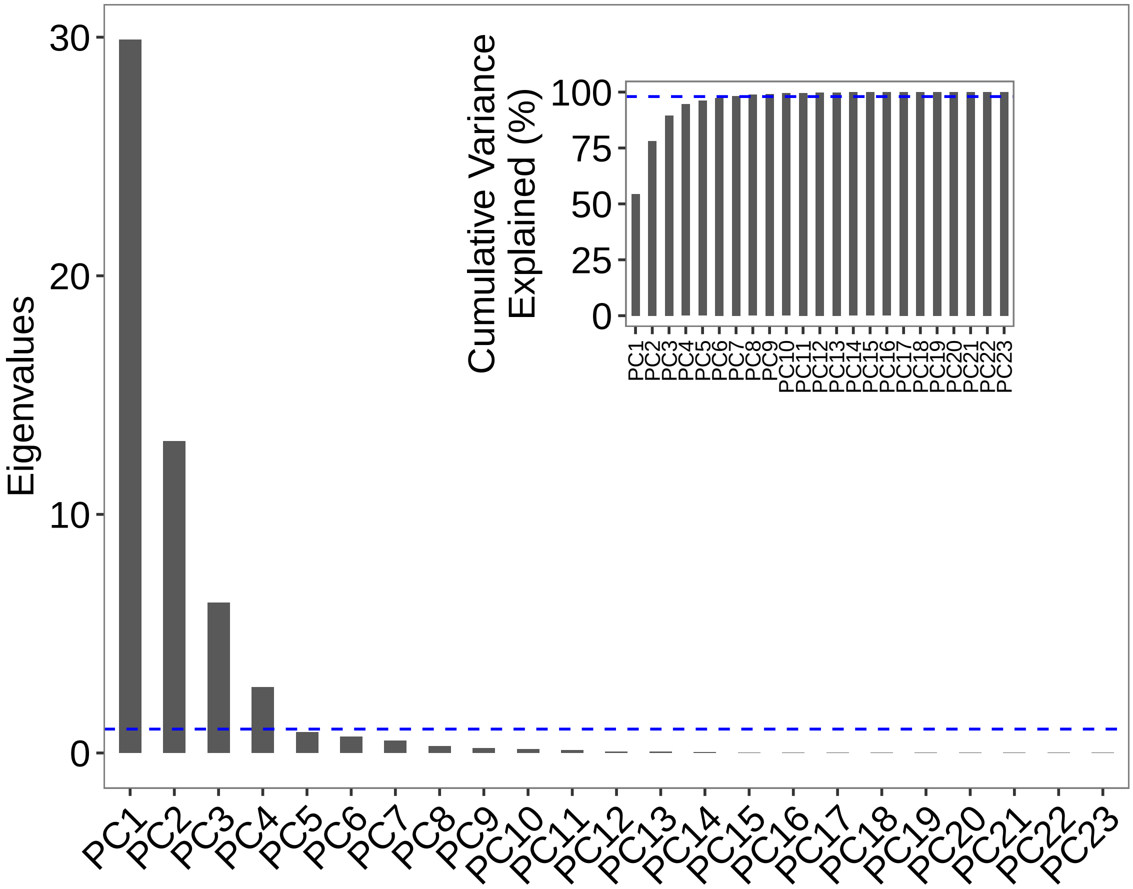

### figureS10_PC_phens.pdf

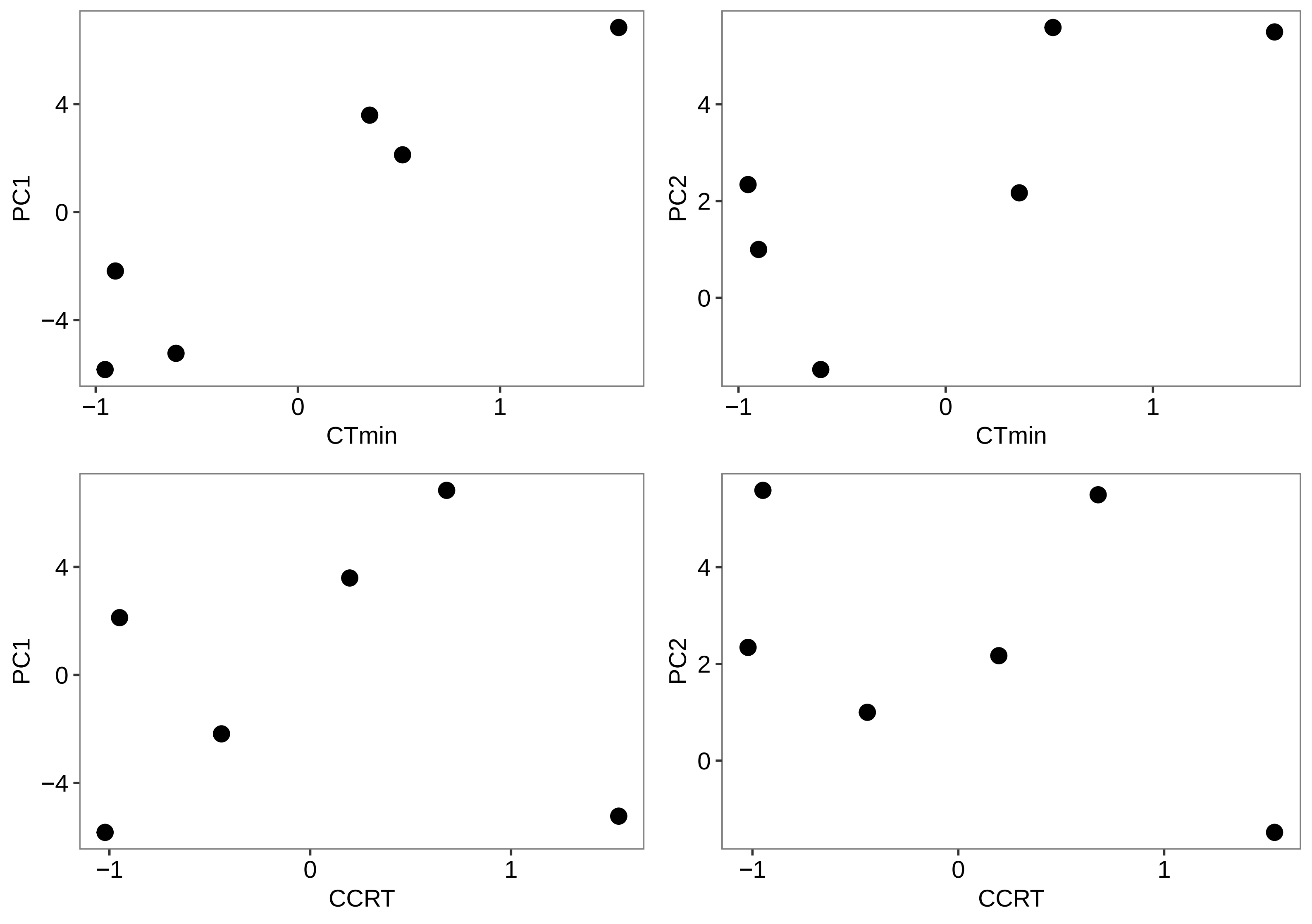

### figureS11.png

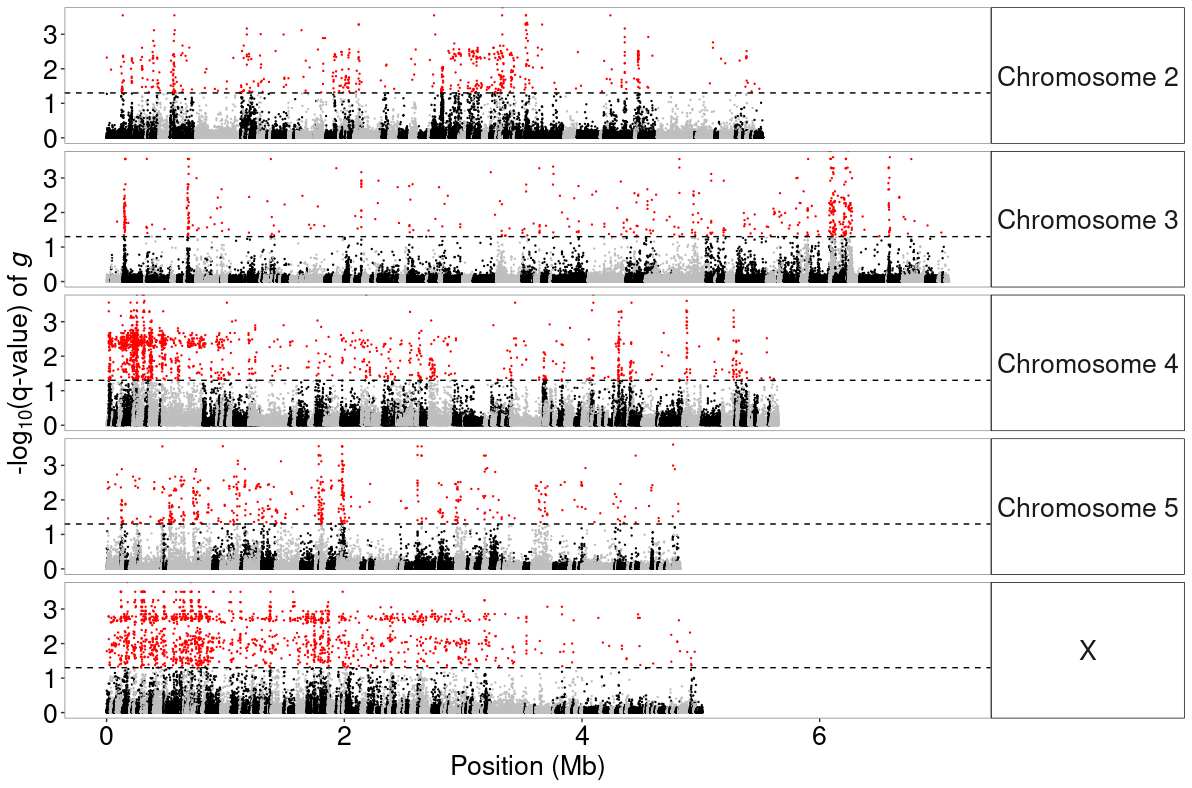

### figureS12.png

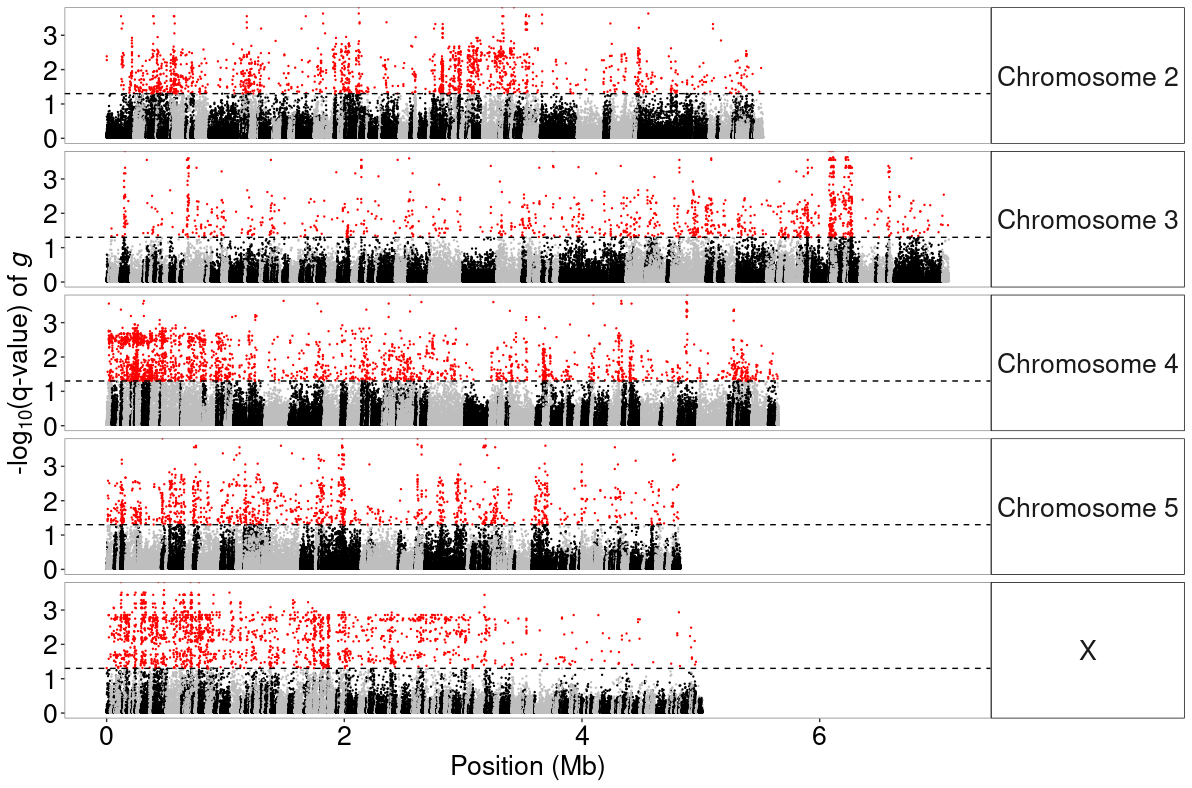

### figureS13.png

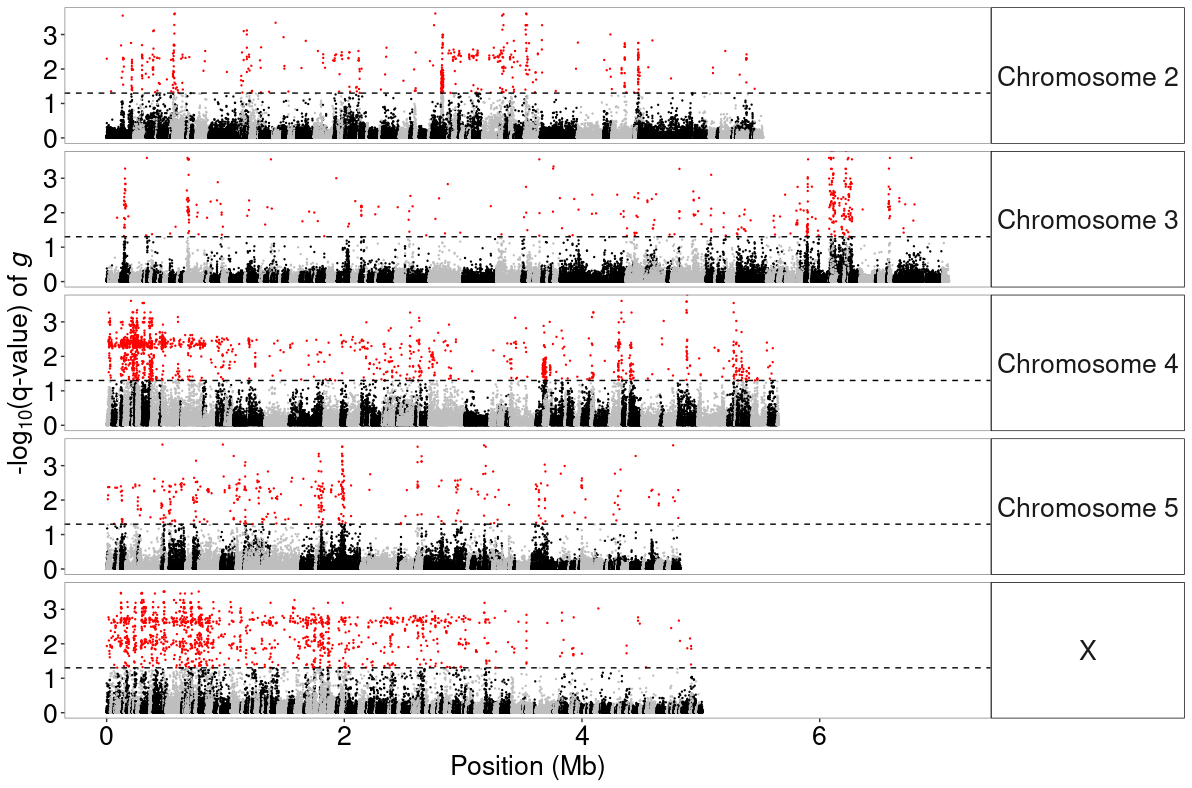

### figureS14.png

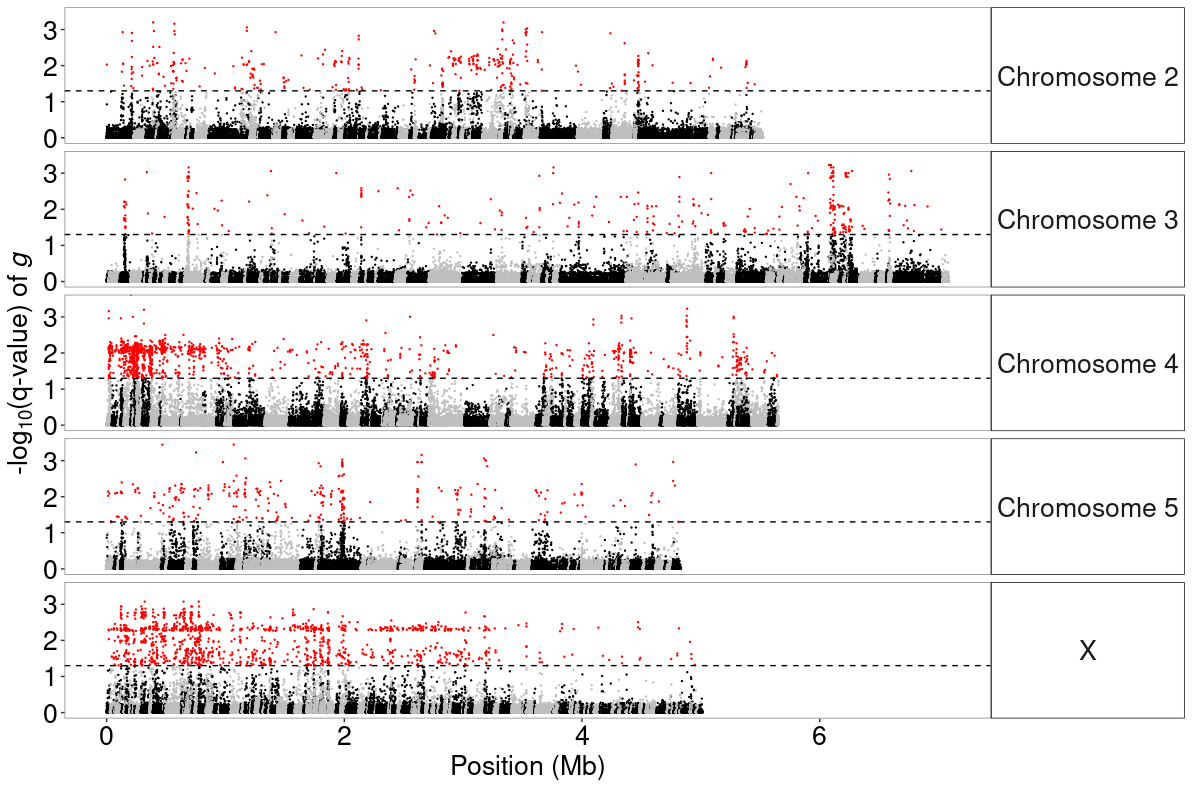

### figureS15.pdf

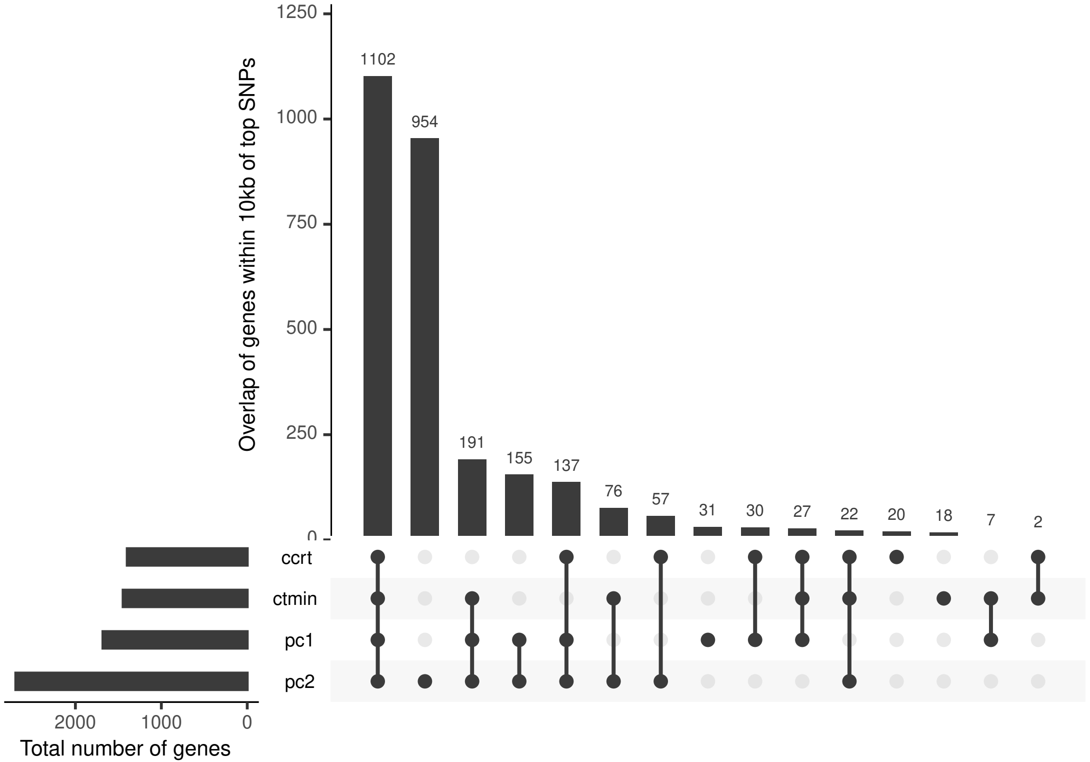

### figureS16.pdf

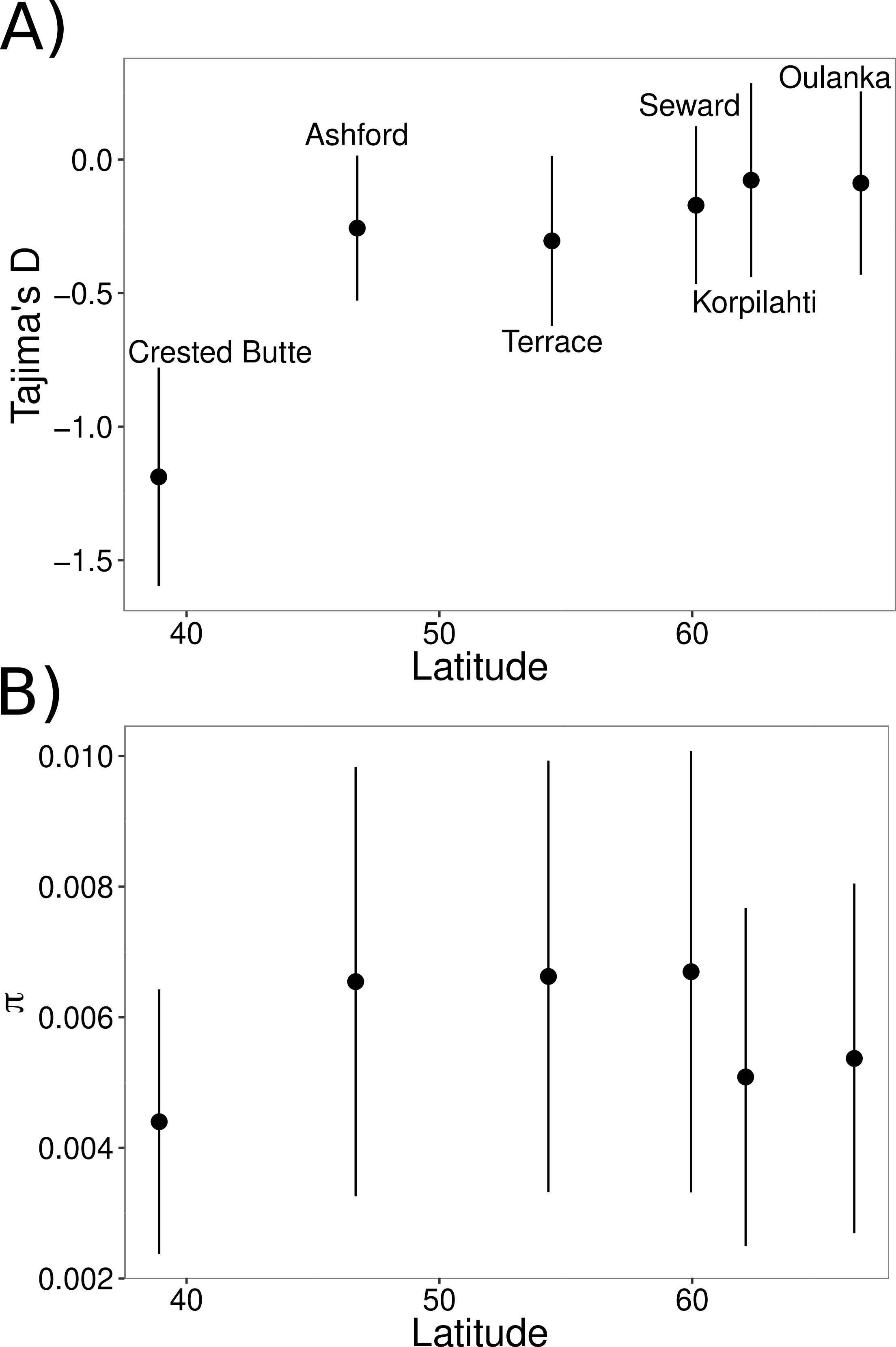

### figureS16A_tajD_by_lat.pdf

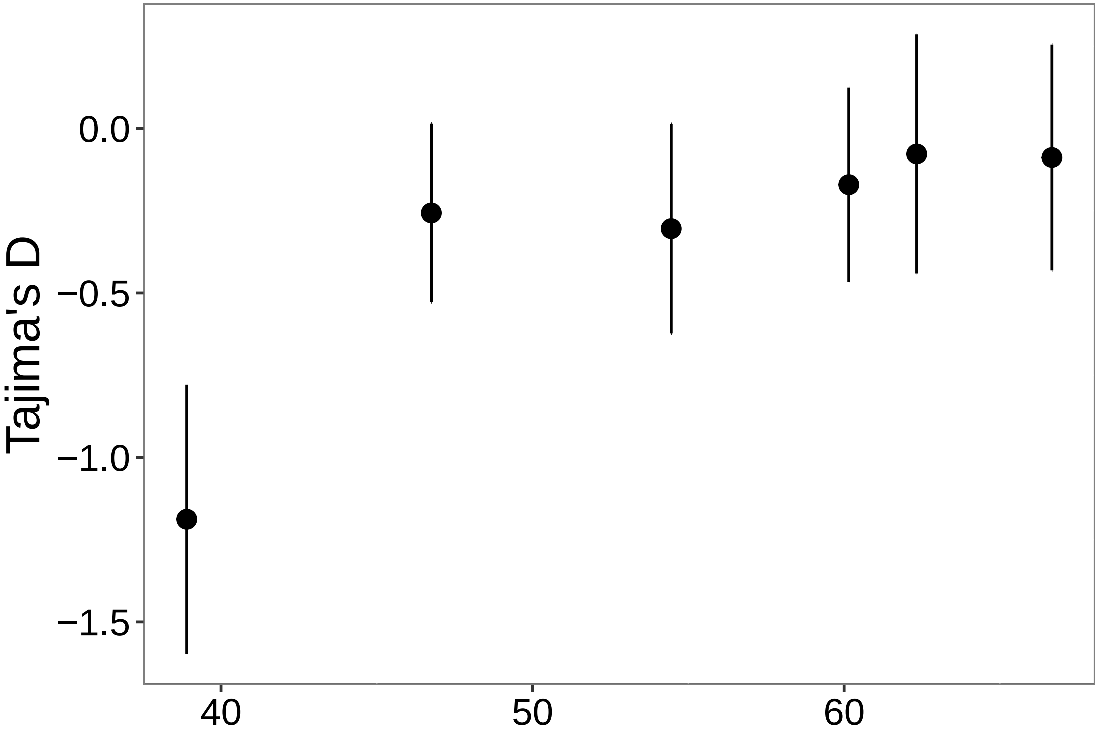

### figureS16B_pi_by_lat.pdf

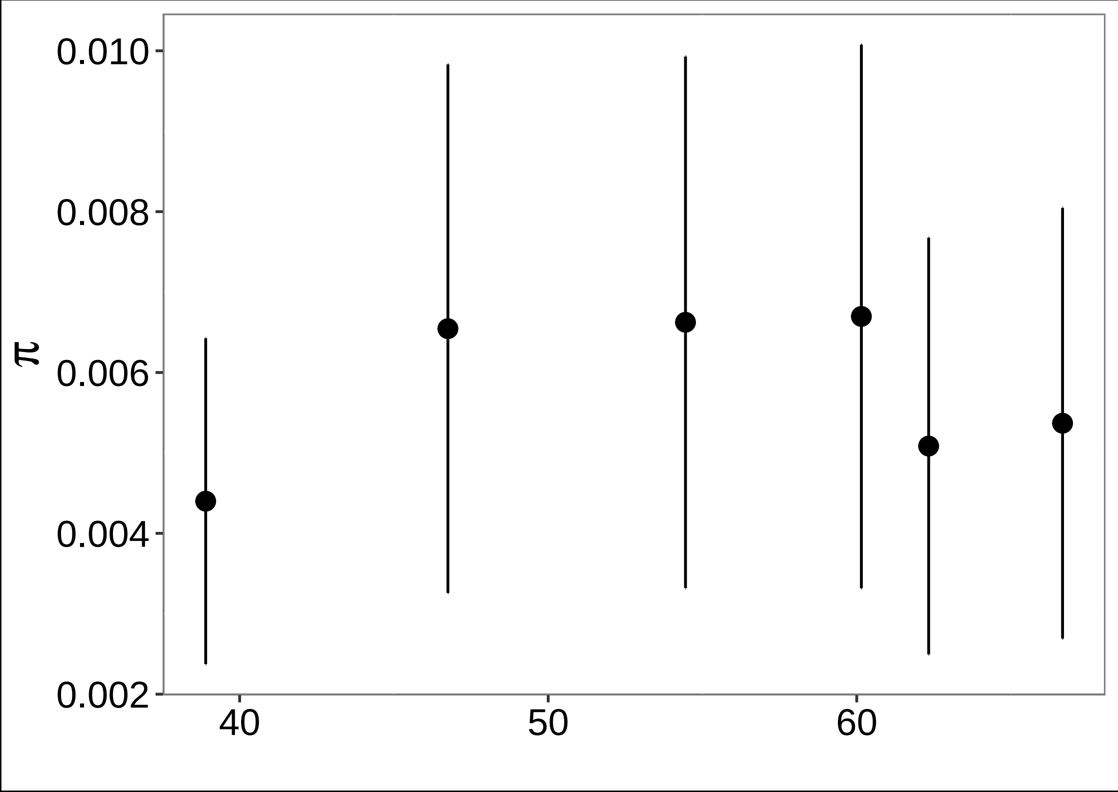
